# AI-assisted Drug Re-purposing for Human Liver Fibrosis

**DOI:** 10.1101/2025.04.29.651320

**Authors:** Yuan Guan, Jakkapong Inchai, Zhuoqing Fang, Jacky Law, Alberto Alonzo Garcia Brito, Annalisa Pawlosky, Juraj Gottweis, Alexander Daryin, Artiom Myaskovsky, Lakshmi Ramakrishnan, Anil Palepu, Kavita Kulkarni, Wei-Hung Weng, Vivek Natarajan, Alan Karthikesalingam, Keran Rong, Yunhan Xu, Tao Tu, Gary Peltz

## Abstract

Liver fibrosis is a severe disease with few treatment options due to the poor quality of the available animal and *in vitro* models. To address this, we investigated whether a hypothesis generating multi-agent AI system (AI co-scientist) could assist in re-purposing drugs for treatment of liver fibrosis and direct their experimental characterization. A multi-parameter image analysis workflow, which enabled anti-fibrotic efficacy and drug toxicity to be serially assessed in multi-lineage human hepatic organoids grown in microwells (i.e., microHOs), was used to assess the effects of 14 drugs. Remarkably, two of the three AI co-scientist-recommended drugs that targeted epigenomic modifiers exhibited significant anti-fibrotic activity. Analysis of the anti-fibrotic effects of five drugs indicated that two inhibited TGFβ-induced intracellular signaling and three drugs altered TGFβ-induced mesenchymal cell differentiation. Since all five of the anti-fibrotic drugs reduced TGFβ-induced chromatin structural changes, epigenomic changes play an important role in the pathogenesis of liver fibrosis. One AI co-scientist recommended drug is an FDA-approved anti-cancer treatment (Vorinostat) that reduced TGFβ-induced chromatin structural changes by 91% and promoted liver parenchymal cell regeneration in microHOs. Hence, the use of AI co-scientist and this microHO platform identified a potential new generation of liver fibrosis treatments that also promote liver regeneration.

---

Liver fibrosis is caused by extracellular matrix (**ECM**) accumulation in response to chronic liver injury, which is often caused by viral infection, non-alcoholic steatohepatitis, or chronic alcohol exposure (*1–3*). This fibrotic state results from an interaction between parenchymal and nonparenchymal liver cells and possibly involves infiltrating immune cells (*4–6*). The key non-parenchymal cells are the myofibroblasts (**MyoF**), which are generated in response to fibrogenic stimuli (especially TGFβ1) and they produce excess fibril-forming collagens and other ECM proteins (*6–11*). Irrespective of the inciting cause, activated MyoF and the ECM proteins they produce drive all forms of liver fibrosis. Patient outcome is determined by the extent of liver fibrosis (*12*), which is a major global cause of death (∼1M per year) (*13*). Although drugs are being studied (*1, 14, 15*), many agents that generated promising results in preclinical studies have failed in clinical trials due to safety concerns or lack of efficacy. The poor quality of available model systems has been the major barrier to identifying anti-fibrotic therapies. Prior *in vitro* models did not have the spectrum of cell types that mediate fibrogenesis nor could they reproduce its key feature (i.e., production of thick collagen filaments (*16*)). Conclusions drawn from animal models are limited by concerns about species-specific differences in fibrotic mechanisms (*17*). Hence, there are limited treatment options (other than when the underlying cause can be treated) for this prevalent and severe disease (*18*).

Here, we use a pioneering approach to identify and characterize new anti-fibrotic agents. First, we investigate whether a hypothesis-generating multi-agent AI system built with Gemini 2.0 (AI co-scientist (*19*)) could identify anti-fibrotic agents that target epigenomic modifiers, and if it could direct their experimental characterization. AI co-scientist (see supplemental note 1) utilizes a set of specialized agents for hypothesis generation and experimental planning using a ‘scientist in the loop paradigm. Second, to examine the anti-fibrotic efficacy of drugs suggested by AI co-scientist we use an enhanced live cell imaging system that enables anti-fibrotic efficacy and drug toxicity to be simultaneously assessed in multi-lineage human hepatic organoids (*20*) grown in microwells (i.e., ‘***microHOs***’) (*21, 22*). microHOs have hepatocytes, cholangiocytes, bile ducts, liver lobule architecture; and thick collagen filaments and MyoF appear in microHOs after TGFβ exposure. microHOs were used to perform the experiments suggested by AI co-scientist to investigate the mechanism of action of five anti-fibrotic agents and to characterize their effects on MyoF, mesenchymal cells and parenchymal cells.

## Results

*The microHO platform for discovery and characterization of anti-fibrotic drugs.* Our previously described microHO system (*21*) and a high content imaging system were used to serially measure the antifibrotic effect of a drug. The collagen-producing cells in microHOs are labeled with a fluorescent intracellular protein, which enables anti-fibrotic efficacy to be serially analyzed. Drugs were added at the same time as a pro-fibrotic stimulus (TGFβ), and the extent of fibrosis developing while the cells differentiate to form a multi-lineage microHO was measured. Addition of a nuclear stain (Hoechst 33342) enabled drug toxicity and antifibrotic efficacy to be simultaneously assessed. Samples for flow cytometry (FCM) and single-cell RNA sequence (scRNA-seq) analysis were collected at the endpoint of each experiment to characterize drug effects on specific cell types (**Fig. 1A**). For example, consistent with our prior finding (*21*), TGFβ receptor (TGFBR1, SB431542), GSK3β (CHIR) and p38 (SB202190) inhibitors blocked TGFβ-induced fibrosis but did not cause significant cellular toxicity in microHOs (**Fig. S1A-B, Tables S1-S2**). A PPARγ inhibitor had a small effect on TGFβ-induced fibrosis in microHOs (**Fig. S1C**). Interestingly, FCM results revealed that the TGFBR1, p38, and GSK3β inhibitors also increased the number of liver parenchymal (EPCAM^+^) cells in TGFβ-treated microHOs (**Fig. S1D, Table S3**). Trichrome staining confirmed that these drugs decreased the amount of ECM and increased the number of liver parenchymal cells in microHOs (**Fig. S1E**). Thus, TGFBR1, p38, or GSK3β inhibitors reduced fibrosis and promoted liver parenchymal cell regeneration in the presence of a pro-fibrotic stimulus.

**Figure 1.**
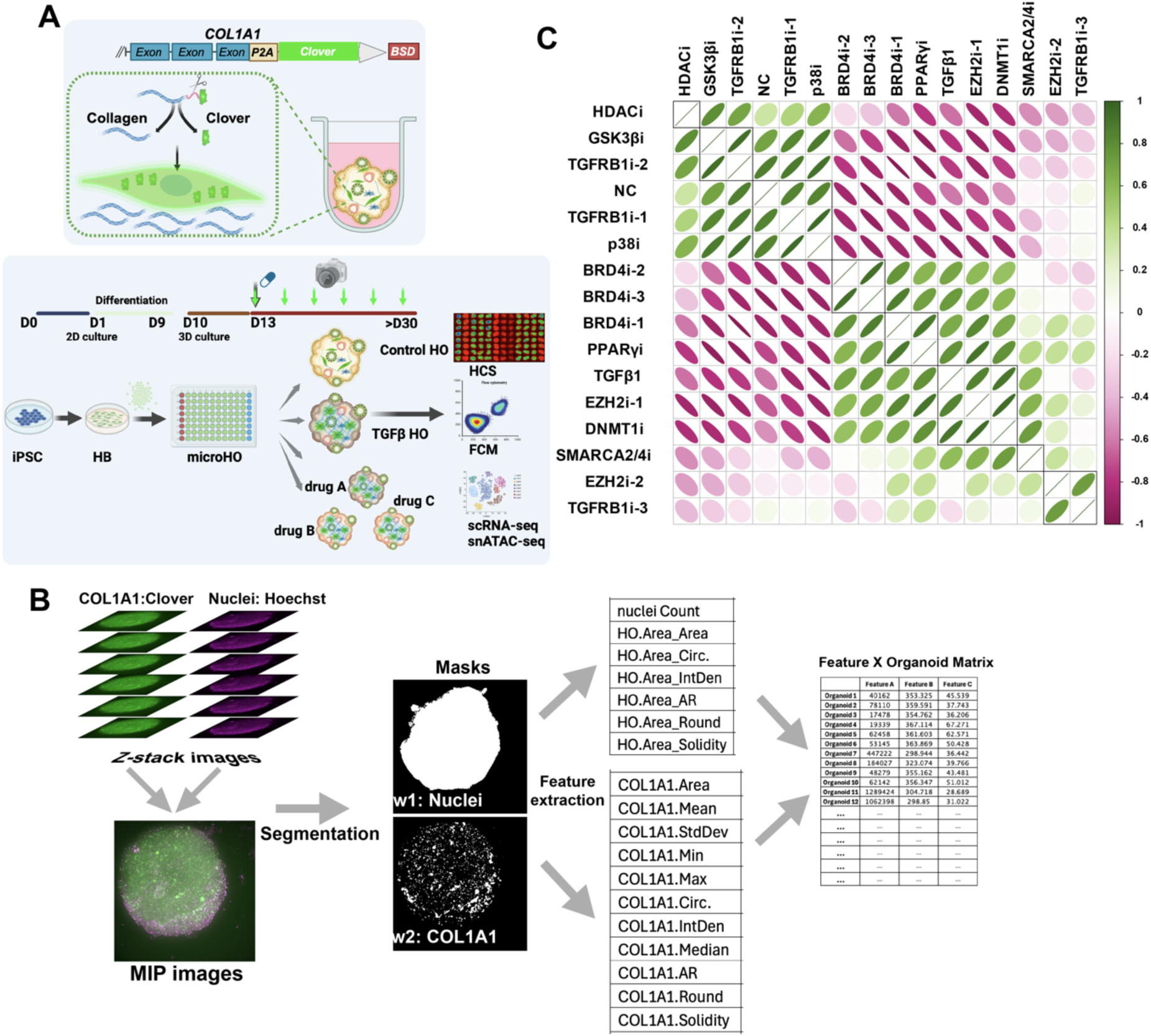
The microHO platform for assessing hepatic fibrosis. (**A**) *Top:* The iPSC line has a P2A-Clover insert at the 3’ end of *COL1A1.* The self-cleaving P2A peptide enables *COL1A1* expressing cells in a microHO to be labelled with a fluorescent intracellular protein (Clover). *Bottom*: A diagram of the microHO platform. Engineered iPSCs are differentiated into hepatoblasts (HB), and 10,000 HBs are placed in each microwell on day 10. TGFβ1 ± drug are added on day 13, and the cultures are further differentiated into hepatic organoids. High content imaging is used to serially measure the fluorescence and cell number in microHOs. At the end of each experiment, samples for flow cytometry (FCM), scRNA-seq and snATAC-seq analyses are collected. (**B**). A diagram of the supervised image workflow for analyzing microHOs. (i) Twenty to forty Z-stacked images per microHO were prepared from the two channels used to analyze the extent of fibrosis (COL1A1:Clover) and cell number (number of nuclei from the Hoechst 33342 stain) for each microHO. (ii) Masks were prepared by region of interest (ROI) segmentation from each channel from the maximum intensity projection (MIP) image. (iii) Then, 18 features from the two channels were extracted to build the ‘Feature X organoid’ matrix. (**C**) A heat map is generated from analysis of the *Pearson’s* correlation coefficients calculated from 18 features measured in normal control (NC) or TGFβ-treated microHOs. The results are compared with those obtained from microHOs treated with TGFβ and 14 other drugs. As shown here, microHOs treated with TGFRB, GSK3β, p38 or HDAC (SAHA) inhibitors have the most similarity with NC microHOs. The TGFβ-treated microHOs are very different from NC microHOs; and microHOs treated with EZH2, DNMT1, or PPARγ inhibitors have the least similarity with NC microHOs. The BRD4 inhibitor (Zen-3964, I-BET762, JQ-1) effects cluster together, and are intermediate between NC and TGFβ-treated microHOs. The bar on the right shows the extent of the correlation (by color) for each square in the diagram: green (or magenta) indicates a positive (or negative) correlation.

A supervised image analysis workflow used the data obtained from the cell number and fibrosis channels to generate an 18 component ‘feature X organoid’ matrix that enhanced our ability to interpret the fluorescence and cell number signals obtained from the twenty to forty Z-stacked images collected from each microHO per time point (**Figs. 1B-C, S2**). Pearson’s correlation coefficients calculated for each feature were used to generate a heat map that enabled the anti-fibrotic efficacy and cellular toxicity of 14 analyzed drugs (Table S1) to be compared with each other, and with normal control (NC) and TGFβ-treated microHOs. For example, the TGFBR1-, GSK3β- or p38 inhibitor-treated microHOs were more like NC microHOs, while PPARγ-treated microHOs were more like TGFβ-treated microHOs (**Fig. 1C**).

*Anti-fibrotic drugs targeting epigenomic pathways*. When AI co-scientist was prompted to generate experimentally testable hypotheses about the role of epigenomic changes in liver fibrosis, it hypothesized that: (i) “*histone modifications, particularly deacetylation, in the promoter regions of genes responsible for MyoF differentiation”; (ii) dynamic changes in DNA methylation patterns around genes involved in the MyoF pathway could be crucial; and (iii) “pharmacological inhibition should be used to test the importance of epigenomic alterations.”* It suggested testing the inhibitors targeting 3 epigenomic modifiers: histone deacetylases (HDACs), which alter gene expression by removing the acetyl moieties from histones to create a more compact chromatin (*23*); DNA Methyltransferase 1 (DNMT1), which produces heritable changes in DNA methylation patterns (*24*); and bromodomain 4 (BRD4), which promotes transcriptional elongation by binding to acetylated lysine residues on histones to promote cell cycle progression and organ development (*25, 26*) (**Supplemental Data File 1**). For comparison, based upon a literature review, one author (GP) selected two other epigenomic targets: EZH2 in the Polycomb Repressive Complex 2 (PRC2) because EZH2 inhibitors reduced liver fibrosis in two murine models (*27*); and SMARCA2/4 in the Switch/Sucrose-Nonfermentable (SWI/SNF) complex because it interacts with transcription factors activated in fibrotic liver (*28*),(*29, 30*) and inhibits a (Hippo) pathway that regulates hepatic stellate cell activation (*31*). However, SMARCA2/4 (PFI-3 (*32*)) and EZH2 (GSK126 (*33, 34*)) inhibitors did not reduce fibrosis; (**Fig. S3**); and microHOs treated with these drugs resembled TGFβ-treated microHOs (Fig. 1).

The drugs used to test the AI co-scientist selected epigenomic targets were: a selective DNA methyl transferase 1 (DNMT1) inhibitor (GSK3685032) that is less toxic and has more durable hypomethylating effects than prior DNMT inhibitors (*35*); an FDA-approved anti-cancer drug that is a pan-histone deacetylase (HDAC) inhibitor (Vorinostat or SAHA) (*36, 37*); and three BRD4 inhibitors: JQ1 (an early BRD4 inhibitor) (*38*), and two (I-BET762 (GSK525762) (*39*), ZEN-3694 (*40, 41*)) with improved risk-benefit profiles. The DNMT1 inhibitor did not reduce fibrosis in microHOs; its cytotoxicity would preclude further clinical development (Fig. S3); and the heatmap shows that GSK3685032-treated and NC microHOs were very different (Fig. 1C). In contrast, fibrosis was blocked by all 3 BRD4 inhibitors and by the HDAC inhibitor at concentrations that did not induce cellular toxicity (**Figs. 2A-C, S4**). The heatmap shows that the effects of the three BRD4 inhibitors clustered together, whereas Vorinostat-treated microHOs were most like NC microHOs (Fig. 1C). FCM results confirmed the anti-fibrotic activity of the HDAC and BRD4 inhibitors (HDACi, BRD4i). Vorinostat significantly increased the parenchymal cell (EPCAM^+^) population in microHOs, while the BRD4i induced an increase in the epithelial population, which did not reach statistical significance (**Fig. 2D**). Trichrome staining confirmed that the HDACi and BRD4i decreased the amount of ECM (p<0.001) and increased the number of epithelial cells in microHOs (p<0.001) (**Fig. 2E**).

**Figure 2.**
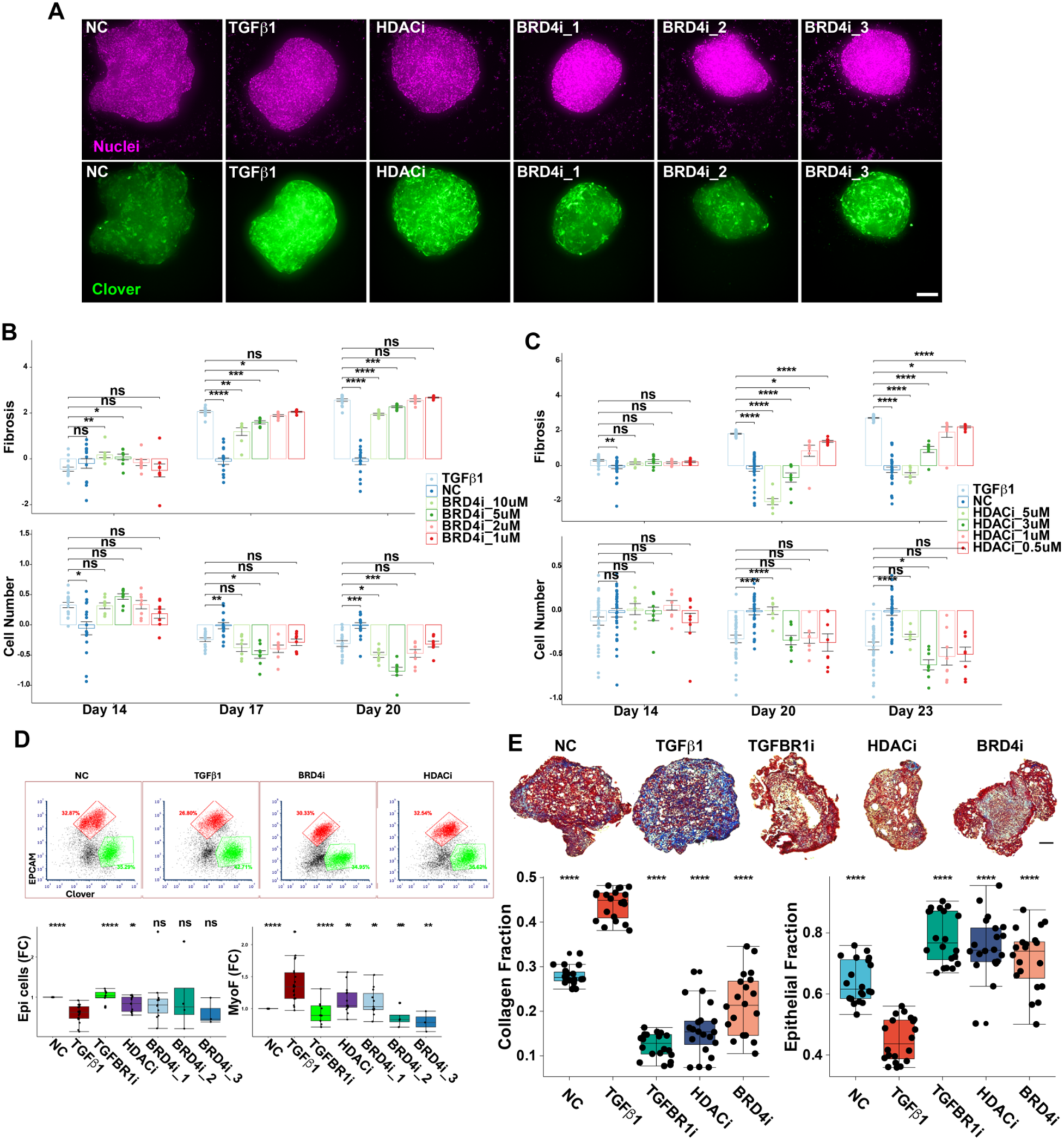
TGFβ-induced fibrosis in microHOs is reduced by BRD4 or HDAC inhibitors. (**A**) Representative MIP images generated from microHOs treated with normal media (NC), 50 ng/mL TGFβ1, or TGFβ1 and the indicated drug. Scale bar, 100 μm. Clover expression is green, and the nuclei stained with Hoechst 33342 are magenta. (B, C) microHOs were treated with either normal media (NC), 50 ng/mL TGFβ, or TGFβ with the indicated concentration of BRD4 (ZEN-3694) (**B**) or HDAC (SAHA) (**C**) inhibitor on day 13. The extent of fibrosis (COL1A1 fluorescence) and cell numbers were serially measured on days 14 through 23. Sixteen microHOs per treatment were assessed per condition. The fibrosis and cell number measurements were normalized relative to NC microHOs. **(D)** FCM results show that HDAC (SAHA) and BRD4 inhibitors decreased MyoFs in microHOs, but only SAHA increased the epithelial cell population. *Top*: Representative FCM results obtained from microHOs receiving the indicated treatment. The x-axis shows the COL1A1 Clover fluorescence, and the y-axis shows the cells stained with an anti-EPCAM antibody. *Bottom*: The change in epithelial cells and MyoFs in microHOs caused by the indicated treatments. Each measurement is made on at least 10 individually prepared batches of microHOs. To calculate the fold change (FC), the percentage of cells in each batch was normalized relative to that in NC microHOs. SB431542 is a TGFBR1 inhibitor; Vorinostat (SAHA) is the HDAC inhibitor; and ZEN-3694, IBET762 and JQ-1 are the BRD4 inhibitors 1 to 3. (**E**) Top panel: Images of Trichrome-stained TGFβ1-treated microHOs show a marked increase in collagen-rich connective tissue (blue-stained regions) relative to control (NC) microHOs, which only had a thin layer of connective tissue. The TGFβ1-induced increased in collagen was markedly inhibited by co-addition of TGFBR1, HDAC, or BRD4 inhibitors. Moreover, the number of epithelial cells (dark red regions) was decreased by TGFβ1, and co-addition of TGFBR1, HDAC, or BRD4 inhibitors prevented the decrease in epithelial cells. Scale bar: 100 μm. *Bottom panel*: Box plots show the area within microHOs that received the indicated treatment (n > 10 per group) occupied by collagen (collagen fraction) or parenchymal cells (epithelial fraction). ns, not significant; *, p-value<0.05; **, p-value<0.001; ***, p-value<0.001; and ****p-value<0.0001.

*Molecular analysis of drug effects*. Since AI co-scientist suggested that *we use “single cell RNA sequencing (scRNA-seq) to assess global transcriptional changes associated with fibrosis and drug effects,”* scRNA-seq data was generated from 613,826 cells obtained from NC (n=320 microHOs), TGFβ- (n=320), and TGFβ plus drug treated (n=32 per drug) microHOs. UMAP plots show that the cellular transcriptomes in NC and TGFβ-treated microHOs were reproducibly different. The sixteen cell clusters identified in microHOs were annotated based upon transcriptomic comparisons with large reference datasets and by canonical marker mRNA expression (**Figs. 3A-B, S5-6; and Supplemental Data File 2**). The scRNA-seq data was examined to investigate how five drugs exerted their anti-fibrotic effects. Transcriptomic comparisons and cell percentage analyses revealed that the TGFBRi and p38i-treated microHOs resembled NC microHOs, which is consistent with their targeted pathways (**Figs. 3B-D**). TGFBR1i block TGFβ-induced intracellular signaling, which also activates the pro-fibrotic p38 Mitogen Activated Protein Kinase (MAPK) signaling pathway (*42–44*), which we previously found was activated in MyoF in microHOs and in cirrhotic human liver (*21*). The p38i effect confirms that p38 pathway activation is essential for TGFβ-induced liver fibrosis.

**Figure 3.**
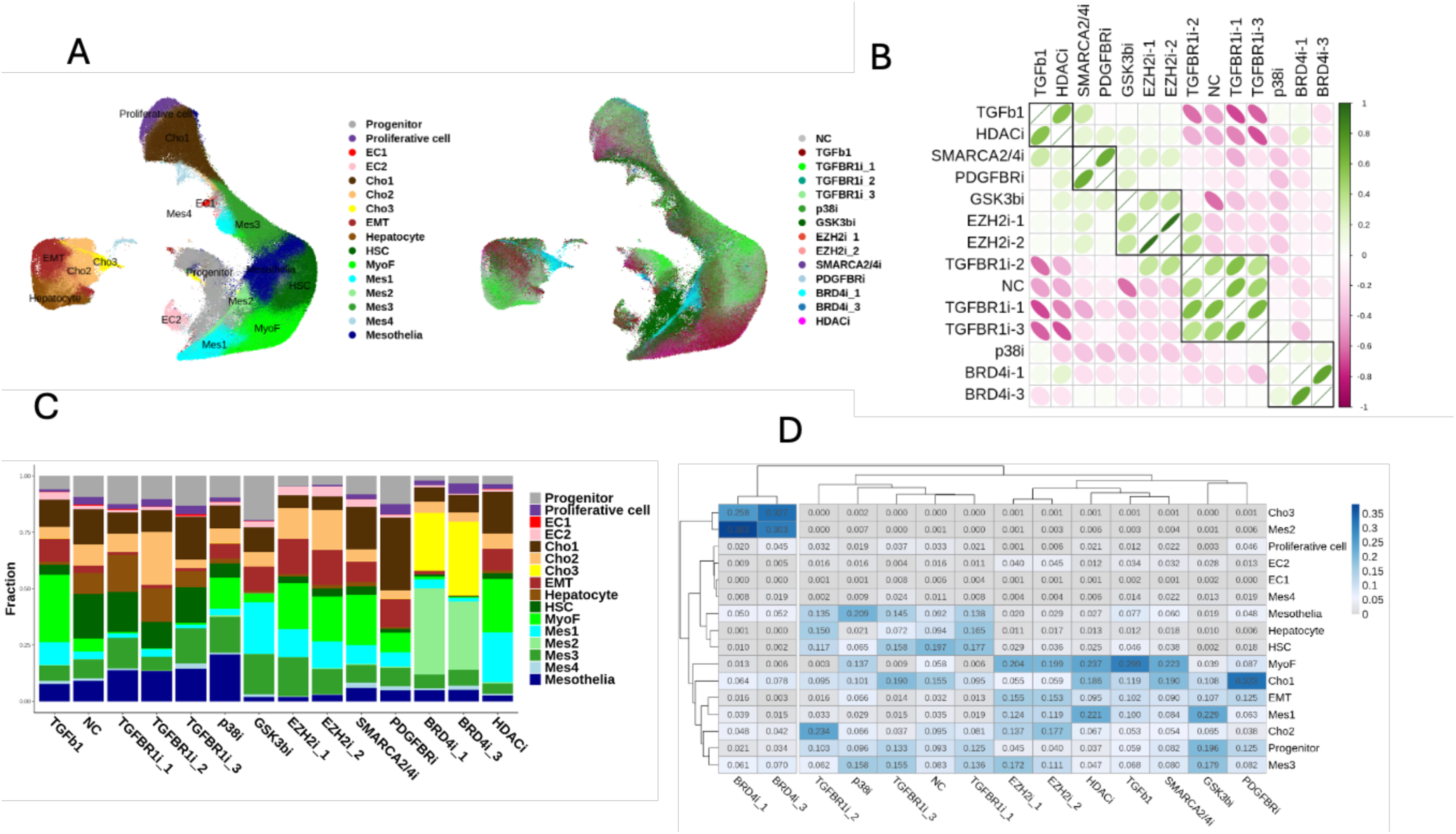
TGFβ1 and drug-induced cellular and transcriptomic changes in microHOs. (**A**) Combined sample UMAP plots are labeled with the 16 cell clusters identified in microHOs (Left) or by the microHOs receiving 14 different treatments (Right). scRNA-seq data was generated from 613,826 cells obtained from microHOs receiving the indicated 14 treatments. (**B**) A heat map showing the relationships between the transcriptomes of normal control (NC), TGFβ-, and TGFβ with plus drug-treated microHOs. The Pearson correlation coefficient for each transcriptomic comparison was calculated using the scRNA-seq data described in (A). The bar on the right shows the extent of the correlation (by color) shown in each square of the diagram (green is positive, magenta is negative). As can be seen, the transcriptomes of the three TGFBR inhibitor-treated and the GSK3β inhibitor-treated microHOs are most like NC microHOs, whereas the transcriptomes of HDAC and SMARCA2/4 inhibitor treated microHOs is like TGFβ-treated microHOs. (**C**) scRNA-seq data obtained from NC, TGFβ-, and TGFβ and drug treated microHOs were analyzed to identify the percentage of cells in each of the 16 cell clusters in each type of microHO. The percentage of each indicated cell type represents the average obtained from multiple experimental batches: 320 NC, 320 TGFβ-treated microHOs, and from 32 TGFβ and drug treated microHOs were evaluated for each drug treatment. (**D**) A heatmap shows the hierarchical clustering of the percentage of each of the 16 cell clusters in NC, TGFβ1 or TGFβ1 plus drug-treated microHOs. The cell type changes in microHOs treated with any of the 3 TGFBR inhibitors or with the GSK3β or p38 inhibitors are most like NC microHOs; while the cell type changes in HDAC, SMARCA2/4, or EZH2 inhibitor-treated microHOs are most like TGFβ1-treated microHOs.

The effects of TGFβ and the GSK3β, p38 and TGFBR inhibitors on the seven mesenchymal clusters were of interest (**Fig. 4A-D**). Transcriptomic comparisons with fetal, adult normal and diseased human liver samples identified clusters resembling MyoF and hepatic stellate cells (HSC) in human liver; along with clusters resembling four mesenchymal cell types (Mes1-4) and mesothelial cells. Since the Mes4 and mesothelial clusters were a minor proportion of the cells and were not changed in response to the various treatments, our analysis focused on five mesenchymal clusters. Among the differentially expressed genes (DEG) identified in the HSC (n=757) and MyoF (n=862) clusters, 449 DEGs were shared between them (Fig. 4C). The DEGs were most highly correlated with ECM and collagen trimer formation, and with the TGFβ response (**Fig. S7**). TGFβ treatment decreased the HSC and increased the MyoF, Mes1 and Mes2 clusters relative to NC microHOs (Fig. 4B). While all 3 drugs prevented the TGFβ-induced increase in MyoF, only the TGFBR1 inhibitor prevented the decrease in HSC and the increase in Mes2 cells. The TGFβ-induced increase in Mes1 was suppressed by the TGFBR1i and p38i: but not by the GSK3β inhibitor, which induced an increase in Mes1. Interestingly, all three drugs increased mesenchymal progenitor-like cells (Mes3) that expresses ribosome biogenesis-related genes (Fig. S7).

**Figure 4.**
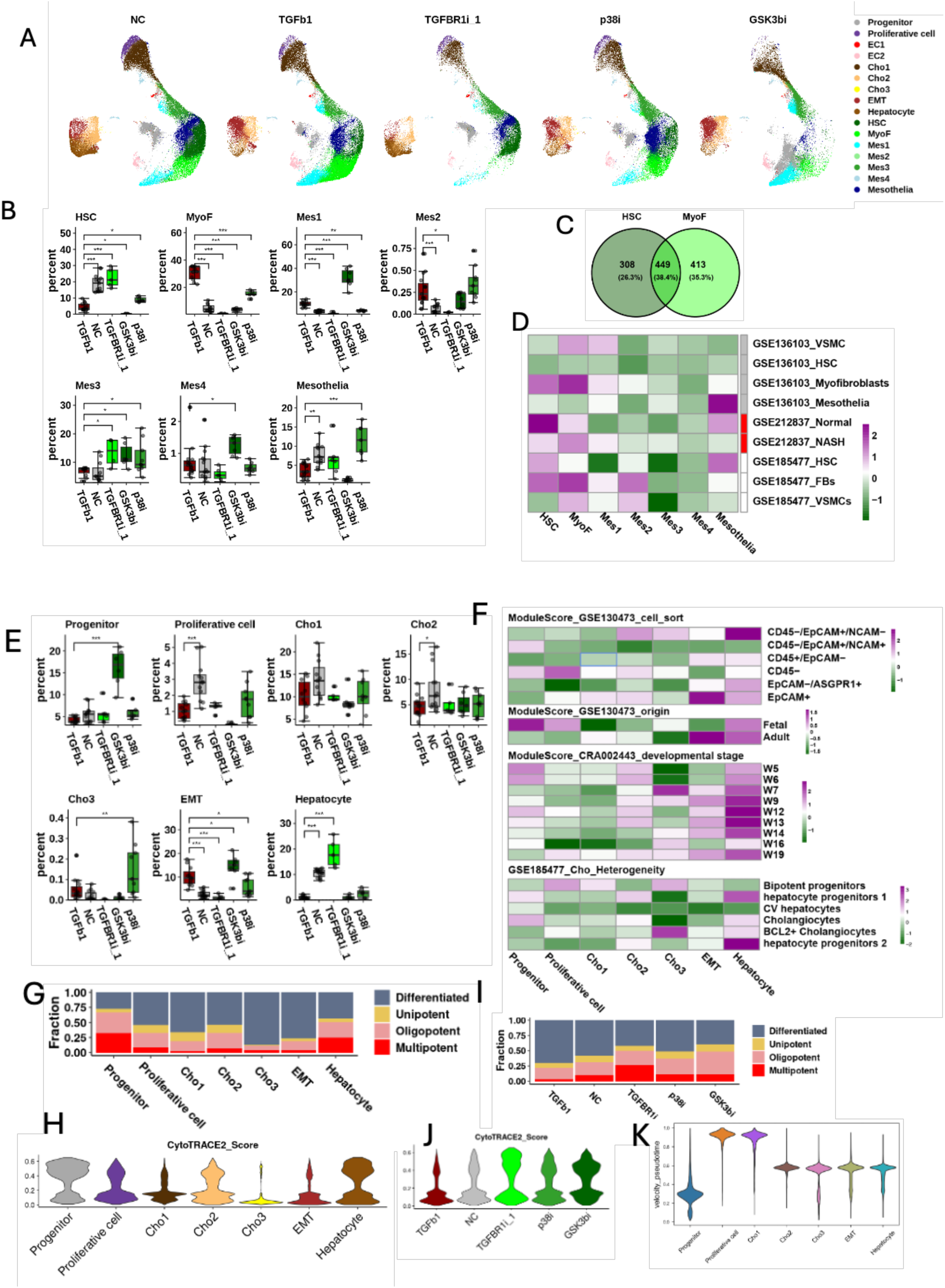
Characterization of the cellular changes in microHOs treated with TGFRB1, p38 or GSK3β inhibitors. (**A**) UMAPs are color coded for each of the 16 cell types present in NC or TGFβ1-treated microHOs, or microHOs treated with TGFβ1 and a TGFRB1, p38 or GSK3β inhibitor. (**B**) Graphs comparing the change in the percentage of the five mesenchymal cell types in microHOs shown in (A). Each group is the average of microHOs prepared from 5 to 10 separate experiments, and the bars show the mean±SEM. ANOVA test p-values for the for the indicated comparisons are *, <0.05; **, <0.01; or ***, <0.001. (**C**) This Venn Diagram shows number of differentially expressed or overlapping genes in the hepatic stellate cell (HSC) and myofibroblast (MyoF) clusters. (**D**) The relationship between the transcriptomes of the seven mesenchymal clusters (HSC, MyoF, Mes1-4 and Mesothelia) in microHOs and human liver disease cell types found in 3 reference datasets was assessed by calculating a module score. The datasets contain HSC in normal and cirrhotic human livers, vascular smooth muscle cells (VSMC), MyoF, and mesothelial cells in GSE136103; mesenchymal cells in normal liver and NASH liver (GSE212837); and HSC, VSMC and MyoF (FB) in GSE185477. (**E**) Graphs comparing the change in the percentage of the seven epithelial cell types in the microHOs shown in (A). (**F**) Module scores assess the relationship between the transcriptomes of the seven epithelial clusters (Progenitor, proliferative cell, Cho1-3, EMT and Hepatocyte) in microHOs with human fetal and adult liver cell types found 3 reference datasets: hepatobiliary hybrid progenitors in fetal liver (GSE130473); hepatogenesis in different developmental stages of fetal liver (CRA002443); or various types of cholangiocytes in human liver (GSE185477). (**G,H**) The ‘Cytotrace’ determined cell potency categories generated for the seven epithelial clusters in microHOs are plotted as percentage bar plot (G) or as a velocity pseudotime plot (H). (**I,J**) The ‘Cytotrace’ cell potency categories are plotted as percentage bar plot (I) or as a velocity pseudotime plot (J) generated from NC, TGFβ-treated. or TGFβ plus drug-treated microHOs. (**K**) The predicted ‘velocity_pseudotime’ for the seven epithelial clusters in microHOs.

TGFβ and the inhibitors also had significant effects (**Fig. 4E**) on the seven parenchymal epithelial clusters identified in microHOs, which include: a MKi67^+^ proliferative cluster, that expressed genes that resembled bi-potent progenitor cells; hepatocytes that expressed lipid metabolism genes; progenitor cells that resembled fetal liver cells; and EMT cells whose gene expression signatures were enriched with those characteristic of the epithelial to mesenchyme transition process, which resembled EPCAM^+^ fetal and adult liver cells. There were three cholangiocyte clusters: Cho1 had similarities with hepatoblasts; Cho2, which had similarities with hepatocyte precursors; and Cho3, which was a differentiated cholangiocyte (**Figs. 4F-H, S8).** While TGFβ significantly decreased the proliferative and Cho2 clusters, their abundance was not affected by the GSK3β, p38 or TGFBR inhibitors. The GSK3βI increased progenitor (but not proliferative) cells (Fig. 4E). While the TGFBRi and p38i suppressed the TGFβ-induced increase in EMT cells, the GSK3β inhibitor expanded this cluster. Only the TGFBR1i was able to revert the TGFβ-induced loss of hepatocytes. Overall, based upon the cell differentiation status determined by the Cytotrace and Velocity analyses, while TGFβ significantly decreased the progenitor pool, the GSK3β, p38 and TGFBR inhibitors were able to revert the loss of the stem cell pool (**Figs. 4I-K**). Also, the GSK3ýi acts via a different mechanism than TGFRB or p38i; while the GSK3ýi does not prevent TGFý intracellular signaling, it alters the TGFβ-induced changes in cellular differentiation. The TGFβ, Wnt/β-catenin and p38 pathways have multiple interaction points in liver (*45*),(*46*) and during development (*47*), which might explain how a GSK3βi alters TGFβ-induced cellular differentiation.

scRNA-seq data also indicates that the HDAC and BRD4 inhibitors had distinct effects on cells in microHOs (**Fig. 5A-B**). While both inhibitors countered the TGFβ-induced increase in MyoF; the BRD4i decreased Mes1 cells, but the HDACi increased Mes1 cells. Notably, the BRD4i idramatically increased the abundance of the most differentiated mesenchymal cluster (Mes2) (**Fig. 5C-F**). These inhibitors also had distinct effects on progenitor and differentiated epithelial cells (**Fig. 5G**). The BRD4 i decreased progenitor cell abundance. The HDAC and BRD4 inhibitors increased Cho1 and Cho3 cell abundance, respectively, which were the most differentiated epithelial cells (Fig. 4G-H). The HDAC inhibitor also increased EMT cells. The HDAC and BRD4 inhibitors did not revert the TGFβ-induced loss of hepatocytes in microHOs. While TGFβ significantly decreased the progenitor cell pool, BRD4i and HDACi increased the level of cell differentiation, and this effect was not due to cell cycle changes (**Fig. 5E-F**). These findings indicate that while the BRD4i and HDACs had different effects on mesenchymal cells, both promoted progenitor cell differentiation into more specialized cell types, which suggests that they could have utility for treatment of liver fibrosis.

**Figure 5.**
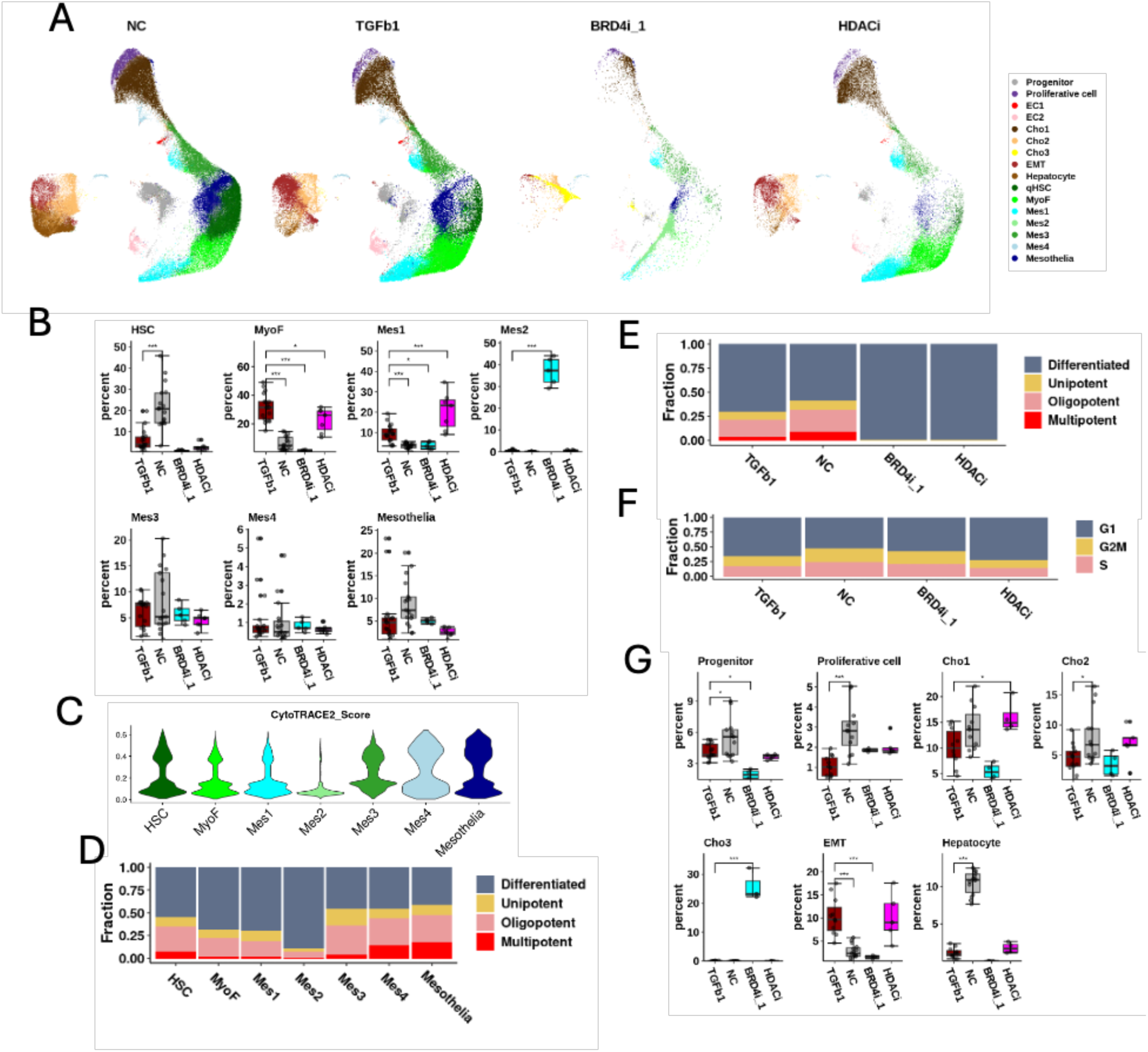
Characterization of the cellular changes in microHOs treated with BRD4 or HDAC inhibitors. (**A**) Individual UMAPs are color coded for each of the 16 indicated cell types in NC or TGFβ1-treated microHOs, or microHOs treated with TGFβ1 plus a BRD4 (Zn-3694) or HDAC (Vorinostat) inhibitor. (**B**) Graphs comparing the change in the percentage of each indicated mesenchymal cell type in the microHOs shown in (A). The bars show the mean±SEM for each group, which is the average of microHOs prepared from 5 to 10 separate experiments. ANOVA test p-values for the indicated comparisons are *, <0.05; **, <0.01; or ***, <0.001. (**C,D**) The ‘Cytotrace’ predicted potency scores are shown as velocity pseudotime plots (C) or as a percentage bar plot (D) for the seven mesenchymal clusters in microHOs. The mesenchymal cell type (Mes2) whose abundance is increased by BRD4 inhibitor treatment is the most differentiated mesenchymal cell type. (**E,F**) The ‘Cytotrace’ cell potency (E) and cell cycle (F) categories are plotted as percentage stacked bar plots for the microHOs with the indicated treatments. TGFβ1 induces an increase in the extent of differentiation of the cells in microHOs. BRD4 and HDAC inhibitors induce a substantial increase in the extent of cellular differentiation without altering the cell cycle. (**G**) Graphs comparing the change in the percentage of the seven mesenchymal cell types in the microHOs shown in (A). The bars and ANOVA p-values are as described in (B).

Since AI co-scientist also recommended that “*the Assay for Transposase-Accessible Chromatin sequencing (ATAC-seq) is used to map TGFβ- and drug-induced effects on accessible chromatin regions,” sn*ATAC-seq was used to analyze NC, TGFβ-treated, and TGFβ plus drug-treated microHOs. Quality control metrics (length/number of fragments, and transcription start site (TSS) enrichment) indicated that the ATAC-seq data was of high quality (**Fig. S9A-C**), and snATAC-seq UMAP plots corresponded with the scRNA-seq UMAP plots (**Fig. 6A**). Since differences in chromatin structure are an indicator of differential gene expression, chromatin structural changes near TSS were identified by calculating differential gene scores when different treatment groups or cell clusters were compared. There were 3527 marker genes with differential gene scores when the 16 cell clusters were compared, and hepatocytes had the highest number of differential gene scores (**Fig. S9D-E**). The dramatically different effects of TGFβ- or TGFβ+drug treatment is seen in a heatmap profiling the differential gene scores (n=399) for each condition. To examine drug effects on TGFβ-induced chromatin structural changes, the differential gene scores for TGFβ- or TGFβ +drug-treated microHOs were individually compared with NC microHOs. From these comparisons, 636 genes had differential gene scores (478 increased, 158 decreased) in the TGFβ-treated vs NC microHOs, and the TGFBR1i blocked all TGFβ-induced chromatin structural changes (**Fig. 6B-C**). Consistent with the importance of the p38 pathway, the p38i reduced the number of TGFβ-induced differential gene scores (21 increased, 9 decreased) by 94.3%. The GSK3βi reduced the number of differential gene scores (90 increased, 13 decreased) by 84%, while the BRD4i (212 increased, 205 decreased) caused a lesser reduction (43% reduction) in TGFβ-induced chromatin structure changes. The HDACi caused a 91% reduction in TGFβ-induced chromatin structural changes (34 increased, 21 decreased). It was noteworthy that the TGFβ- and drug-induced changes in the different cell types determined by analysis of snATAC-Seq data reflected those measured using scRNA-Seq data (**Fig. 6D-F**). Specifically, TGFβ induced a significant increase in the percentage of MyoFs, which was blocked by all five tested drugs. TGFβ also caused a significant decrease in the percentage of hepatocytes, which was countered by the TGFBR1, GSK3β and HDAC inhibitors; and the percentage of proliferative cells was increased by the BRD4i and HDACi. Hence, analysis of drug-induced changes in chromatin structure indicates that multiple mechanisms underlie the anti-fibrotic effects of the HDACi and BRD4i.

**Figure 6.**
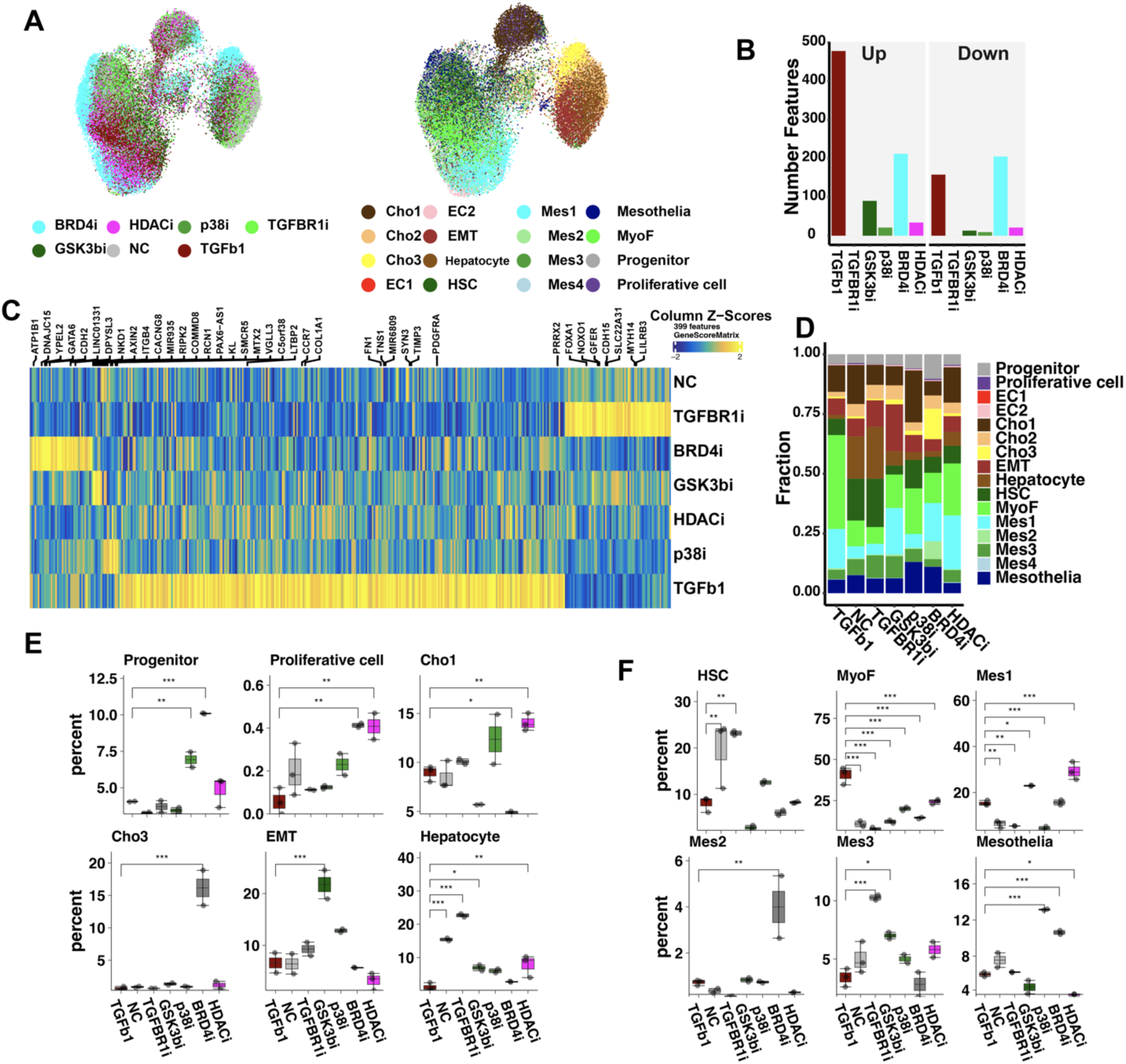
TGFβ1 and drug-induced epigenetic changes in microHOs. (**A**) Combined sample UMAP plots of the snATAC-seq data generated from microHOs that received the indicated treatments (Left); or with labeling of the 16 cell clusters, which were determined by transfer of the scRNA-seq data coordinates (Right). The data for each group was generated from three different experimental batches of samples, which have 10 or more microHOs in each batch. (**B**) These bars plot illustrates the number of differential marker genes identified by comparing the gene scores between each specified treatment and normal control (NC) microHOs. These comparisons are made to ensure that the TSS (Transcription Start Site) enrichment ratio and the log10-transformed number of fragments are similar between the treatment and NC groups. The “up” or “down” bars indicate whether the gene scores in the treatment group were higher or lower than in the NC group. The Wilcoxon test was used to analyze the differences in the gene scores between the treatment and NC groups for each gene. This statistical test identifies genes with significant differences in their gene scores, which highlights those that are differentially affected by treatment with TGFβ or TGFβ and drug. (**C**) A heatmap shows the z-score scaled ‘differential gene score’ calculated from snATAC-seq data for 248 features, which are organized by the indicated treatment. (**D**) The average percentage of each cell type present in NC, TGFβ- or TGFβ+drug-treated microHO. (**E, F**) Graphs comparing the change in the percentage of six epithelial (E) or mesenchymal (F) cell types in microHOs receiving the indicated treatment. Each group is the average obtained from analysis of snATAC-Seq data generated from 3 separate experimental batches of microHOs (with ∼30 microHOs per condition per batch), and the bars show the mean±SEM. ANOVA test p-values for the for the indicated comparisons are *, <0.05; **, <0.01; or ***, <0.001.

## Discussion

This study provides a first demonstration that a compound, multi-agent system, which was designed to mirror the reasoning process underlying scientific discovery, can assist in re-purposing drugs for treating a disease with limited therapeutic options. Two of three types of epigenomic modifier drugs recommended by AI co-scientist exhibited significant anti-fibrotic activity, and both had features that make them excellent candidates for treating liver fibrosis. We recently demonstrated that gene-phenotype analyses performed by a LLM could lead to novel genetic discoveries in mice and genetic diagnoses in humans (*48*). The results presented here demonstrate that a next generation LLM system (AI co-scientist) can analyze the vast amounts of scientific literature covering a specified area, identify hidden connections, and generate novel hypotheses that could produce new therapeutic approaches. Remarkably, AI co-scientist also guided the research steps used for testing the hypothesis. Due to the inherent complexity of biomedical research, AI-generated hypotheses require rigorous experimental validation by scientists. The purpose of these AI tools is to augment (and not replace) human scientific reasoning; their use will increase the ability of researchers to generate discoveries, while allowing them to retain intellectual oversight of the discovery process. Nevertheless, this work represents a significant milestone for AI, it indicates how AI-enabled science could significantly accelerate discoveries that could advance many biomedical fields.

Our results demonstrate that the microHO platform provides a robust, high-throughput platform for evaluating the anti-fibrotic and cytotoxic effects of drug candidates in a developing hepatic organoid. Moreover, the FCM, scRNA-seq and ATAC-Seq data provide key information about drug-induced effects on parenchymal cell regeneration, cell differentiation state, and cell composition in a fibrotic liver-like environment. The TGFBR1, GSK3β, p38 and HDAC inhibitors reduce fibrosis and increased the number of hepatic parenchymal cells after TGFβ exposure.

Transcriptomic analyses revealed that the anti-fibrotic effect of these drugs are mediated by different mechanisms. TGFBR1 and p38 inhibitors blocked TGFβ-induced intracellular signaling; while GSK3β, BRD4 and HDAC inhibitors altered TGFβ-induced mesenchymal cell differentiation. Vorinostat (Mes5), GSK3β (Mes5) and BRD4 (Mes6) inhibitors caused a marked increase in mesenchymal cell populations that were not present in NC or TGFβ-treated microHOs. While BRD4 inhibitors are still in clinical trials; Vorinostat (*36, 37*) is an FDA approved anti-cancer drug, which did not cause toxicity in microHOs nor did it cause hepatitis or clinically apparent liver injury in treated subjects (*49*). Vorinostat is being considered as a treatment for other types of cancers (*50, 51*), but it had not previously been considered for treatment of liver fibrosis. A very limited amount of prior experimental data supported its potential efficacy in liver fibrosis: it suppressed hepatic stellate cell activation *in vitro* (*52*), and reduced liver injury (*53*) and fibrosis (*54*) in rodent models. Since our microHOs do not have immune cells, the observed anti-fibrotic effects result from decreased production of activated myofibroblasts. Because epigenomic modifications effect gene expression and cellular differentiation, they play a role in the pathogenesis of multiple human diseases (*55, 56*) (including liver fibrosis (*57*)), and HDAC-mediated effects on histones play a particularly important pathogenetic role (*58*). Vorinostat and BRD4 inhibitors caused substantial reductions in TGFβ-induced chromatin structure changes, which indicates that this is likely to be a major contributor to their anti-fibrotic effect. It was also noteworthy that Vorinostat and BRD4 inhibitors potently induced cellular differentiation in microHOs. Anti-cancer drugs, including Vorinostat and BRD4 inhibitors, have been used as cellular differentiating agents for treatment of several types of cancer (*59–61*); and Vorinostat can induce mesenchymal cell differentiation (*62*). Our findings indicate that epigenomic modifying drugs should be given serious consideration as treatments of liver fibrosis and could form part of a new generation of anti-fibrotic agents that also promote liver parenchymal cell regeneration.

**Supplemental Data File 1**. The AI co-scientist prompt and the output generated in response to the prompt asking about the role that epigenetic changes play in Liver Fibrosis.

**Supplemental Data File 2**. The cell number within different treatments for all batches of microHOs combined are shown in table A. The cell number for different treatments per batch of microHOs generated from the scRNA-seq data is shown in Table B. The statistics used for comparing the changes in cell number in response to the different treatments in all cell clusters are shown in Tables C-D. Table 2E shows the number of single nuclei and the median number of fragments for each of the 24 samples used for snATAC-seq. Table 2F shows the list of genes from ‘differential gene score’ calculated from the snATAC-seq data organized by conditions. Table 2G shows the list of genes from ‘differential gene score’ calculated from the snATAC-seq data organized by cell cluster.

## Data availability

All raw and processed single cell RNA-seq data were deposited in the Gene Expression Omnibus (GEO) and are available under accession (GSE228214 and # pending). Processed scRNA-seq and snATAC-seq data are also available at 10.5281/zenodo.14996331.

## Code availability

We do not open-source the model code and weights due to the safety implications of unmonitored use of AI co-scientist. In the interest of responsible innovation, we will be working with research partners, regulators, and providers to validate and explore safe onward uses of AI co-scientist, and we expect to make them available via Google Cloud APIs. For reproducibility, we have documented technical deep learning methods while keeping the paper accessible to a clinical and general scientific audience. Our work builds upon Google Gemini whose technical details have been described extensively in a technical report (*63*).

## Author contributions

GP and YG formulated the project. YG, TT and GP wrote the paper with input from all authors. YG, JI, ZF, JL, and AAGB generated experimental data. APaw, JG, AD, AM, APal, LR, KK, WW, VN, AK, KR, YX, TT, and GP analyzed the data. All authors have read and approved of the manuscript.

## Acknowledgements

This work was supported by NIH awards (1R56DK12930901A1, 1R01DC021133 and 1 R24 OD035408) to GP and by an award from Google Cloud. We thank Dr. Bob Lewis for advice and careful review of this manuscript.

## Supplemental Information

### AI-assisted Drug Re-purposing for Human Liver Fibrosis

*Supplemental note 1: AI co-scientist description.* AI co-scientist ^1^ is a multi-agent system built on Gemini 2.0 that is designed to mimic the scientific method and accelerate scientific discovery using a “scientist-in-the-loop” paradigm. It utilizes a coalition of specialized agents that work asynchronously and collaboratively to help domain experts augment their hypothesis generation and experimental planning processes. The specialized agents used by AI co-scientist include:

**Generation Agent:** Generates initial hypotheses and research plans through literature exploration, simulated scientific debates, iterative assumption identification, and research expansion.

**Reflection Agent:** Acts as a scientific peer reviewer, critically assessing hypotheses for correctness, novelty, quality, and safety.

**Ranking Agent:** Employs an Elo-based tournament to rank hypotheses based on pairwise comparisons and scientific debates.

**Proximity Agent:** Calculates hypothesis similarity and builds a proximity graph to aid in de-duplication and idea exploration.

**Evolution Agent:** Refines and improves existing hypotheses by grounding them in literature, combining ideas, simplifying concepts, and out-of-box thinking.

**Meta-review Agent:** Synthesizes insights from all reviews, identifies recurring patterns, and generates meta-reviews to improve the performance of other agents and create research overviews.

Scientists can interact with the AI co-scientist system using natural language to define research goals (including the constraints on the hypothesis and the criteria for evaluating hypothesis quality), provide feedback, and refine hypotheses. The system uses self-play strategies, including a scientific debate and a tournament-based evolution process, to iteratively refine hypotheses and research proposals. It leverages a significant scaling of test-time compute to allow for iterative reasoning, evolution, and improvement of outputs, mimicking the iterative nature of scientific discovery. Additionally, the co-scientist uses various tools, including web search, specialized AI models, domain-specific databases (PubMed) to enhance grounding and the quality of generated hypotheses.

## Methods

The chemicals, drugs and antibodies used in this study are shown in Tables S1-S3.

*Generation of a collagen reporter iPSC line and microHO generation.* The human iPSC line (C3 COL1A1-P2A-Clover) used in this study was prepared as previously described ^2^. The biopsy sample used to generate it was obtained according to a protocol (number 10368) approved by the Institutional Review Board at Stanford. In brief, a clover expression cassette (**P2A-Clover**) was inserted at the COOH terminus of the endogenous *COL1A1* gene of the C3 iPSC line without a STOP codon using a sgRNA (TTGGGATGGAGGGAGTTTAC). To produce microHOs, 50∼60% confluent iPSC cultures were cultured in a series of differentiation media as previously described ^2^. First, a definitive endoderm differentiation media that consisted of Advanced DMEM/F-12 (Gibco, 12634010) + ITS (Gibco, 41400045) supplemented with 0.1 mM nonessential amino acids, 1 mM pyruvate and 2 mM L-alanyl-L-glutamine dipeptide (GlutaMAX). On days 1 to 2, 100 ng/ml Activin-A (Peprotech, Rocky Hill, NJ), 10 ng/ml BMP4 (Peprotech), 100 ng/ml bFGF (Peprotech), 3 mM CHIR99021 (Sellechchem, Houston, TX) and 10 uM LY294002 (Sellechchem) were added the culture. On day 3, 100 ng/ml Activin-A and 100 ng/ml bFGF were added the definitive endoderm differentiation medium. From days 4 to 9, 20 ng/ml FGF10 (Peprotech) and 20 ng/ml BMP4 were added to the hepatoblast medium, which consisted of which consisted of Advanced RPMI 1640 Medium (Gibco) that was supplemented with GlutaMAX and ITS. After day 9, the hepatoblasts were dissociated to single cells in Accutase (Invitrogen) medium with 10 uM Y-27632 (Santa Cruz Biotechnology, Dallas, Texas). Then, 10,000 cells/well were re-aggregated in low-cell-adhesion Nunclon™ Sphera™ 96 well Microplates (ThermoFisher) that contained serum-free HO growth and differentiation medium, which consisted of William’s E medium supplemented with 0.1% Polyvinyl alcohol (Sigma-Aldrich), 0.1 mM nonessential amino acids, 1 mM pyruvate, 2 mM L-alanyl-L-glutamine dipeptide (GlutaMAX), 10 mM Y-27632, 100 ng/ml EGF, 10 ng/ml HGF, 10 mM Dexamethasone and 10 mM Hydrocortisone (Sigma, St. Louis MO). On day 13, 50 ng/ml of TGFβ1 (Peprotech, 100-21; or Sinobiological, 10804-HNAC) and drugs at the indicated concentration were added to each microwell. The microHO cultures were allowed to differentiate over 21 to 31 days.

*Supervised image analysis*. High content screening data was collected from microHOs as previously described ^2^. In brief, for the COL1A1:Clover and Hoechst 33342 (Invitrogen, H3570) channels, z-stacked images were captured from day 14 to day 31 cultures with a Molecular Devices ImageXpress Micro Confocal system. For each 10X confocal z-stack field, maximum intensity projections (MIPs) were generated from all acquired z-planes (20 to 40 per microHO). The MIPs from each channel from the same microHO were used for region of interest (ROI) segmentation. Then, 18 features from two channels were extracted using customized FIJI macro code ^3^ to build the ‘Feature X organoid’ matrix with the features shown in Figure S2 ^4^.

This matrix is used to compute *Pearson’s* correlation coefficients that were used to generate the principal component analyses (PCA) and heatmaps shown.

*Flow cytometry.* Single cell suspensions were prepared from microHOs by protease digestion as described ^5^. Flow cytometry was performed using at a BD Accuri C6 flow cytometer and the conjugated antibodies listed in Table S2. The data were analyzed using FCS Express (denovosoftware.com) software. A density plot of forward scatter height (FSC-H) vs. forward scatter area (FSC-A) was used to exclude doublets.

***Trichrome staining***. At the experimental end point (between days 20 to 30), the NC, TGFβ1 and TGFβ1 drug treated microHOs were harvested, allowed to settle by gravity, and were embedded in low melting point agarose (IBI Scientific, Dubuque, Iowa). The embedded organoids were then processed by sectioning of paraffin-embedded microHOs to produce 10-micron sections. To assess the amount of collagen in the microHOs, trichrome staining of the sections was performed using the MASSON’S 2000 TRICHROME STAIN KIT according to the manufacturer’s instructions. Image segmentation and quantitative measurement of collagen rich areas was performed using our previously described methods ^6^. In brief, Fiji (2.1.0) implementation of ImageJ was used to quantify the areas of positive staining. The ‘Trainable Weka Segmentation’ plug-in was used to train the classifiers and calculate the test experimental image. Statistical analysis was performed using a one-way ANOVA to assess the overall differences between group means, and Tukey’s post-test was then used for pairwise comparisons to identify groups with significant differences in their means.

***scRNA-sequencing.*** Control, TGFβ1 and TGFβ1+drug-treated microHOs (n>30 per group) were harvested from multiple independently prepared sets of cultures, and single cell suspensions were prepared by protease digestion as described ^5^. Single cell suspensions were visually inspected under a microscope, cells were counted using a Scepter™ 2.0 Handheld Automated Cell Counter (EMD Millipore, Burlington, MA), and then resuspended in PBS with 0.01% BSA. scRNA-seq libraries were prepared by using the spit-pool based scRNA-seq method (Evercode™ WT mini/WT/Mega from Parse biosciences) according to the manufacturer’s instructions. In brief, 1500/4000/10,000 cells from each group were loaded for the first round of sample barcoding respectively according to which format of kit was used, and the final pooled cells were divided into 2/8/18 equal libraries for sequencing. The expression matrix was generated using the ‘Parse Biosciences analysis pipeline.’ A total of 60,701 features that were generated from 613,826 cells, which passed quality control for processing of the scRNA-seq data obtained from 17 experimental batches.

***scRNA-seq data analysis*.** The scRNA-seq data was imported into ‘Seurat’ ^7^ for the subsequent analysis steps. (i) Cells with unique gene counts <200 or >7000 or where the percentage of mitochondrial mRNAs was >25% were removed. (ii) The 17 batches of data were integrated by ‘RPCA’ processing methods using the Seurat integration function. (iii) 30 principal components were used to construct the shared nearest neighbor (SNN) graph, and the parameter regulating the resolution of the ‘Find Clusters’ program, was set to 0.5. This resulted in the identification of 17 unique clusters for all 613,826 cells. To identify the cell type of each cluster, differentially expressed genes for the pre-defined cell types were computed using the ‘FindAllMarkers’ function within the Seurat Package with the following parameters: only.pos = F, min.pct = 0.25, logfc.threshold = 0.25. Seurat then identifies the differentially expressed genes using the non-parametric Wilcoxon rank sum test. The top 50 upregulated DEGs were used for the GO biological process analysis, which was performed using the ‘clusterProfiler’ for GO over-representation analysis ^8^.

***Module score calculation***. Module scores were calculated to assess the relationship between the transcriptomes of microHO clusters and the different types of cells in human liver tissue. scRNA-seq data obtained from normal and cirrhotic human liver tissue (GSE136103) ^9^ was used to identify the different types of mesenchymal cells in liver (myofibroblast, hepatic stellate cell (HSC), vascular smooth muscle cell (VSMC) and mesothelia (Meso)). The differentially expressed genes for each mesenchymal cell type were used to calculate module score for each of the clusters in the microHOs. A subset of the HSC scRNA-seq data obtained from GSE212837 ^10^ was used to identify the normal and NASH HSC types. A subset of the mesenchymal cell scRNA-seq data obtained from GSE185477 ^11^ was used to identify normal HSCs, myofibroblasts (FB) and VSMC. For the epithelial cell clusters, scRNA-seq data obtained from GSE130473 ^12^ was used to identify the subpopulations of hepatocytes and cholangiocytes in fetal and adult liver. Also, hepatic lineage cells from different developmental stages in the human liver scRNA-seq data obtained from CRA002443 ^13^ was used to identify cell lineages in fetal liver. A subset of the cholangiocyte scRNA-seq data obtained from GSE185477 ^11^ was used to identify normal hepatic progenitors and the subpopulations of cholangiocytes.

***Cell type annotation***. R package Azimuth ^14^ was used to annotate microHO scRNA-seq data to the reference dataset. Azimuth uses a ‘reference-based mapping’ pipeline that inputs a count matrix; and then performs the normalization, visualization, cell annotation, and assesses differential expression of the data. This human reference consists of 79,492 integrated liver cells from data collected in 5 different studies ^9,15-18^.

***Cell status analysis***. The ‘Cellular potency categories’ and the absolute developmental potential were assessed using ‘CytoTRACE 2’ ^19^ and the microHO scRNA-seq data. The predicted ‘potency score’ provide a continuous measure of developmental potential; and they range from 0 (differentiated) to 1 (totipotent). The raw count from the Seurat object is used to directly to compute CytoTRACE score. ‘**RNA velocity**’ analysis infers the cell state by measuring the ratio of un-spliced to spliced mRNA transcripts, which provides information about the transcriptional activity of a gene and its direction. The transcript assignment file (tscp_assignment.csv.gz from the Parse pipeline output directory) contains the splicing information for each transcript identified in a Parse assay. The splicing information is used to generate the splice matrices required to run scVelo ^20^. The Anndata file, which is required for scVelo, was generated using the Seurat object that contains the metadata, and splicing matrix. RNA velocity was estimated with the ‘stochastic model’ (using second-order moments).

***Cluster pathway enrichment***. The pathways associated with the DEGs in is microHO cell cluster was performed using the ‘clusterProfiler’ package ^8^. The top 100 DEGs, which were ranked by their ‘avg_log2FC’ using the Seurat ‘FindAllMarkers’ function, are the input gene list for calculating the over-representation analysis (ORA). The databases used in this study are: Gene Ontology (GO, http://www.geneontology.org); Kyoto Encyclopedia of Genes and Genomes (KEGG, https://www.genome.jp/kegg/); ReactomePA (http://bioconductor.org/packages/ReactomePA); and the Molecular Signatures Database (MSigDb, http://software.broadinstitute.org/gsea/msigdb).

***snATAC-seq***. snATAC-seq used the same single cell samples as those prepared for scRNA-seq. For the snATAC-seq analyses, 24 batches of microHO samples were prepared. Single nuclei were isolated from those samples using the nuclei Isolation kit (Nuclei EZ Prep, Sigma Aldrich, NUC101-1KT) according to the manufacturer’s instructions and the samples were pre-indexed using the scATAC Pre-Indexing Kit (ScaleBio, San Diego, CA) according to the manufacturer’s instructions. The pooled single nuclei were loaded onto the 10xChromium Next GEM Single Cell v2 workflow.

***snATAC-seq Data Processing and analysis*.** The raw sequencing data were demultiplexed, and FASTQ files were generated using the the <u>ScaleTagToolkit</u> from the Scale Bio seq Suite Suite (https://github.com/ScaleBio/ScaleTagToolkit). This suite processes the data from demultiplex samples, and the 150 bp paired-end (PE) snATAC-seq reads were aligned to the GRCh38 (hg38) reference genome. The mapped Tn5 insertion sites (fragments.tsv files) from scaletagtoolkit were read into the ArchR (v1.0.3) R package ^21^. To ensure high-quality sequencing and signal-to-background ratio, cells with fewer than 1,000 unique fragments or TSS enrichment below 4 were filtered out. After filtering, there were 34,698 nuclei with an average of 7000 reads per nuclei. To calculate TSS enrichment, the genome-wide Tn5-corrected insertions were aggregated ±2,000 bp relative to each TSS (TSS-strand-corrected). This profile was normalized to the mean accessibility ±1,900–2,000 bp from the TSS, smoothed every 51 bp, and the maximum smoothed value was reported as TSS enrichment in R.

The Latent Semantic Indexing (LSI) dimensionality reduction was performed by appending fragment files from all 24 samples of microHOs analyzed. Chromatin-derived gene accessibility scores were computed by aggregating the snATAC-seq reads for each cell, which were weighted by their distance from each gene within its cis-regulatory domain. The Leiden clustering algorithm ^22^ was used to identify cell clusters in the LSI subspace. Canonical correlation analysis (CCA), implemented in Seurat ^7^, was used to align and match cells from the scRNA-seq and snATAC-seq experiments. Log2-transformed gene accessibility scores served as proxies for gene expression in snATAC-seq cells. For integration, we selected the 2,000 most variable genes from each modality and used Seurat’s FindTransferAnchors() function with CCA as the reduction method and k.anchor = 10. Each scRNA-seq cell was mapped to its nearest neighbor in snATAC-seq using a nearest-neighbor search in the joint CCA L2 space. Nearest neighbors were identified with the FNN R package (<u>FNN</u>), using the ‘kd_tree’ algorithm and Euclidean distance. The labels from the scRNA-seq data were transferred to snATAC-seq data for further chromatin accessibility analysis. Differential gene analysis was performed using the getMarkerFeatures function in ArchR, based on the gene scores computed from chromatin accessibility near gene coordinates. Differential genes between different treatment conditions and normal control cells were identified using FDR ≤ 0.05 and Log2FC ≥ 0.25 thresholds.

---

**Table S1.** Chemicals and reagents used in this study.

| Name | Company | Catalog # | Location |
|---|---|---|---|
| Matrigel | BD | 354234 | Milpitas, CA |
| BMP-4 (carrier-free) | Biolegend | 795606 | San Diego, CA |
| Thiazovivin | Medchemexpress | HY-13257 | Monmouth Junction, NJ |
| PFI-3 | Medchemexpress | HY-12409 | Monmouth Junction, NJ |
| (+)-JQ-1 | Medchemexpress | HY-13030 | Monmouth Junction, NJ |
| Galunisertib | Medchemexpress | HY-13226 | Monmouth Junction, NJ |
| GSK126 | Medchemexpress | HY-13470 | Monmouth Junction, NJ |
| GSK503 | Medchemexpress | HY-12856 | Monmouth Junction, NJ |
| Ipatasertib | Medchemexpress | HY-15186 | Monmouth Junction, NJ |
| Alantolactone | Medchemexpress | HY-N0038 | Monmouth Junction, NJ |
| I-BET-762 (Molibresib) | Medchemexpress | HY-13032 | Monmouth Junction, NJ |
| PFI-3 | Medchemexpress | HY-12409 | Monmouth Junction, NJ |
| (+)-JQ-1 | Medchemexpress | HY-13030 | Monmouth Junction, NJ |
| Galunisertib | Medchemexpress | HY-13226 | Monmouth Junction, NJ |
| GSK126 | Medchemexpress | HY-13470 | Monmouth Junction, NJ |
| GSK503 | Medchemexpress | HY-12856 | Monmouth Junction, NJ |
| Ipatasertib | Medchemexpress | HY-15186 | Monmouth Junction, NJ |
| Alantolactone | Medchemexpress | HY-N0038 | Monmouth Junction, NJ |
| I-BET-762 (Molibresib) | Medchemexpress | HY-13032 | Monmouth Junction, NJ |
| Sirolimus (Rapamycin) | Medkoo | 100766 | Durham, NC |
| Imatinib mesylate | Medkoo | 100470 | Durham, NC |
| FGF10 | Peprotech | 100-26 | Cranbury, NJ |
| BMP-4 | Peprotech | 120-05 | Cranbury, NJ |
| OSM | Peprotech | 300-10 | Cranbury, NJ |
| A-83-01 | Santa Cruz Biotechnology | sc-203791 | Santa Cruz, CA |
| PD0325901 | Santa Cruz Biotechnology | sc-205427 | Santa Cruz, CA |
| SB431542 | Sellechchem | S1067 | Houston, TX |
| SB202190 (FHPI) | Sellechchem | S1077 | Houston, TX |
| DAPT (GSI-IX) | Sellechchem | S2215 | Houston, TX |
| LDN-193189 | Sellechchem | S2618 | Houston, TX |
| SAG | Sellechchem | S7779 | Houston, TX |
| IWR1-en | Sellechchem | S7086 | Houston, TX |
| GSK3685032 | Sellechchem | E1046 | Houston, TX |
| Vorinostat | Sellechchem | S1047 | Houston, TX |
| ZEN-3694 | Sellechchem | E1517 | Houston, TX |
| GSK3685032 | Sellechchem | E1046 | Houston, TX |
| Vorinostat | Sellechchem | S1047 | Houston, TX |
| ZEN-3694 | Sellechchem | E1517 | Houston, TX |
| SB525334 | sigmaaldrich | S8822-5MG | St. Louis, MO |
| Glitazone | sigmaaldrich | CDS021593-50MG | St. Louis, MO |
| SB525334 | sigmaaldrich | S8822-5MG | St. Louis, MO |
| Glitazone | sigmaaldrich | CDS021593-50MG | St. Louis, MO |
| FGF2 | sinobiological | 10014-HNAE | Newark, DE |
| Activin A | sinobiological | 10429-HNAH | Newark, DE |
| EGF | sinobiological | 10605-HNAE | Newark, DE |
| HGF | sinobiological | 10463-HNAS | Newark, DE |
| FGF10 | sinobiological | 10573-HNAE | Newark, DE |
| TGF beta 1 | sinobiological | 10804-HNAC | Newark, DE |
| PI-103 | Tocris | 2930 | Minneapolis, MN |
| CHIR99021 | Tocris | 4423 | Minneapolis, MN |

**Table S2.** Information about the 15 drugs examined in this study.

| Drug | Target | Active Concentration |
|---|---|---|
| PFI-3 | SMARCA2/4 | 10 µM |
| SB202190 | p38 | 10 µM |
| CHIR-99021 (Laduviglusib) | GSK3β | 3 µM |
| GSK126 | EZH2 | 10 µM |
| GSK503 | EZH2 | 10 µM |
| GSK3685032 | DNMT1 | 10 µM |
| Galunisertib | ALK5 | 10 µM |
| SB431542 | ALK5 | 10 µM |
| Vorinostat (SAHA) | Pan-HDAC | 1 µM |
| ZEN-3694 | BRD4 | 10 µM |
| JQ-1 | BRD4 | 1 µM |
| I-BET-762 (Molibresib) | BRD4 | 10 µM |
| SB525334 | Alk5 | 10 µM |
| Glitazone | PPARγ | 10 µM |
| Imatinib | PDGFRB | 10 µM |

**Table S3.** The conjugated antibodies used in this study were all obtained from Biolegend.

| Conjugated abs | clone | vendor | Cat # |
|---|---|---|---|
| Alexa Fluor® 647 anti-human CD326 (EpCAM) Antibody | 9C4 | Biolegend | 324212 |
| APC anti-human CD133 Antibody | W6B3C1 | Biolegend | 397905 |
| APC anti-human CD26 Antibody | BA5b | Biolegend | 302709 |
| Alexa Fluor® 647 anti-human CD324 (E-Cadherin) Antibody | 67A4 | Biolegend | 324112 |
| Alexa Fluor® 647 anti-human CD29 Antibody | TS2/16 | Biolegend | 303017 |
| Alexa Fluor® 647 anti-human CD90 (Thy1) Antibody | 5E10 | Biolegend | 328115 |
| APC anti-human CD140b (PDGFRβ) Antibody | 18A2 | Biolegend | 323608 |

**Figure S1.**
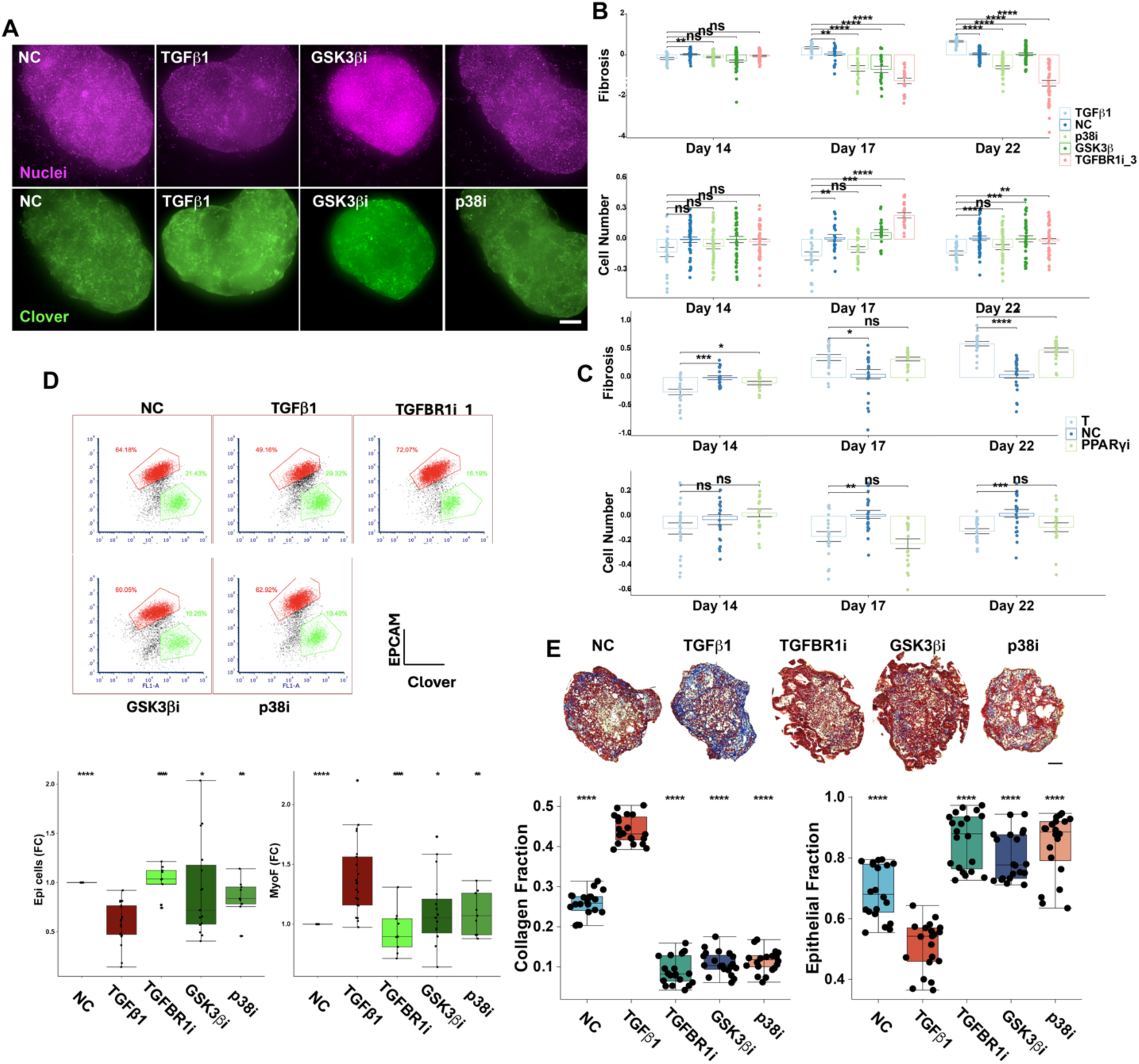
TGFBR1, p38 and GSK3β inhibitors block TGFβ1-induced fibrosis in microHOs. (**A**) Representative maximal intensity projection images obtained from microHOs treated with no addition (NC), 50 ng/mL TGFβ1, or TGFβ1 and the indicated drug. Scale bar, 100 μm. Clover expression is green, and nuclei stained with Hoechst 33342 are magenta. (**B**) microHOs were treated with either normal media (NC); 50 ng/mL TGFβ; or TGFβ with 10 μM of either p38 (SB202190) 3 μM GSK3β (CHIR-99021) or TGFBR1 (SB525334) inhibitor on day 13. The fluorescent signal from COL1A1^+^ cells and the total number of cell nuclei within a microHO were serially measured on days 14 through 22. **(C)** TGFβ-induced fibrosis in microHOs is not blocked by PPARγ inhibitors. microHOs were treated with either normal media (NC), 50 ng/mL TGFβ, or TGFβ and PPARγ inhibiter μM (10 μM) on day 13. The fluorescent signal from COL1A1^+^ cells and the total number of cell nuclei within a microHO were serially measured on days 14 through 22. In panels A-B, >30 microHOs were assessed per condition analyzed. **(D)** FCM results show that TGFBR1, p38 and GSK3β inhibitors decreased the MyoFs and increased the epithelial cell (EPCAM^+^) population in microHOs. *Left*: Representative FCM results obtained from microHOs receiving the indicated treatment. The x-axis shows the COL1A1 Clover fluorescence, and the y-axis shows the cells stained with an anti-EPCAM antibody. *Right*: The change in epithelial cells and MyoFs in microHOs caused by the indicated treatments. Each measurement is made on at least 10 individually prepared batches of microHOs. SB431542 (10 μM) is a TGFBR1 inhibitor; the GSK3β inhibitor is SB202190 (3 μM); and the p38 inhibitor is SB202190 (10 μM). To calculate the fold change (FC), the percentage of cells in each batch was normalized relative to that in NC microHOs. (**E**) Top panel: Images of Trichrome-stained TGFβ1-treated microHOs show a marked increase in collagen-rich connective tissue (blue-stained regions) relative to control (NC) microHOs, which only have a thin layer of connective tissue. The TGFβ1-induced increased in collagen was markedly inhibited by addition of the TGFBR1, GSK3β, or p38 inhibitors. The number of epithelial cells (dark red regions) was decreased by TGFβ1, but addition of TGFBR1, GSK3β, or p38 inhibitors prevented the TGFβ1-induced decrease in epithelial cells. The scale bar: 100 μm. Lower panel: Box plots show the area in microHOs with the indicated treatments (n > 10 per group) that was occupied by collagen (collagen fraction) or parenchymal (Epithelial fraction) cells. In all panels: ns, not significant; *, p-value<0.05; **, p-value<0.001; ***, p-value<0.001; and ****p-value<0.0001.

**Figure S2.**
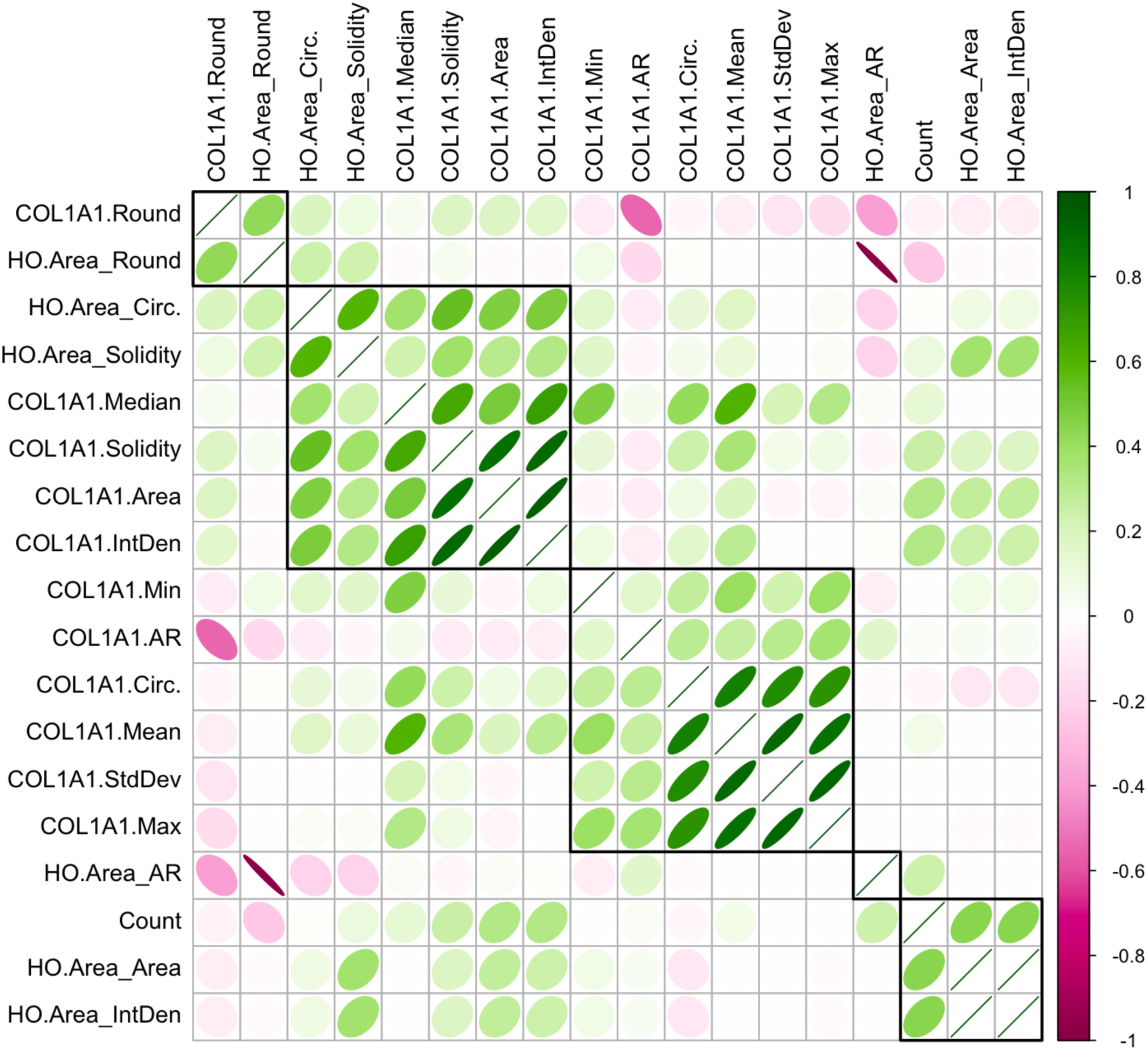
A heatmap formed using the Pearson’s correlation coefficients calculated from the 18 features (shown on each axis), which were generated from the COL1A1:Clover and nuclear (Hoechst 33342) staining channels, obtained from all organoids and all conditions examined in in this paper. The bar on the right shows the extent of the correlation (by color) that is shown in each square of the diagram (green is positive, magenta is negative). As can be seen, the features obtained from same channel cluster together. However, sometimes features measured using different channels will cluster together, which indicates that those features were similarly affected by the drugs.

**Figure S3.**
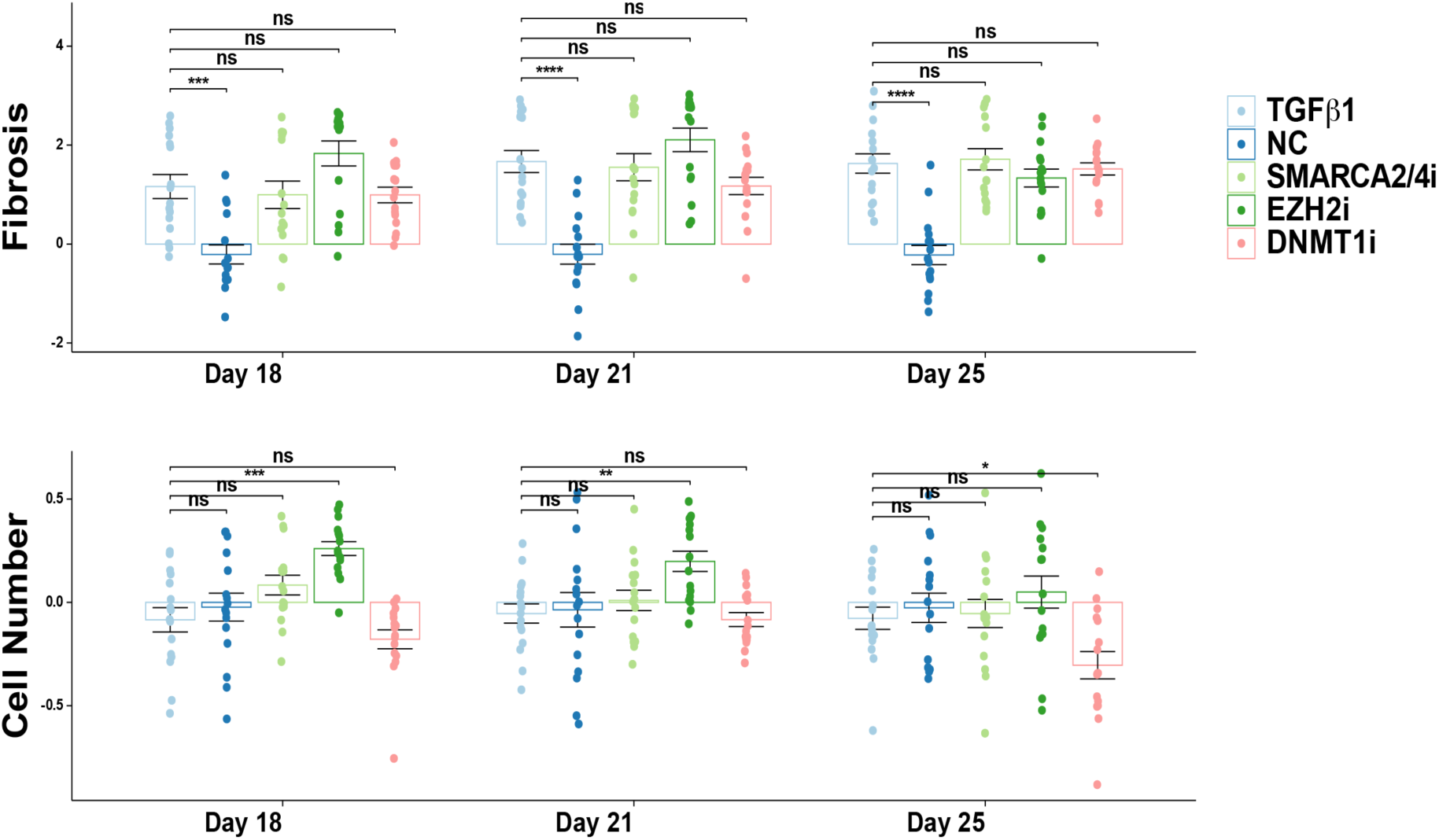
TGFβ-induced fibrosis in microHOs is not blocked by SMARCA2/4, EZH2 or DNMT1 inhibitors. microHOs were treated with either normal media (NC), 50 ng/mL TGFβ, or TGFβ and either 10 μM SMARCA2/4 (PFI-3), 10 μM EZH2 (GSK126) or 10 μM DNMT1 (GSK3685032) inhibitor on day 13. The fluorescent signal from COL1A1^+^ cells and the total number of cell nuclei within a microHO were serially measured on days 18 through 25, and 16 microHOs were assessed per condition analyzed. The DNMT1 inhibitor caused significant toxicity. ns, not significant; *, p-value<0.05; **, p-value<0.001; ***, p-value<0.001; and ****p-value<0.0001.

**Figure S4.**
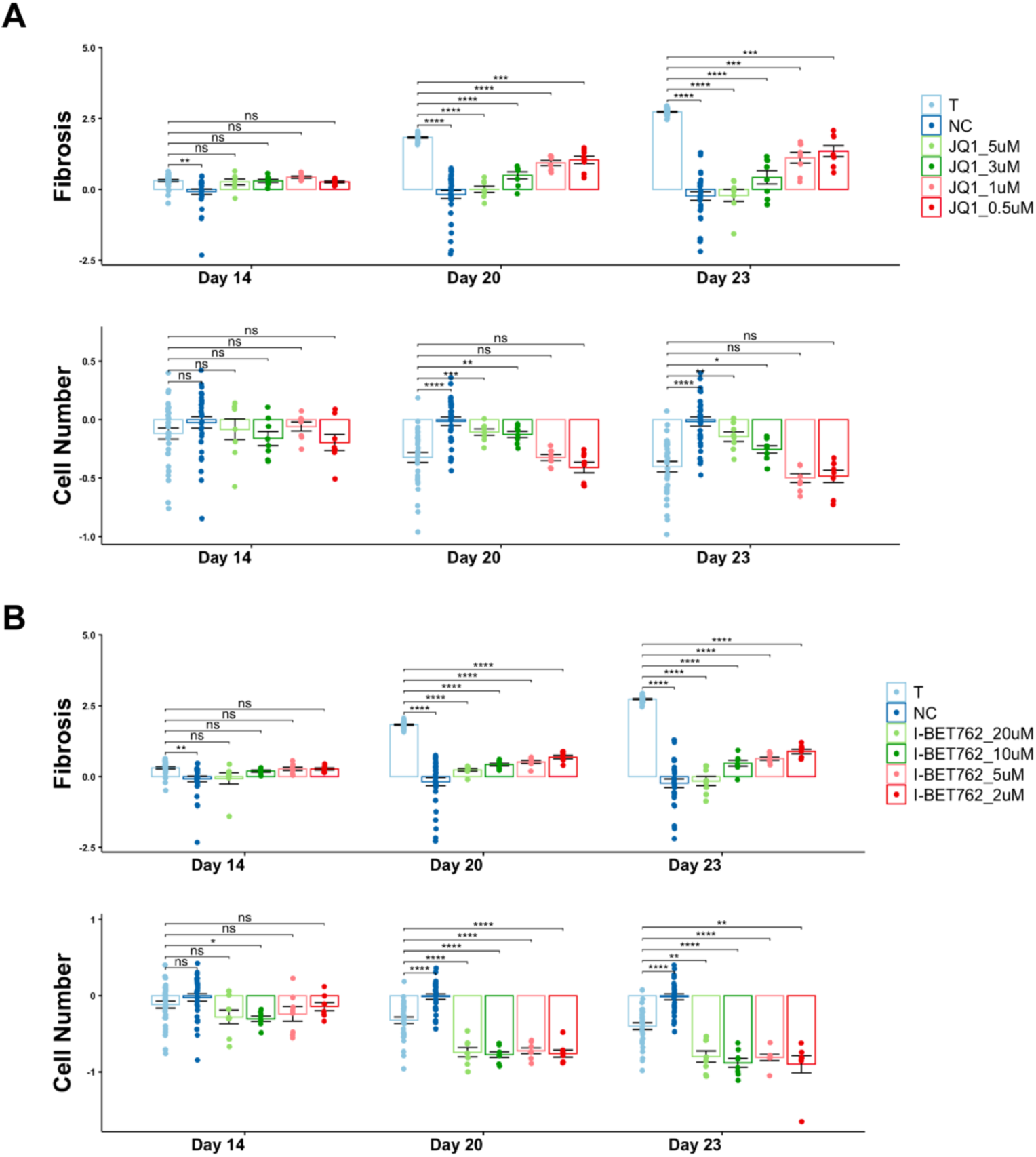
TGFβ-induced fibrosis in microHOs is blocked by BRD4 inhibitors. microHOs were treated with either normal media (NC), 50 ng/mL TGFb, or TGFb and the indicated concentration of JQ-1 (**A**) or I-BET762 (**B**). The fluorescent signal from COL1A1^+^ cells and the total number of cell nuclei within a microHO were serially measured on days 14 through 23, and 32 microHOs were assessed per condition analyzed. ns, not significant; *, p-value<0.05; **, p-value<0.001; ***, p-value<0.001; and ****p-value<0.0001.

**Figure S5.**
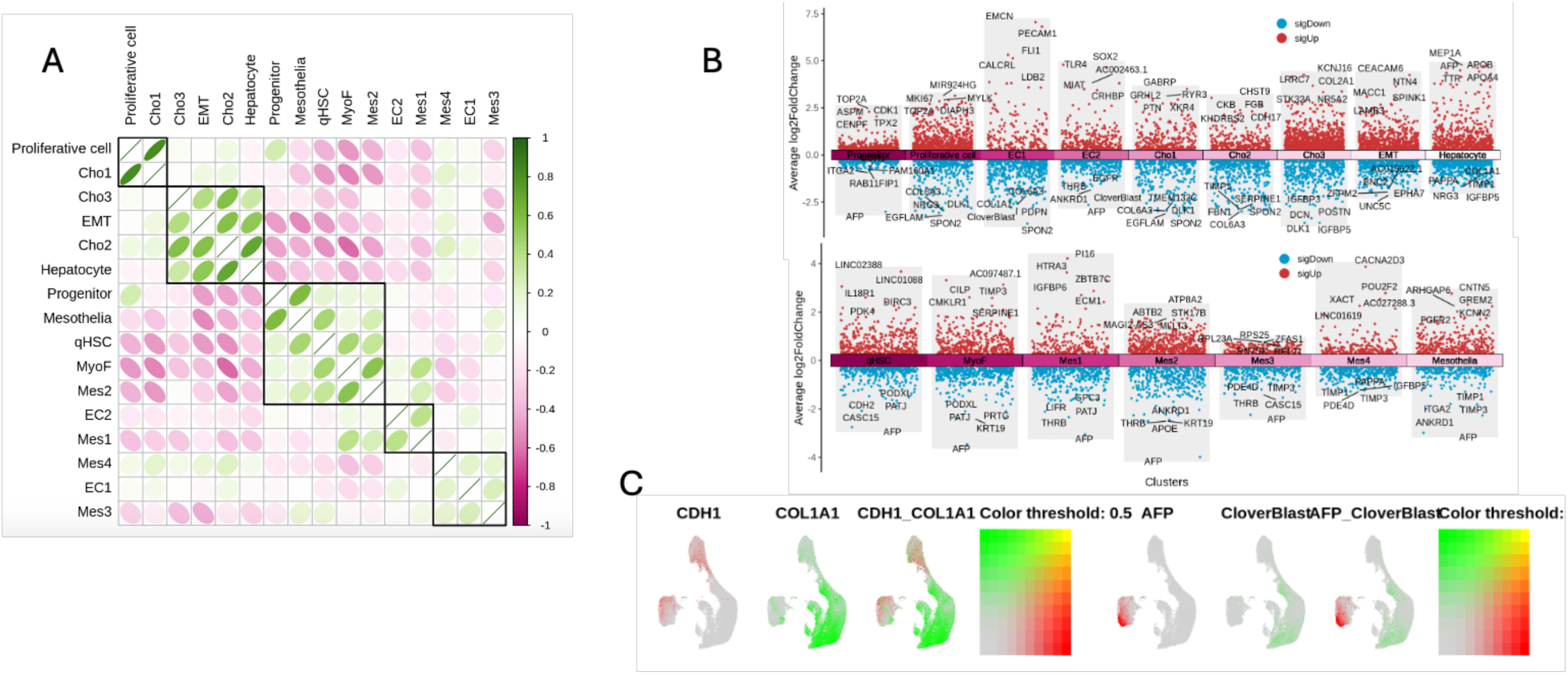
(**A**) A heatmap showing the Pearson correlation coefficients calculated for comparing the average transcriptome of each of the 16 cell clusters using the scRNA-seq data described in Figure 3A. The bar on the right shows the extent of the correlation (by color) that is shown in each square of the diagram (green is positive, magenta is negative). Note that the mesenchymal cells cluster together and are separated from the epithelial cell types, and vice versa. (**B**) A volcano plot showing the top 5 differentially expressed genes among the 16 cell clusters. As examples: the hepatocyte cluster (Epi4) expresses *AFP, TTR, CDH1(E-Cadherin)* and *APOB* mRNAs; the endothelial cell cluster (EC) express *PECAM1* mRNA; and the myofibroblast cluster (Mes1) expresses *TIMP3, COL1A1* and *Clover* mRNAs. (**C**) A blended feature plot showing the epithelial and mesenchymal cells in microHOs, which are separated by *CDH1* vs *COL1A1* or *AFP* vs *Clover* mRNA expression.

**Figure S6.**
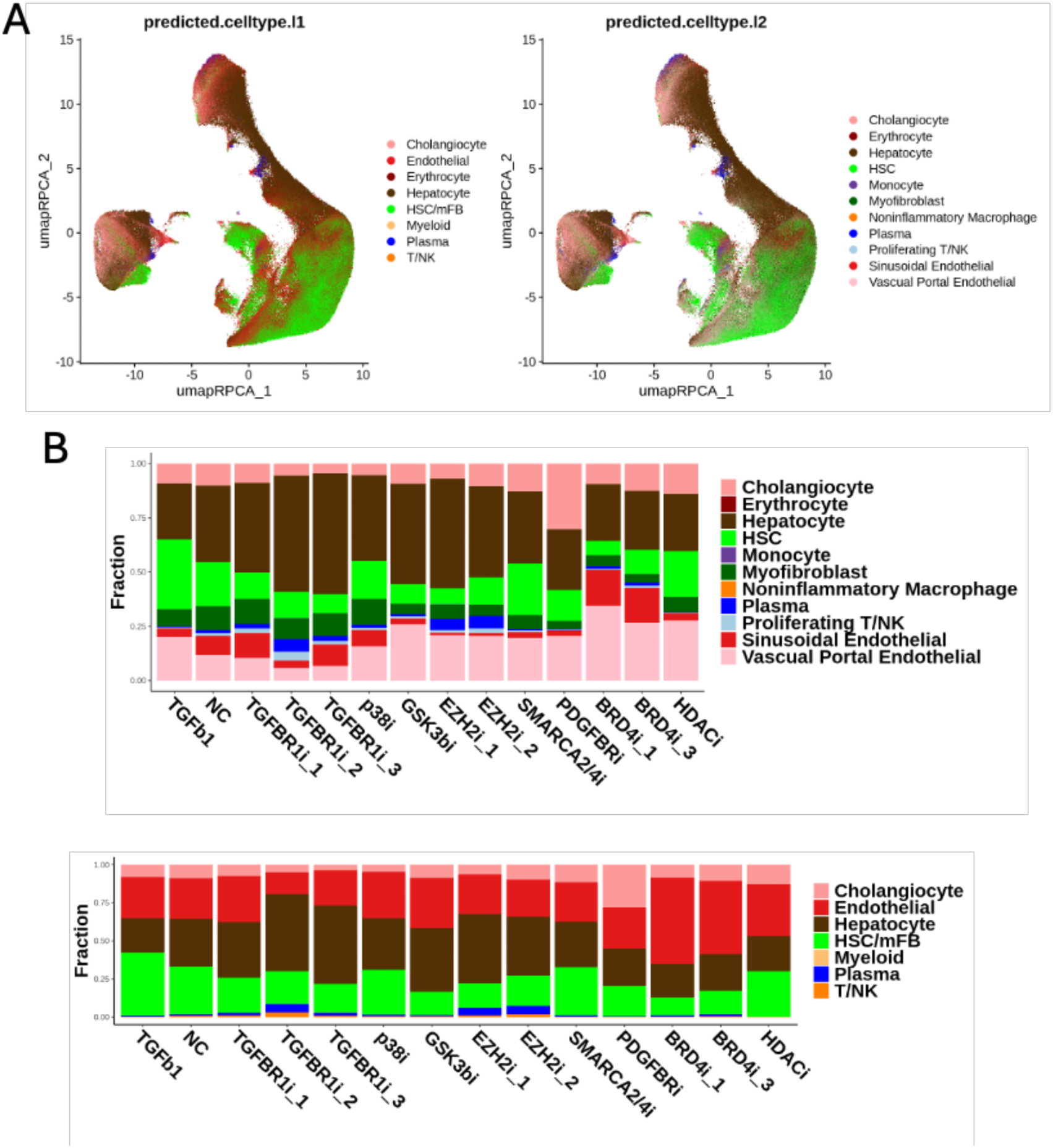
Cell type annotation of the clusters in the scRNA-seq data was determined using the Azimuth liver reference. (**A**) These the liver cell annotations were projected onto the UMAP of the microHOs. Transcriptomic comparisons, which were performed using with two different groups of annotated cells (l1, left; l2, right) in the Azimuth reference, were used to determine the cell types. (**B**) The percentage of each predicted cell type represents the average obtained from analysis performed on ten batches of microHOs, which were used to generate the scRNA-seq data for each of indicated condition.

**Figure S7.**
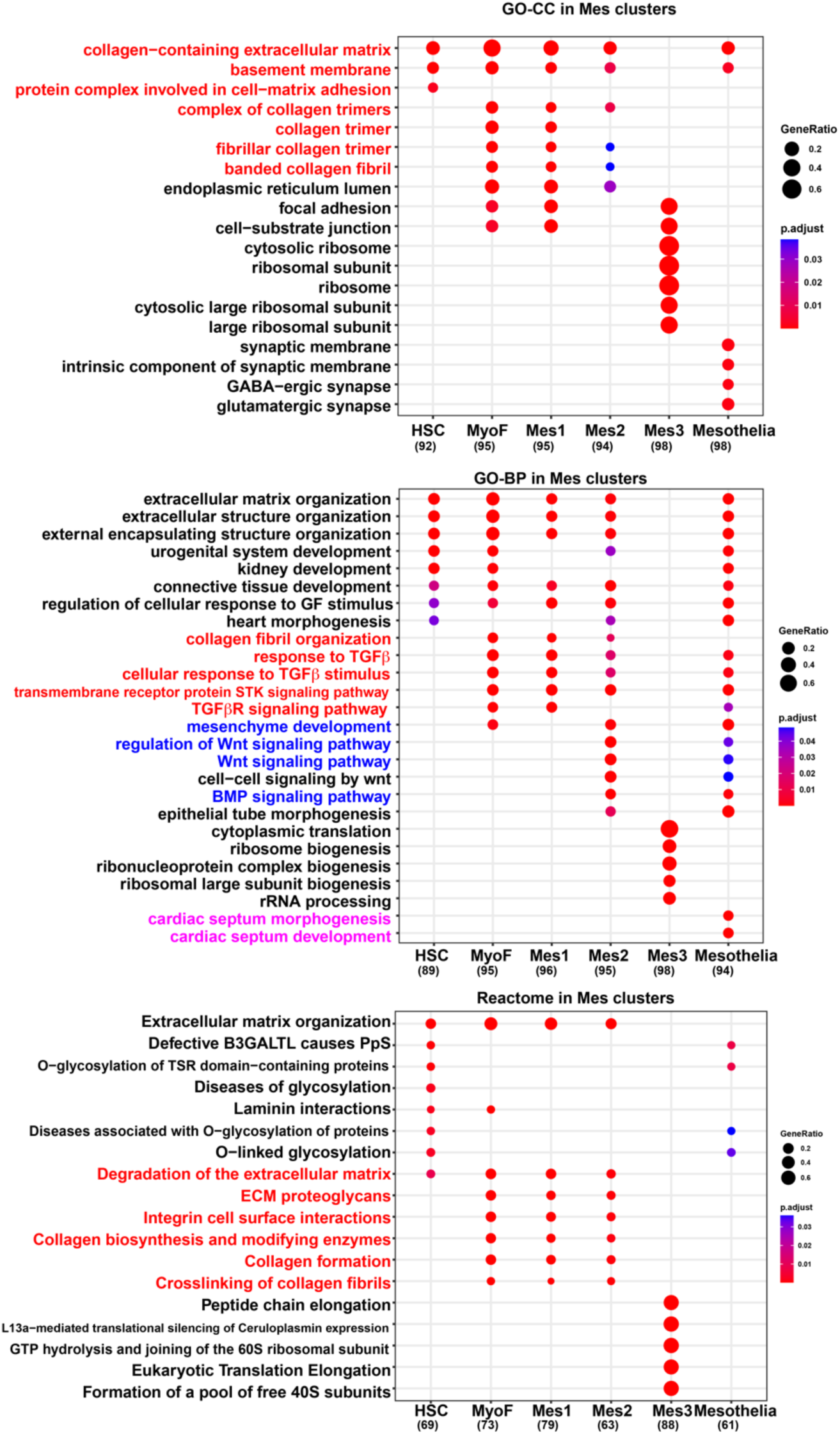
Pathway enrichment analysis for six mesenchymal clusters present in microHOs. The pathways associated with the top 100 DEGs for each cell type were annotated using the GO-CC, GO-BF and Reactome databases. The insets on the right show the gene ratio, which represents the proportion of the genes in the input gene list that are associated with a particular term; and the color-coded adjusted p-value for the statistical significance of the enrichment for a particular term in a given cluster of genes.

**Figure S8.**
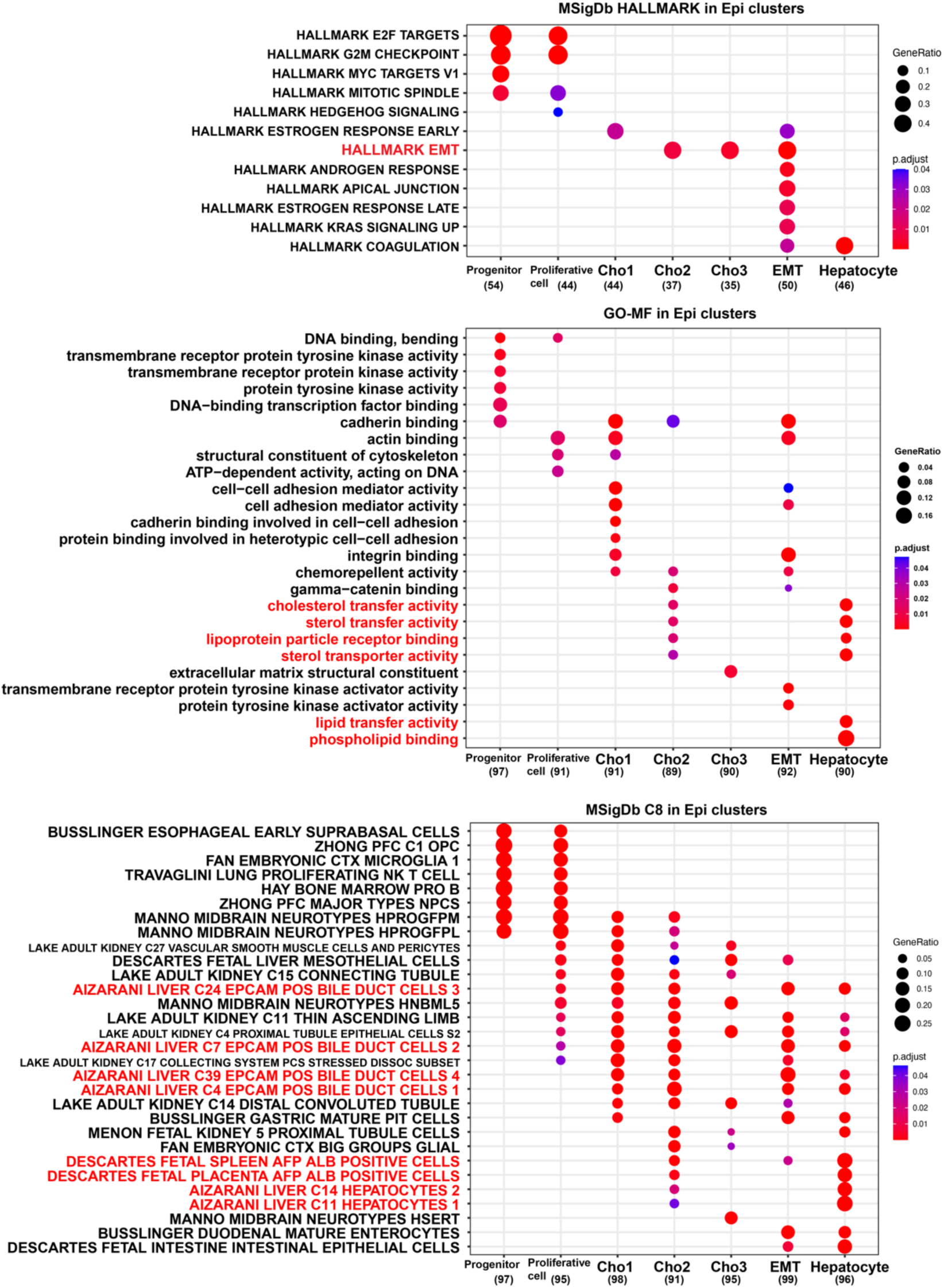
Pathway enrichment analysis for seven epithelial clusters present in microHOs. The pathways associated with the top 100 DEGs for each cell type were annotated using the MSigDb Hallmark, GO-MF and MSigDb C8 databases. The insets on the right show the gene ratio, which represents the proportion of the genes in the input gene list that are associated with a particular term; and the color-coded adjusted p-value for the statistical significance of the enrichment for a particular term in a given cluster of genes.

**Figure S9.**
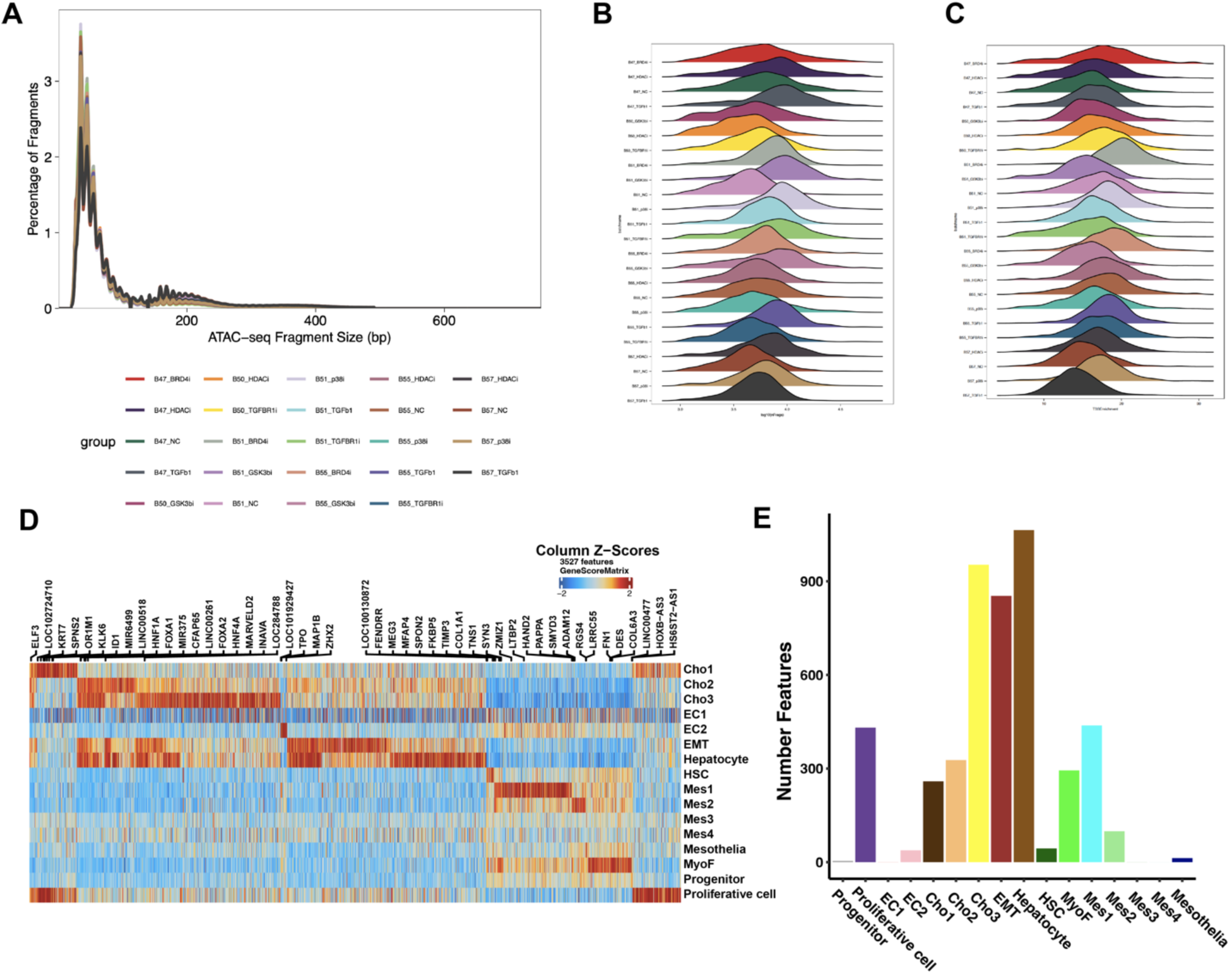
Quality control information for the ATAC-seq data. The panels show the Fragment length (**A**), the (log10 transformed) number of fragments (**B**), and TSS enrichment scores (**C**) in each microHO sample analyzed. **(D**) A heatmap showing the z-score scaled ‘differential gene score’ for 3527 features calculated from the snATAC-seq data, which are organized by the 16 cell types that were identified using the scRNA-seq data. (**E**) A bar plot showing the number of features identified in (C) for each of the 16 cell types.

## Supplemental Data File 1

### AI co-scientist Prompt and Output for the Role of Epigenetic Changes in Liver Fibrosis

**Input prompt:** Develop hypotheses about the genes and epigenomic changes required for myofibroblast generation in liver fibrosis and indicate what drugs should we test as new treatments for liver fibrosis. Novel experiments performed in hepatic organoids are preferred. At the end: judge the novelty of the hypotheses and the experiments, were similar experiments already done, and could any liver fibrosis researcher easily come up with the same idea?

*Background text provided (from our prior papers)*. Liver fibrosis is caused by extracellular matrix accumulation (ECM) in response to chronic liver injury. This fibrotic state results from an interaction between parenchymal and nonparenchymal liver cells and possibly involves infiltrating immune cells. The key non-parenchymal cell is the myofibroblast, which is generated when a fibrogenic stimulus causes hepatic stellate cells (HSC) to transdifferentiate into Myofibroblast that produce fibril-forming collagens. TGFβ1 and PDGF are key growth factors that drive Myofibroblast transdifferentiation. TGFβ1 binding to its type I receptor induces phosphorylation of downstream SMAD proteins, which ultimately promotes the transcription of ECM components. PDGF is a well-known mitogen for HSCs. Although it is most commonly an acquired condition caused by viral infection, non-alcoholic steatohepatitis (NASH), or chronic alcohol exposure; a few genetic diseases can also cause it. The rate of progression and histological features can vary in response to different inciting causes but activated myofibroblasts and the collagens (and other extracellular matrix proteins they produce are essential drivers of all forms of liver fibrosis. Patient outcome is determined by the extent of the fibrosis, which is a major global cause of death (∼1M per year). Although several drugs are being studied, there are no effective treatments (other than if the underlying cause can be treated) and the poor quality of available model systems has been a barrier to identifying anti-fibrotic therapies. Prior in vitro models did not have the spectrum of cell types that mediate fibrogenesis nor could they reproduce its key feature (i.e., generation of thick collagen filaments). Conclusions drawn from animal models are limited by concerns about species-specific differences in fibrotic mechanisms. Hence, the ability to analyze a fibrosis like that in human patients in a multi-lineage human hepatic organoid (HO) in a microwell could advance liver fibrosis research.

We developed a model system that differentiates human iPSCs into a multi-lineage 3-D HO that has liver lobule architecture with hepatocytes, cholangiocytes, and other cell types. We engineered disease-causing mutations into the genome of iPSCs and demonstrated that hepatic organoids provide a unique system for characterizing the impact of disease-causing genetic mutations. Transcriptomic and metabolomic analysis of early hepatic organoid development identified an essential biosynthetic pathway for early liver development, and we demonstrated that a commonly used over-the-counter medication (meclizine) that inhibits the rate-limiting enzyme in this pathway could be repurposed for treatment of liver cancer. scRNA-Seq analysis revealed that HOs contain cells of other lineages (macrophages, endothelial cells, and fibroblasts) and formed more complex structures than was appreciated in our earlier studies.

Hepatic organoids expressing an ARPKD mutation developed ARPKD liver pathology (abnormal bile ducts, hepatic fibrosis) in 21 days. Second harmonic generation microscopy (SHG) results demonstrated that ARPKD hepatic organoids had a marked increase in collagen and in thick collagen fiber formation. scRNA-Seq analysis indicated that ARPKD organoids had a markedly expanded population of collagen-producing MyoF (29% vs 3% in control) with an activated PDGF receptor β (PDGFRβ)-STAT3 signaling pathway. To study liver fibrosis, we developed a live cell-imaging method that enables serial monitoring for the appearance of collagen producing cells in hepatic organoids. We recently developed protocols for growing HOs in microwells (i.e., ‘microHOs’) and demonstrated that they form thick collagen filaments after PDGF or TGFβ exposure. Liver fibrosis can best be studied using this innovative live cell microHO imaging system with a Myofibroblast-driven fibrosis characterized by thick collagen filament formation.

### AI co-scientist Output

**Example experiment 1** (score=novelty+details+usefulness+pairwise rank=11):

### ## Prompt: A Novel Hypothesis Regarding Myofibroblast Generation in Liver Fibrosis

We propose a novel hypothesis that specific epigenetic alterations contribute to myofibroblast (MyoF) formation in liver fibrosis. Unlike previous approaches that primarily focused on canonical TGFβ1 and PDGF pathways, we hypothesize that changes in the cell’s epigenetic landscape play a critical role in regulating MyoF differentiation and subsequent ECM production.

Specifically, our hypothesis centers on the following key elements:

- **Histone Modifications:** We theorize that during fibrogenesis, alterations in histone modifications, particularly deacetylation, in the promoter regions of genes responsible for MyoF differentiation occur. For example, reduced acetylation of Histone 3 lysine 9 (H3K9ac) could promote the recruitment of repressive transcriptional machinery, silencing pro-MyoF factors. Alternatively, increased acetylation on certain histones might facilitate the binding of transcription factors that drive MyoF identity.
- **DNA Methylation:** Our hypothesis also posits that dynamic changes in DNA methylation patterns around genes involved in the MyoF pathway could be crucial. For instance, we predict that demethylation of CpG islands near regulatory elements of pro-MyoF genes might lead to enhanced transcription. Conversely, hypermethylation of the same regions might prevent the binding of critical activators, thereby inhibiting MyoF transition.

These two mechanisms, working in concert, could provide a complex regulatory system governing MyoF generation and subsequent ECM deposition impacting liver fibrosis.

### ## Groundbreaking Experiment to Test the Hypothesis

To test our hypothesis, we propose an innovative experiment utilizing the human micro-HO system:

Experiment Setup:

**1. micro-HO Generation:**

* Human micro-HOs will be generated from iPSCs according to the established protocol. These micro-HOs faithfully recapitulate the architecture and cellular composition of the human liver, including hepatic stellate cells (HSCs).

**2. Fibrosis Inducement:**

* micro-HOs will be exposed to a relevant fibrotic stimulus (e.g., TGFβ1 or PDGF) to induce the fibrotic response and MyoF generation.

* A parallel group will be kept untreated as controls.

**3. Epigenetic Profiling:**

* **ChIP-Seq**: At different time points during the fibrosis induction process, we will perform chromatin immunoprecipitation with sequencing (ChIP-Seq) to map the genome-wide landscape of relevant histone modifications such as H3K9ac, and relevant histone methylases/demethylases.

* **Methylation analysis**: Using bisulfite sequencing or similar approaches, we will assess the

DNA methylation status of critical MyoF-associated genes.

**4. Candidate Identification:**

* The epigenomic analysis will guide the identification of regions with significant epigenetic changes during fibrosis development.

* Bioinformatic analysis will be conducted to correlate the epigenetic alterations with candidate genes involved in MyoF differentiation.

**5. Functional Validation:**

* The epigenetic modifications and associated candidate genes will be validated using targeted approaches. This includes:

* **CRISPR/Cas9 Epigenetic Editing:** To directly confirm the role of specific epigenetic changes, we will manipulate the epigenome using CRISPR/Cas9-based tools directly or by employing dCas9 systems fused to epigenetic writer/eraser domains.

* **Pharmacological Modulation:** To assess the therapeutic potential, pharmacological targeting of identified epigenetic modifiers will be performed. **Drugs like inhibitors of DNA methyltransferases (e.g., Azacitidine) and histone deacetylase (HDAC) inhibitors (e.g., Vorinostat) could be tested for their impact on MyoF generation and ECM production in the micro-HO model.**

## Expected Outcomes

* Identification of key transcriptional regulators with specific epigenetic alterations driving MyoF formation and liver fibrosis.

* Validation of epigenetic markers as new diagnostic tools for liver fibrosis.

* Discovery of potential therapeutic targets through specific epigenetic modifiers.

## Additional Techniques

* **Transcriptomic Analysis:** Alongside epigenomic data, RNA sequencing will be performed throughout the experiment to track the dynamics of gene expression during the fibrotic process.

* **Live Cell MyoF Imaging:** The micro-HO platform, in combination with collagen-specific fluorescent probes and second harmonic generation (SHG) microscopy, will allow for real-time live cell monitoring of MyoF emergence and ECM generation, providing invaluable insights into the kinetics of fibrosis progression.

**## Judgments and Discussion Novelty and Impact:**

This hypothesis and experiment hold significant novelty as they depart from a sole focus on signaling pathways and explore the role of epigenetic mechanisms in MyoF development. The human micro-HO model, alongside powerful techniques like ChIP-Seq and live cell imaging, provide unprecedented access to a detailed understanding of fibrogenesis in a human context. This research could have a profound impact on liver fibrosis research and therapeutic advancement.

### Uniqueness of this Approach

Investigating epigenetic modifications in MyoF generation during liver fibrosis hasn’t been extensively explored previously. While some studies investigated epigenetic changes related to HSC activation, the comprehensive and dynamic analysis we propose using micro-HOs is a significant step forward. This approach holds high potential for a deeper understanding of a complex and crucial cellular phenomenon in liver fibrosis.

**Example experiment #2** (score=novelty+details+usefulness+pairwise rank=11): ## Novel Hypothesis Regarding Myofibroblast Differentiation in Liver Fibrosis:

We hypothesize that specific epigenetic alterations, in addition to known growth factor signaling, are essential for driving the transdifferentiation of hepatic stellate cells (HSCs) into myofibroblasts – the primary collagen-producing cells responsible for fibrosis. Specifically, we propose that activation of HSCs during liver injury triggers changes in DNA methylation patterns and histone modifications at critical gene loci. These epigenetic modifications promote the expression of genes involved in:

* **Collagen production and ECM remodeling:** Genes like ‘COL1A1’, ‘COL3A1’, and matrix metalloproteinases (MMPs).

* **Myofibroblast phenotype acquisition:** Transcription factors like ‘SNAI1’, ‘TWIST1’, and smooth muscle actin (αSMA).

* **Fibrotic signaling pathways**: Genes encoding components of TGFβ, PDGF, and Hedgehog pathways.

Furthermore, based on our preliminary scRNA-seq data from ARPKD hepatic organoids, we posit that these epigenetic modifications might be driven, in part, by the STAT3 signaling pathway which is downstream of PDGFRβ activation.

\## Experiment Targeting the Epigenetic Landscape:

**Objective:** To determine if epigenetic modifiers can block myofibroblast transdifferentiation and reduce collagen production in the microHO live-cell fibrosis model.

## Experimental Setup

**1. MicroHO Preparation:**

* Hepatic organoids will be generated from human iPSCs and differentiated into microHOs using established protocols.

* To mimic chronic liver injury, microHOs will be exposed to low, continuous doses of TGFβ1 and/or PDGF for a defined period (e.g., 7 days).

**2. Epigenetic Modifier Treatment:**

* Based on prior research and known mechanisms, we will select various classes of epigenetic modifiers for testing, including:

* **DNA Methyltransferase Inhibitors (DNMTis):** Examples include 5-Azacytidine and Decitabine, which deplete DNA methylation and reactivate silenced genes.

* **Histone Deacetylase Inhibitors (HDACis):** Examples like Trichostatin A (TSA) and Vorinostat increase histone acetylation leading to a more open chromatin state and favoring gene expression.

* **Bromodomain Inhibitors:** Such as JQ1, targeting epigenetic readers that recognize acetylated histones, thereby disrupting transcription.

**3. Assessment of Myofibroblast Transdifferentiation:**

* **Live-cell imaging**: microHOs will be monitored daily for the emergence of collagen-producing cells utilizing our established live-cell imaging system.

* **Second Harmonic Generation (SHG) microscopy:** Visualization of thick collagen fiber formation as a hallmark of fibrosis will be conducted.

* **Immunofluorescence staining:** Quantification of myofibroblast markers like αSMA and Col1A1 expression in fixed microHOs.

* **scRNA-seq analysis:** Global transcriptomic changes upon epigenetic modifier treatment will be evaluated to assess:

* Alterations in the myofibroblast gene expression signature.

* Expression changes in TGFβ, PDGF, and STAT3 pathway components

* Identification of specific genes activated or repressed by epigenetic modifications.

**4. Evaluation of Collagen Production and ECM deposition:**

* **Quantitative analysis of collagen content:** Utilizing biochemical assays like Sircol assay or hydroxyproline quantification.

* **Measurement of MMP activity:** Assays will assess the ECM remodeling dynamics influenced by epigenetic modifications.

### Expected Outcome and Impact

This experiment has the potential to:

* Uncover novel epigenetic changes driving myofibroblast transdifferentiation and collagen production in liver fibrosis.

* Determine the efficacy of epigenetic modifiers in reducing or reversing hepatic fibrosis in the microHO model.

* Identify promising drug targets within the epigenetic landscape of fibrosis.

* Provide a valuable platform for drug discovery and validation for liver fibrosis treatments.

### Novelty and Significance

* **Innovative Hypothesis:** Linking specific epigenetic changes to myofibroblast transdifferentiation represents a novel departure from the prevailing focus solely on growth factor signaling.

* **Advanced Experimental Platform:** The utilization of human microHOs offers a physiologically relevant and species-specific model system to study fibrosis.

* **Detailed Experimental Design:** The multi-faceted approach employing live-cell imaging, molecular biology, and transcriptomics provides a comprehensive analysis of myofibroblast behavior and collagen deposition.

### Similarity assessment

While studies have explored epigenetic factors in broad cellular models of fibrosis, this experiment’s strength lies in its specific focus on myofibroblast generation and collagen production within the context of the innovative microHO liver fibrosis model. This detailed approach encompassing multiple analytical techniques provides a more nuanced understanding of the epigenetic landscape. While other groups may explore aspects of epigenetics in fibrosis, the proposed combination of hypothesis, methodology, and model system likely distinguishes it from existing research.

### cell# per condition

| NC | TGFb1 | TGFBR1i_1 | TGFBR1i_2 | TGFBR1i_3 | p38i | GSK3bi |
|---|---|---|---|---|---|---|
| 135048 | 118302 | 60056 | 5909 | 29459 | 77500 | 80381 |
| EZH2i_1 | EZH2i_2 | SMARCA2/4i | PDGFBRi | BRD4i_1 | BRD4i_3 | HDACi |
| 4712 | 6867 | 15323 | 11440 | 22337 | 8998 | 37494 |

### cell# per condition per Batch

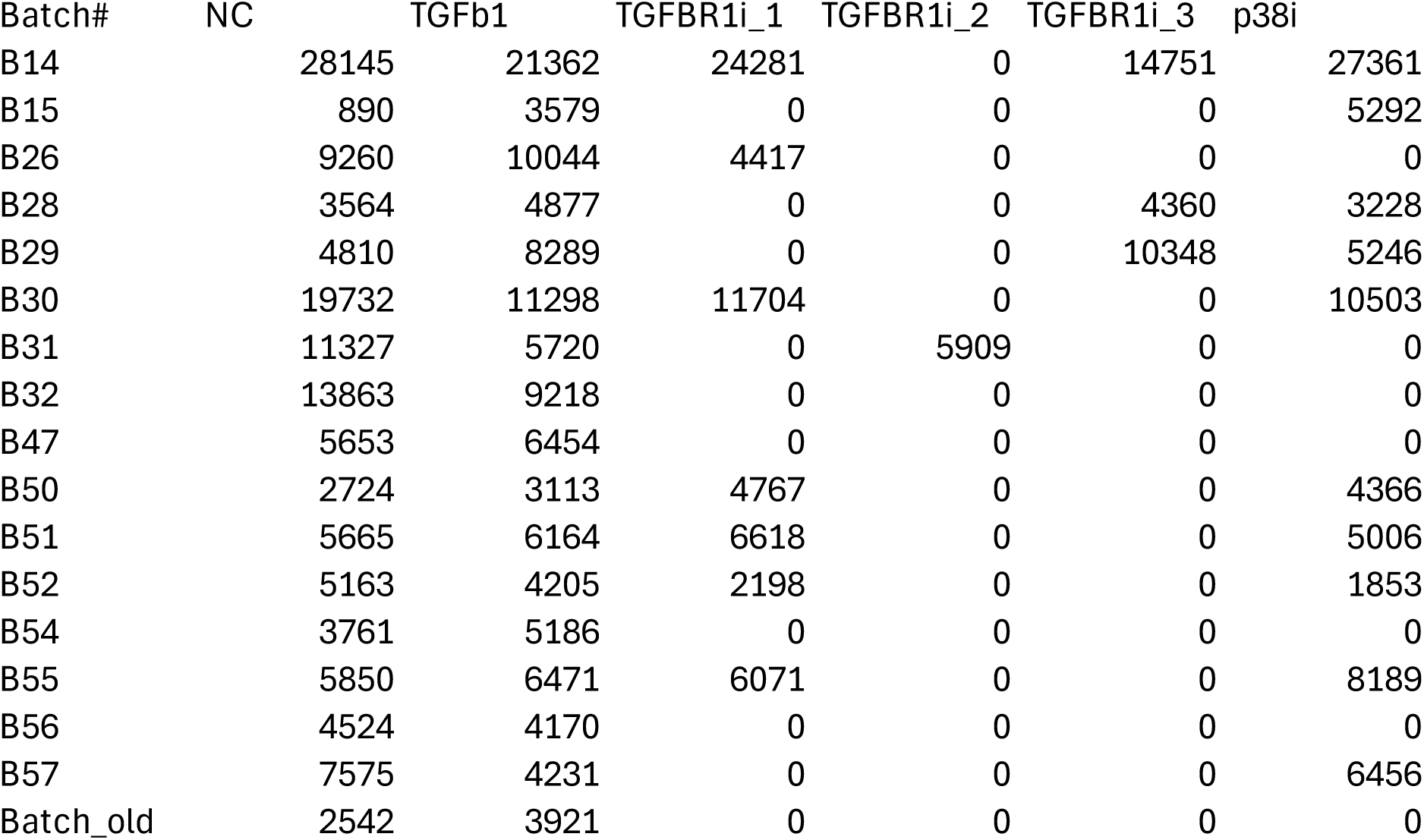
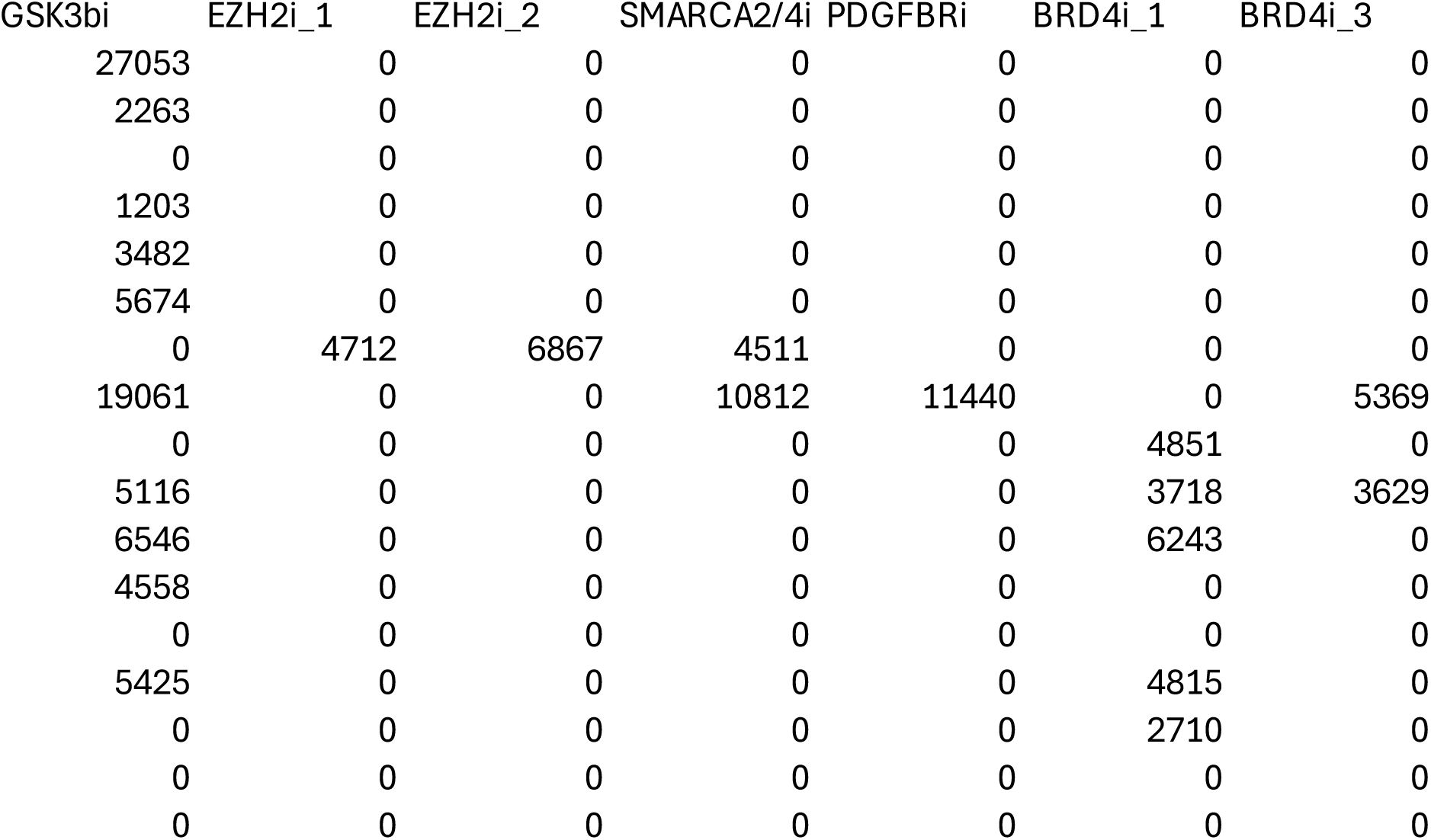
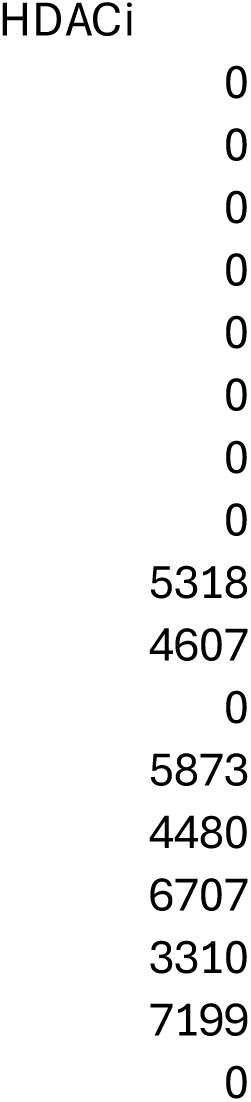

### cell# per condition per Batch

| cluster | term | df | sumsq | meansq | statistic | p.value |
|---|---|---|---|---|---|---|
| Progenitor | group | 9 | 1026.74873 | 114.083192 | 11.1960639 | 5.70364965451576e-10 |
| Progenitor | Residuals | 59 | 601.185237 | 10.1895803 | NA | NA |
| Proliferative cell | group | 9 | 63.6677939 | 7.07419933 | 4.55610555 | 0.00012572 |
| Proliferative cell | Residuals | 62 | 96.266505 | 1.55268556 | NA | NA |
| EC1 | group | 9 | 1.43214822 | 0.15912758 | 4.38383149 | 0.00018665 |
| EC1 | Residuals | 62 | 2.25052218 | 0.03629874 | NA | NA |
| EC2 | group | 9 | 52.4342633 | 5.82602925 | 3.12327128 | 0.00355007 |
| EC2 | Residuals | 64 | 119.383121 | 1.86536126 | NA | NA |
| Cho1 | group | 9 | 919.99404 | 102.22156 | 2.67042299 | 0.01069368 |
| Cho1 | Residuals | 64 | 2449.8665 | 38.2791641 | NA | NA |
| Cho2 | group | 9 | 261.041678 | 29.0046309 | 1.89325593 | 0.06988725 |
| Cho2 | Residuals | 61 | 934.518394 | 15.3199737 | NA | NA |
| Cho3 | group | 9 | 5180.1333 | 575.570367 | 84.0653012 | 5.38631725540313e-32 |
| Cho3 | Residuals | 64 | 438.189158 | 6.84670559 | NA | NA |
| EMT | group | 9 | 1090.96626 | 121.218473 | 5.50032174 | 1.45678175092213e-05 |
| EMT | Residuals | 63 | 1388.42129 | 22.0384331 | NA | NA |
| Hepatocyte | group | 9 | 2013.1956 | 223.6884 | 37.8631882 | 1.06312174044478e-21 |
| Hepatocyte | Residuals | 61 | 360.376213 | 5.90780678 | NA | NA |
| HSC | group | 9 | 5144.01344 | 571.557049 | 32.3960741 | 1.89261969615749e-19 |
| HSC | Residuals | 58 | 1023.28167 | 17.6427874 | NA | NA |
| MyoF | group | 9 | 9412.86265 | 1045.87363 | 38.814053 | 2.17895866510121e-22 |
| MyoF | Residuals | 63 | 1697.58202 | 26.9457464 | NA | NA |
| Mes1 | group | 9 | 5840.70449 | 648.967165 | 30.3395401 | 1.75912536854279e-19 |
| Mes1 | Residuals | 62 | 1326.189 | 21.3901451 | NA | NA |
| Mes2 | group | 9 | 7827.77927 | 869.753252 | 310.997507 | 7.26837628810936e-50 |
| Mes2 | Residuals | 65 | 181.782683 | 2.79665666 | NA | NA |
| Mes3 | group | 9 | 575.332112 | 63.9257902 | 2.83578274 | 0.00724387 |
| Mes3 | Residuals | 63 | 1420.18101 | 22.5425556 | NA | NA |
| Mes4 | group | 9 | 11.3891323 | 1.26545914 | 3.33212502 | 0.00217904 |
| Mes4 | Residuals | 63 | 23.9258508 | 0.37977541 | NA | NA |
| Mesothelia | group | 9 | 823.744204 | 91.5271338 | 10.2399883 | 2.00831790696435e-09 |
| Mesothelia | Residuals | 61 | 545.230621 | 8.9382069 | NA | NA |

### dunnet_stats for 16 cell clusters

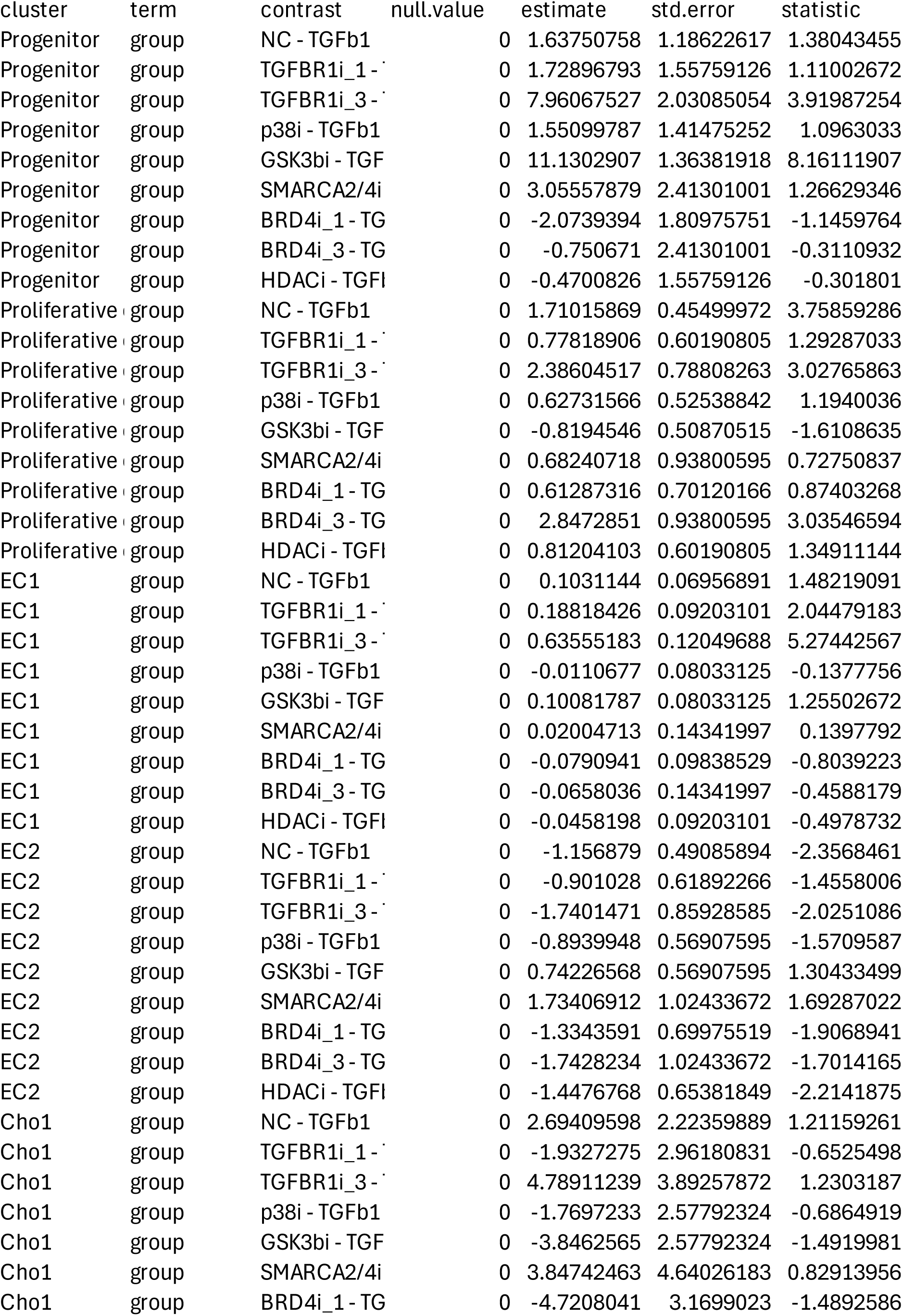
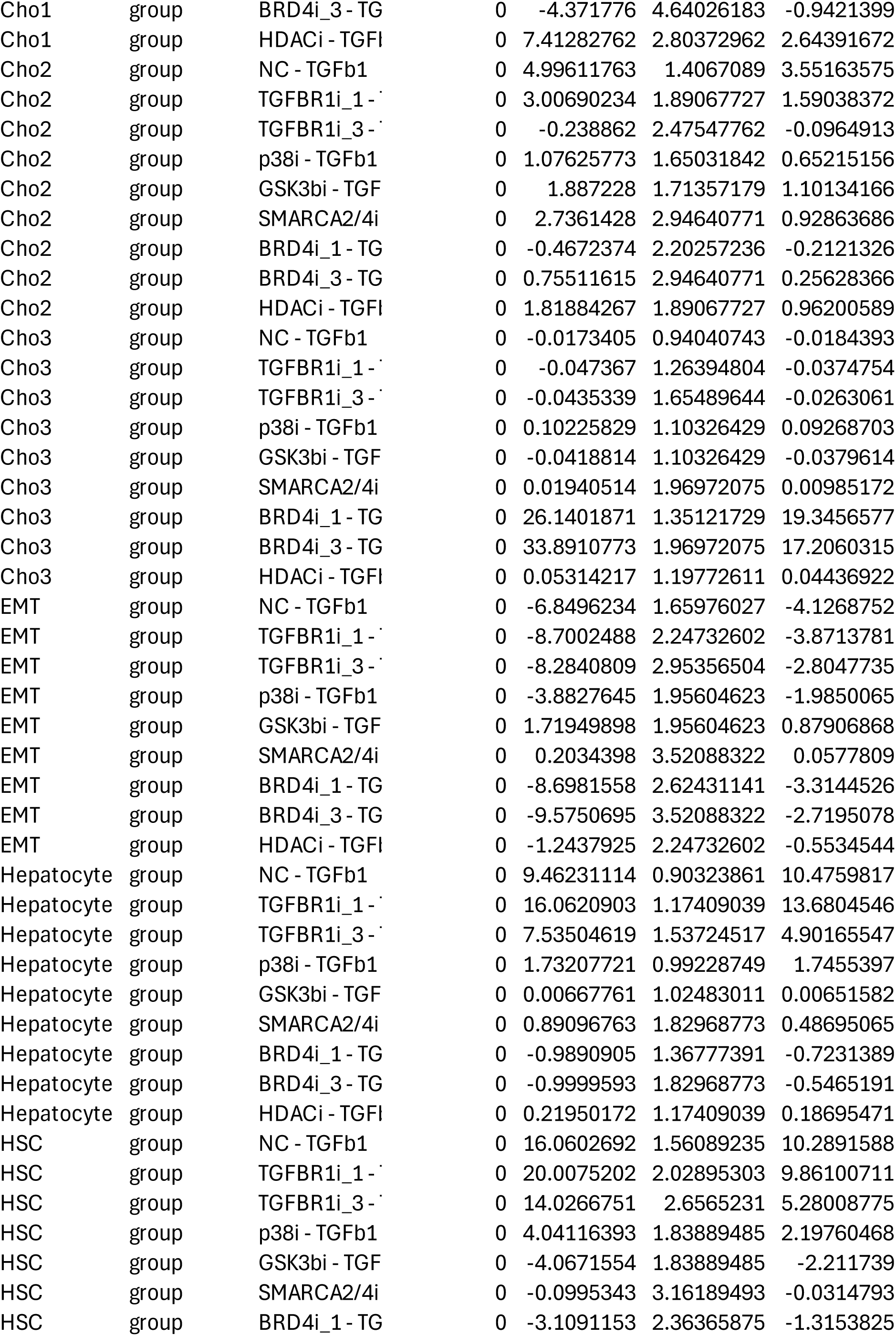
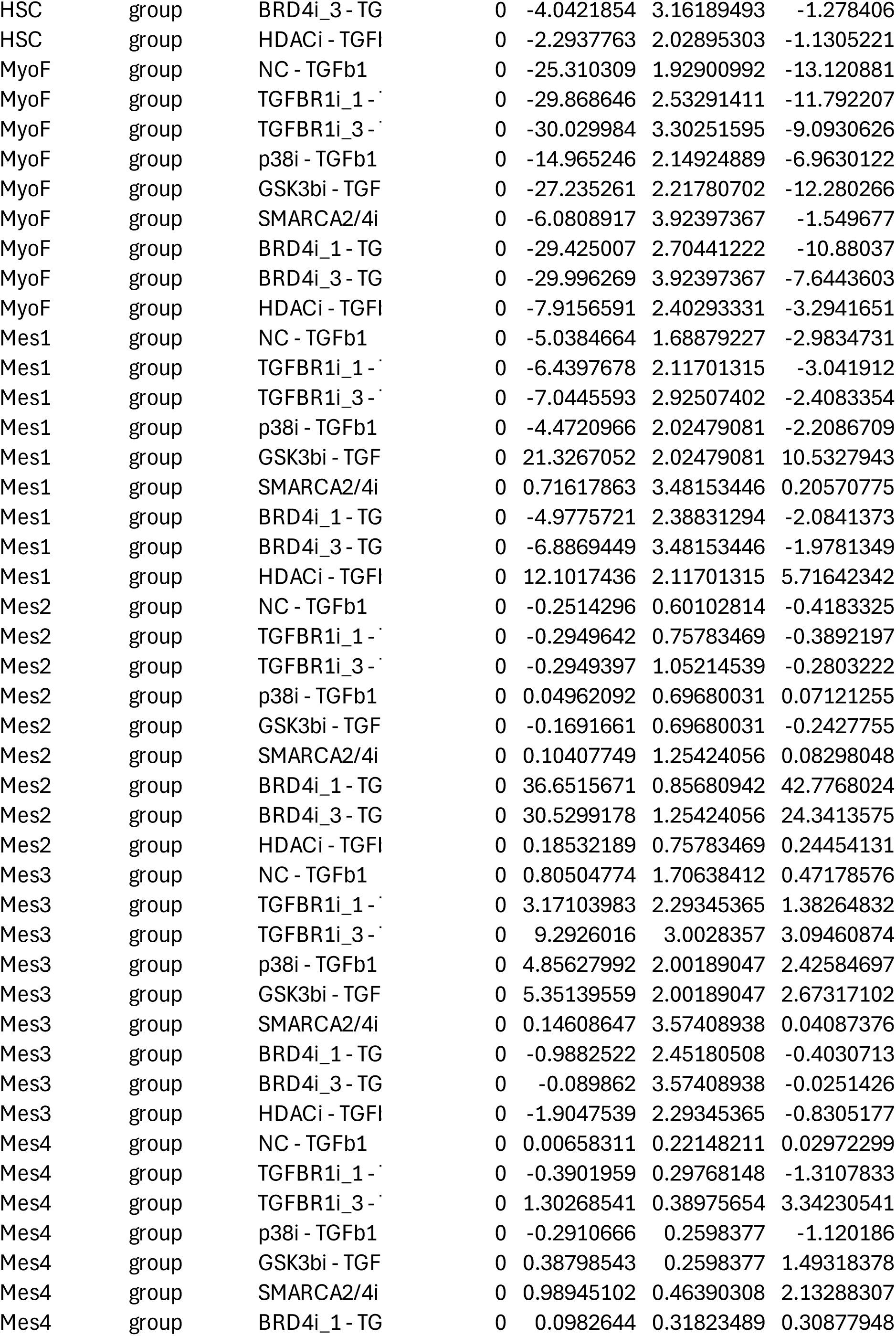
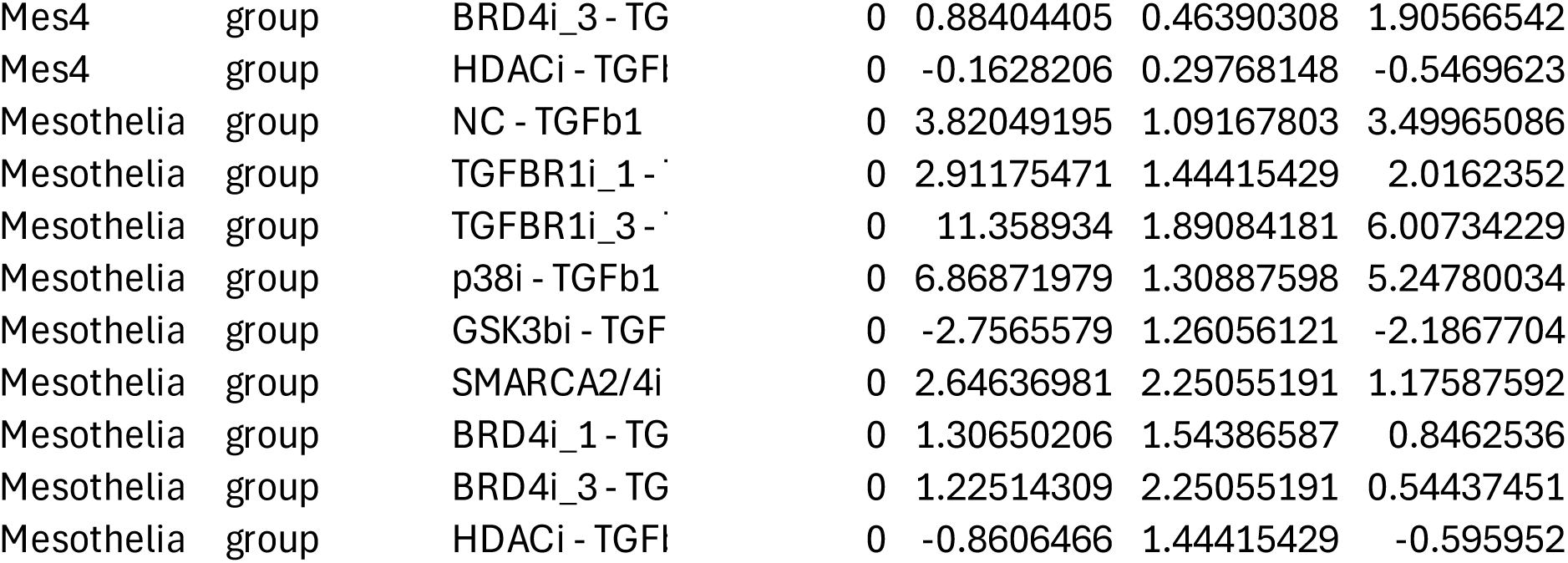
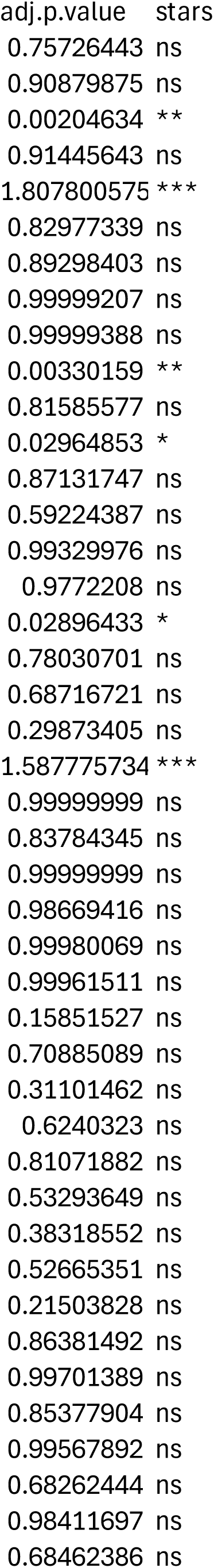
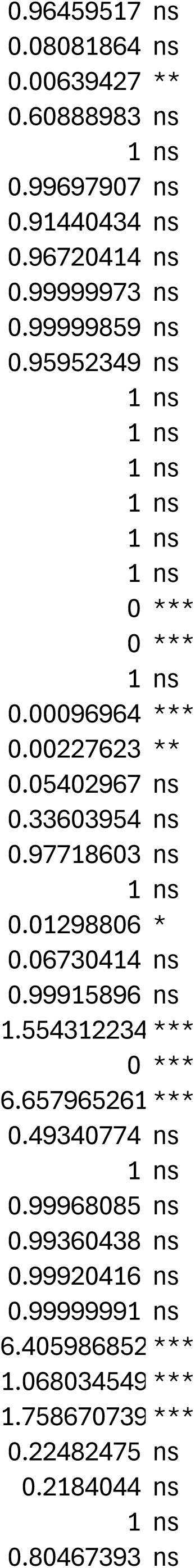
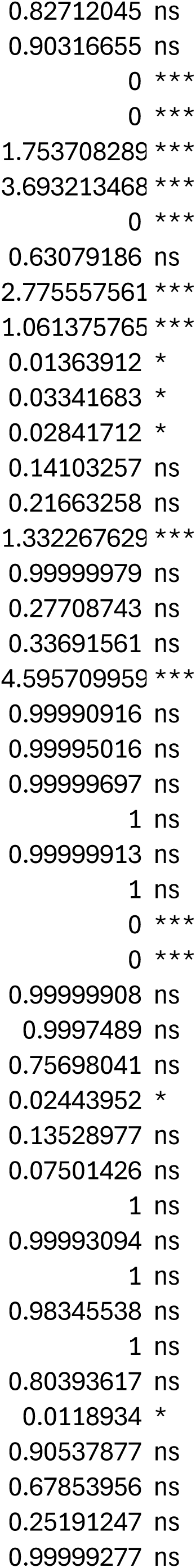
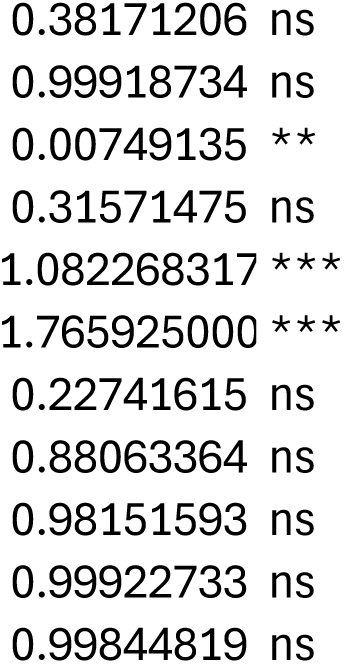
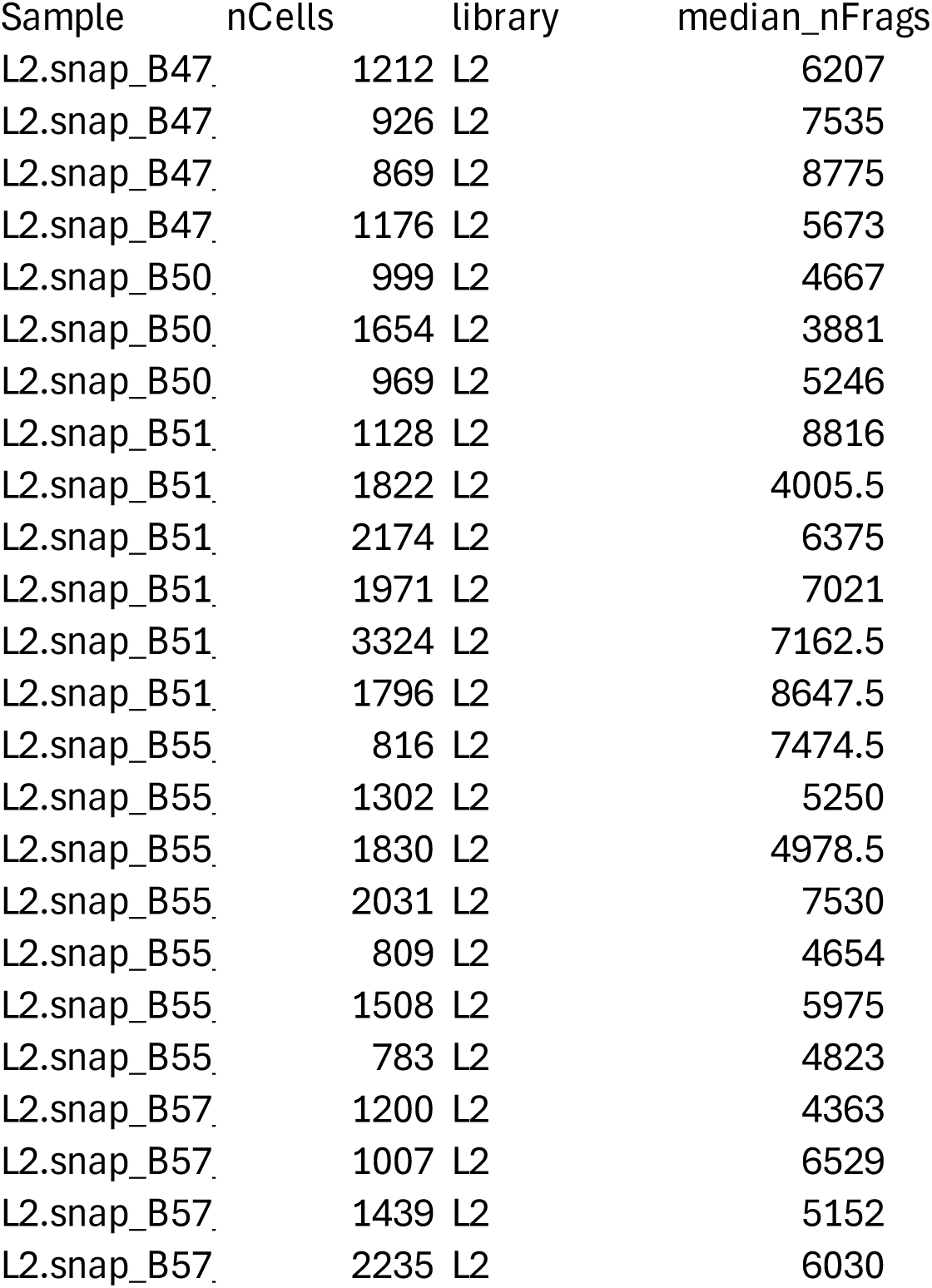
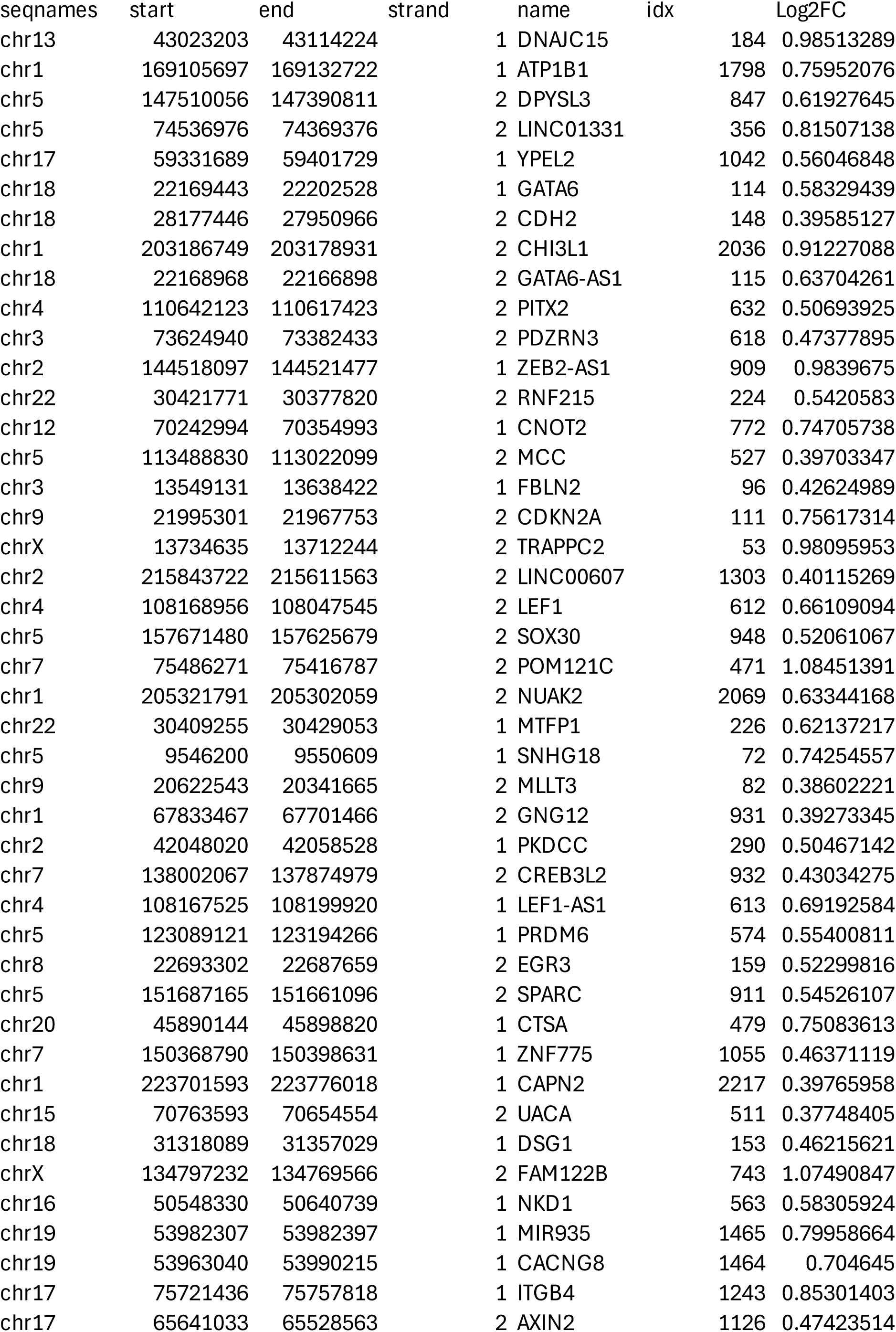
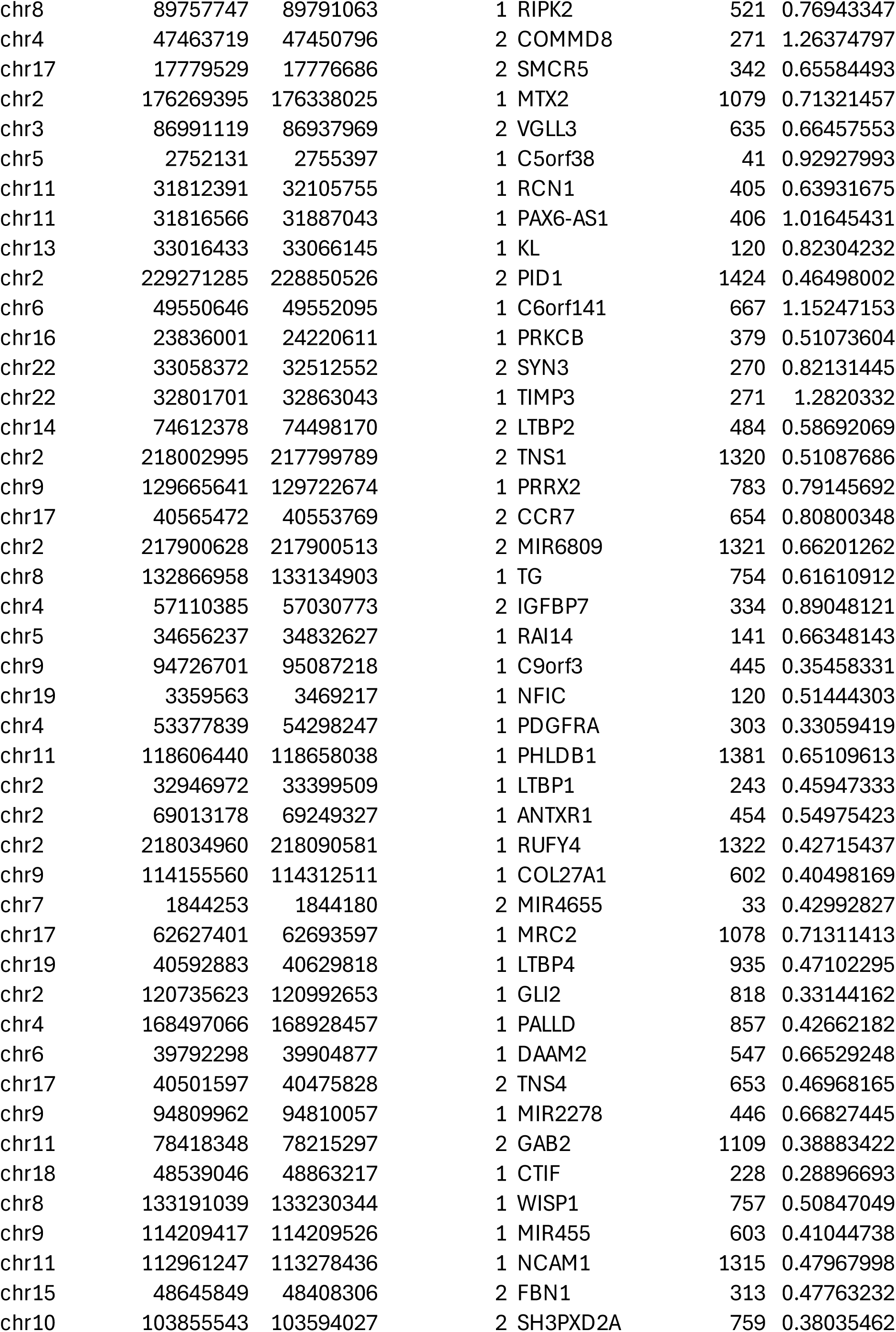
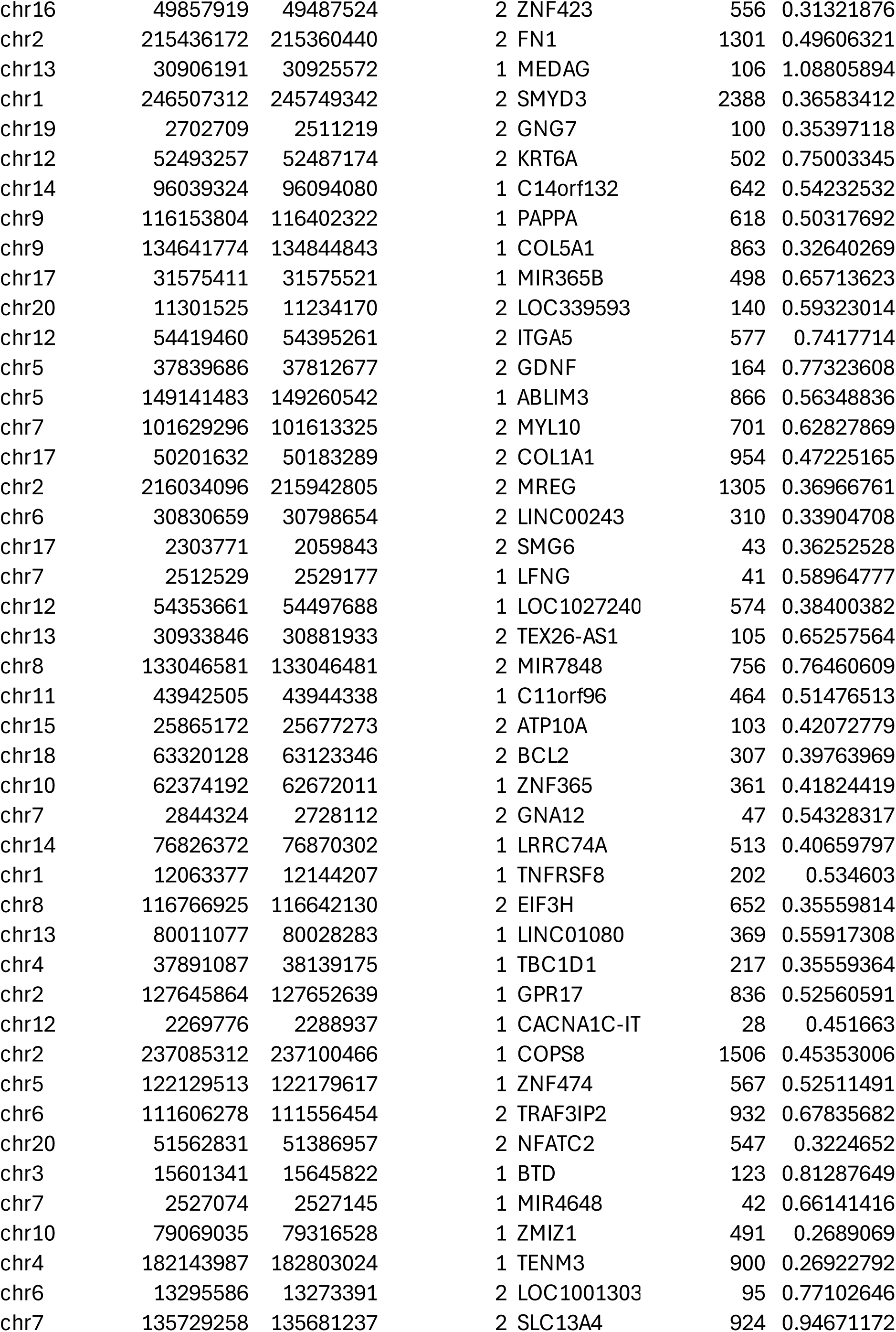
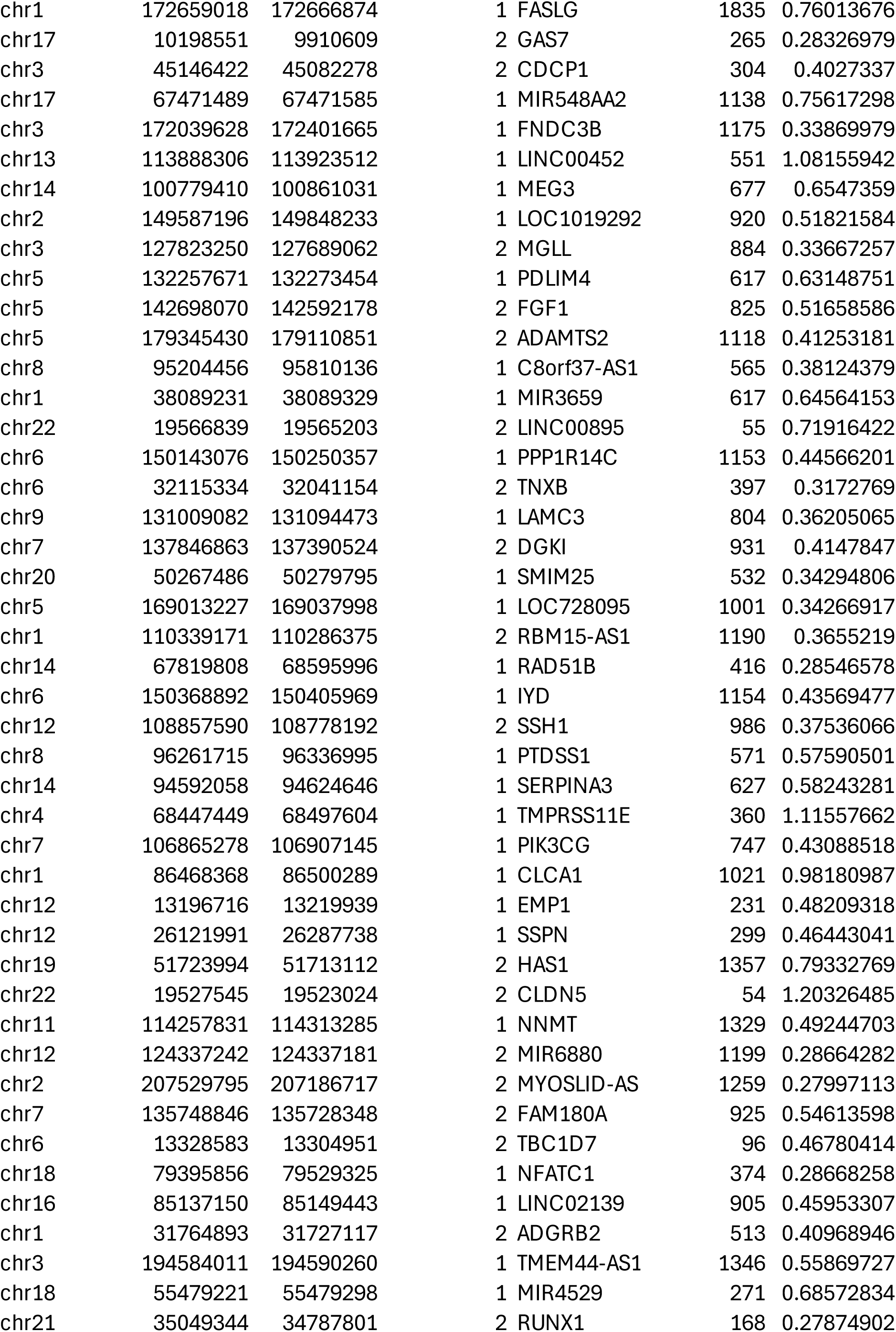
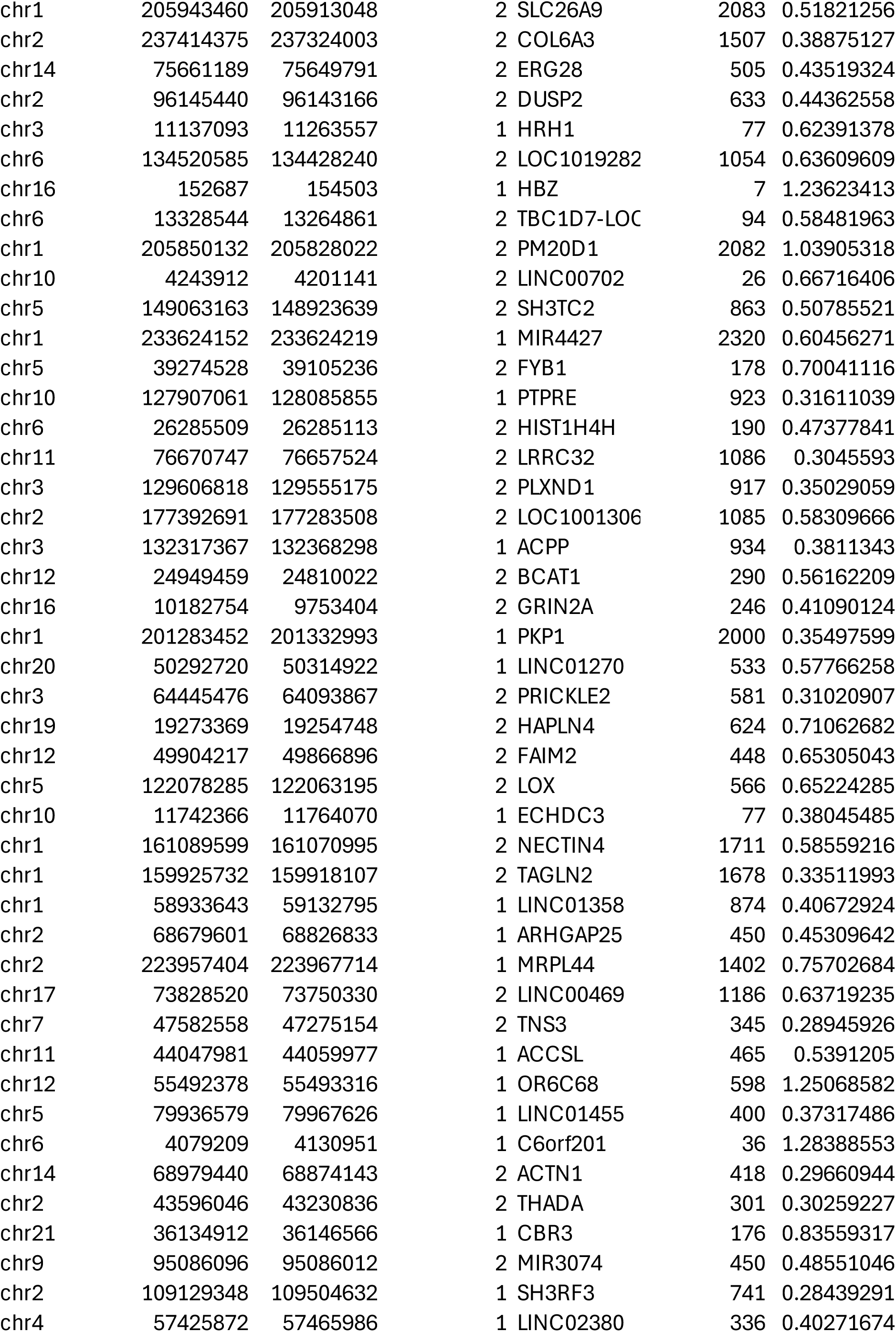
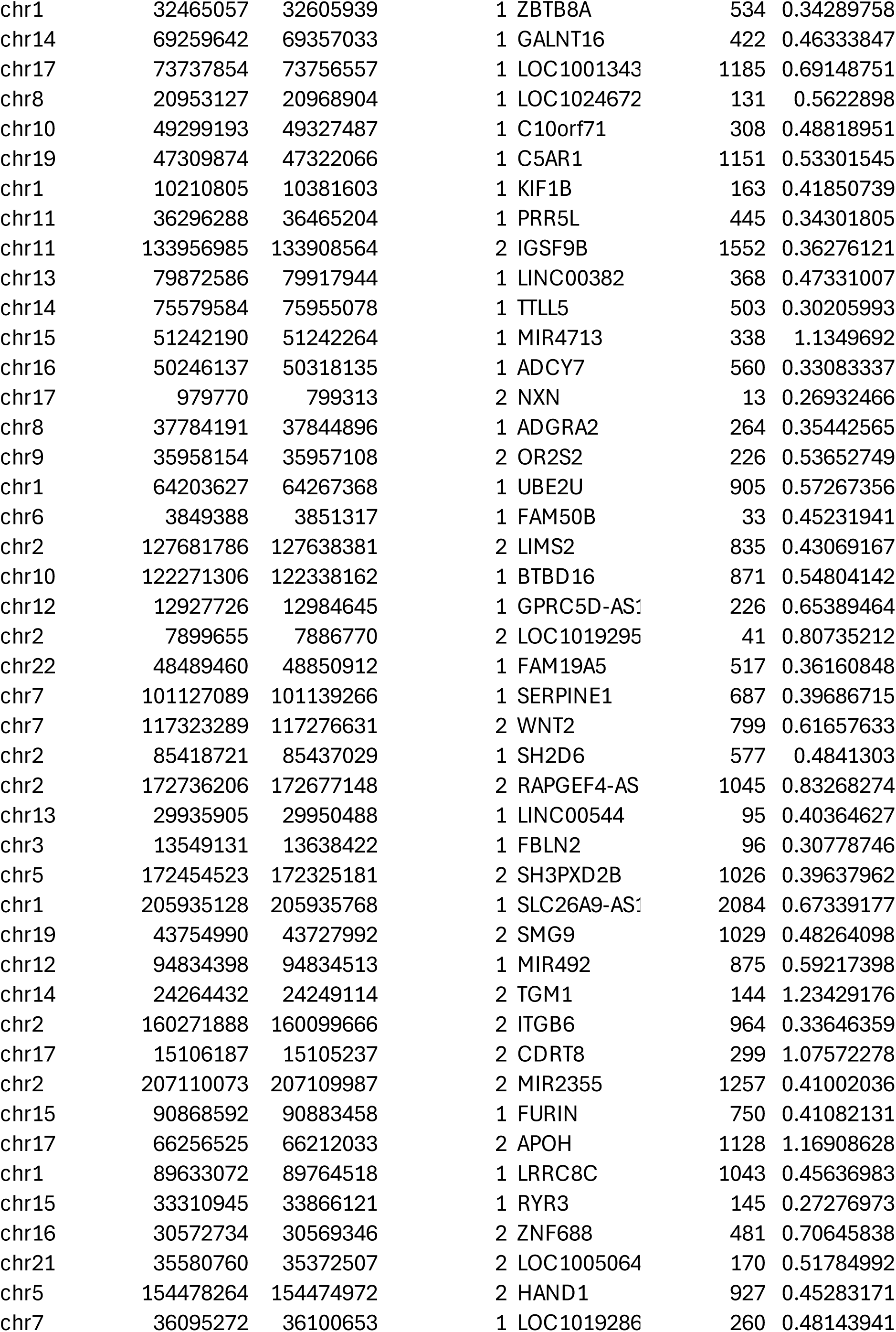
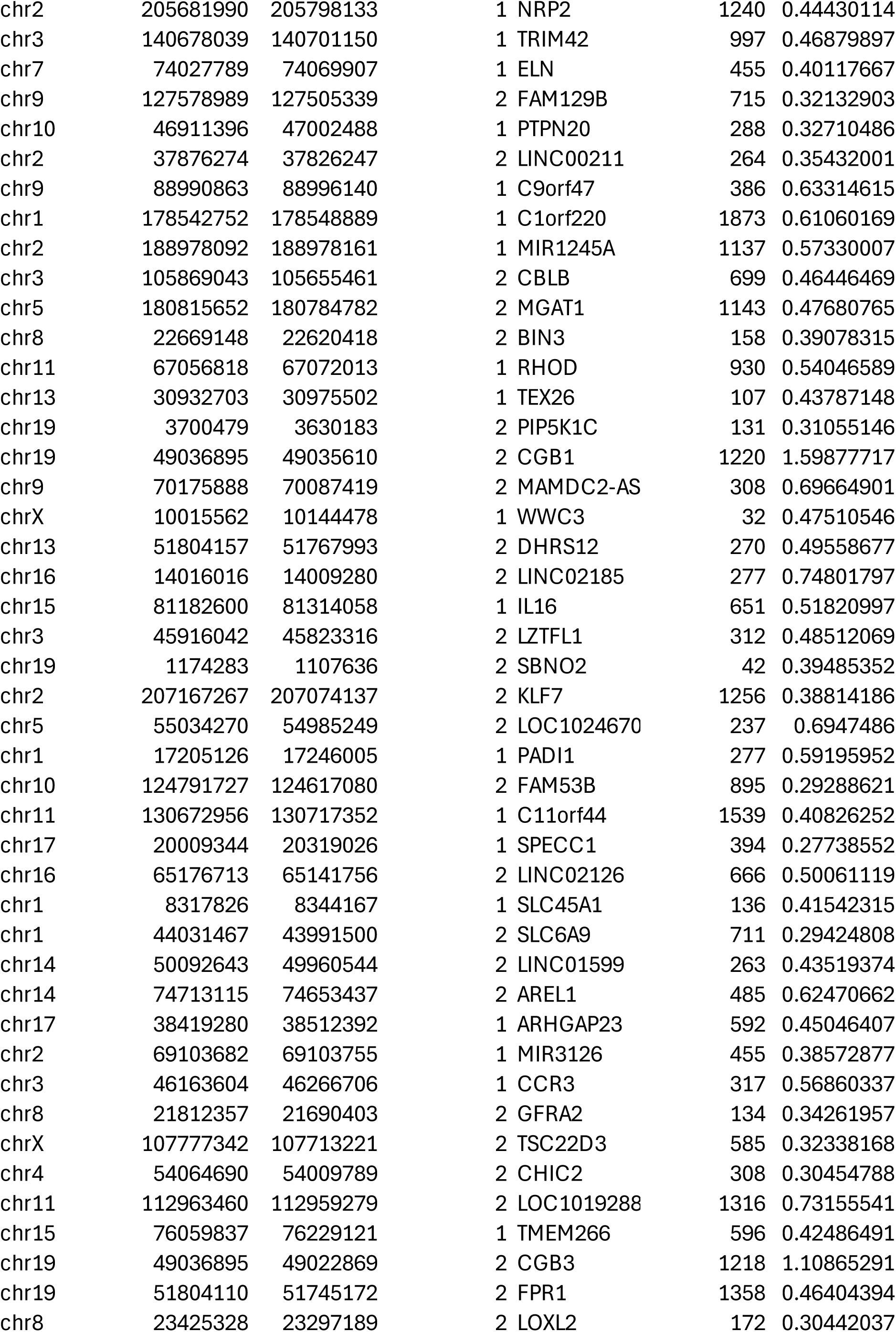
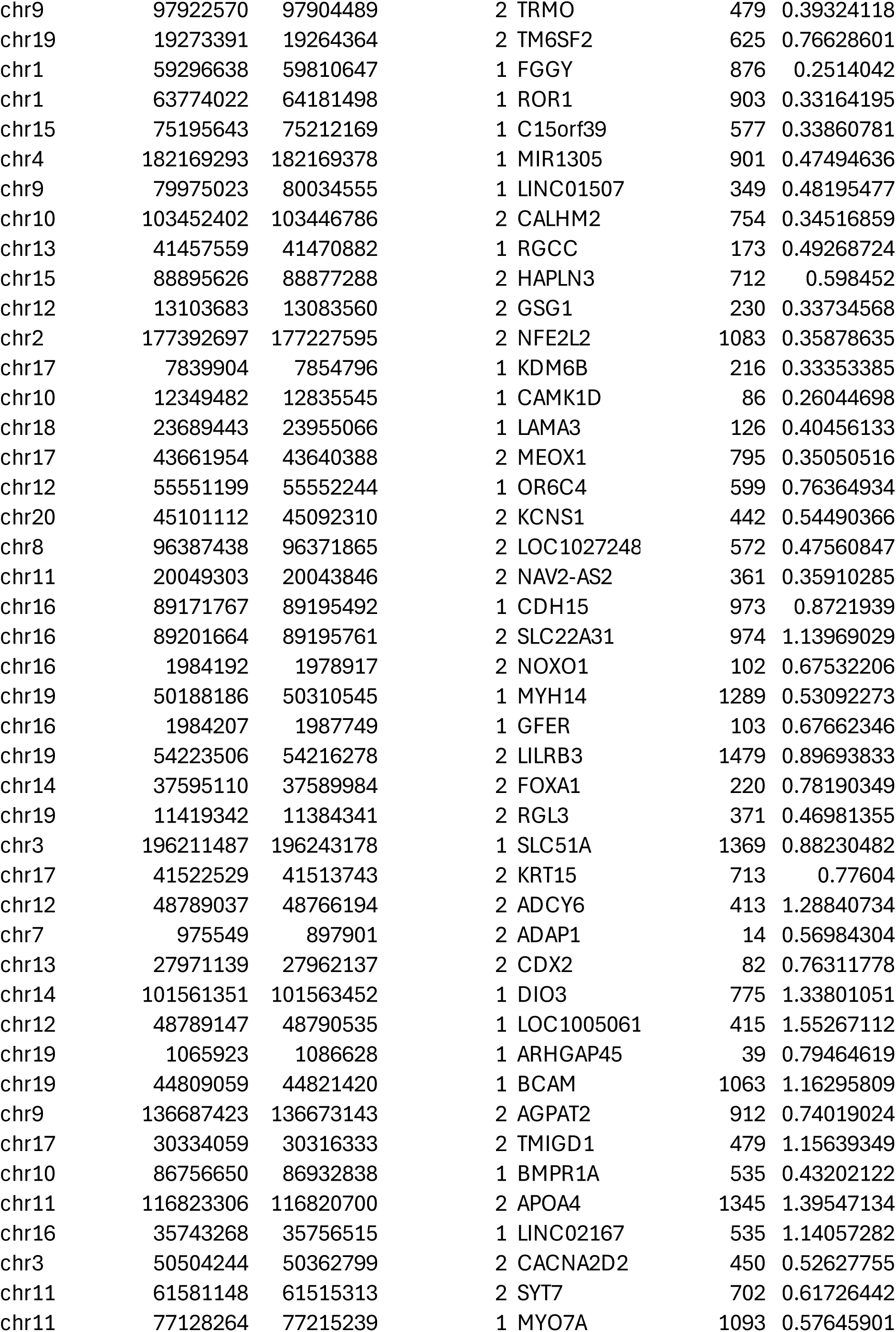
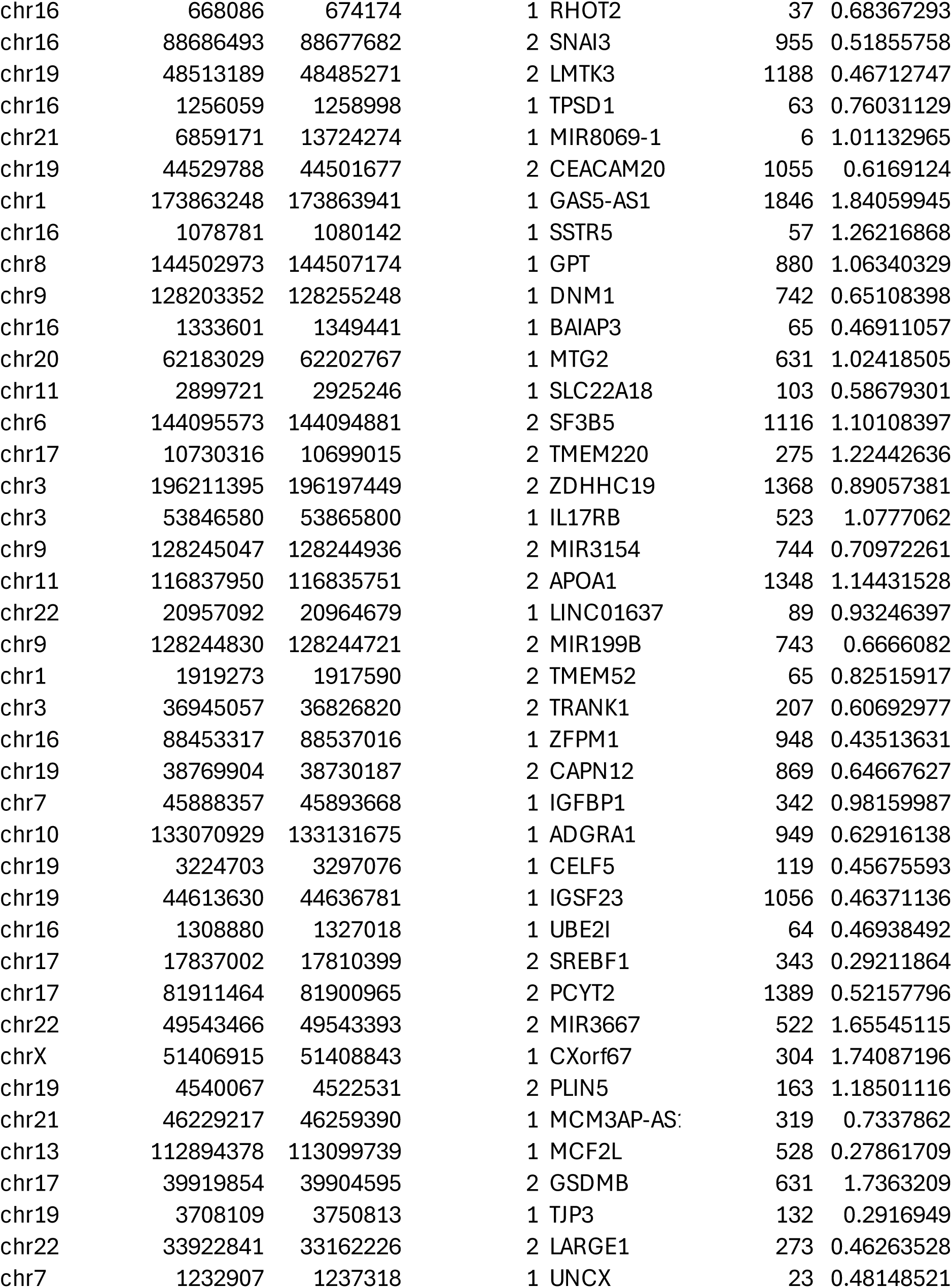
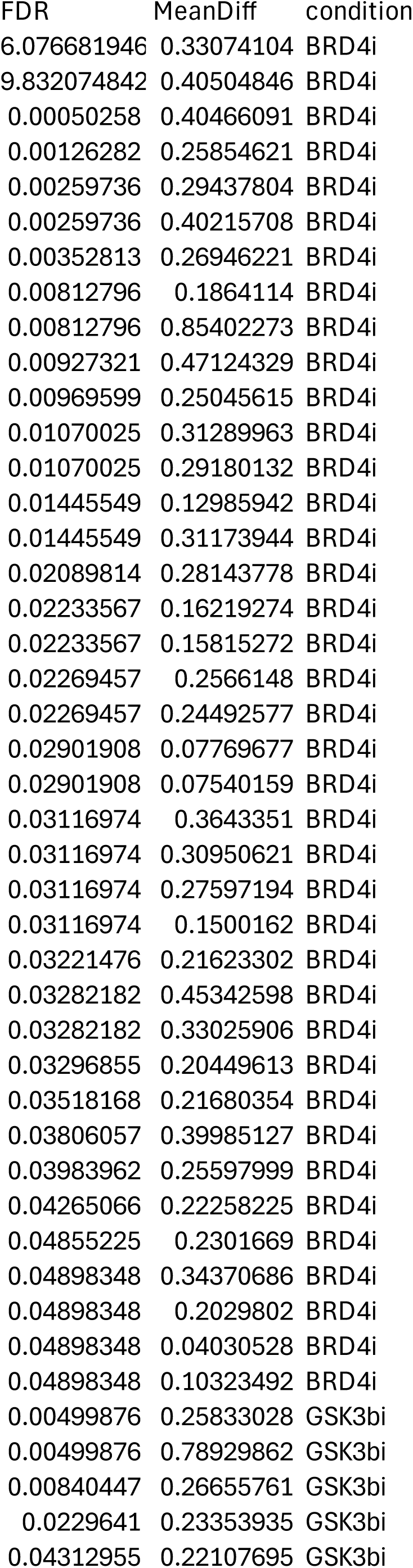
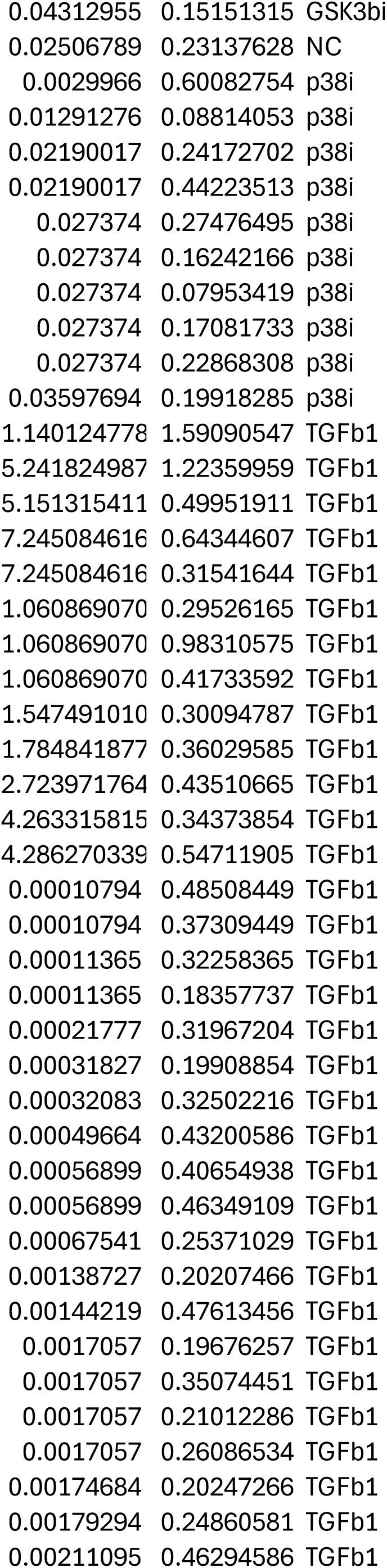
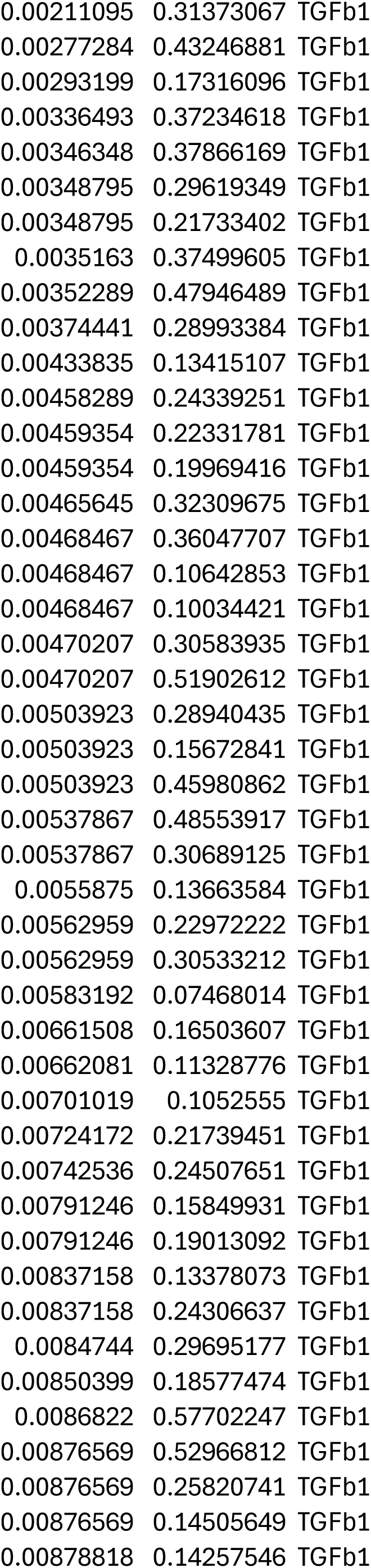
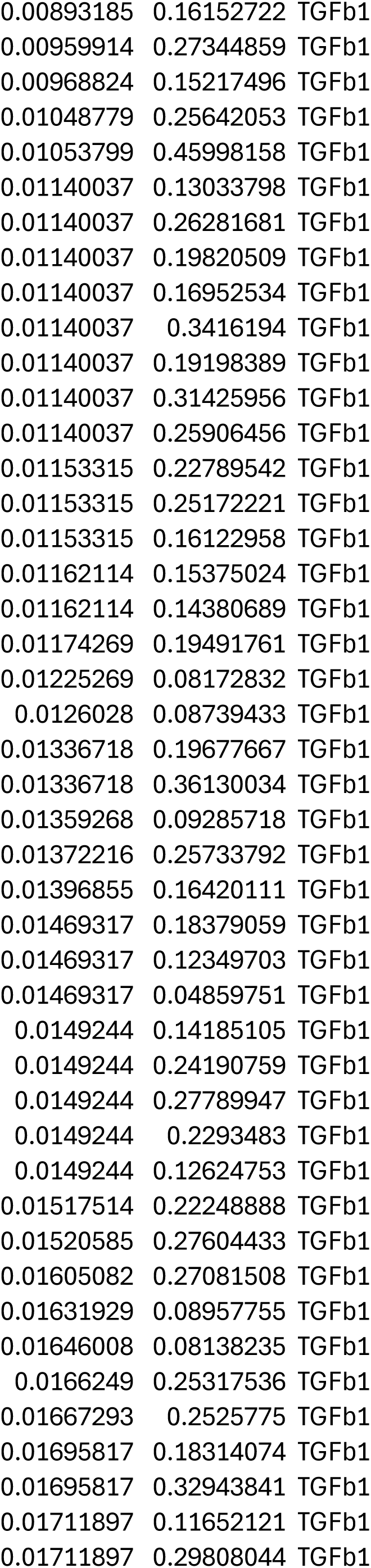
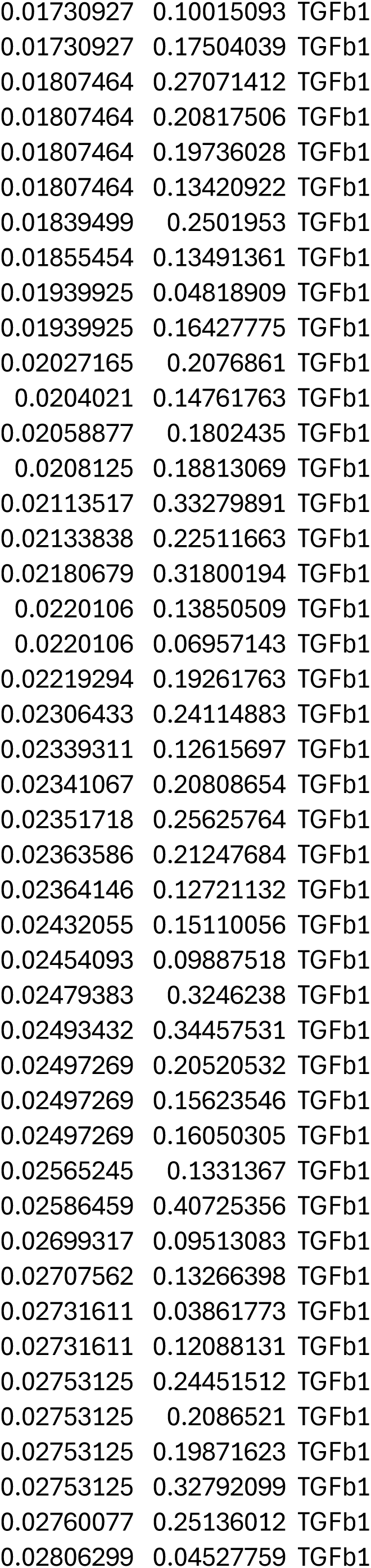
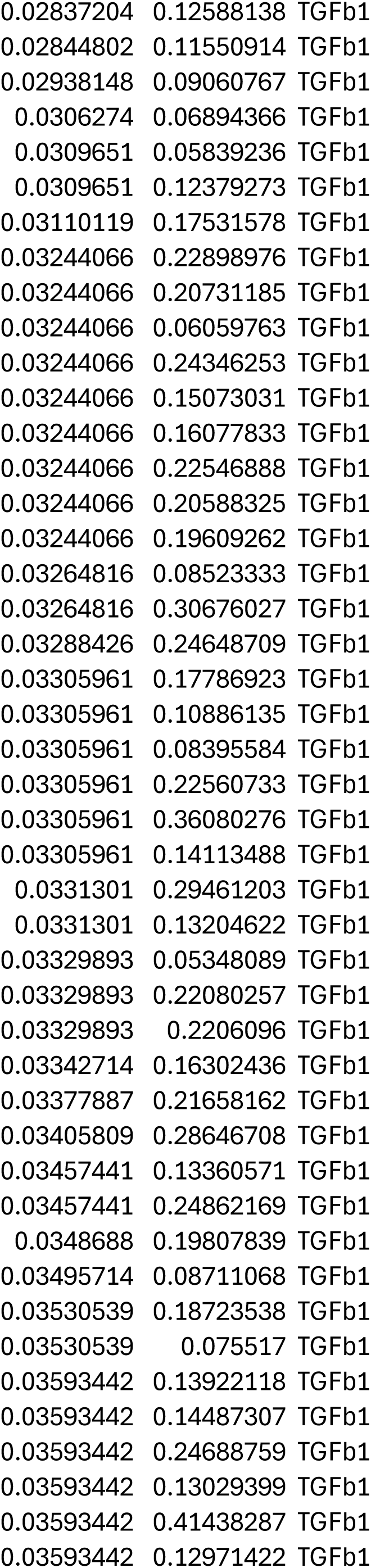
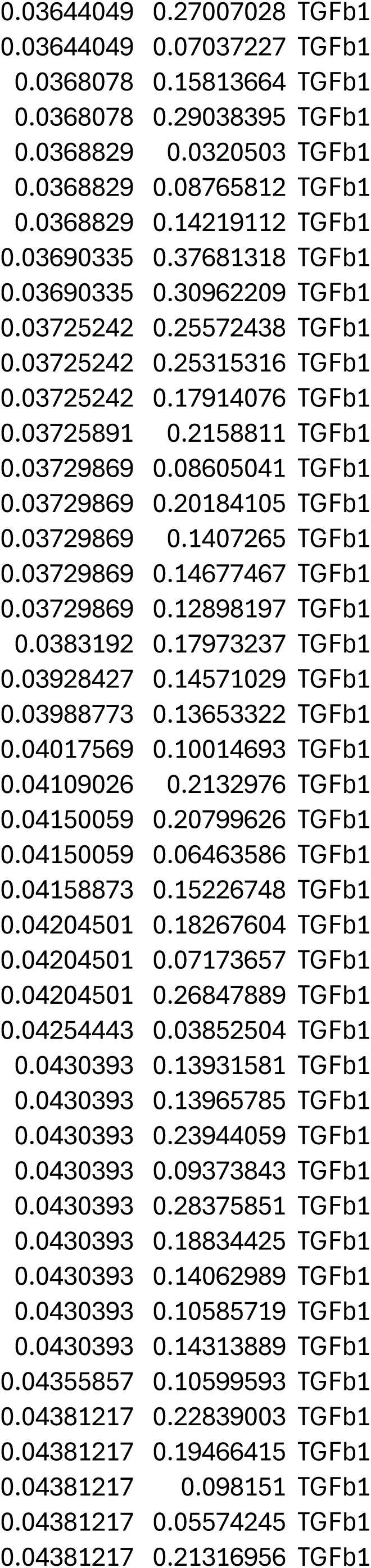
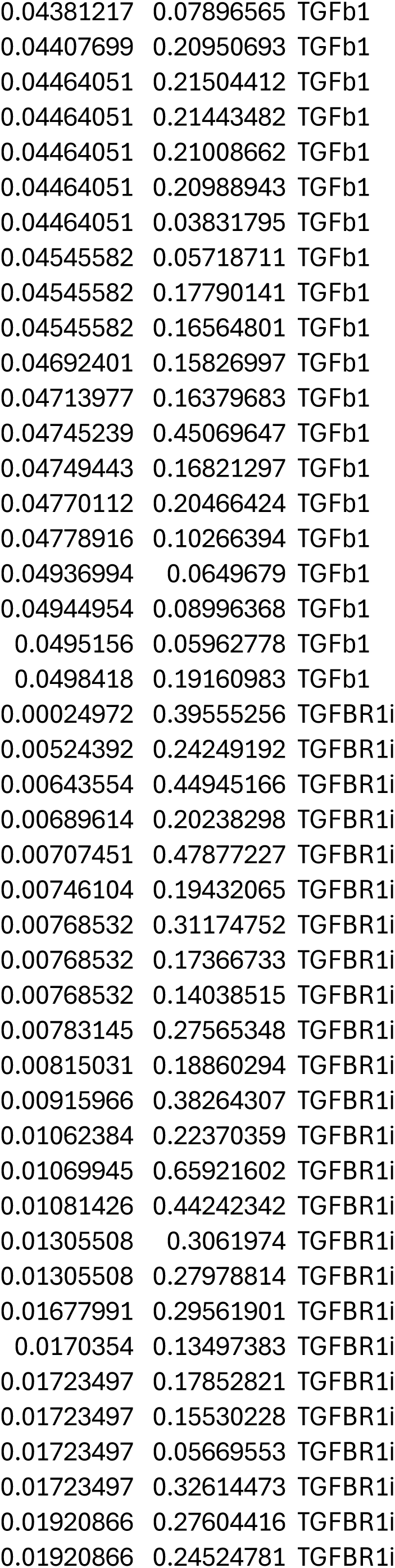
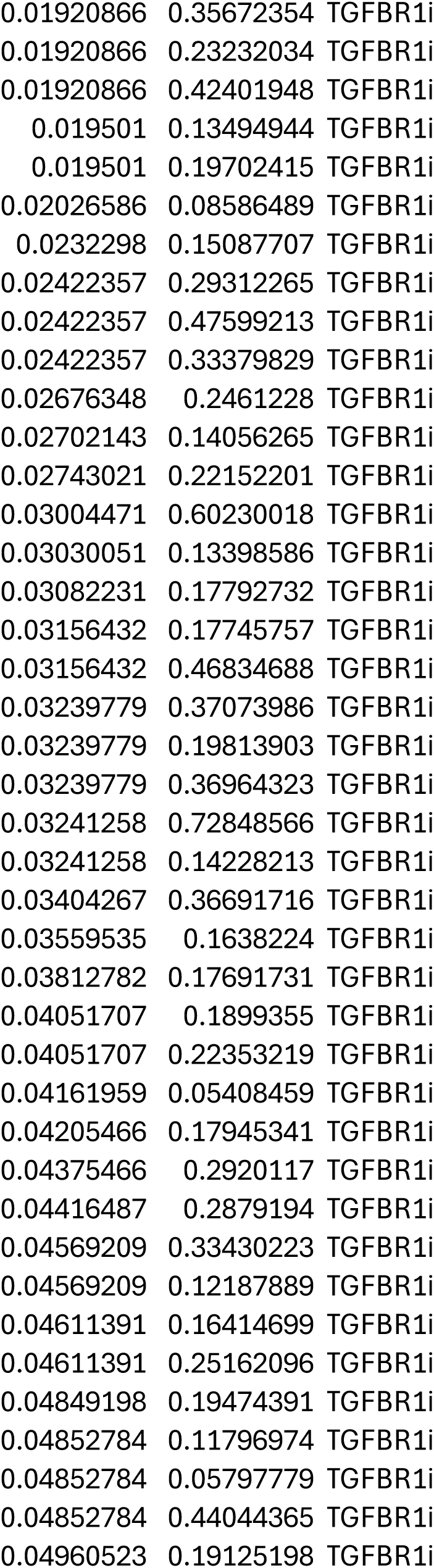
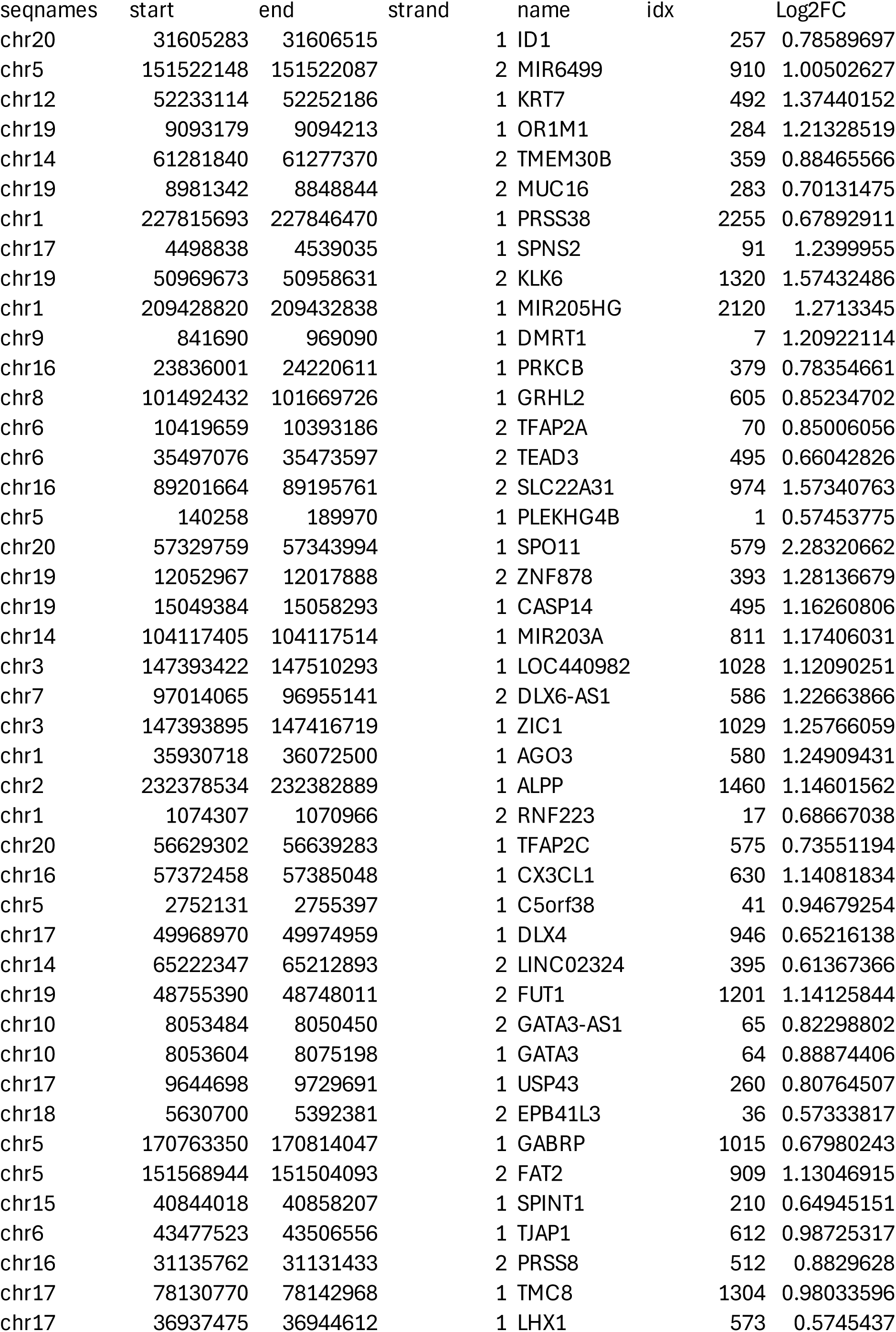
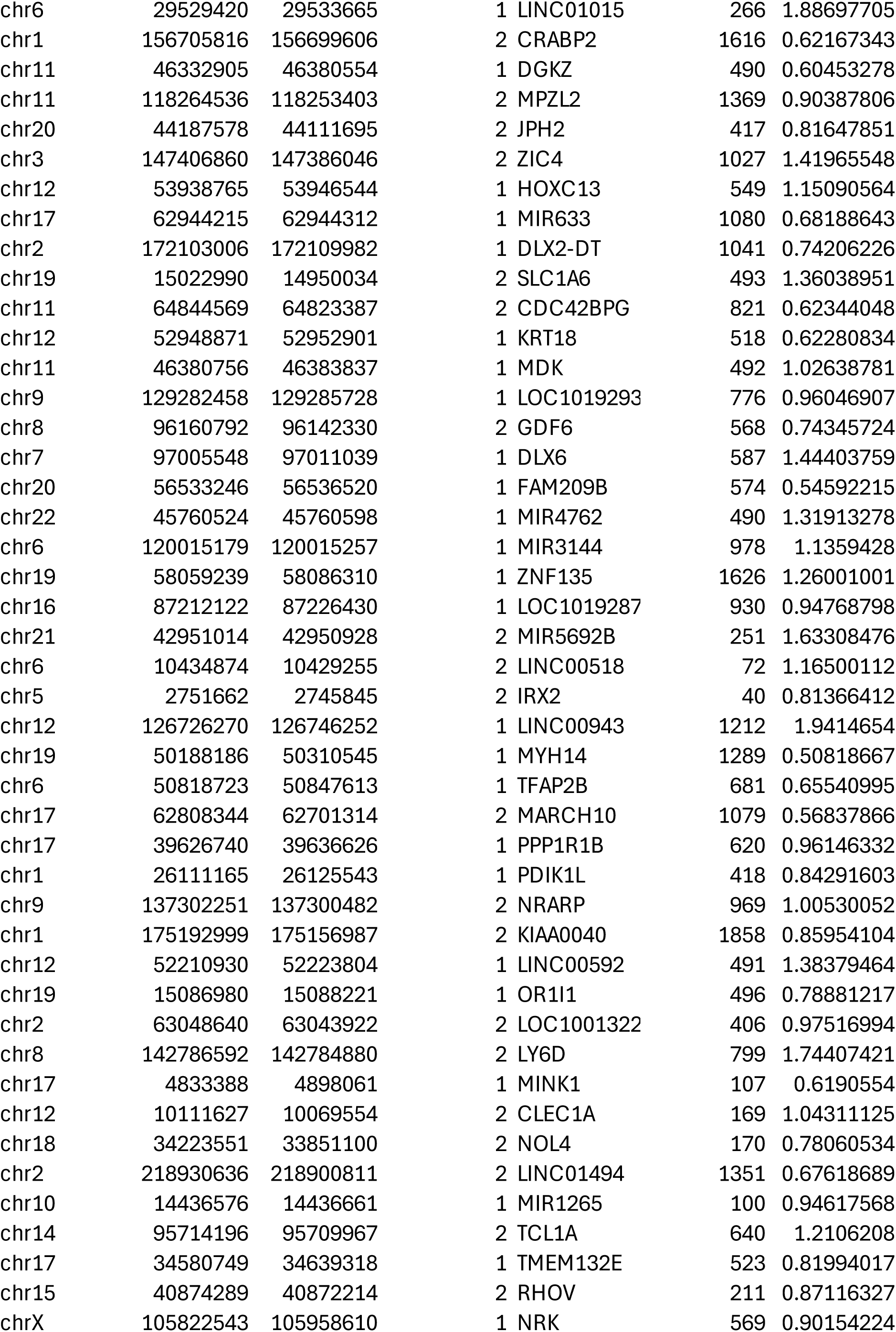
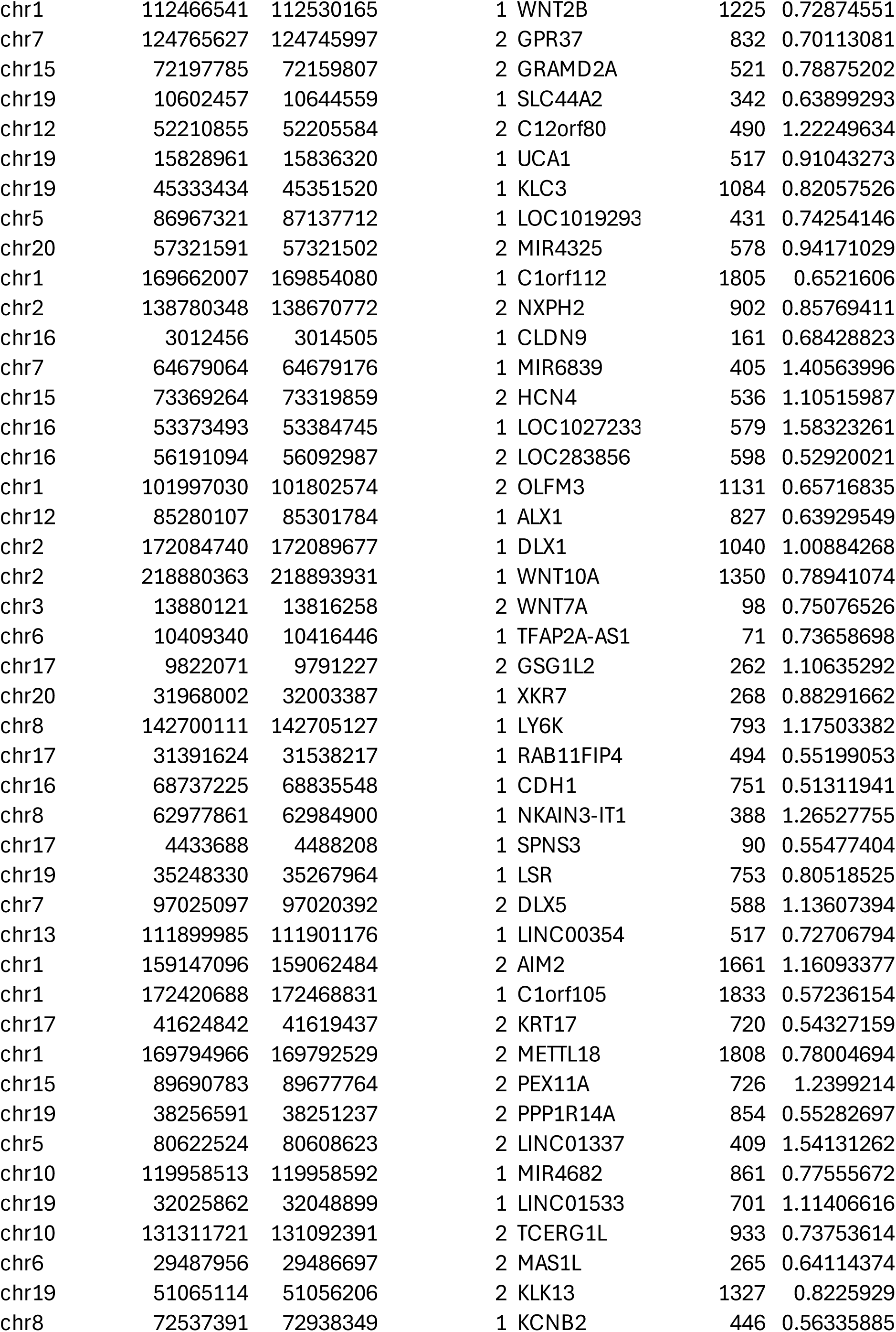
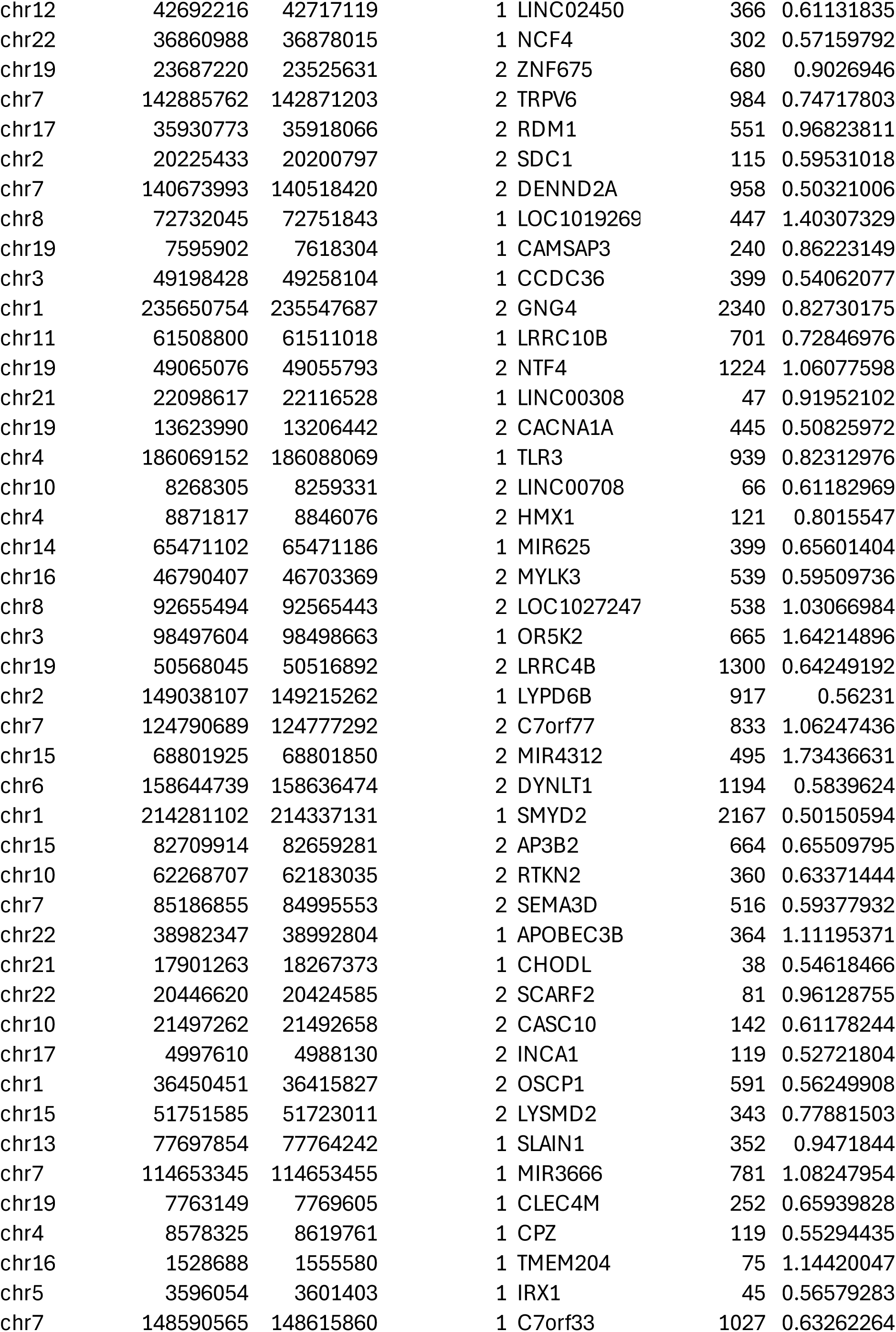
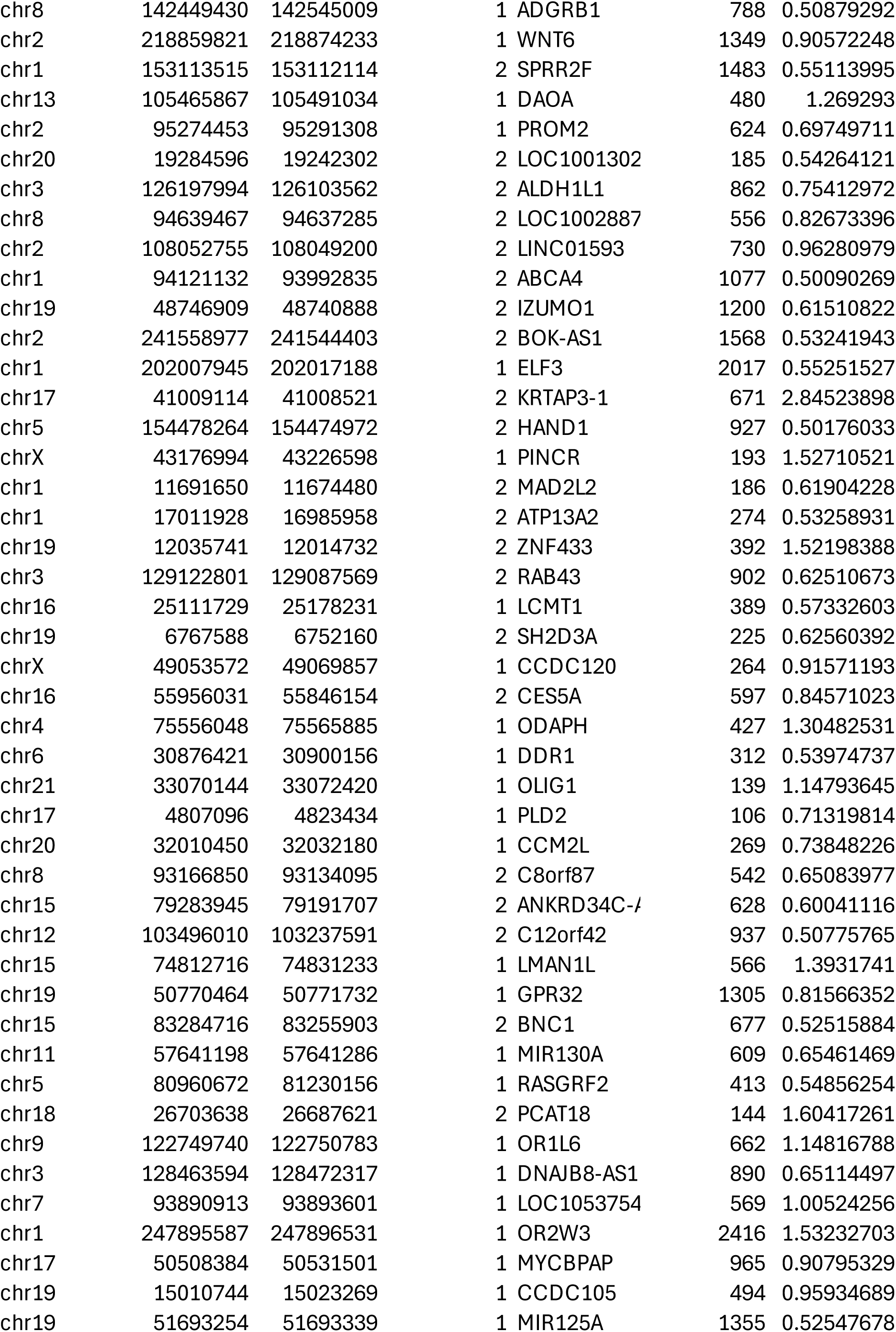
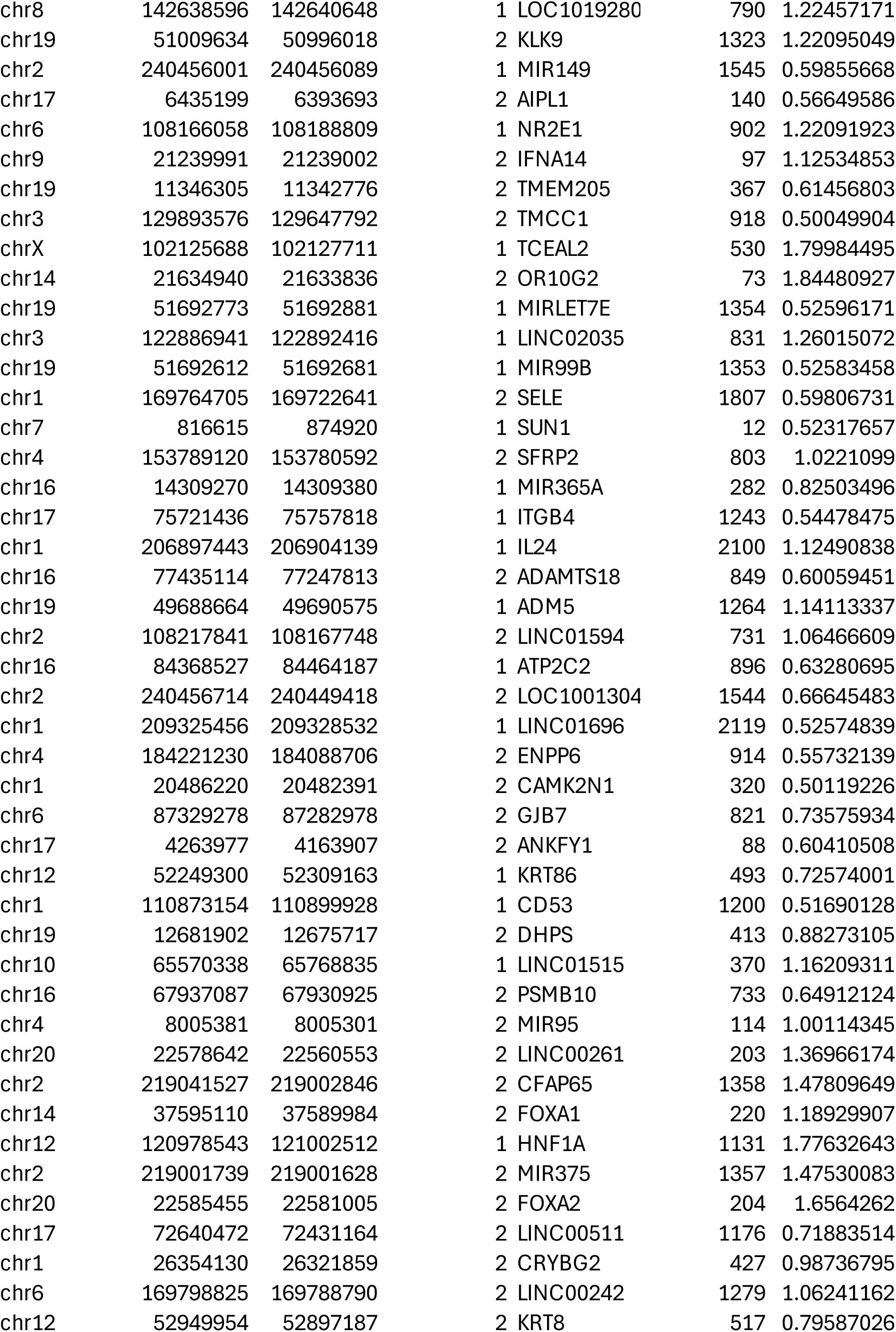
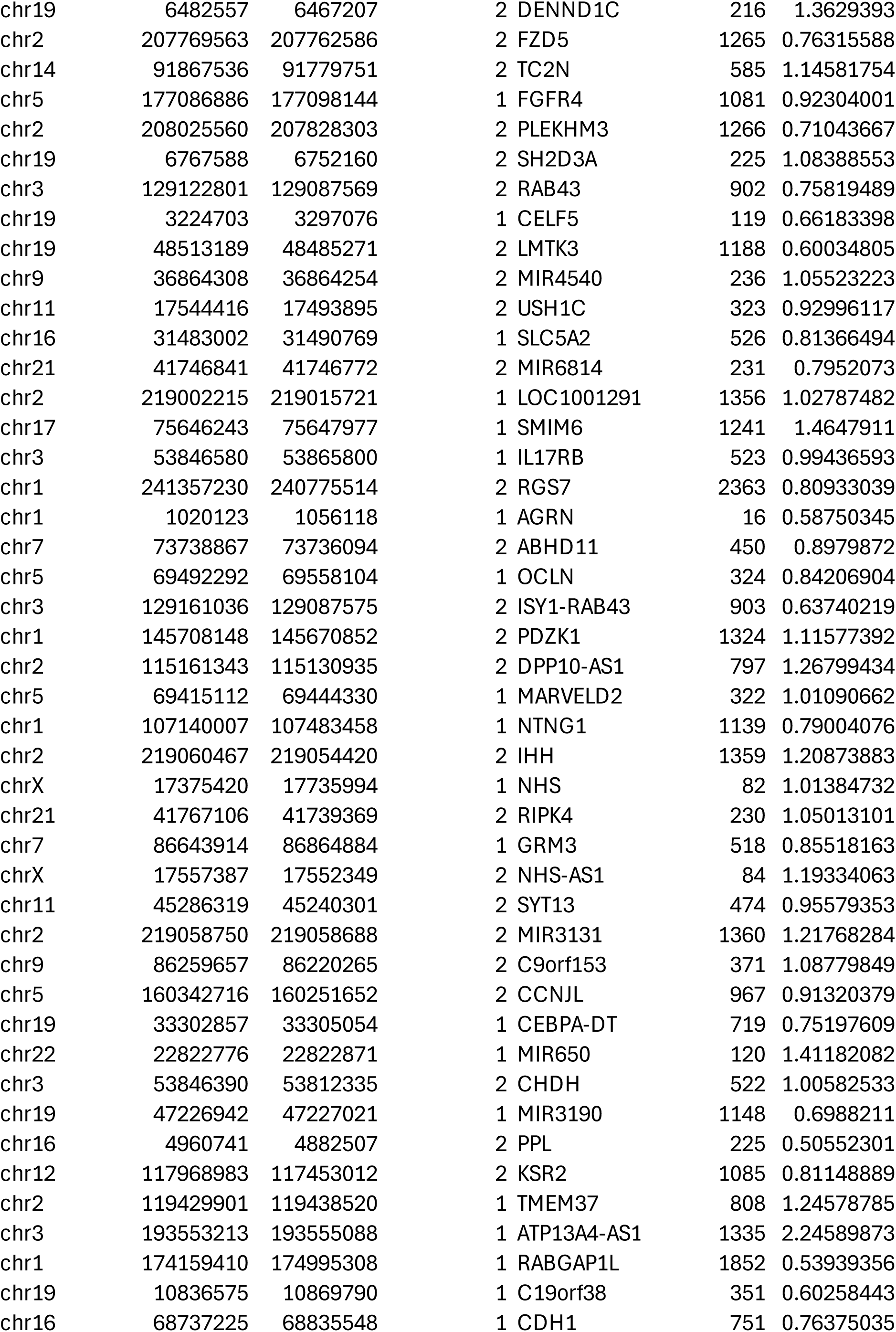
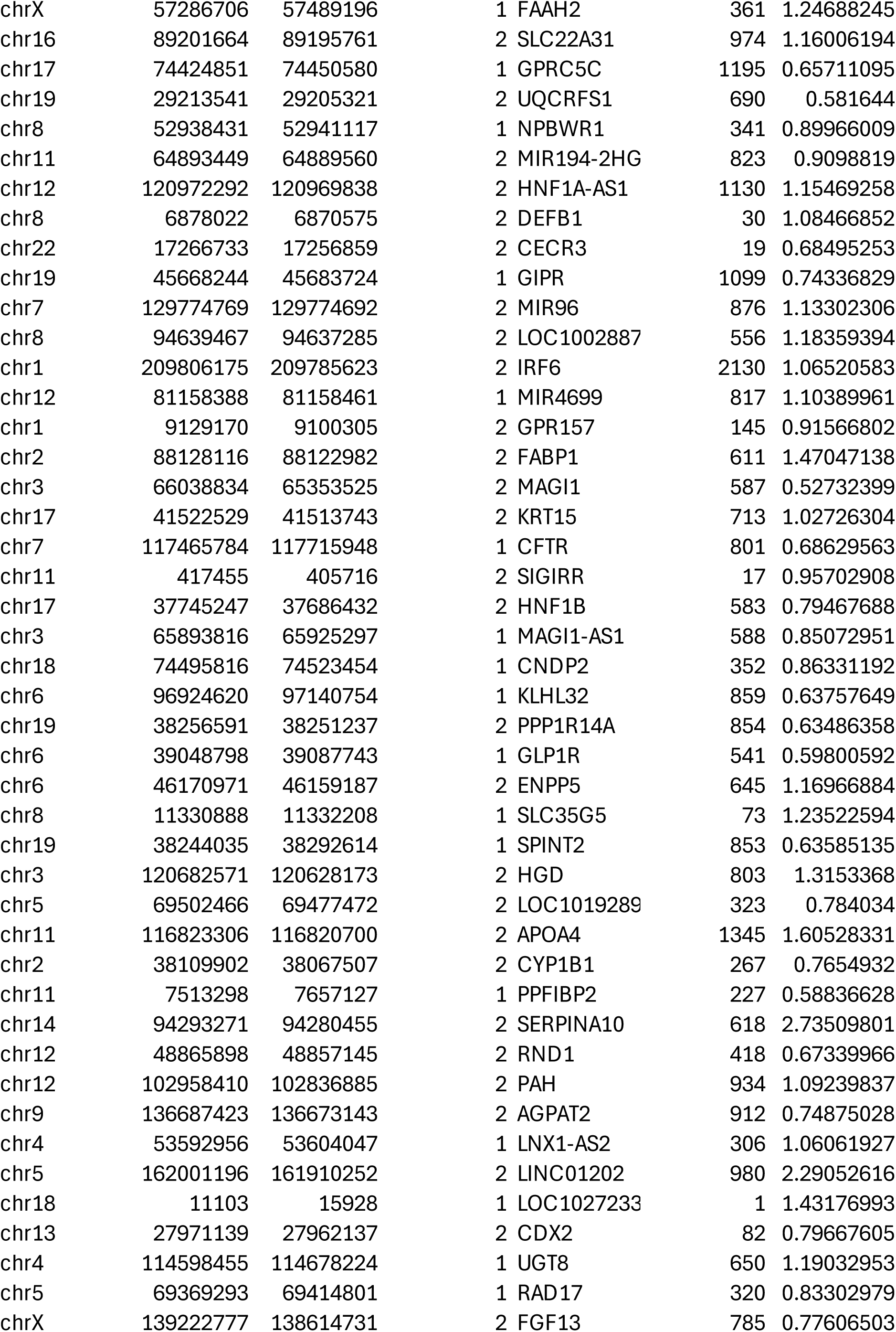
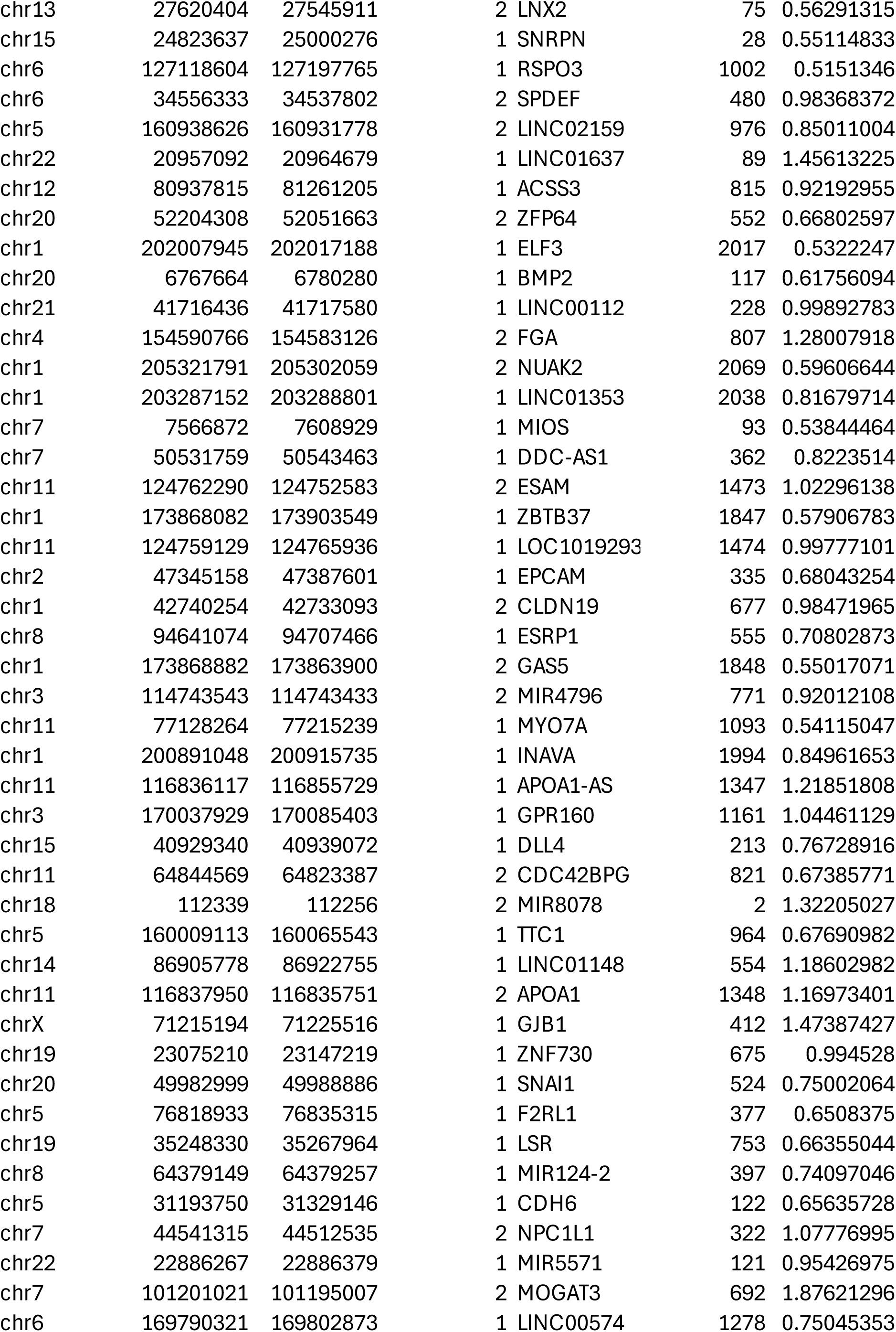
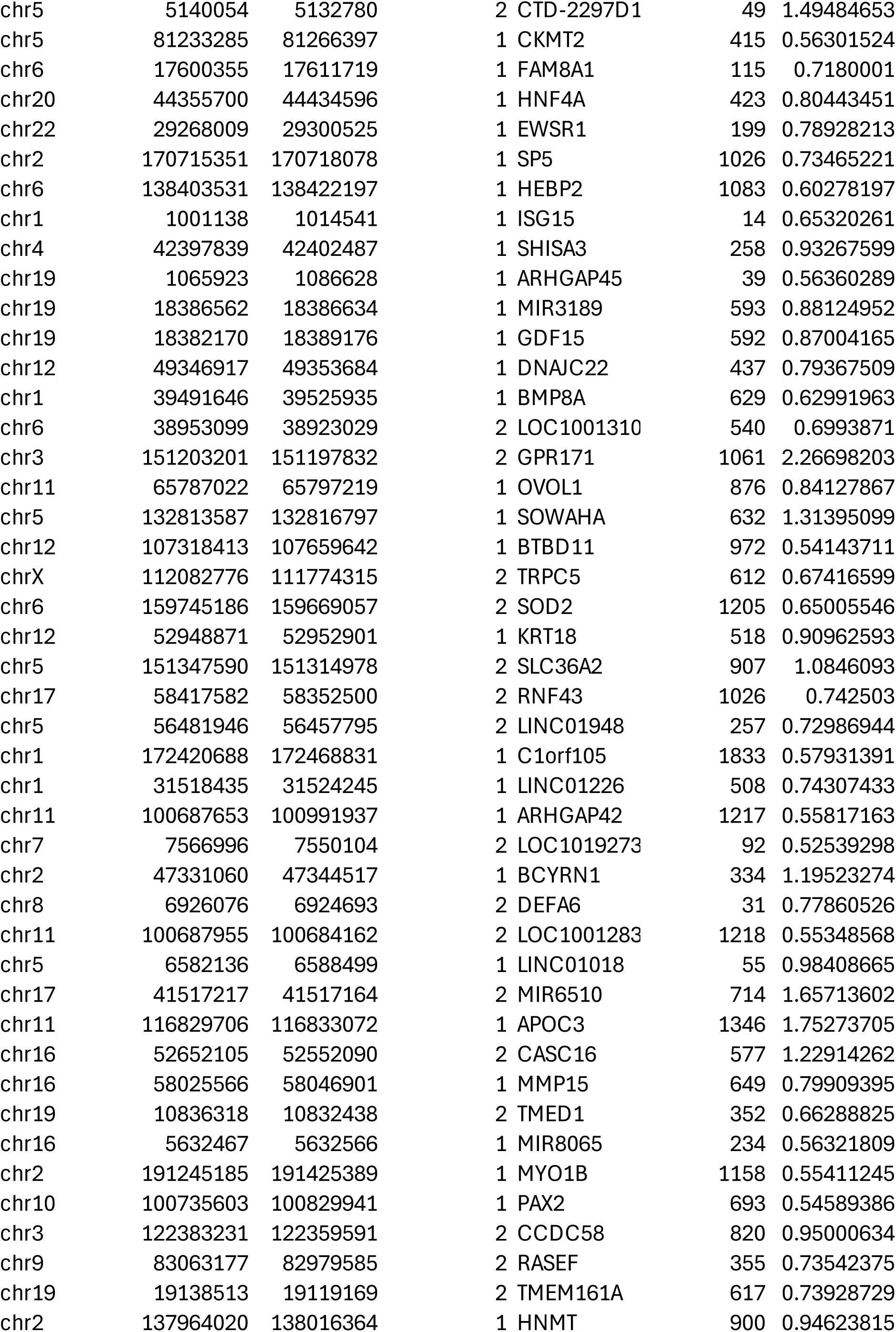
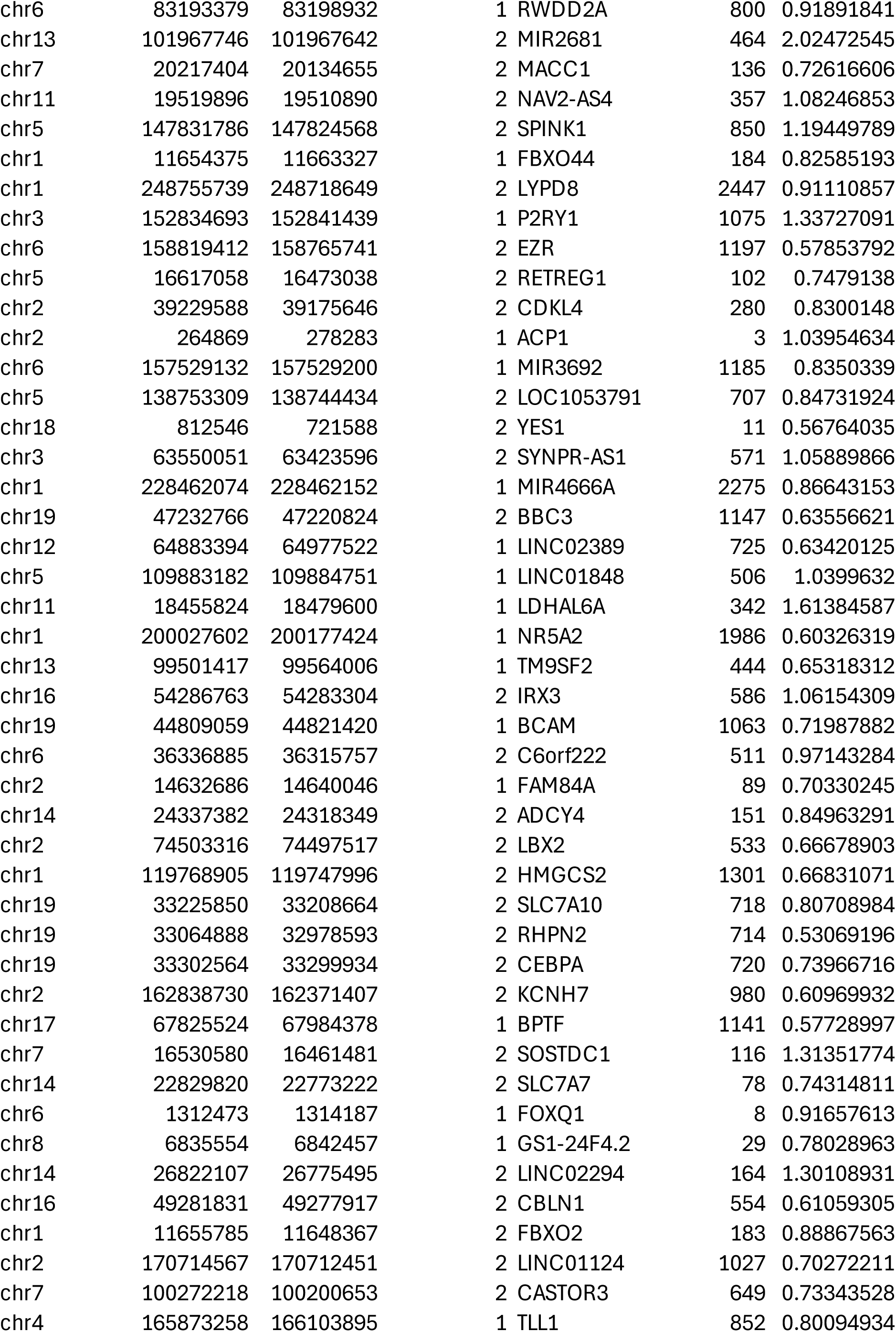
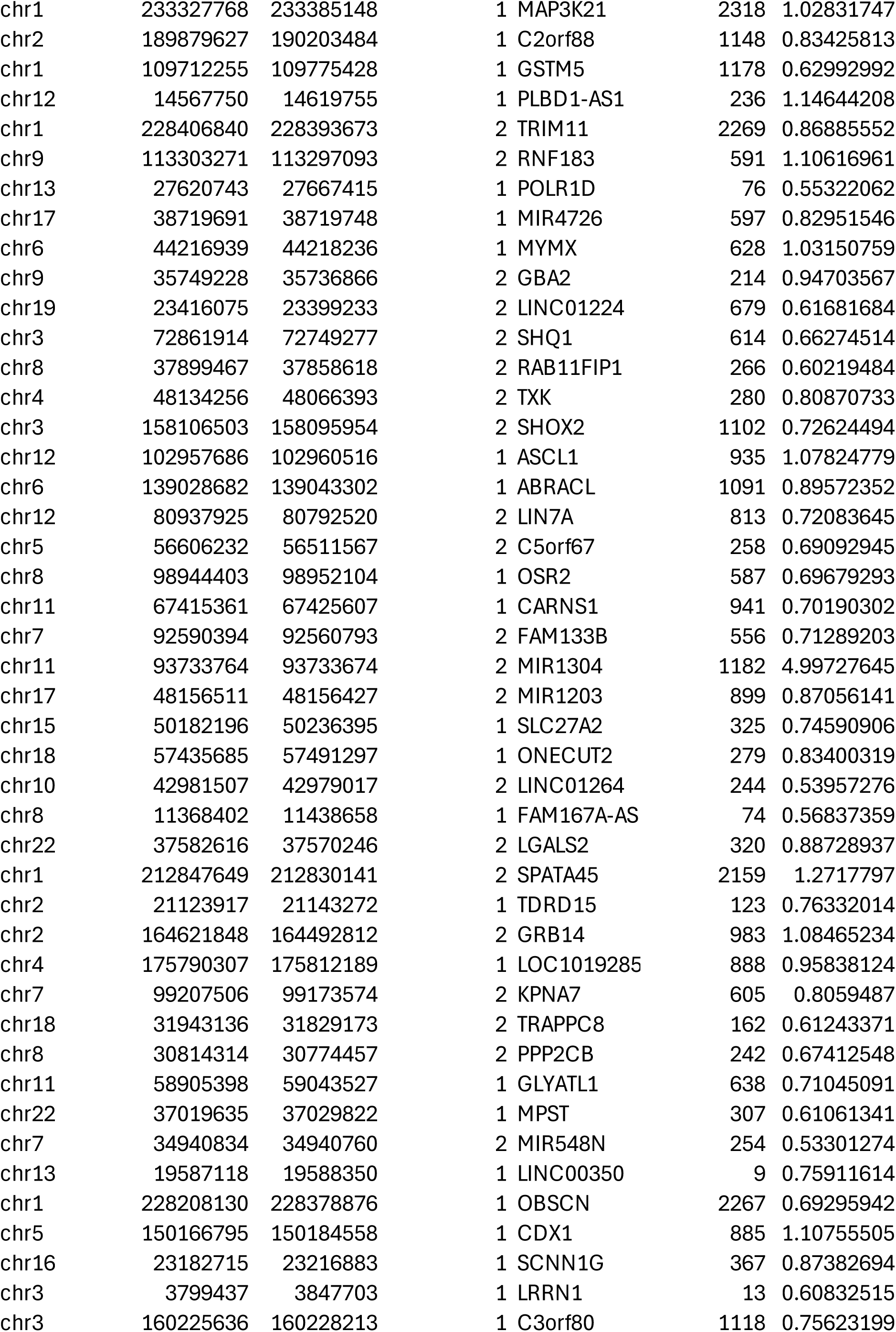
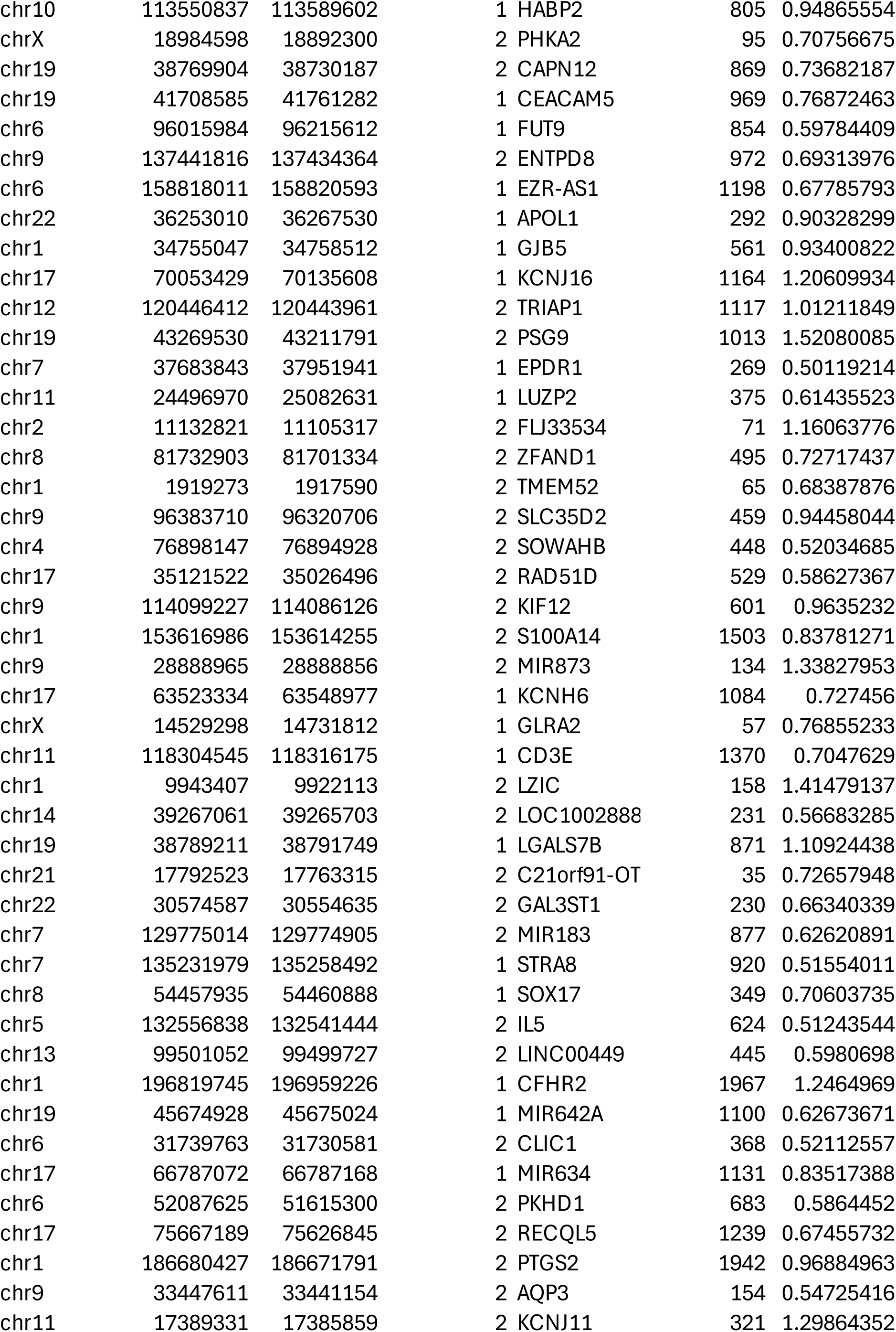
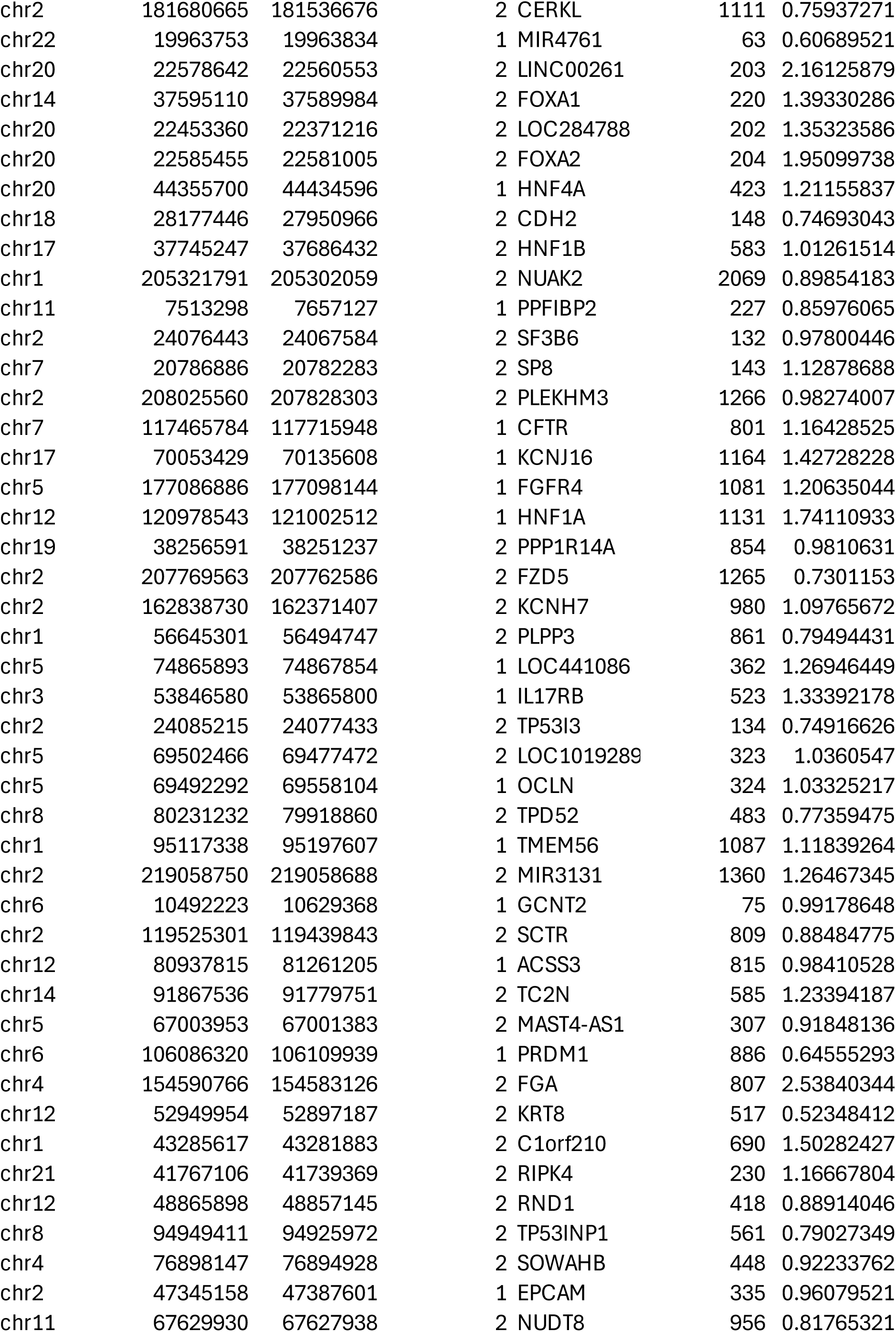
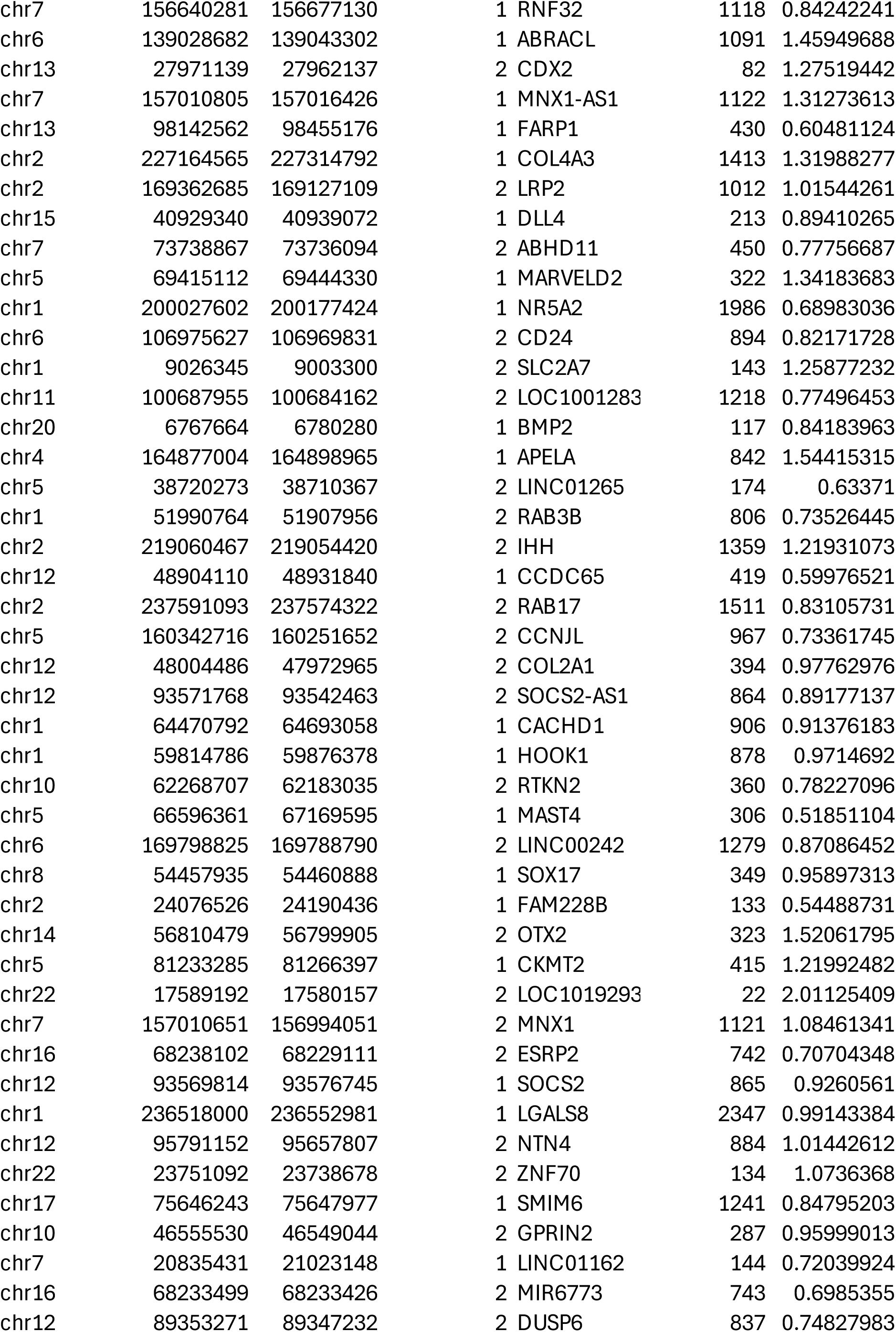
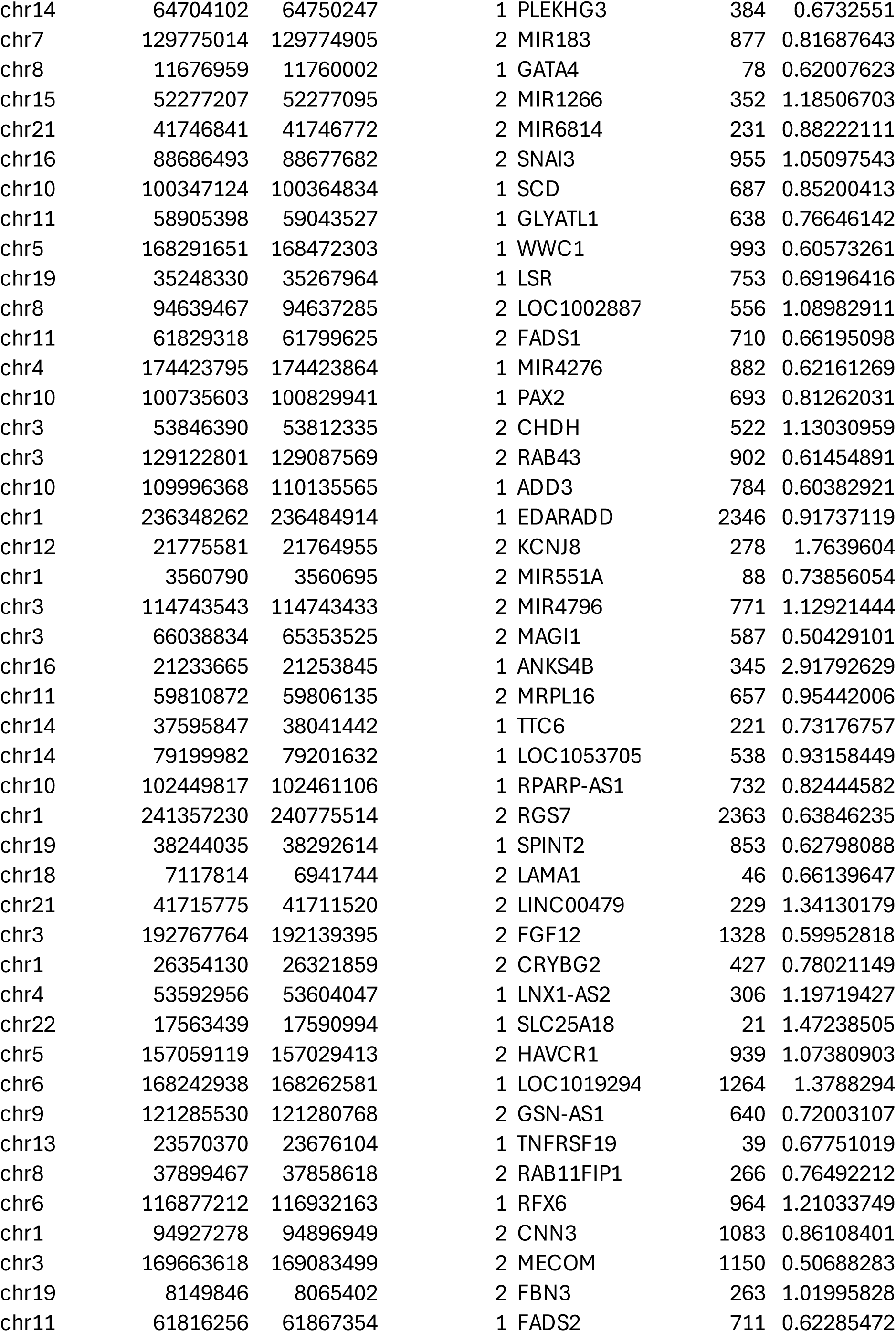
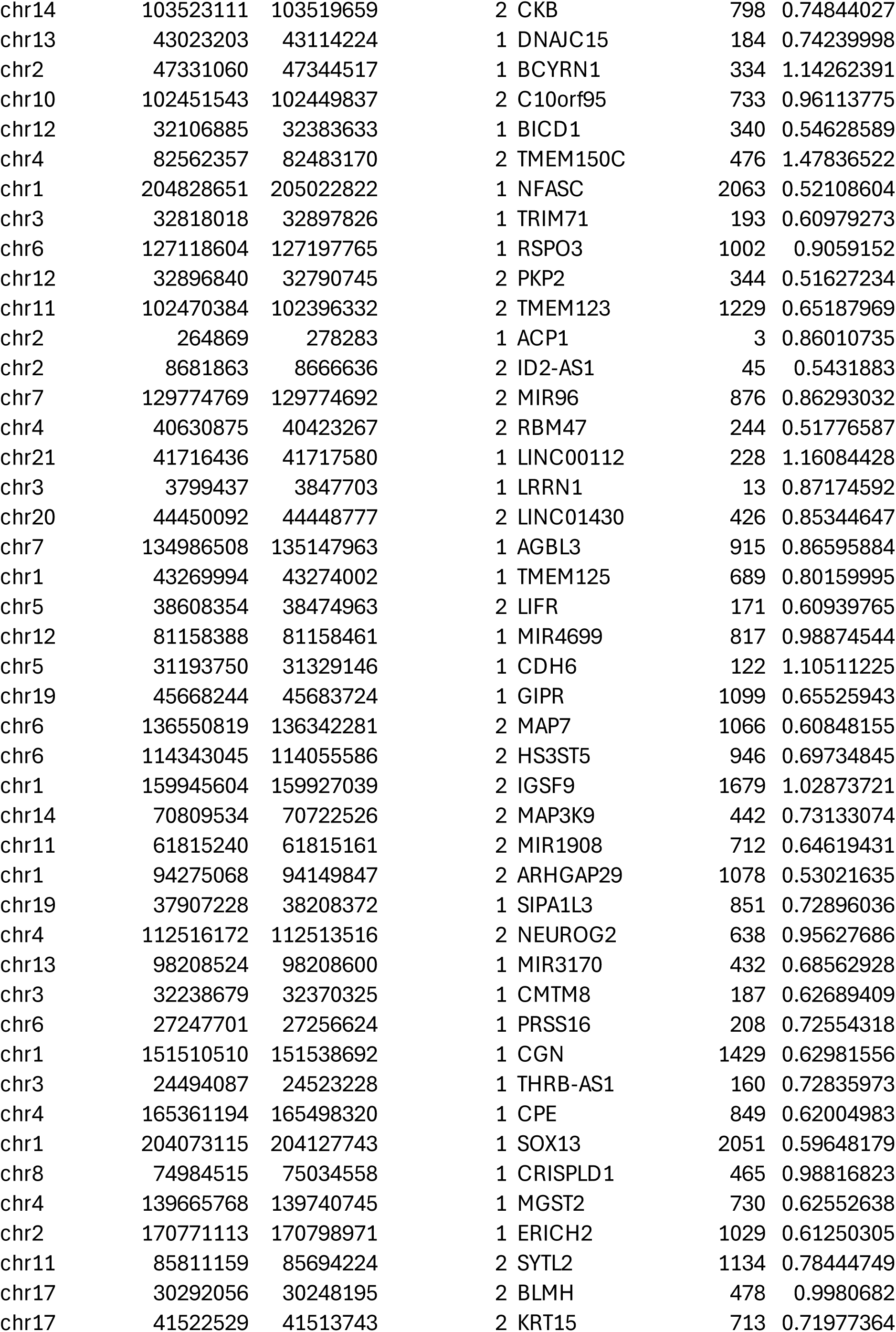
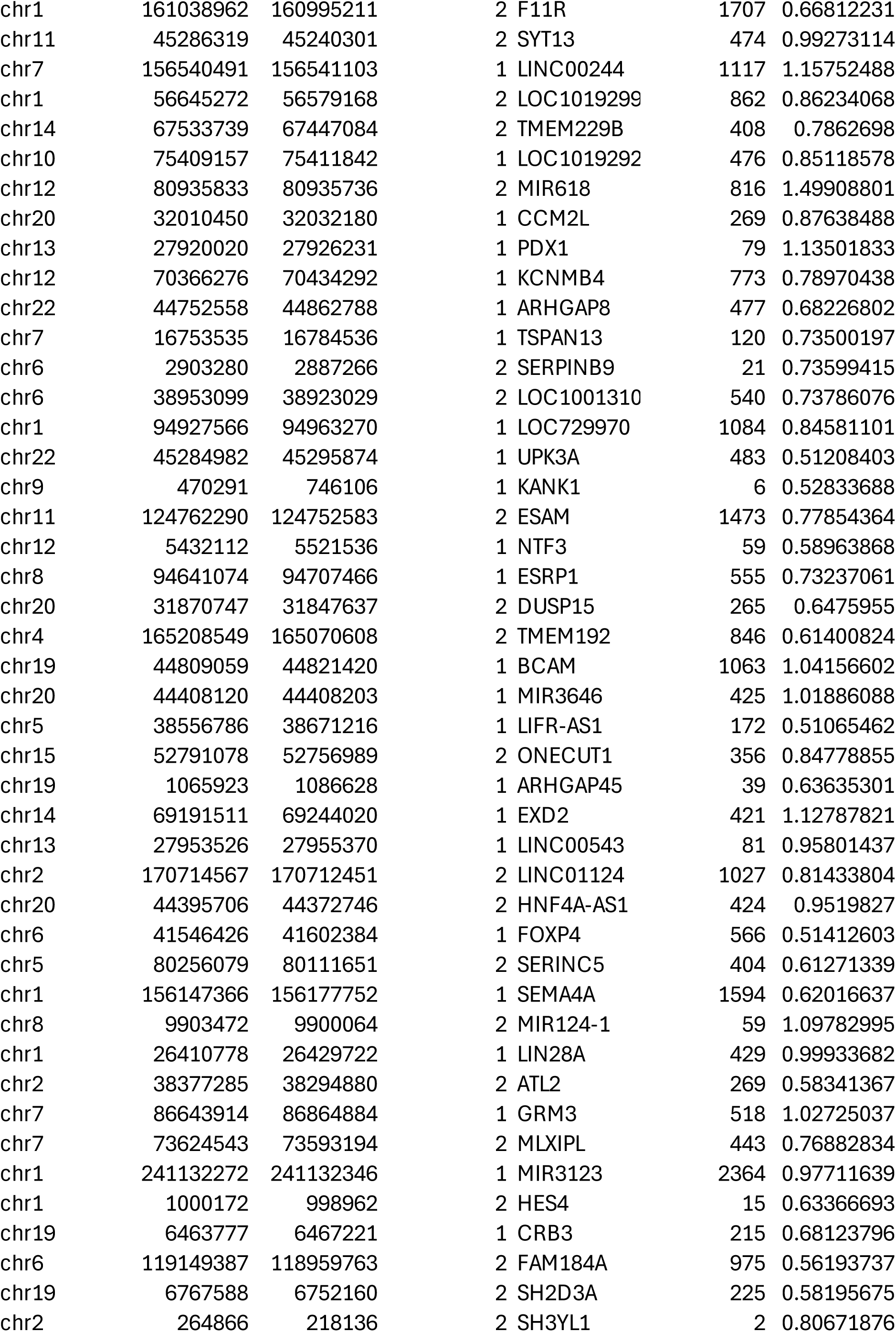
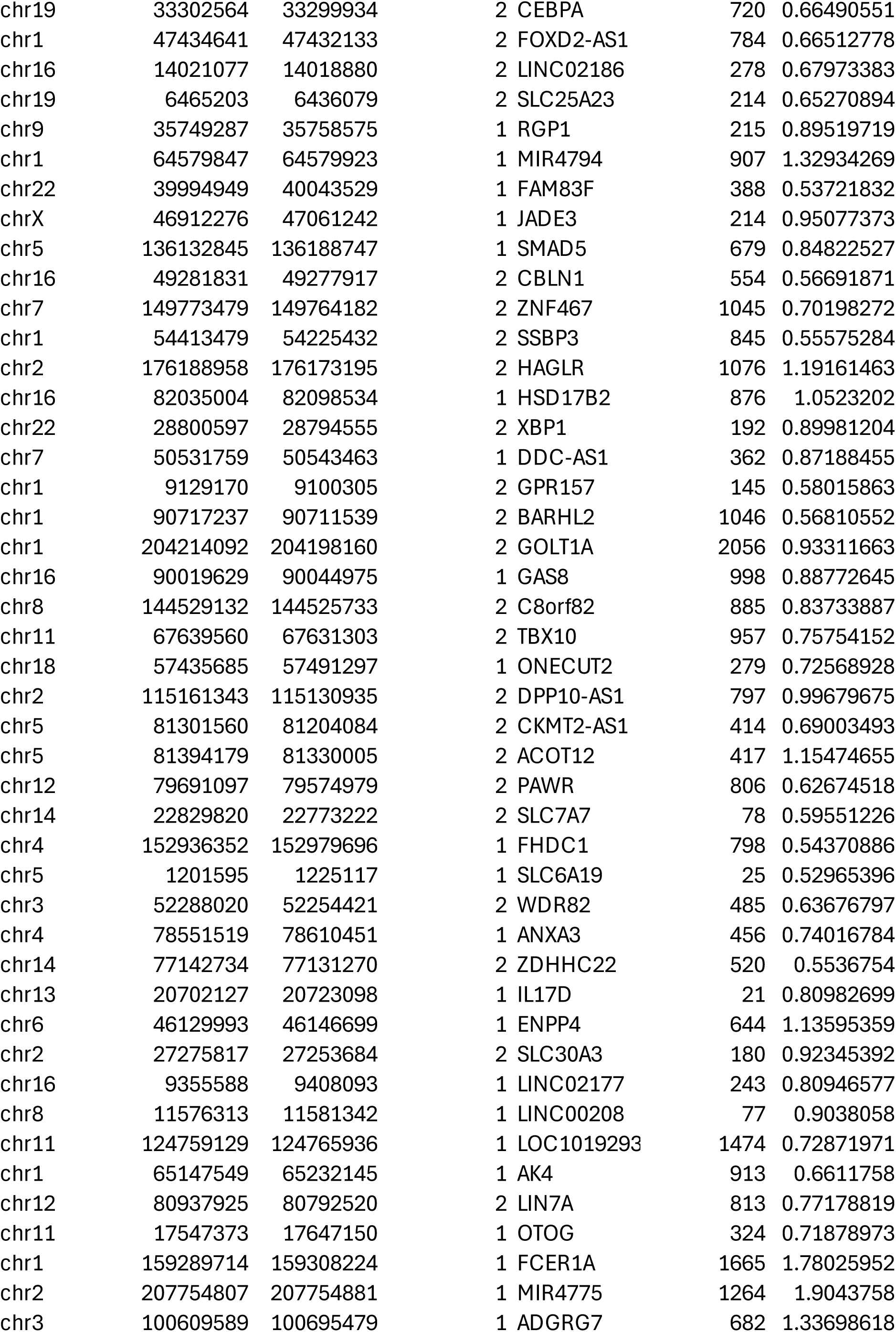
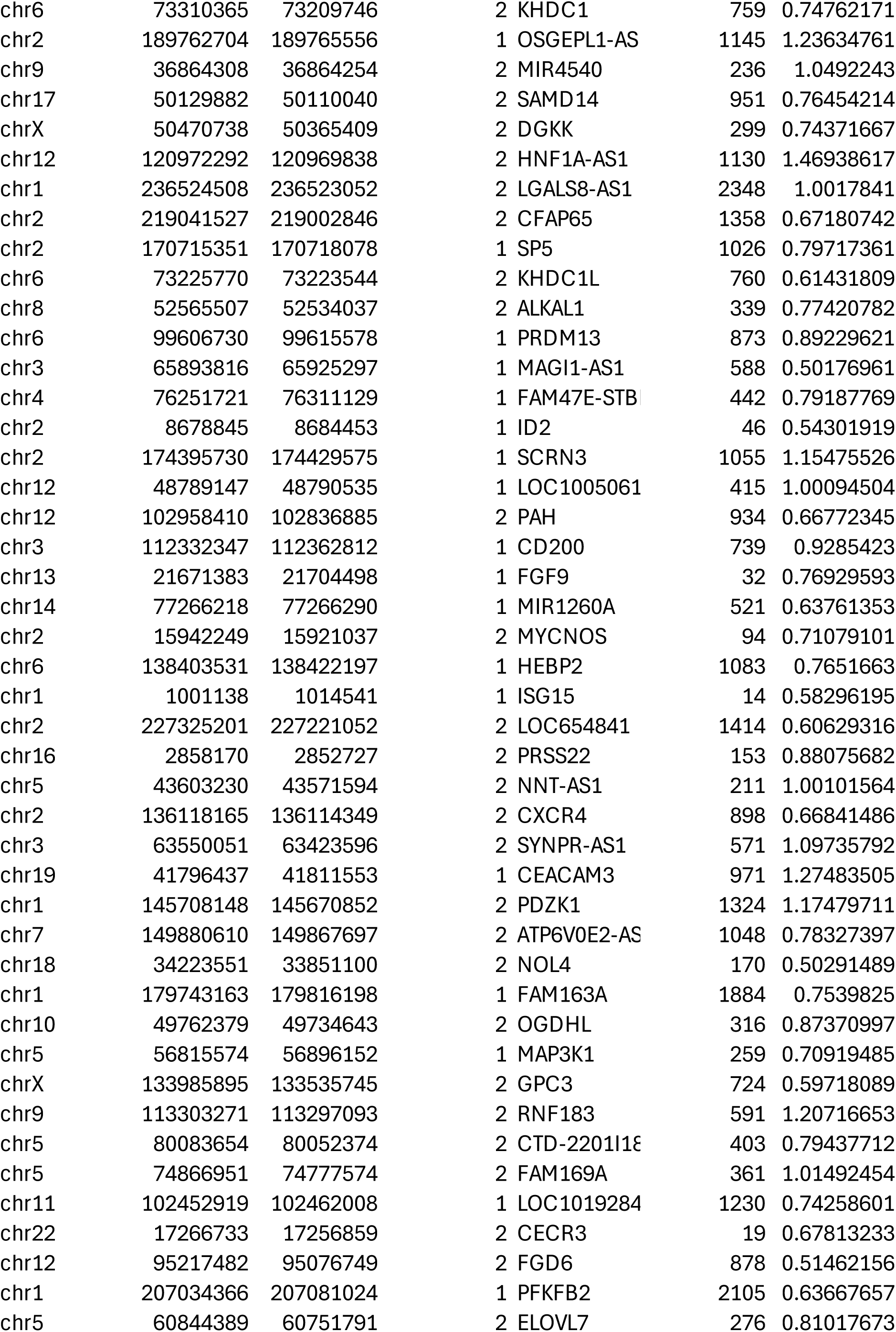
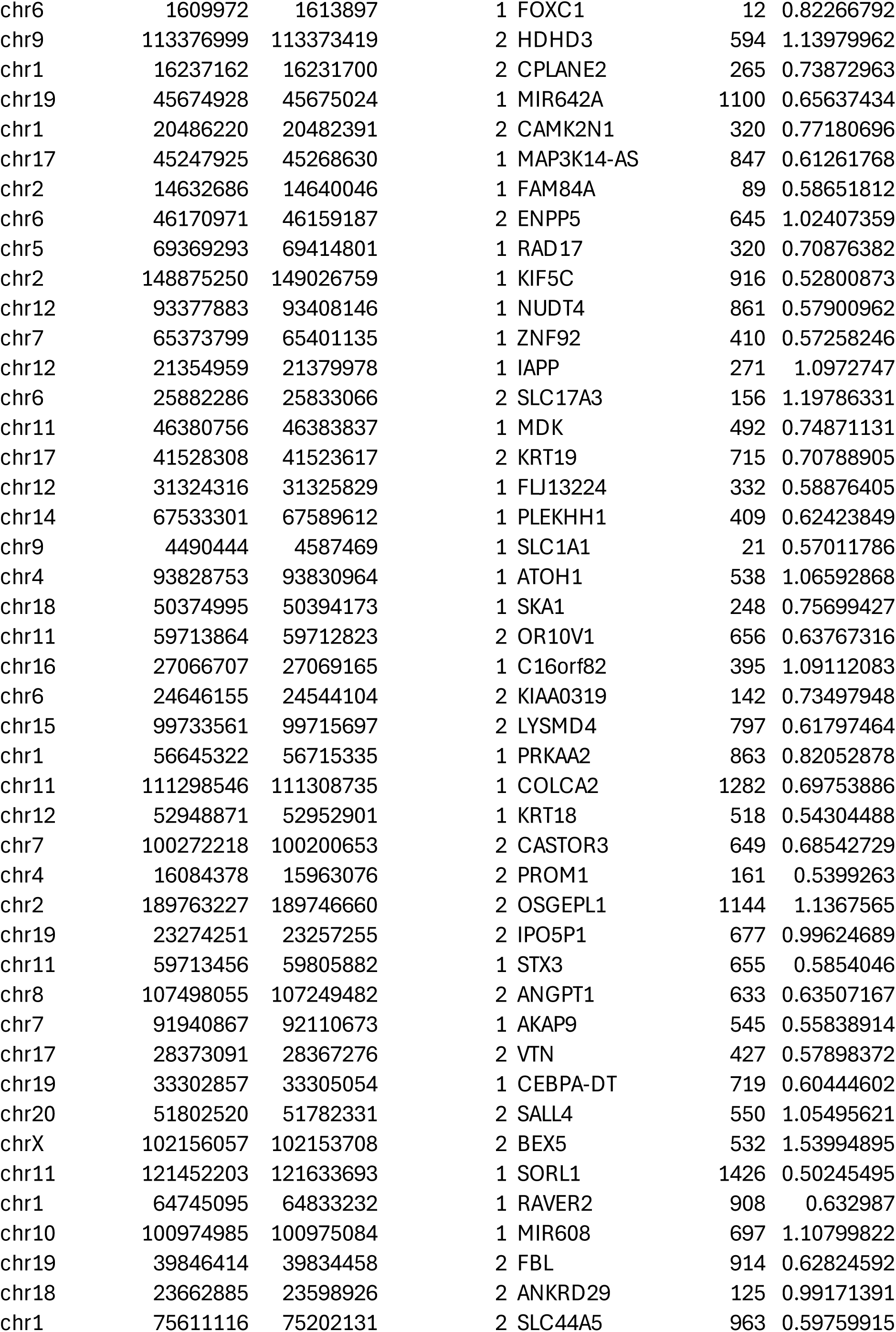
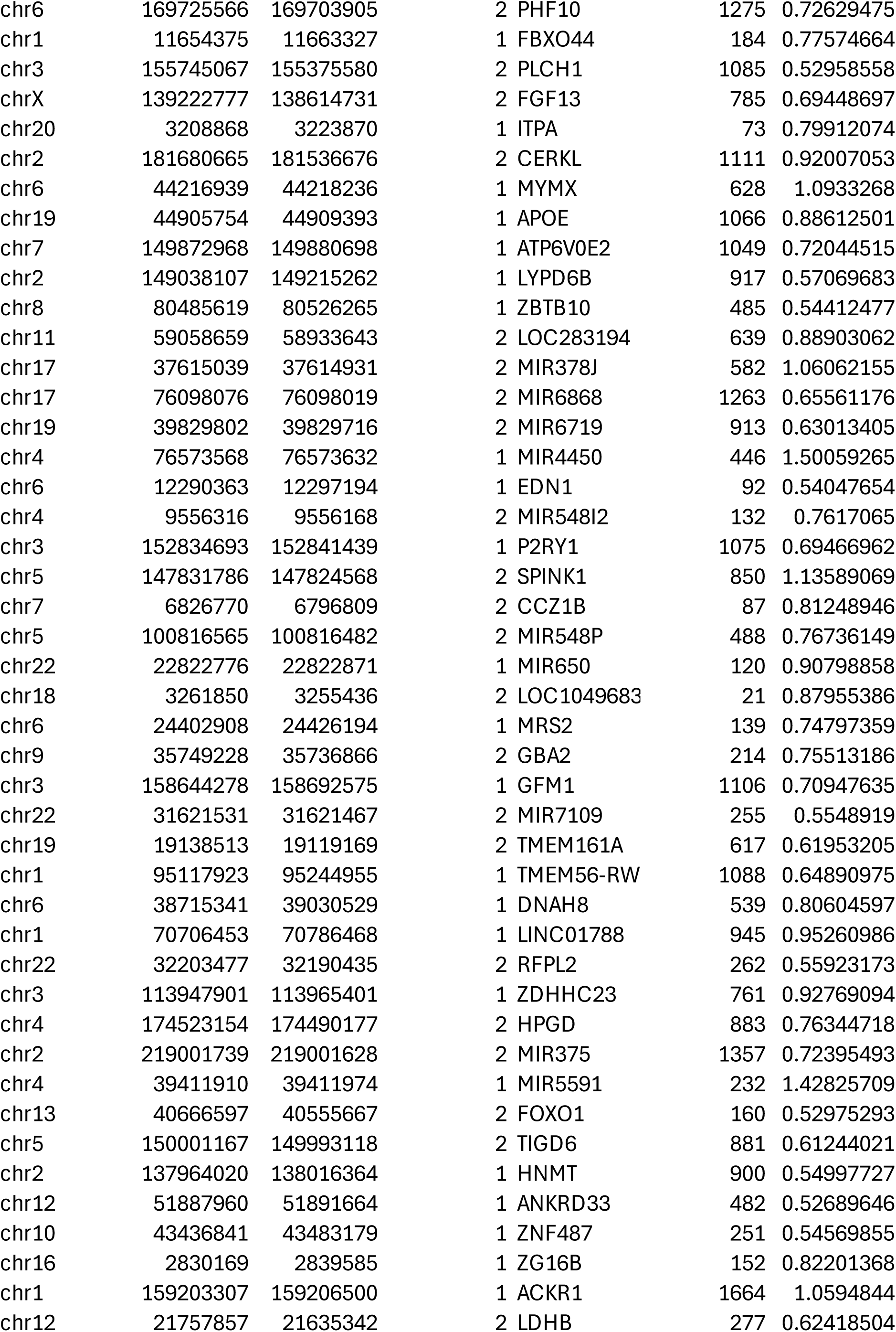
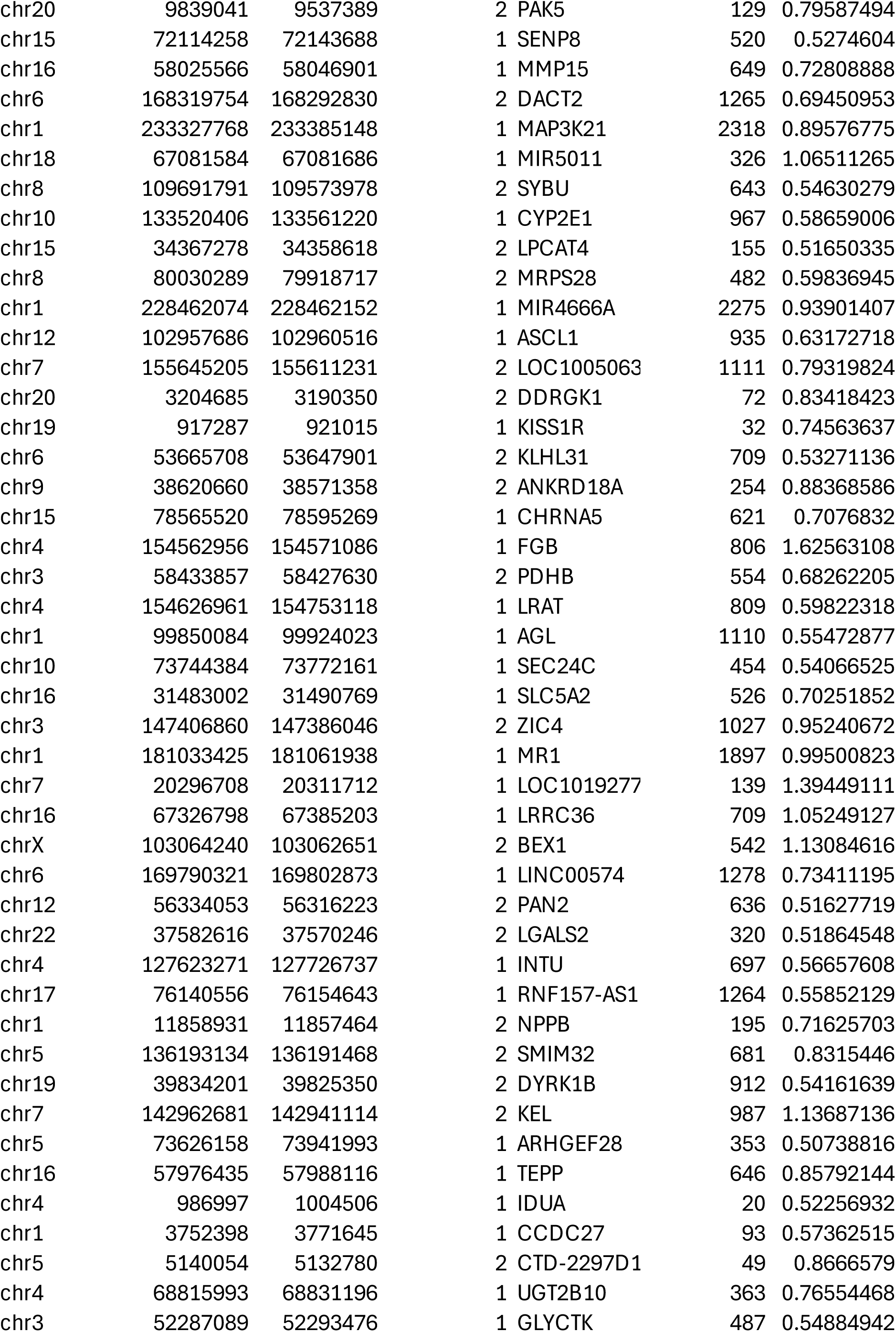
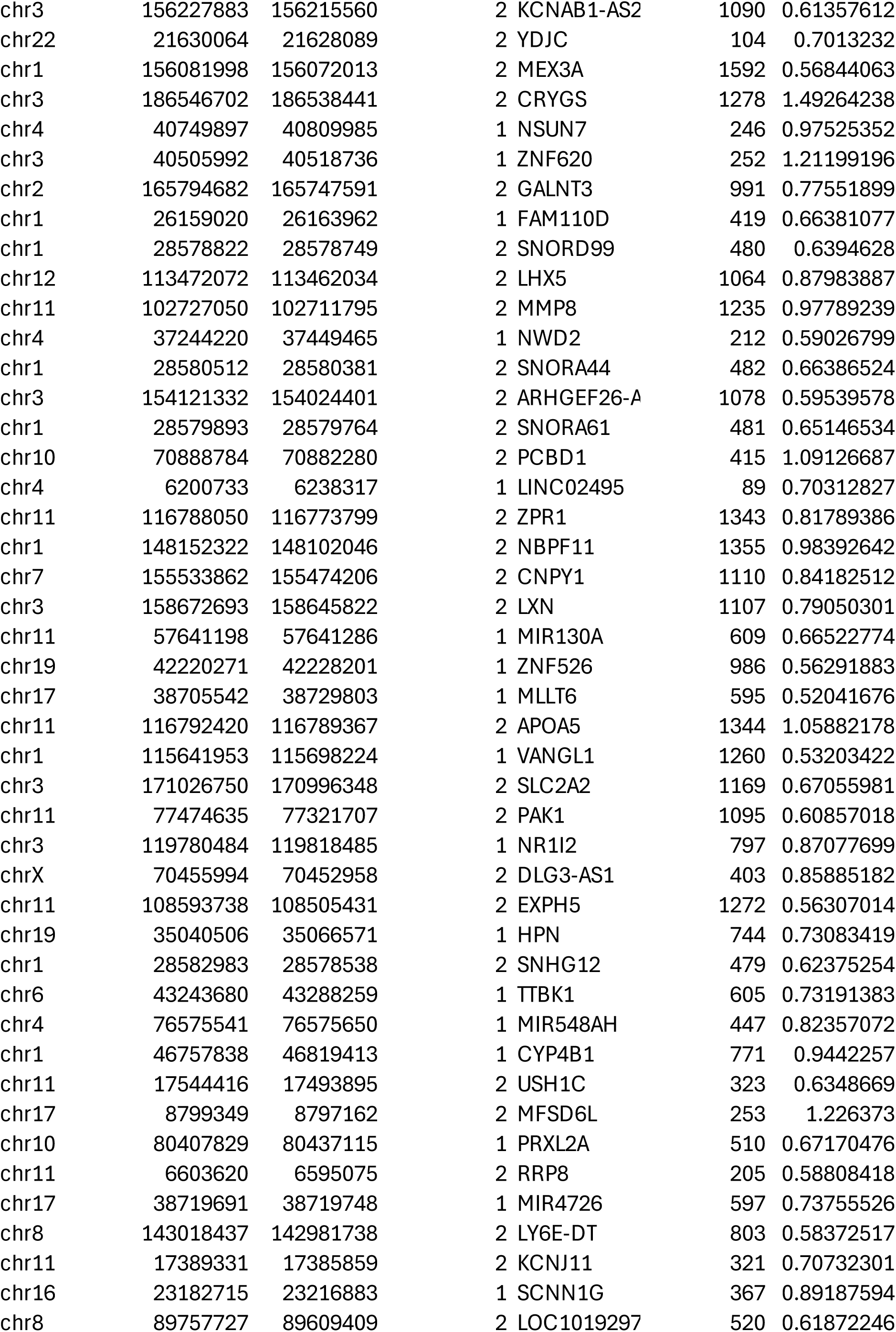
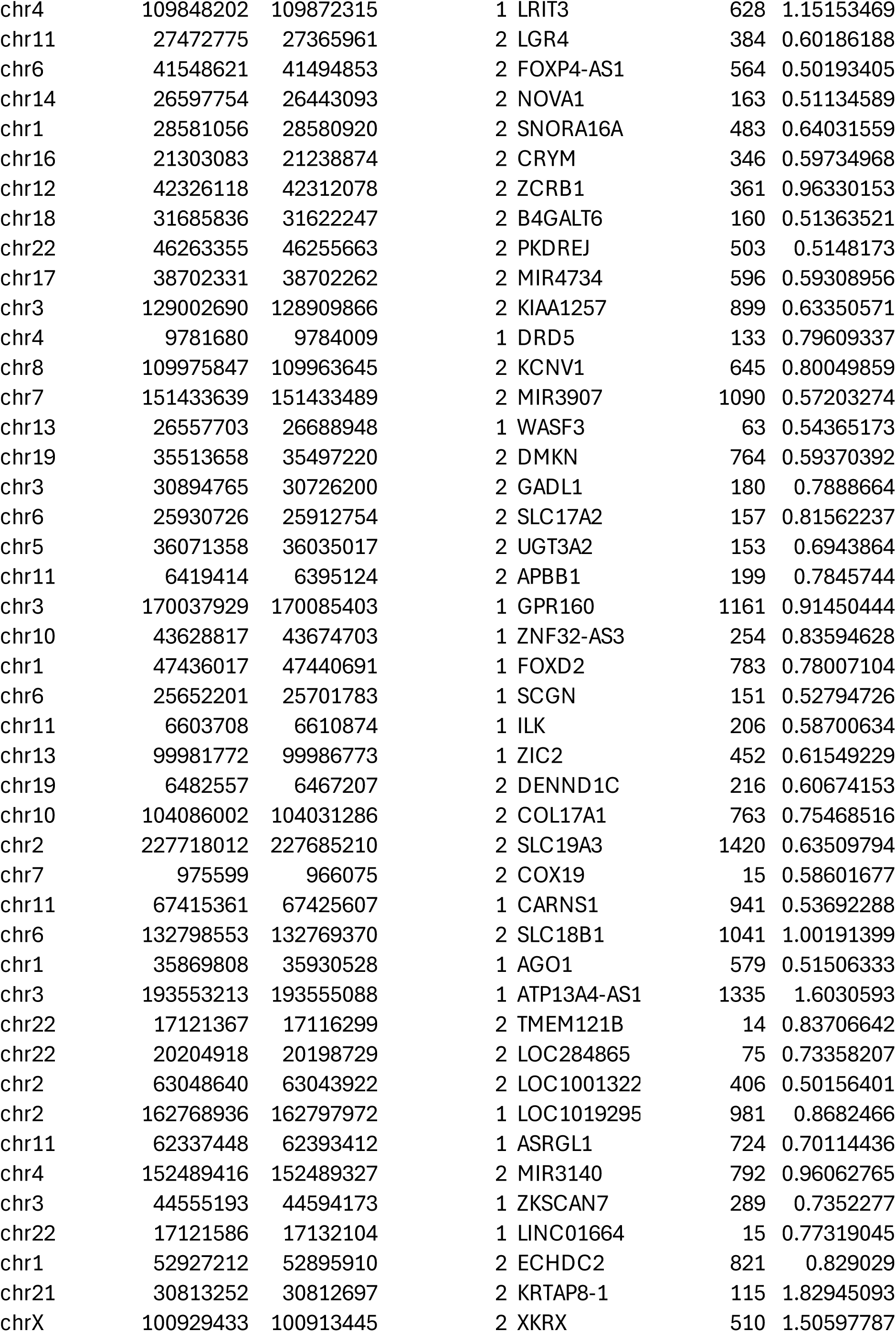
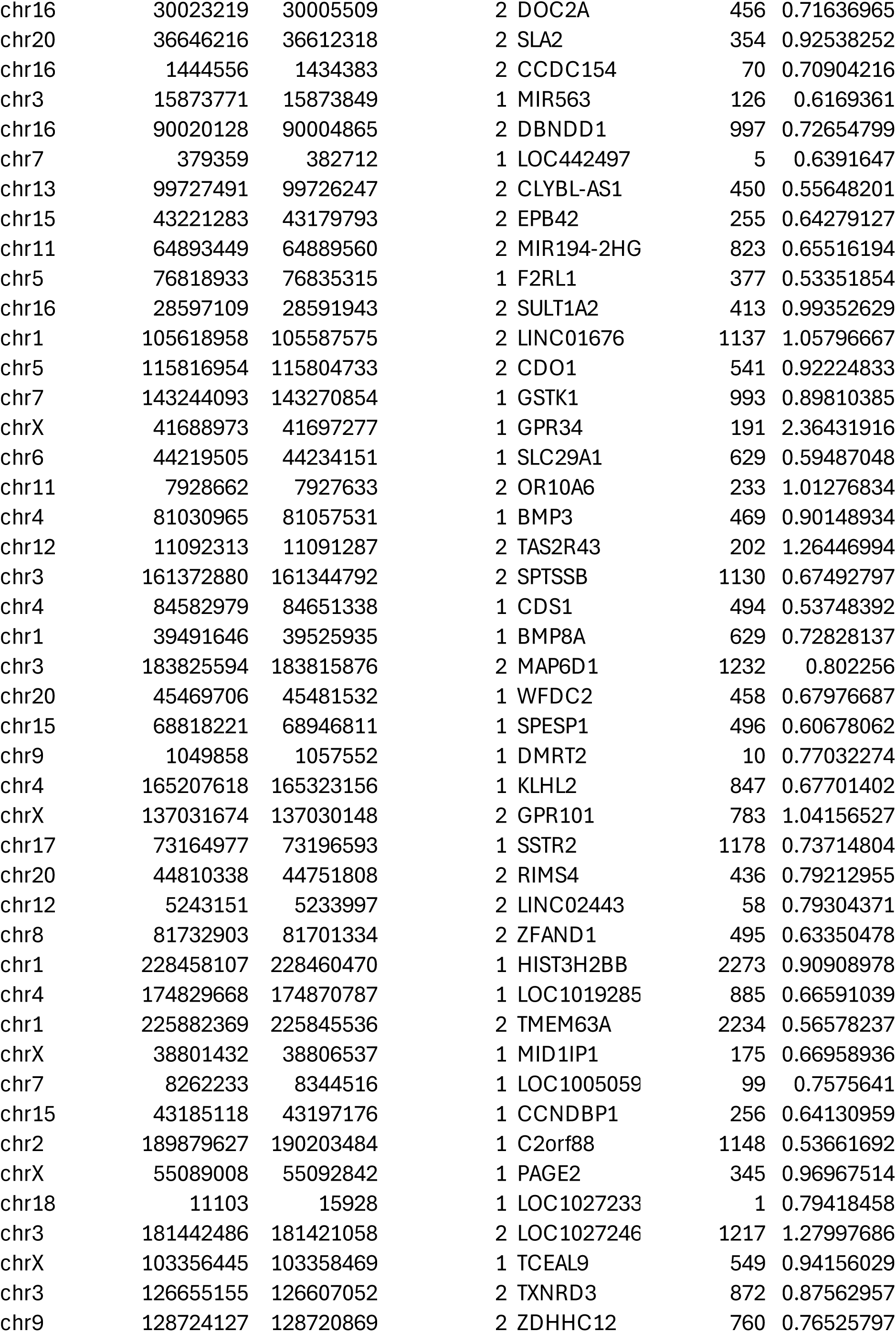
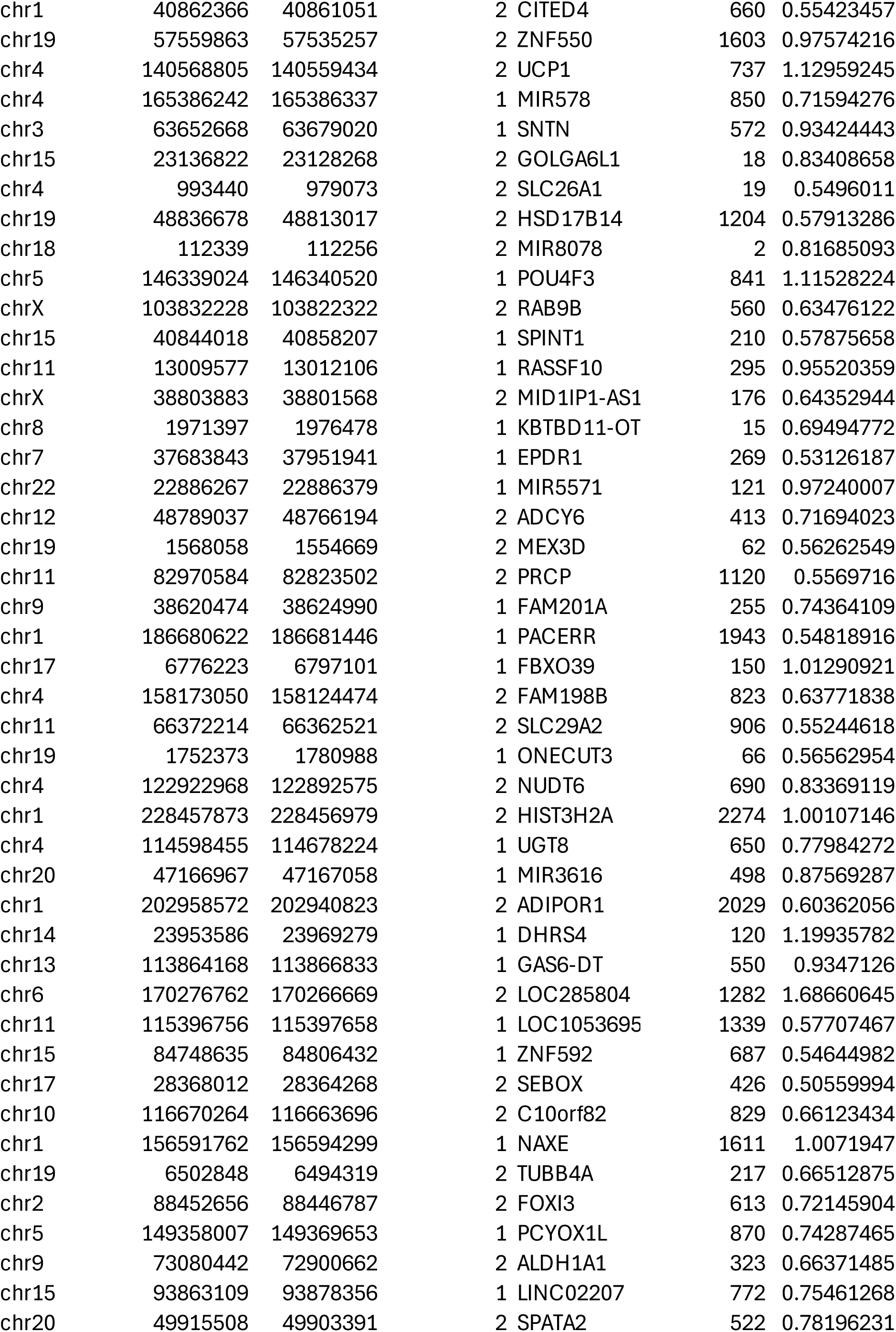
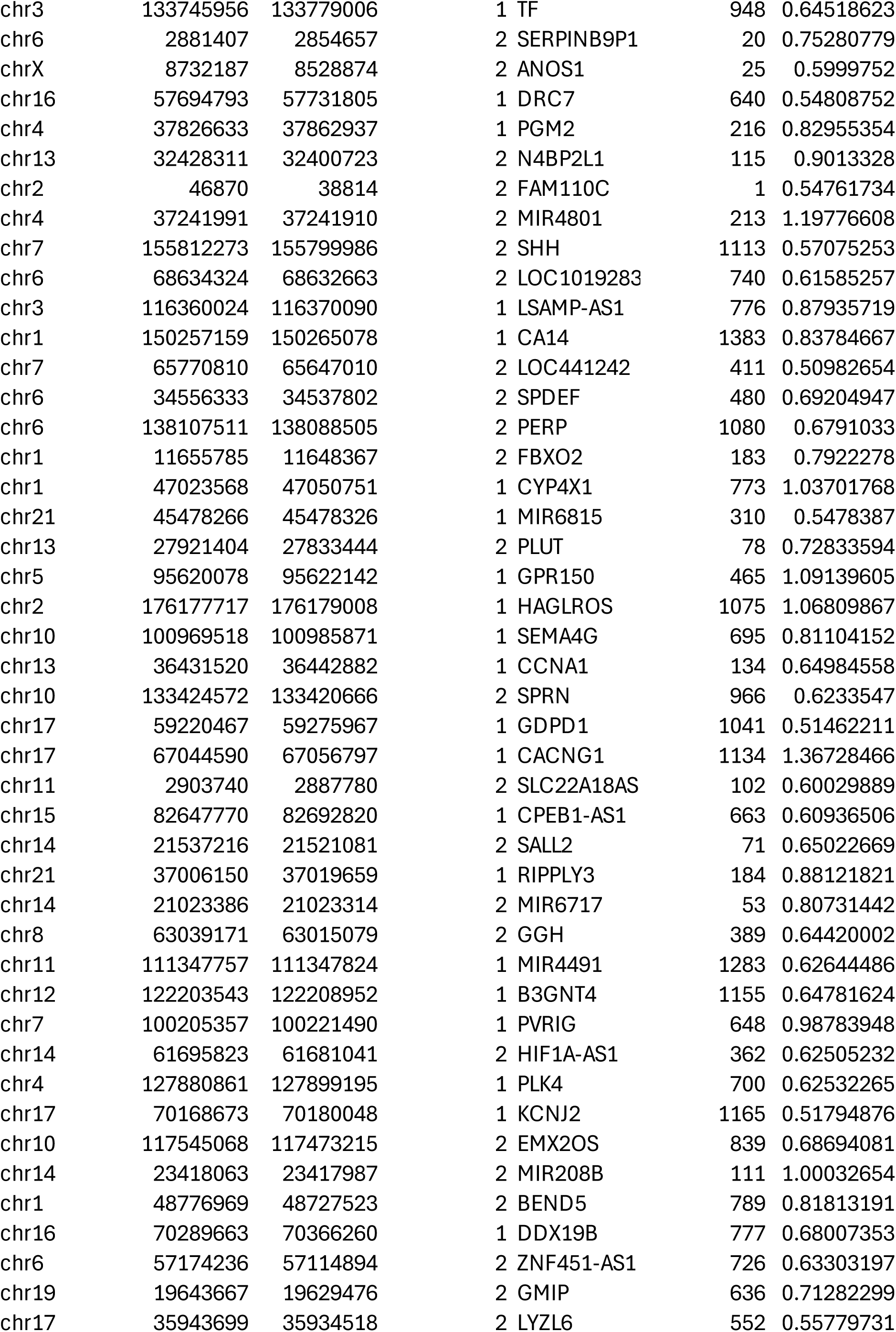
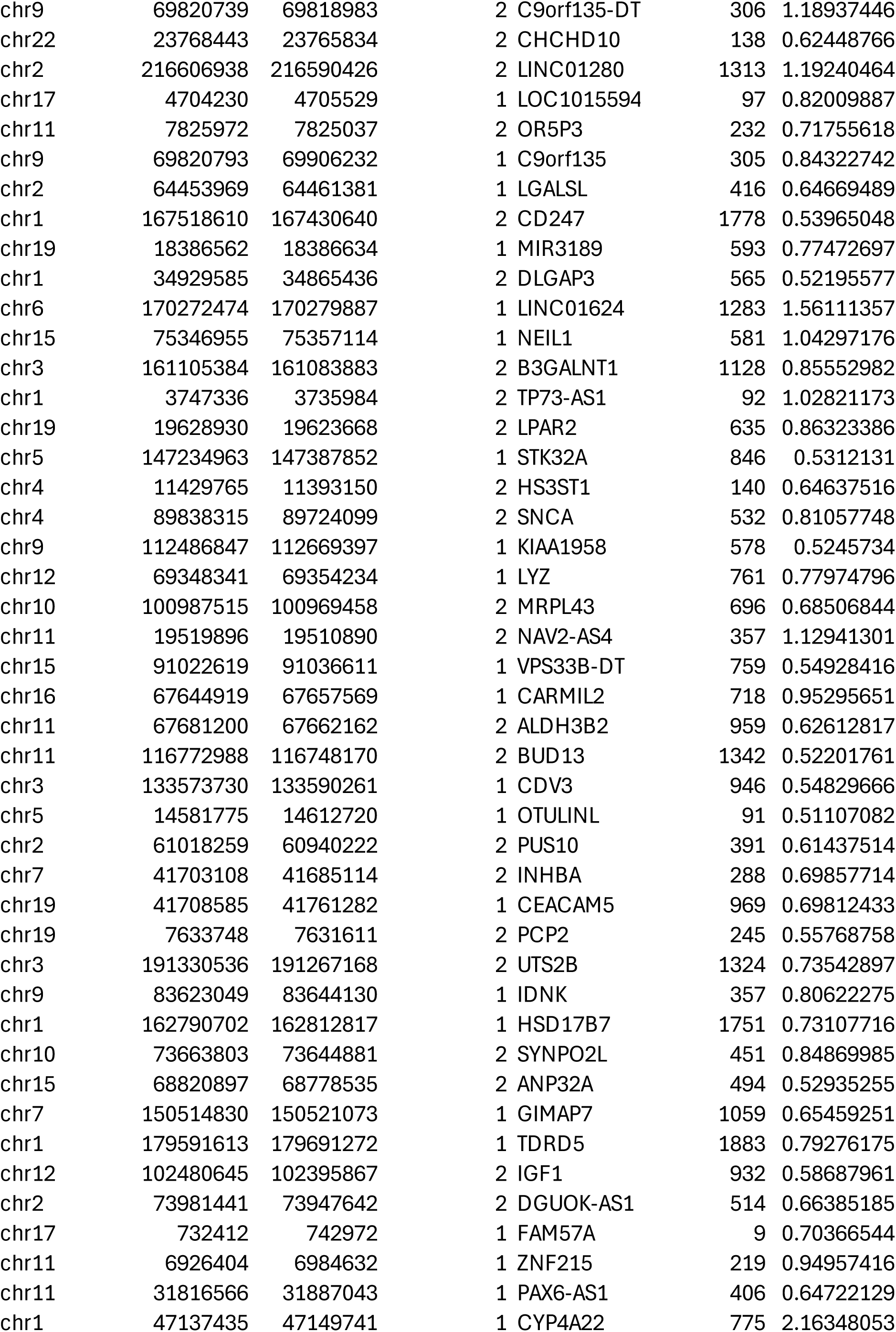
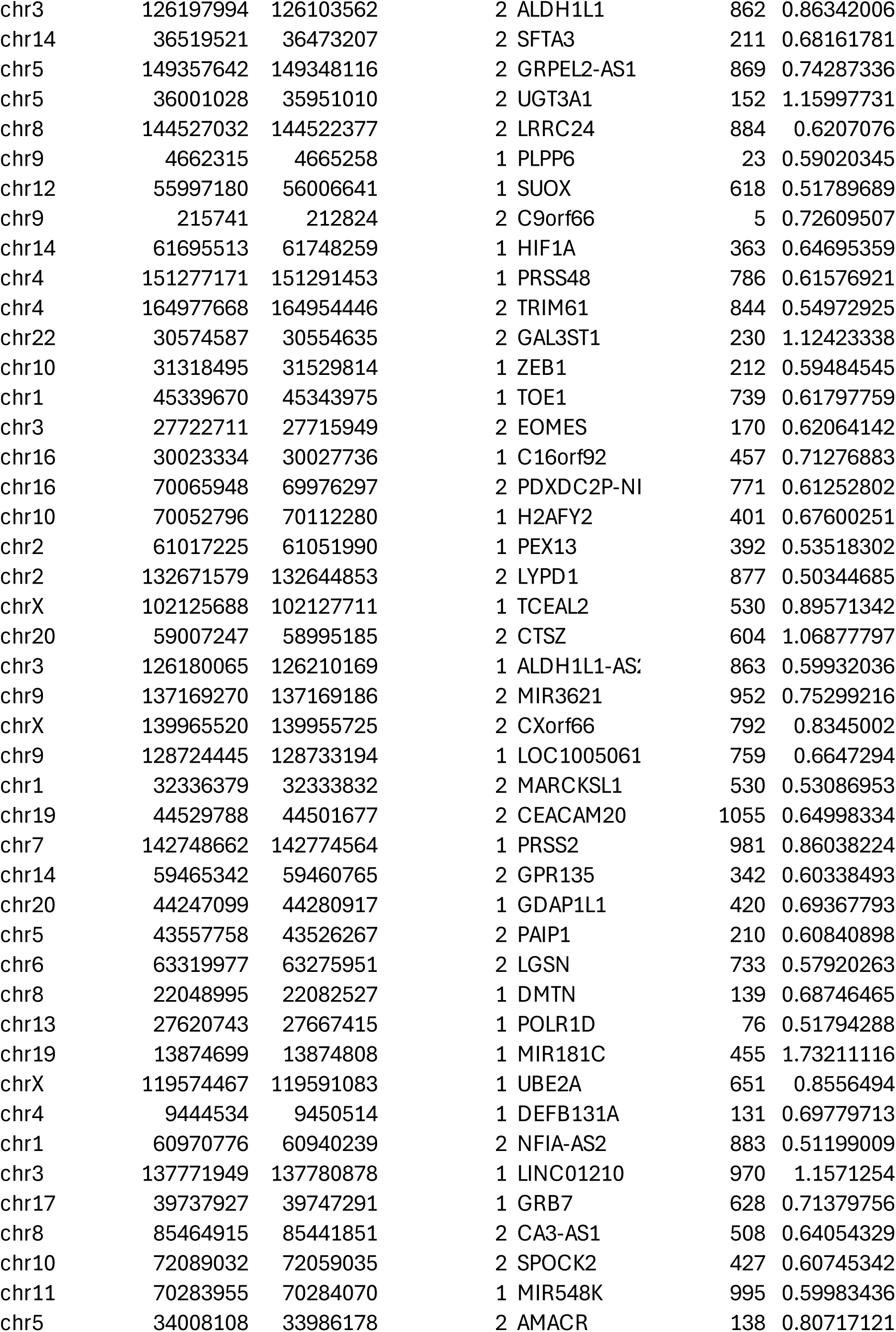
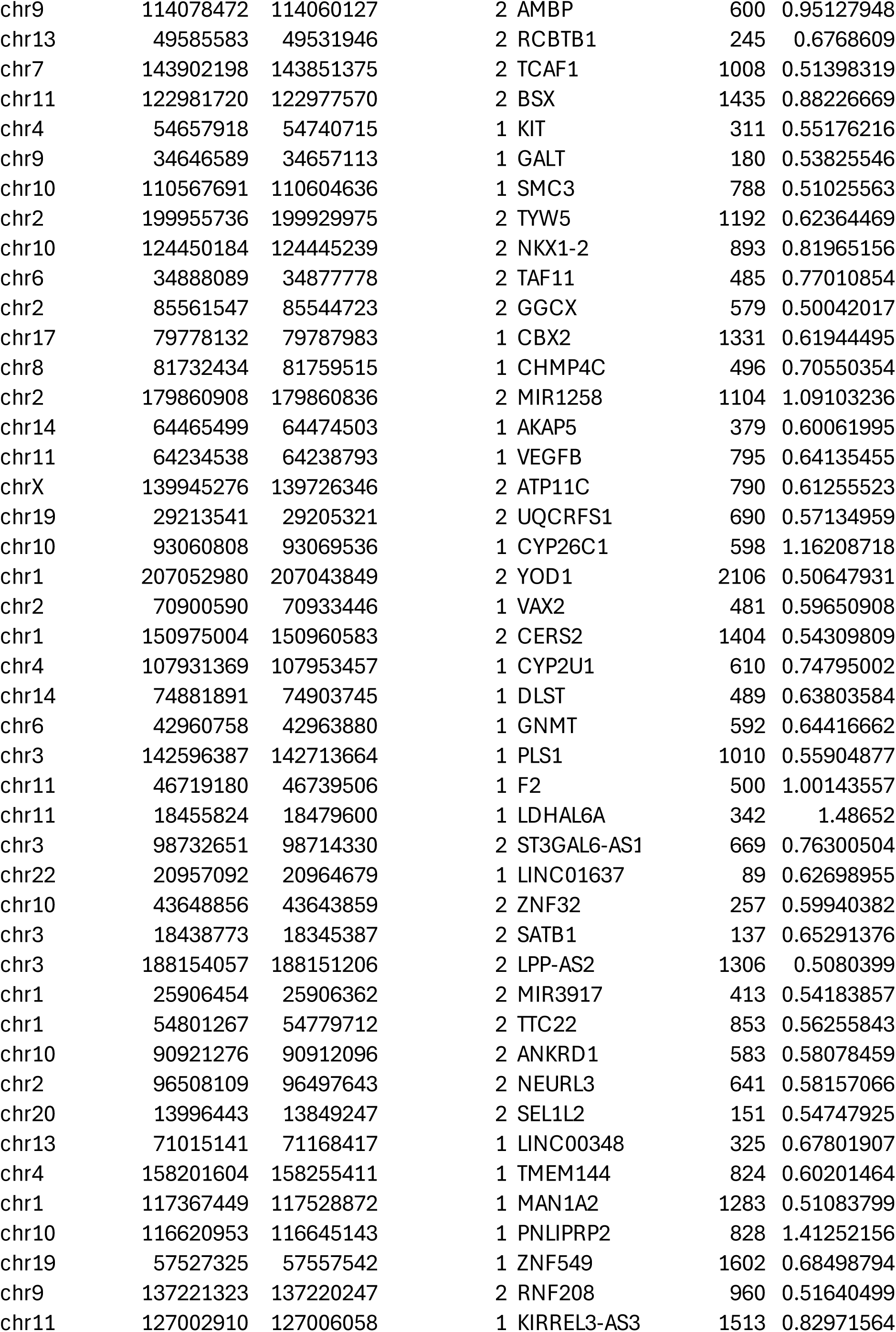
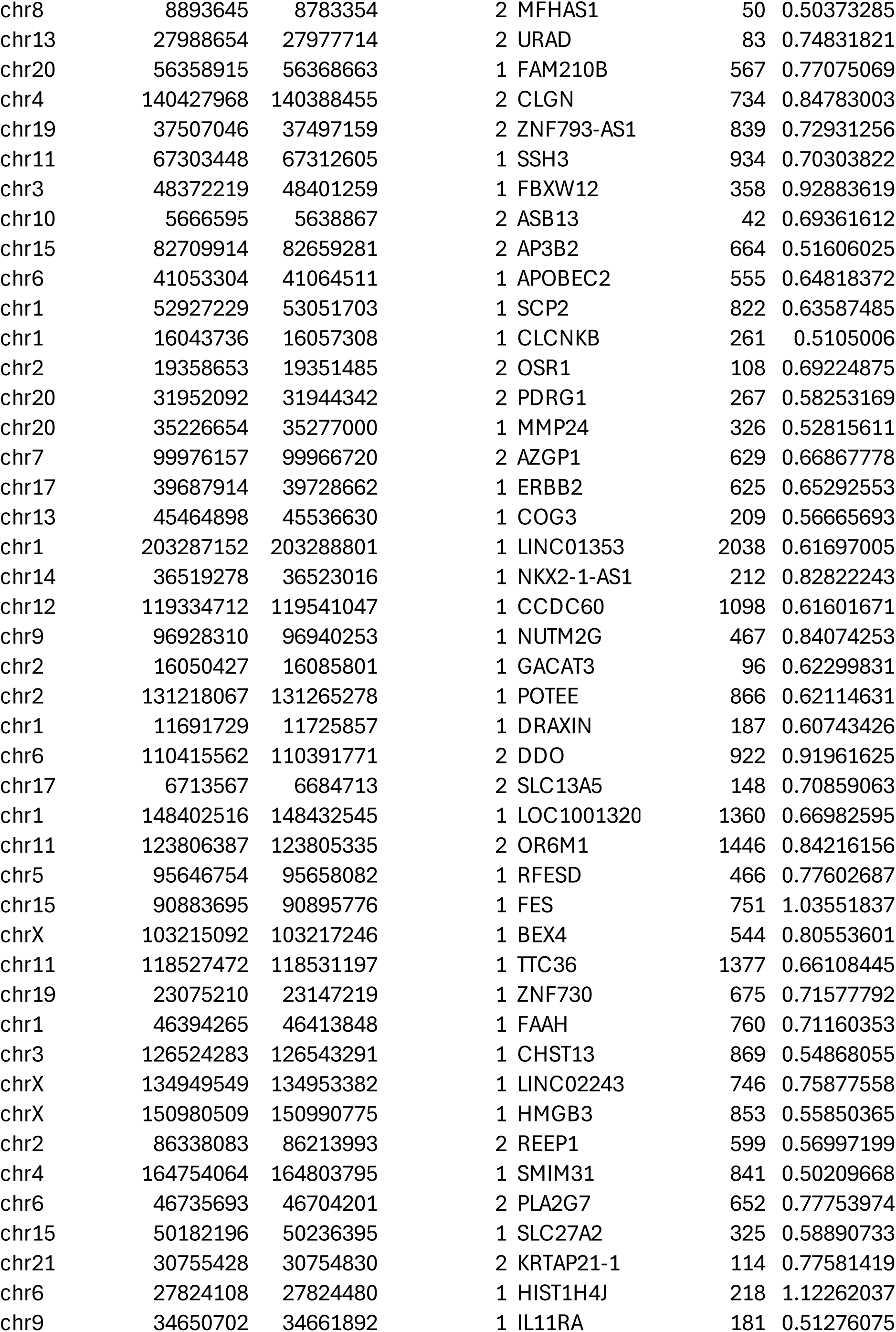
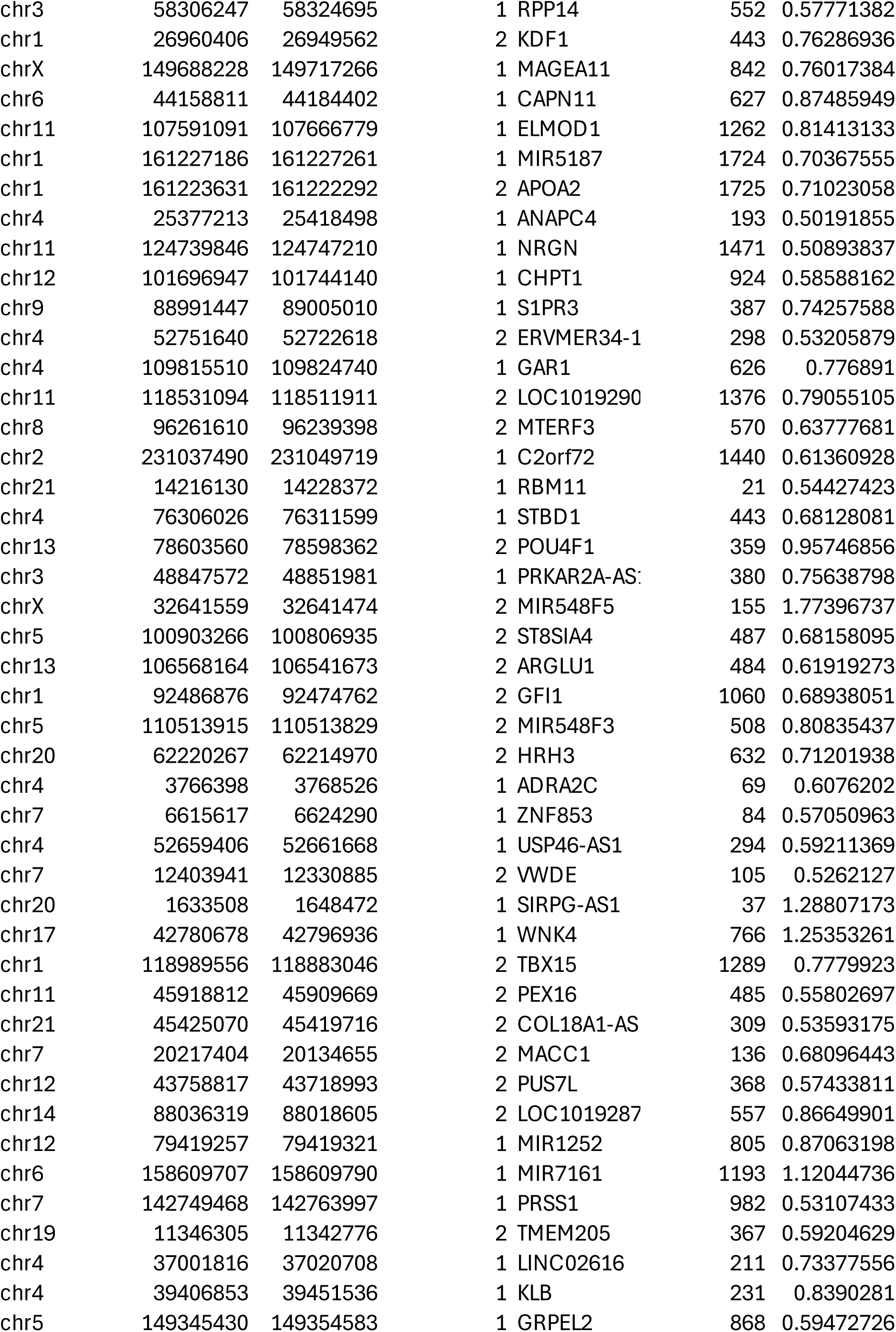
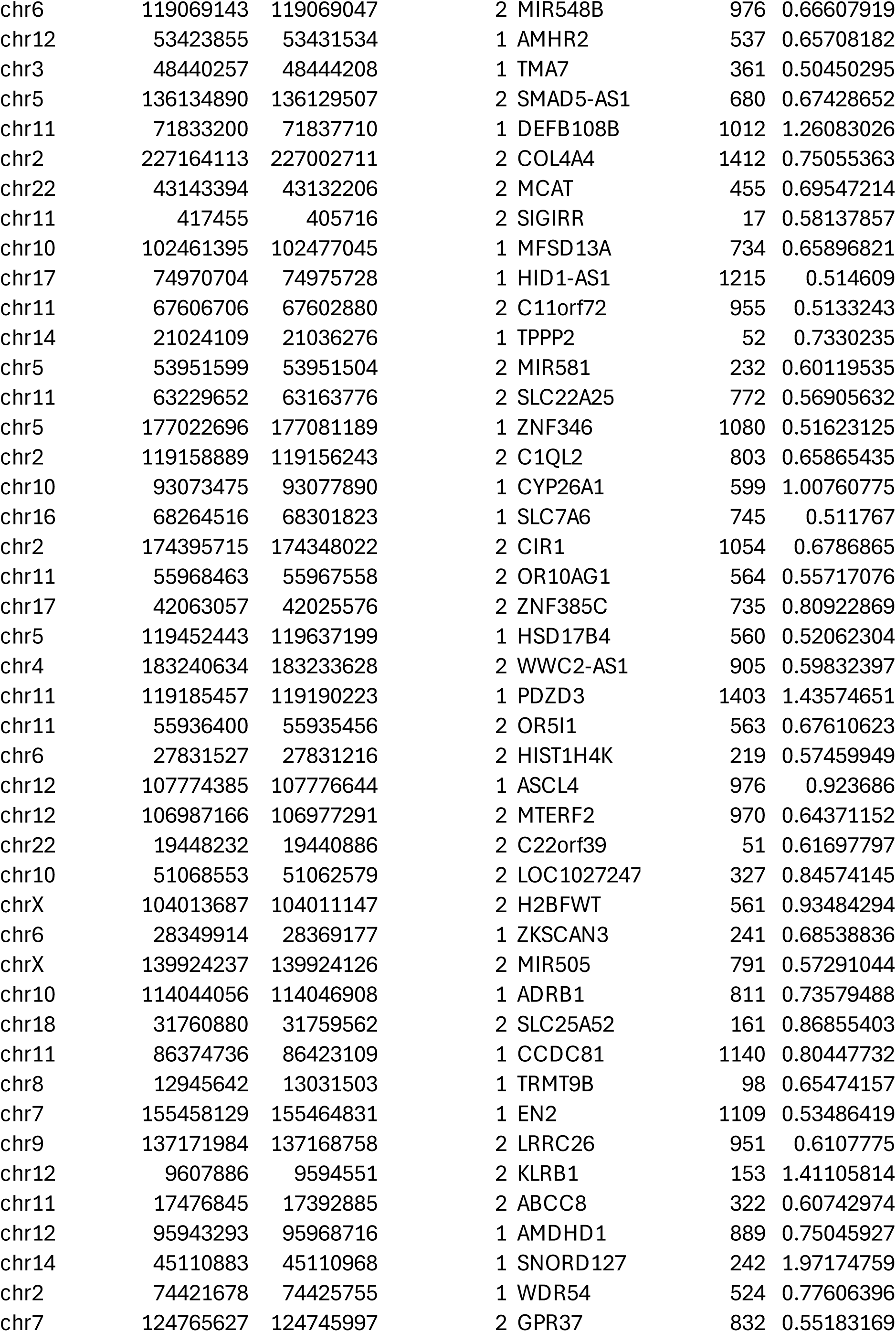
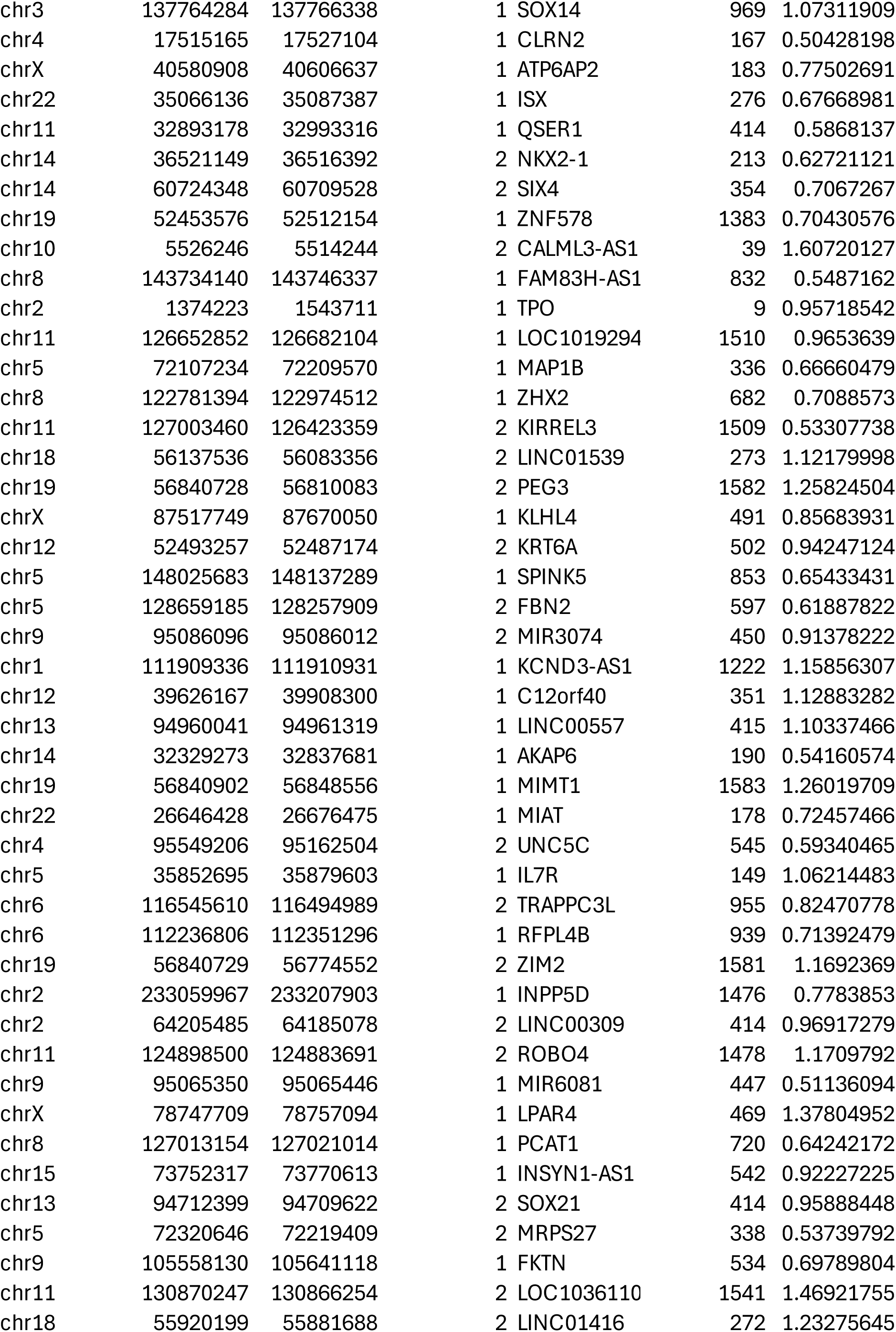
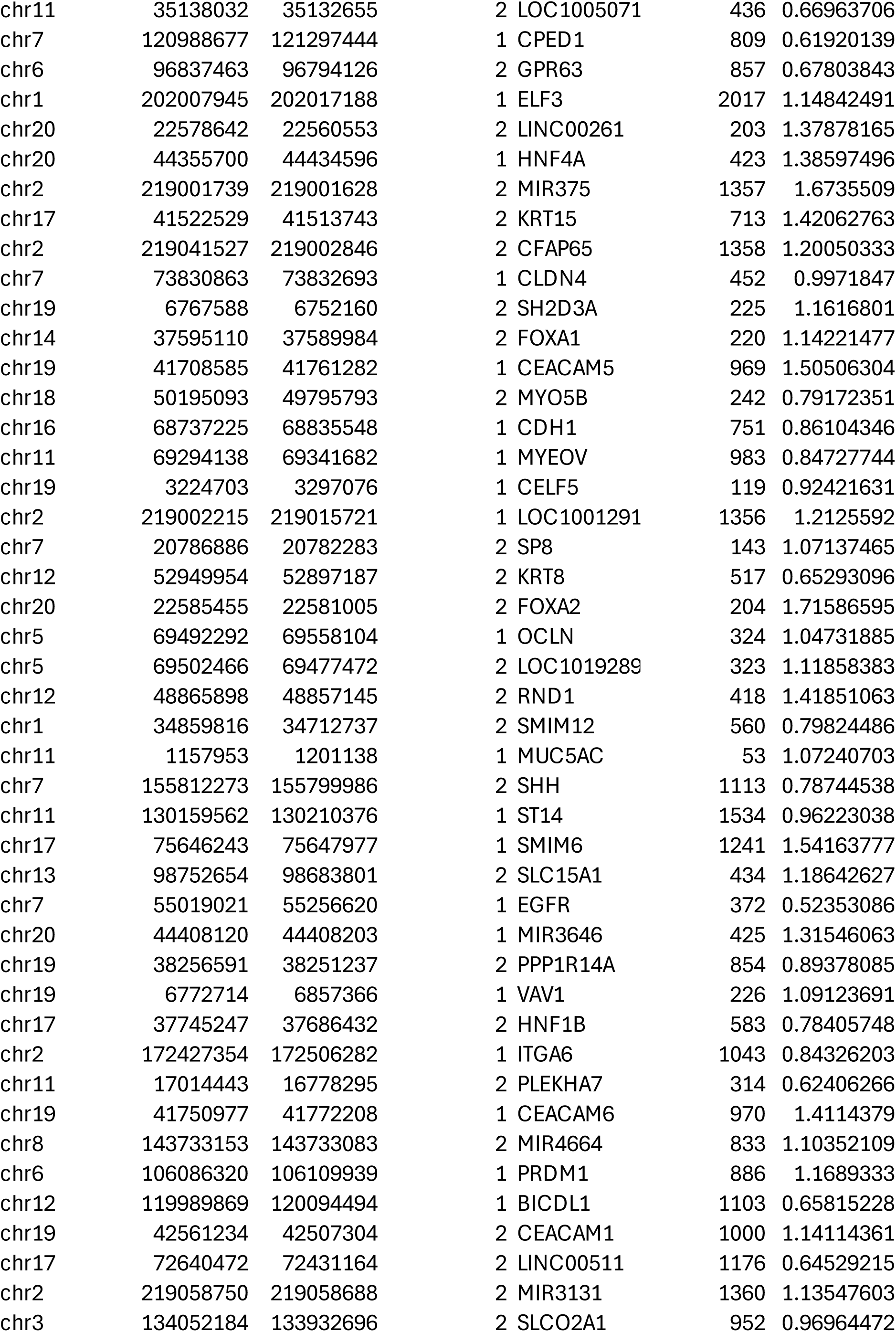
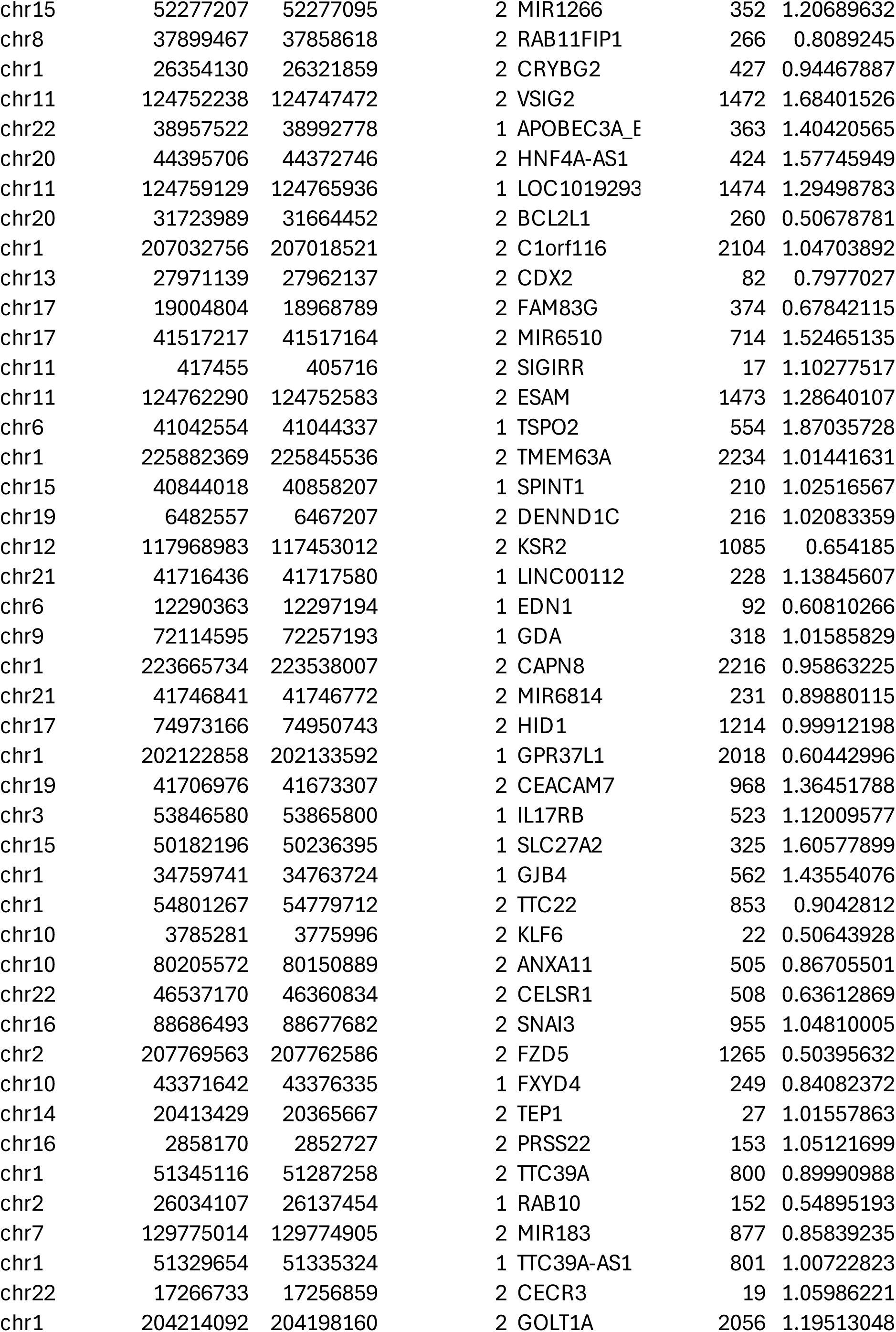
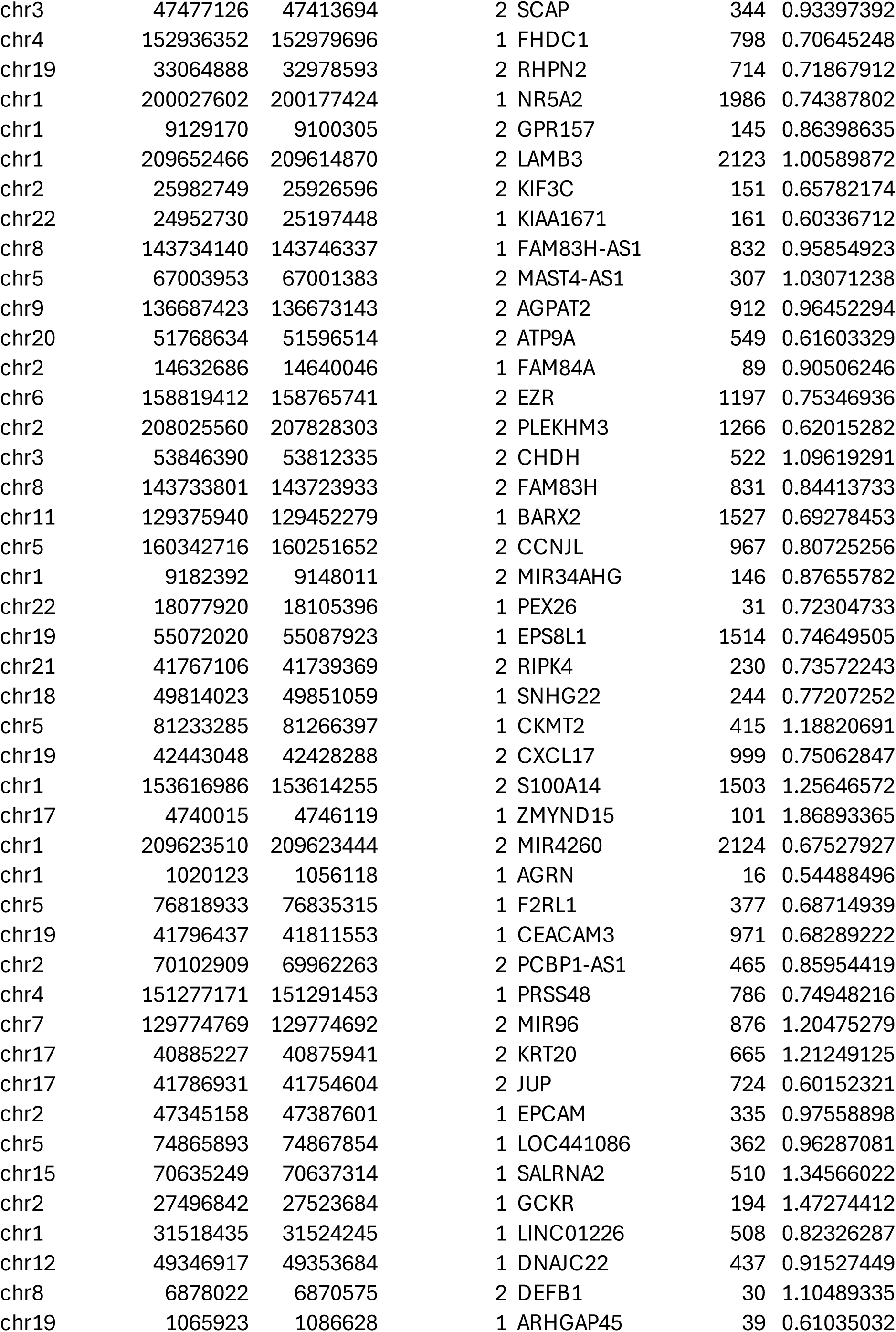
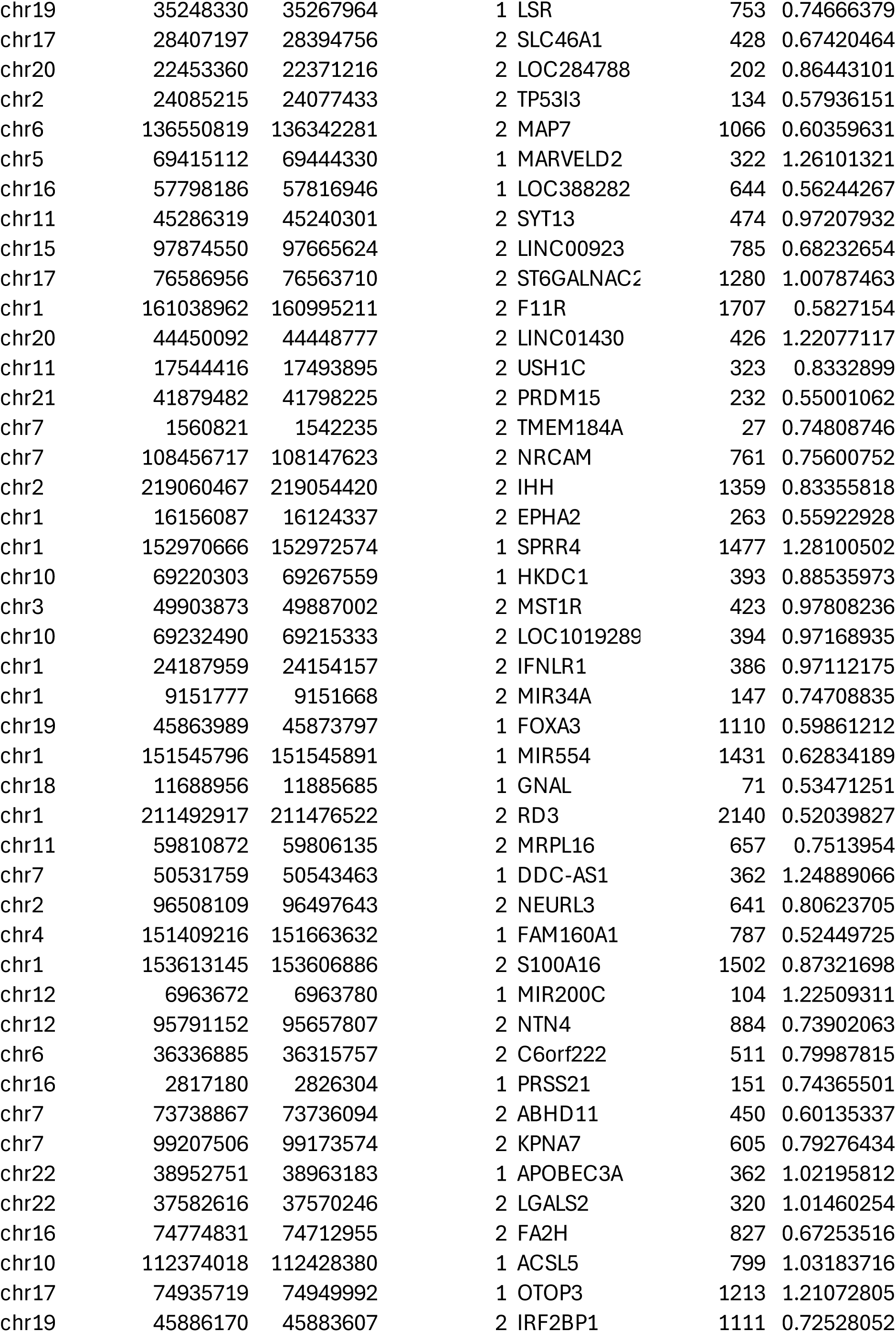
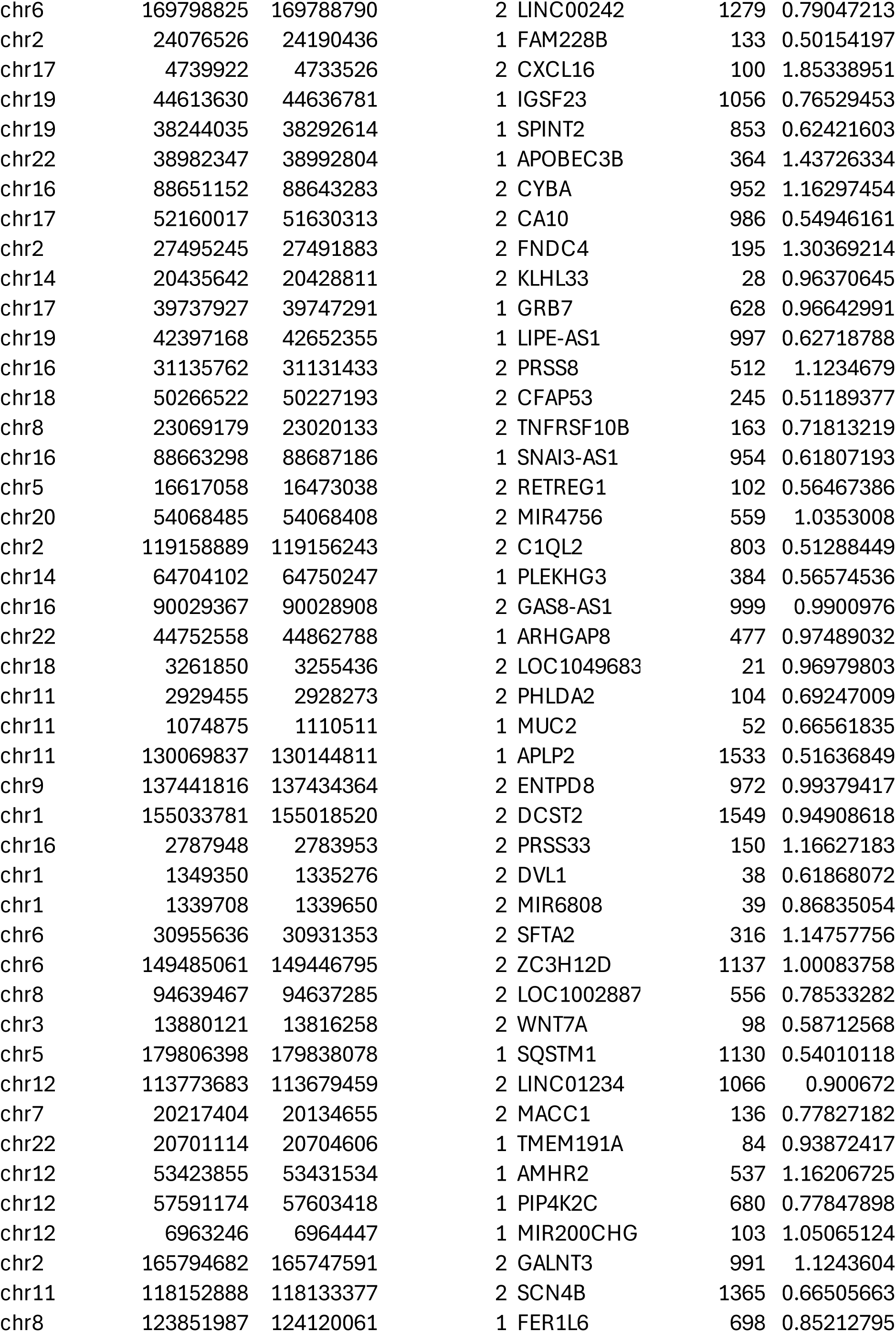
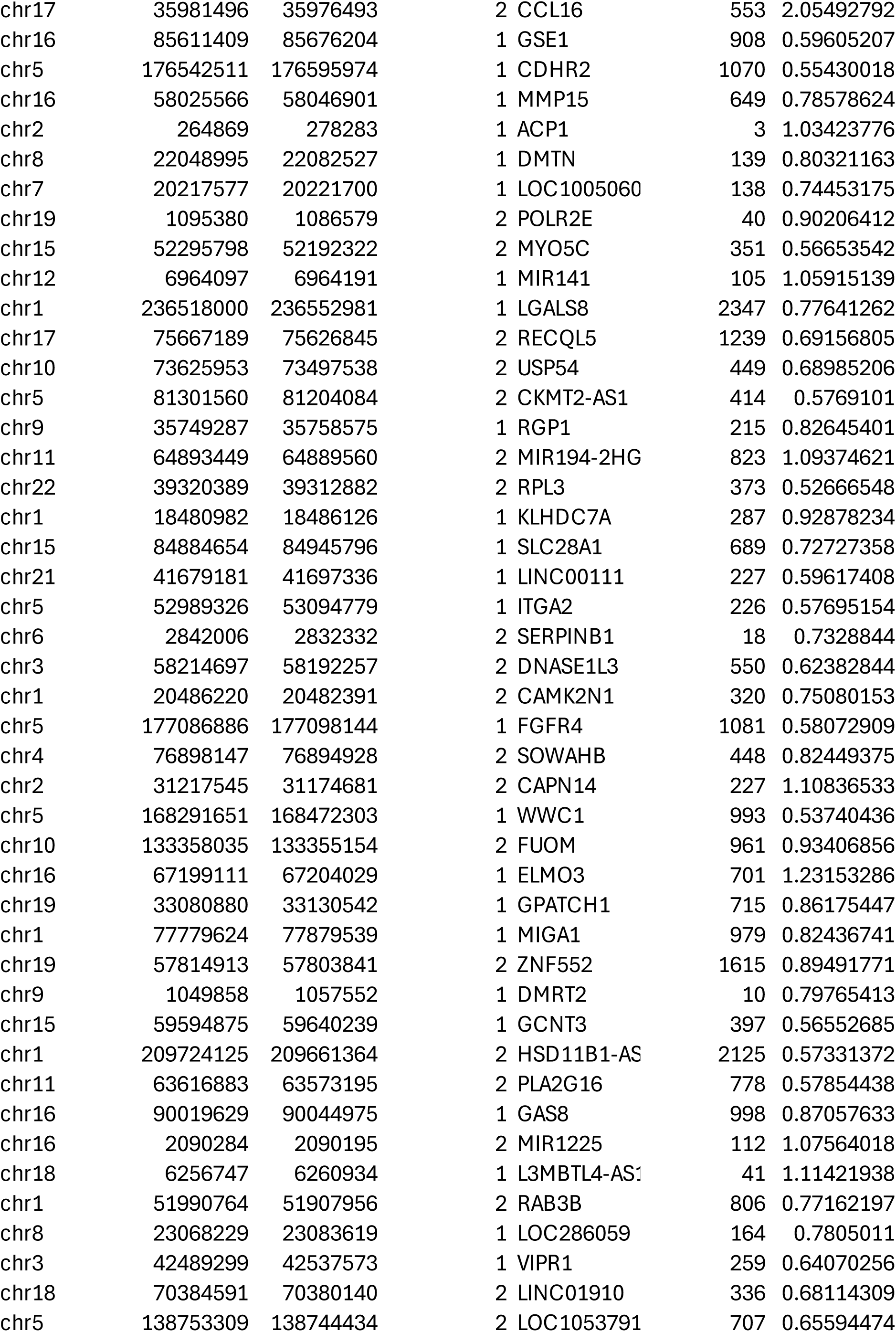
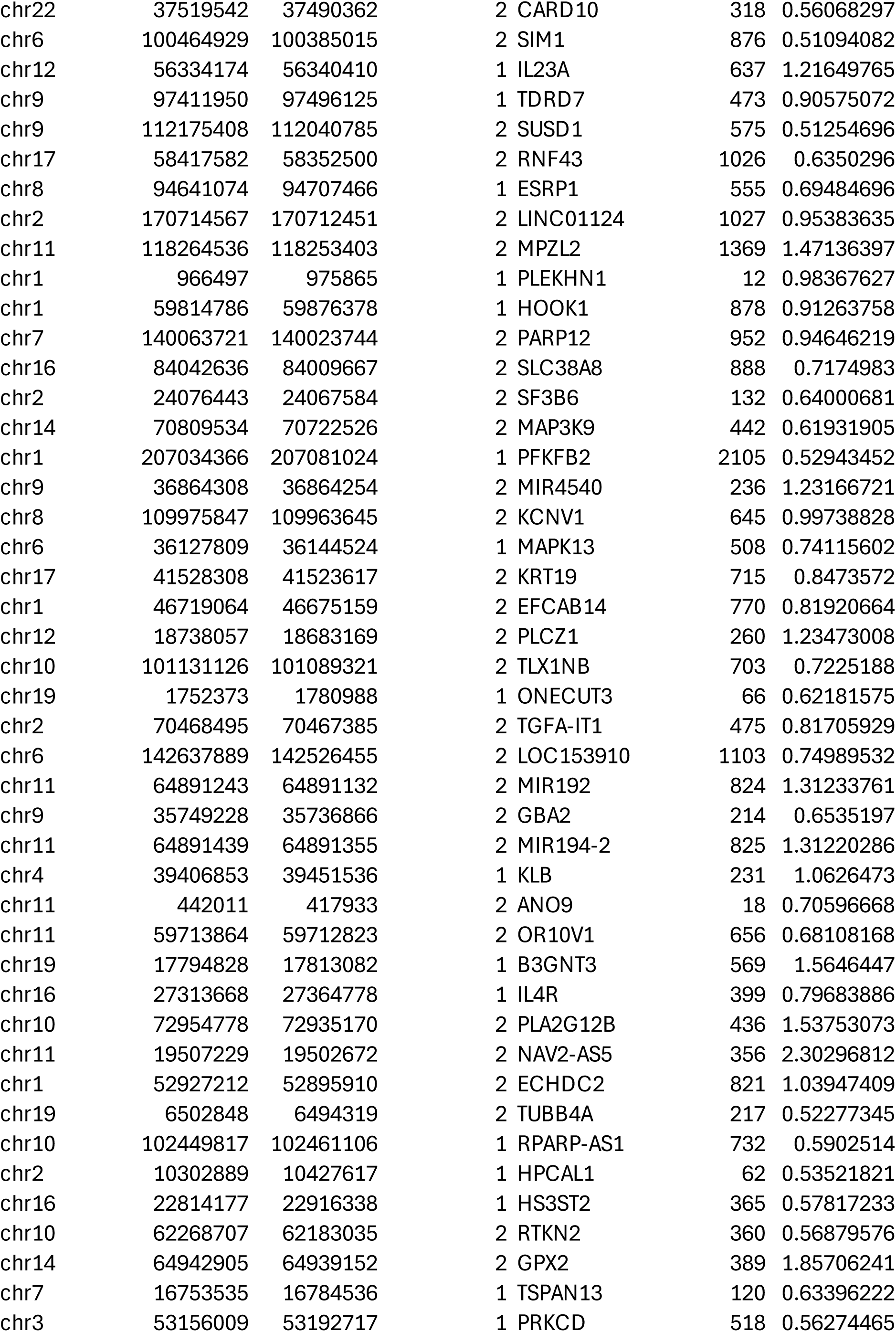
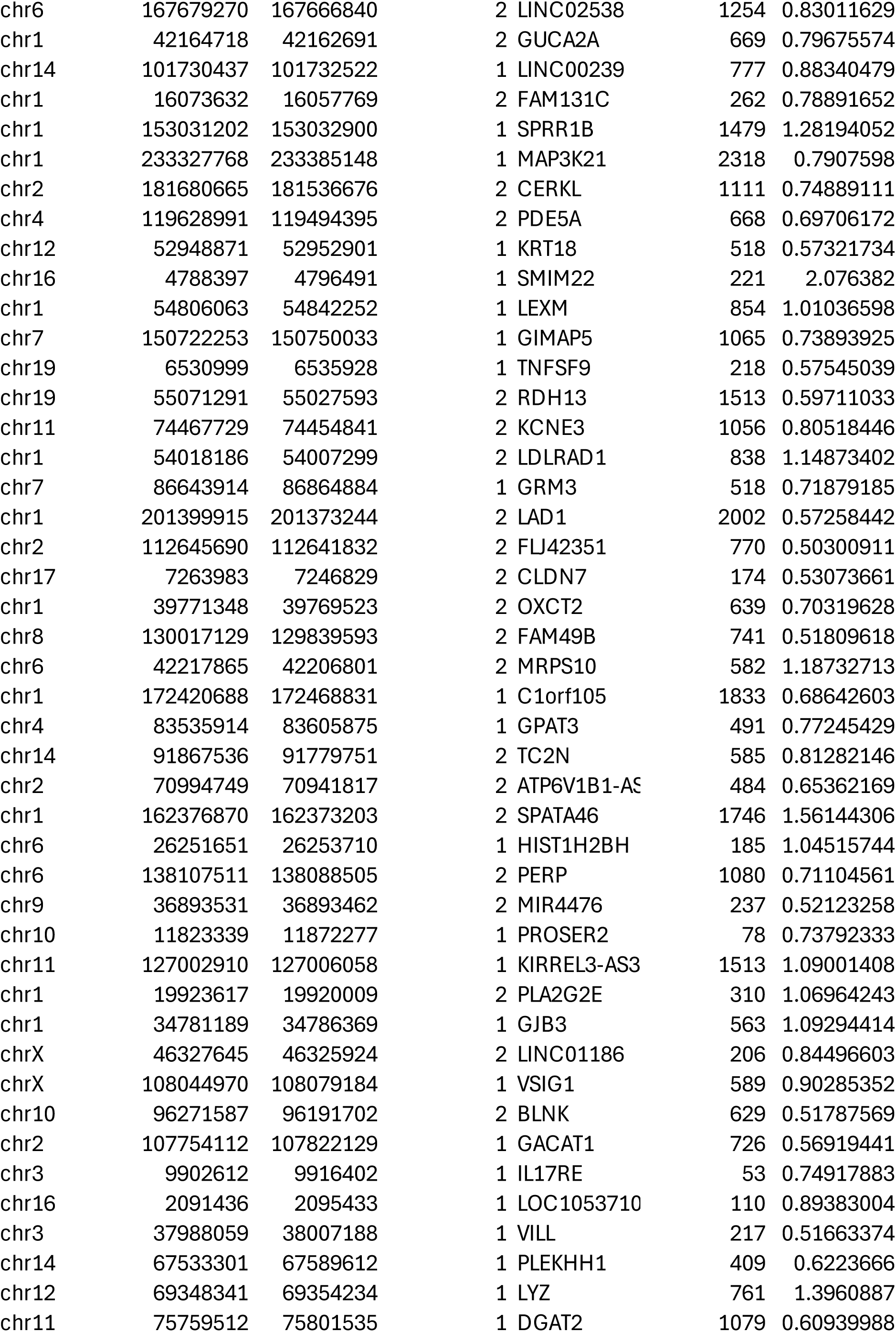
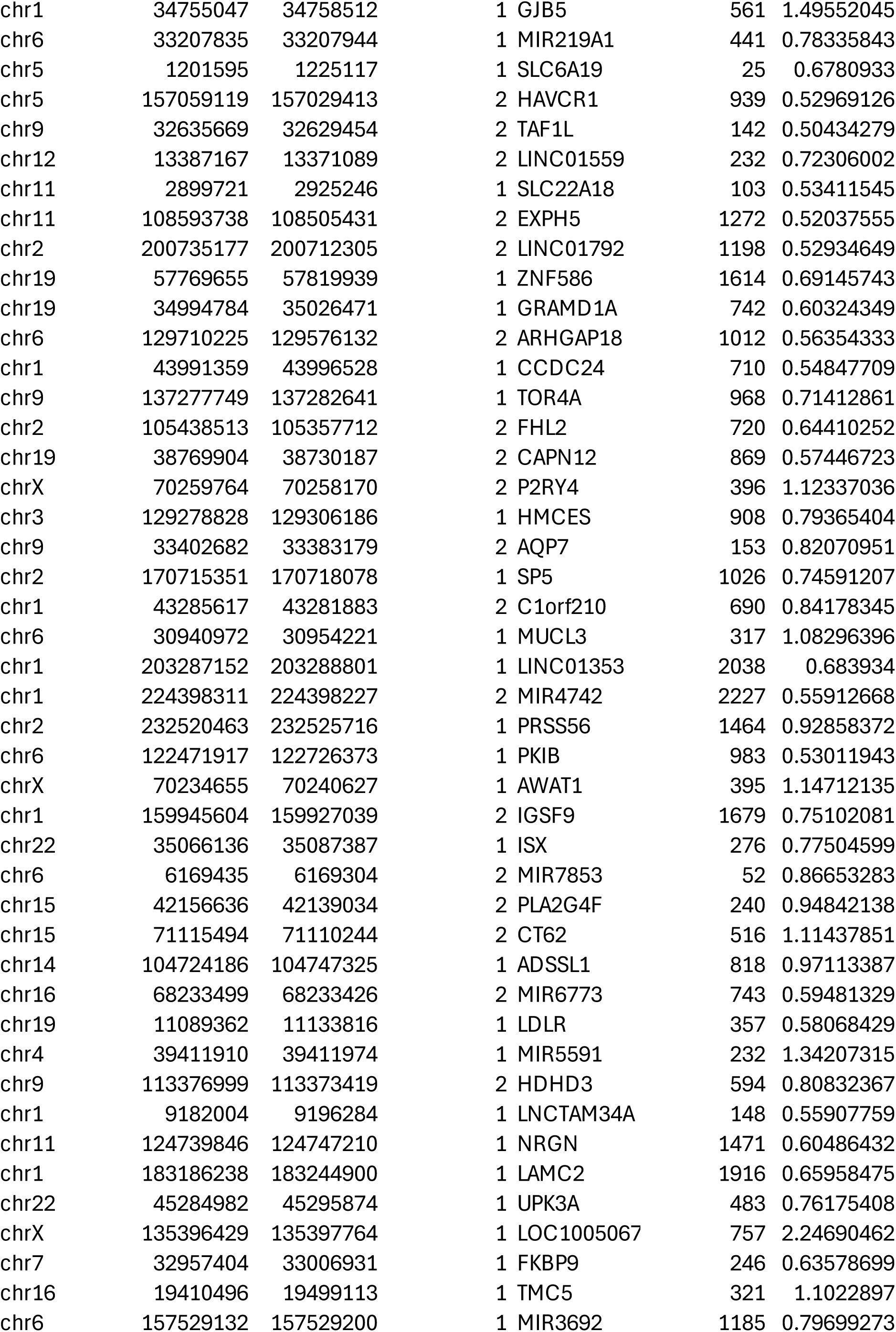
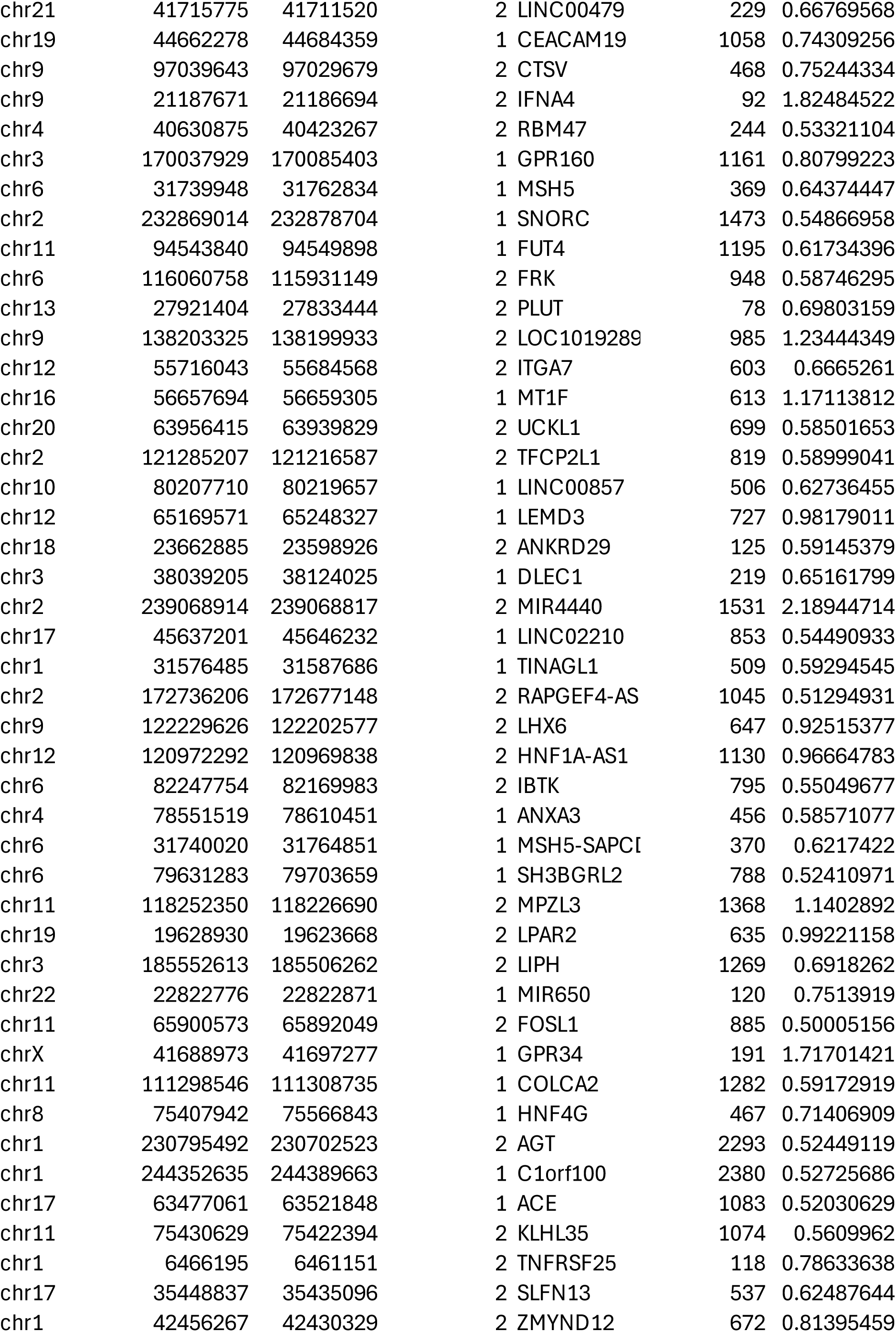
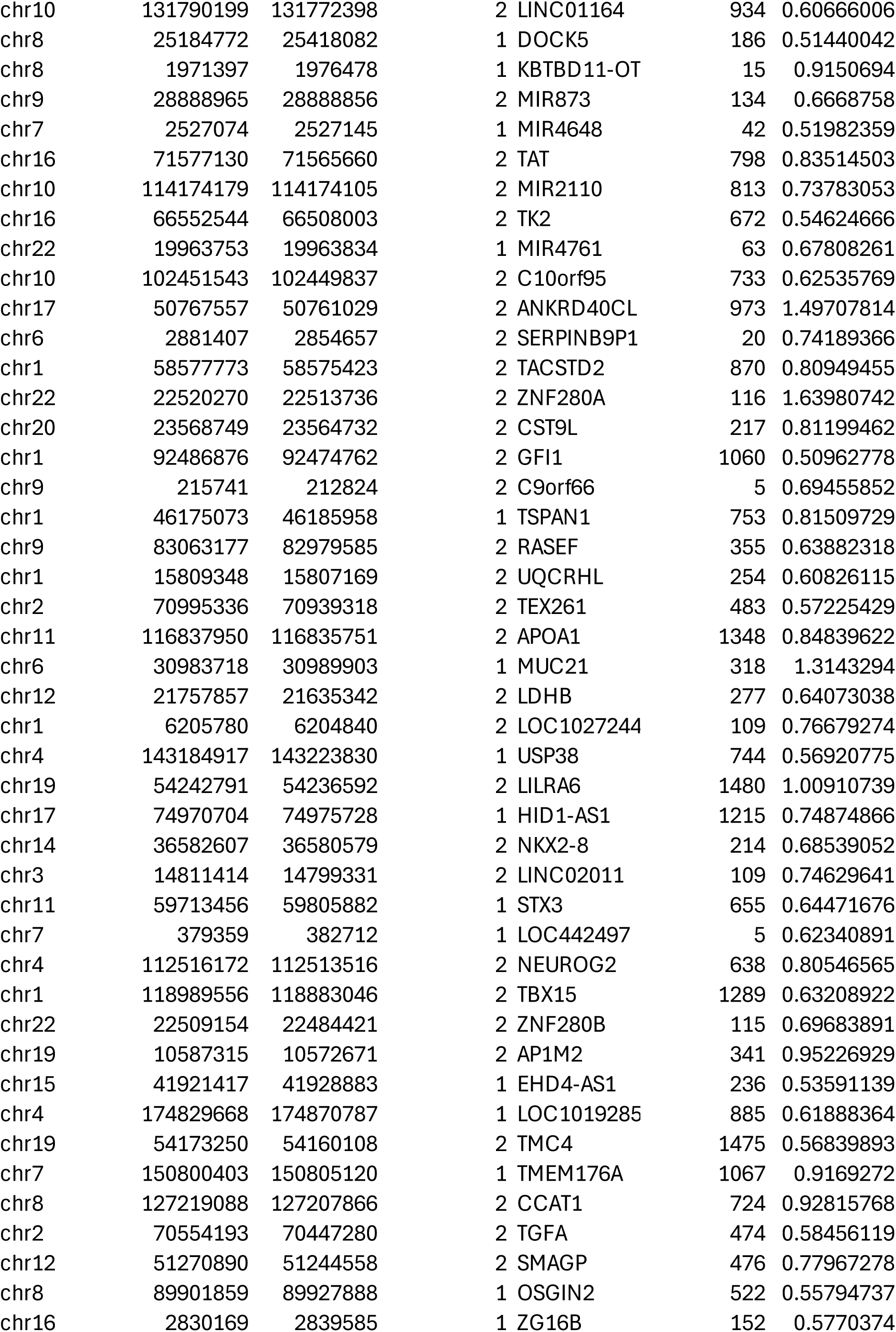
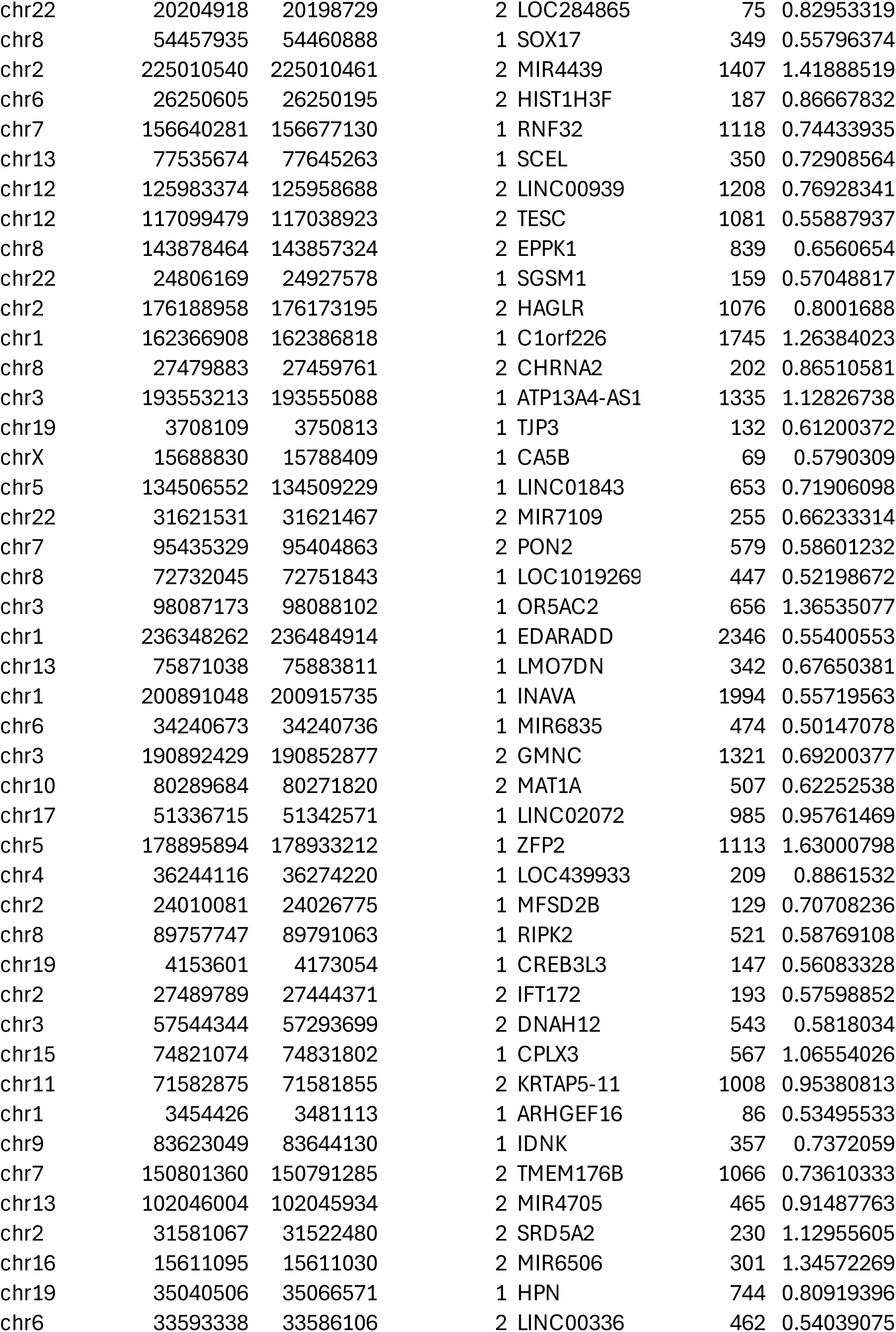
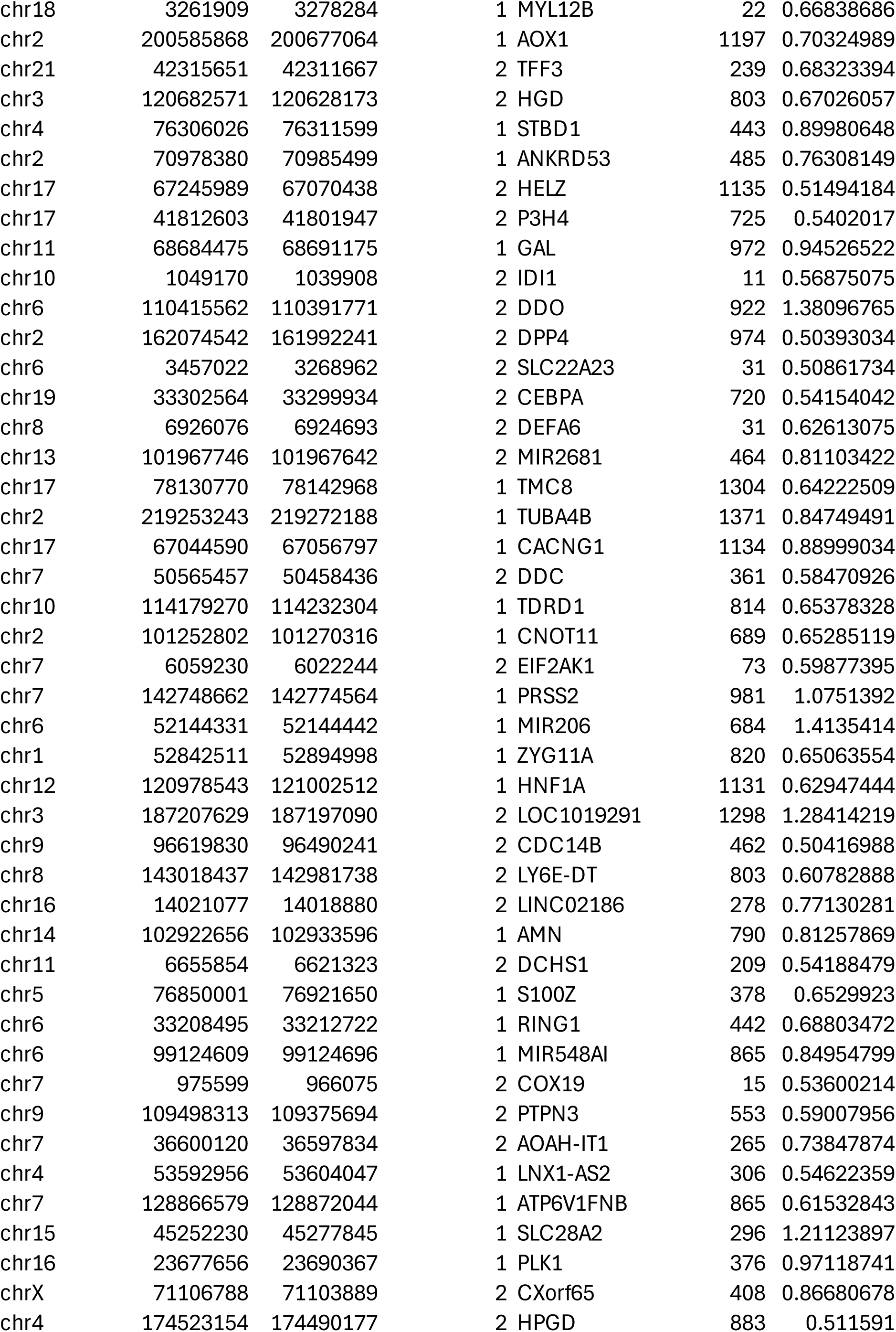
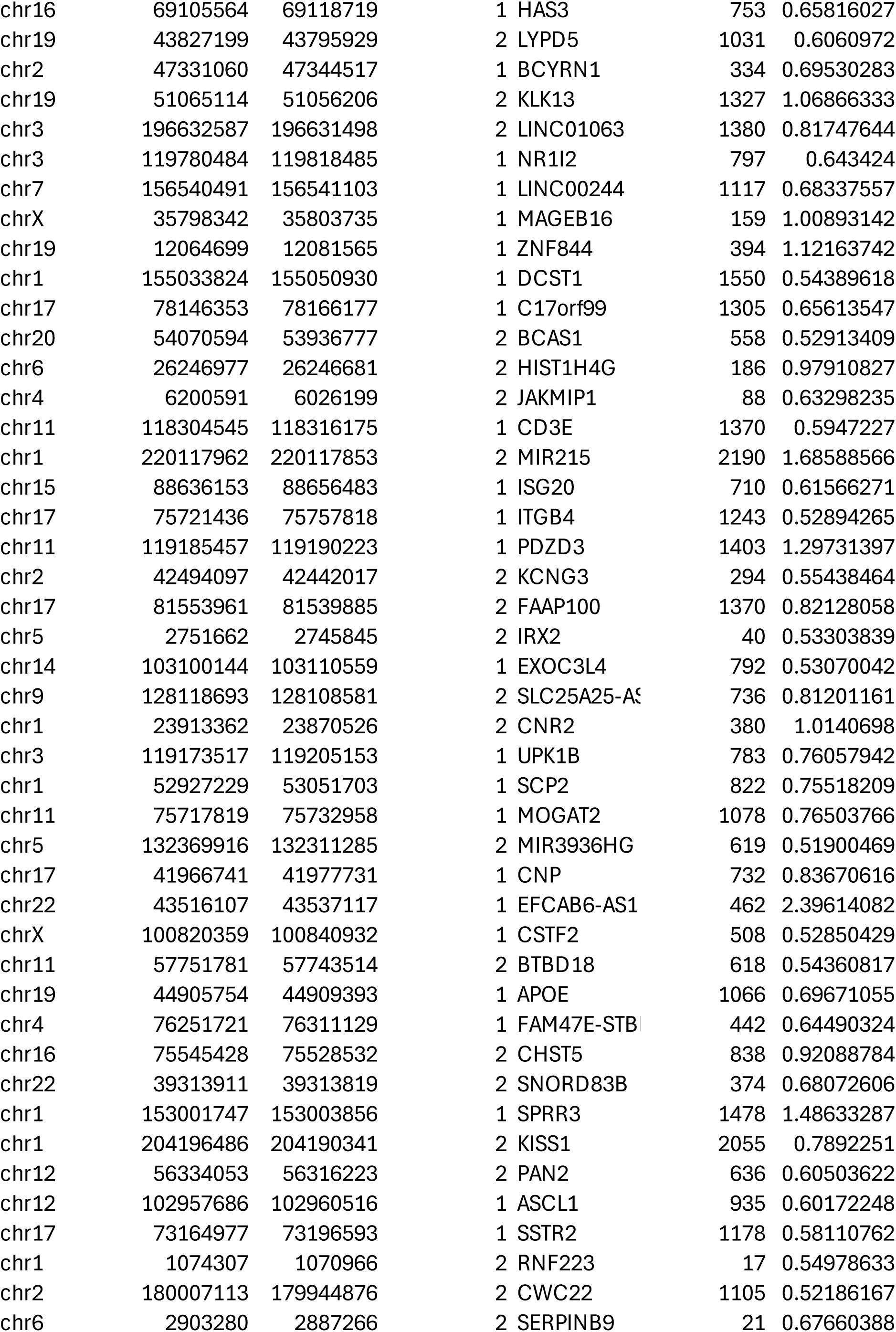
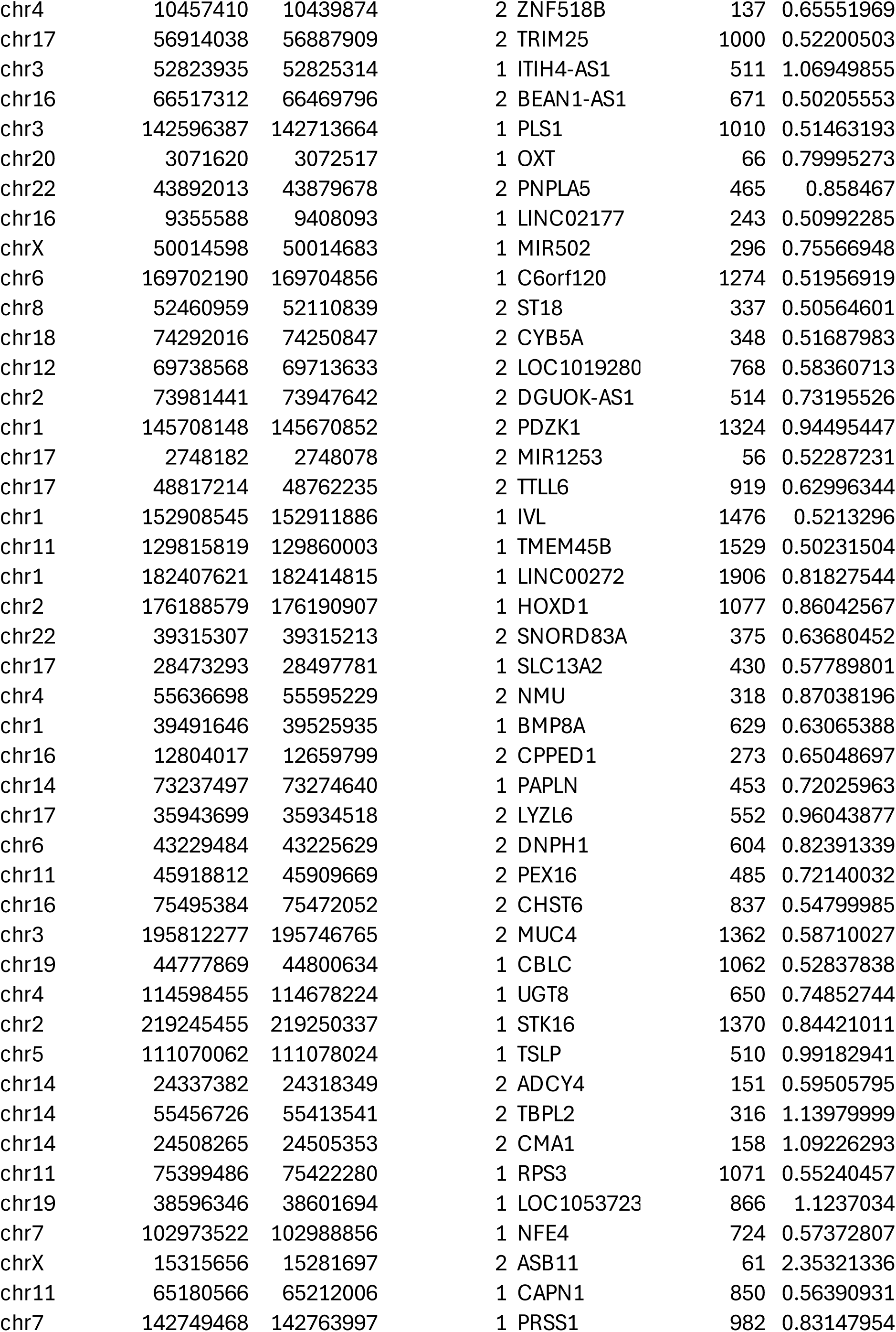
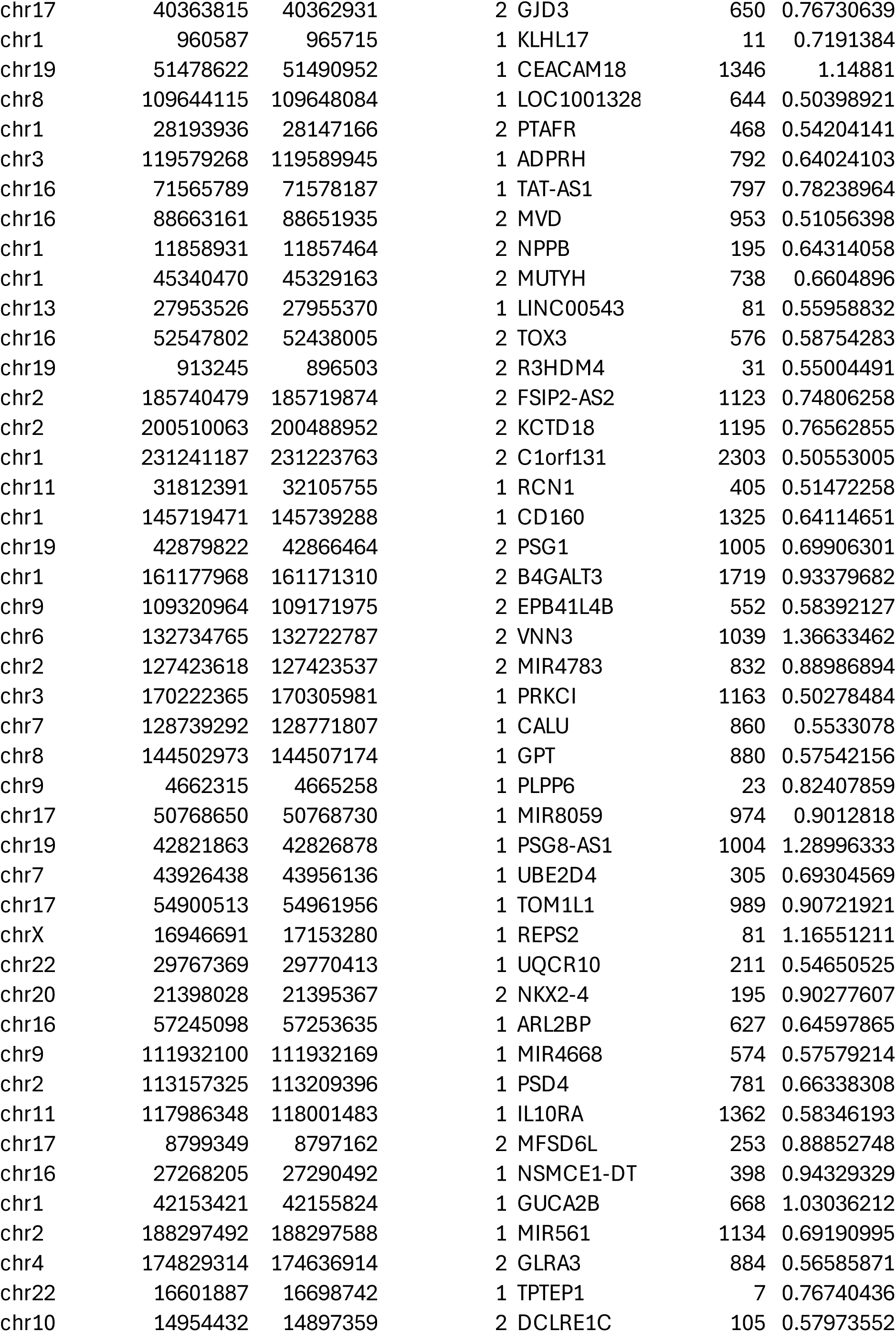
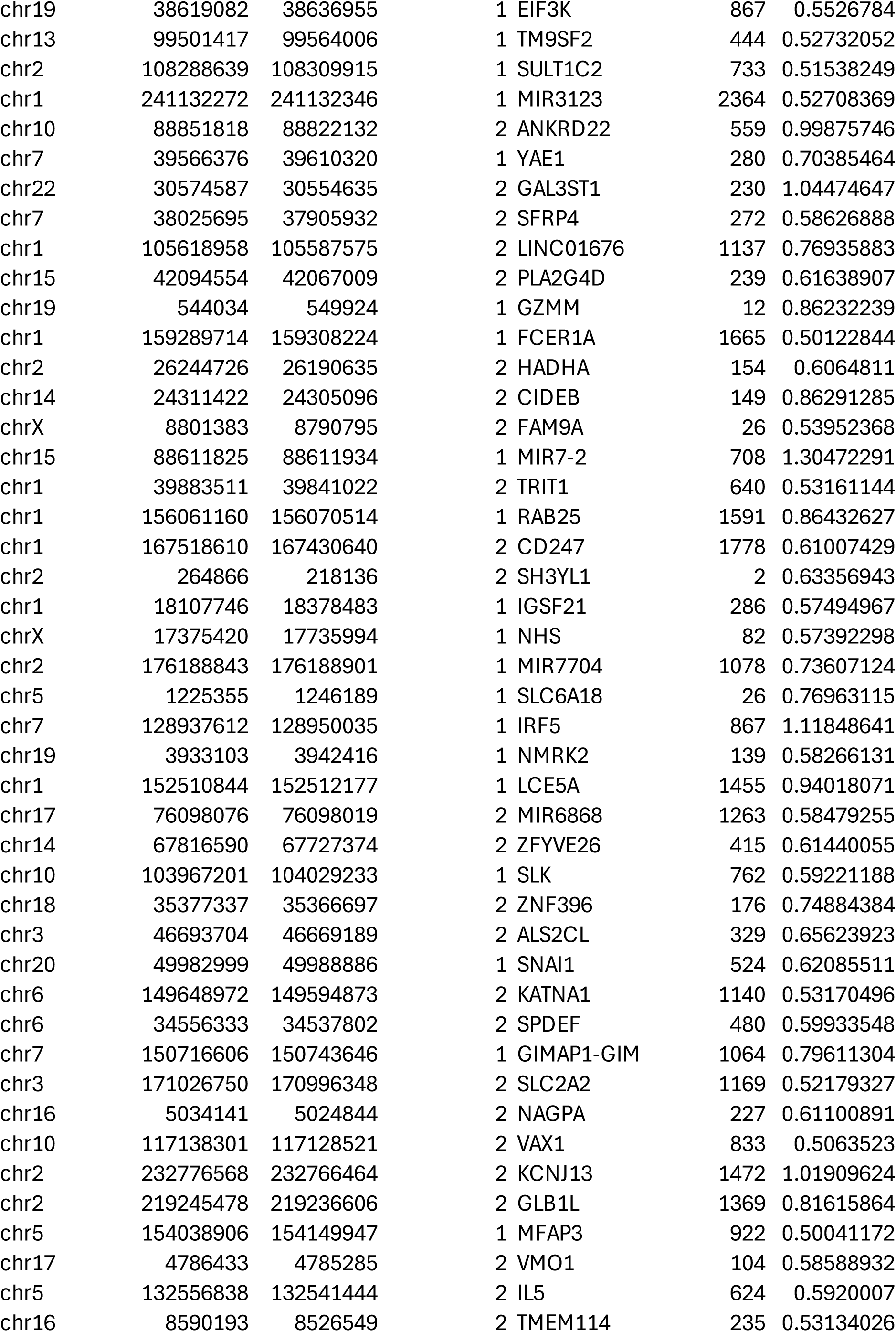
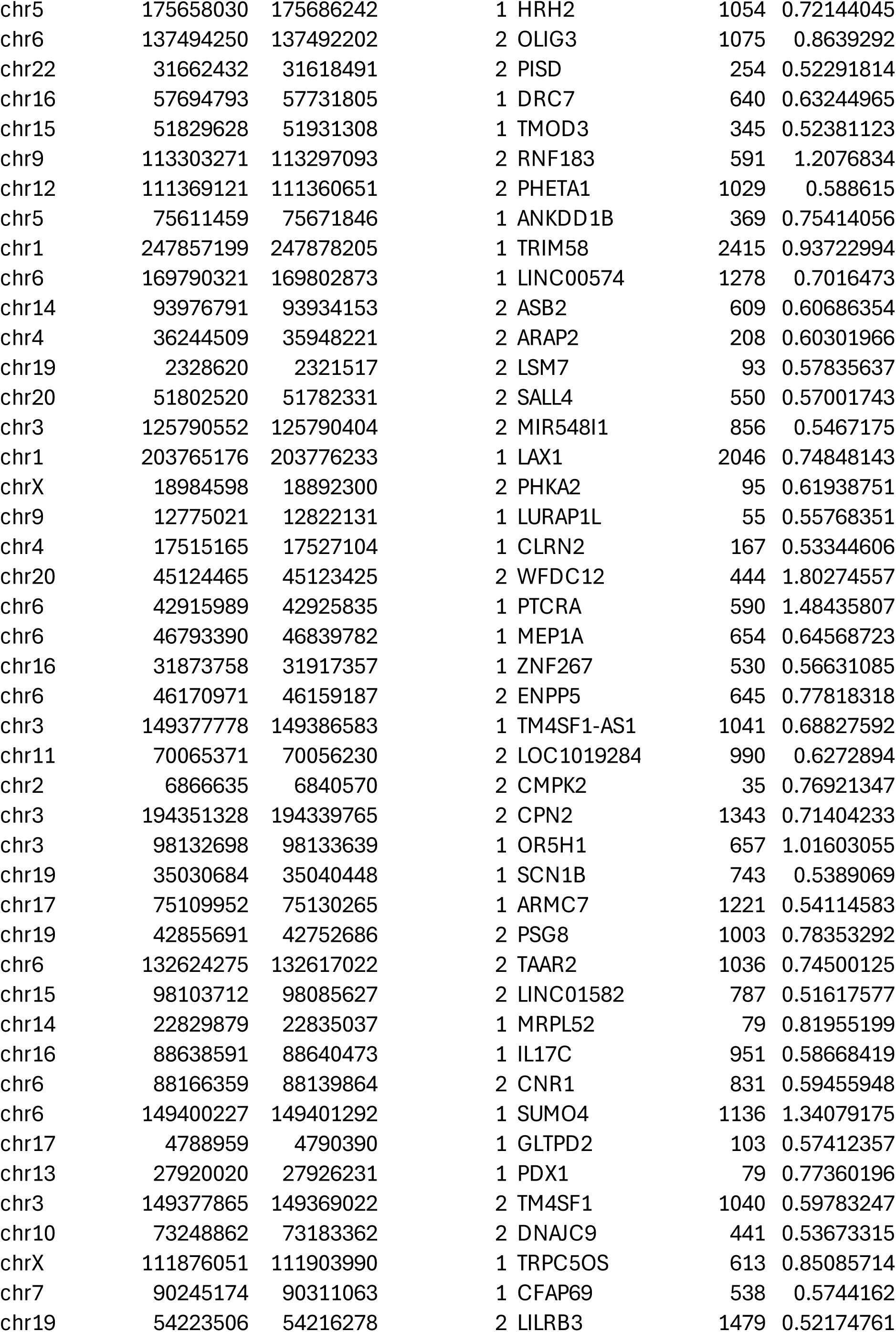
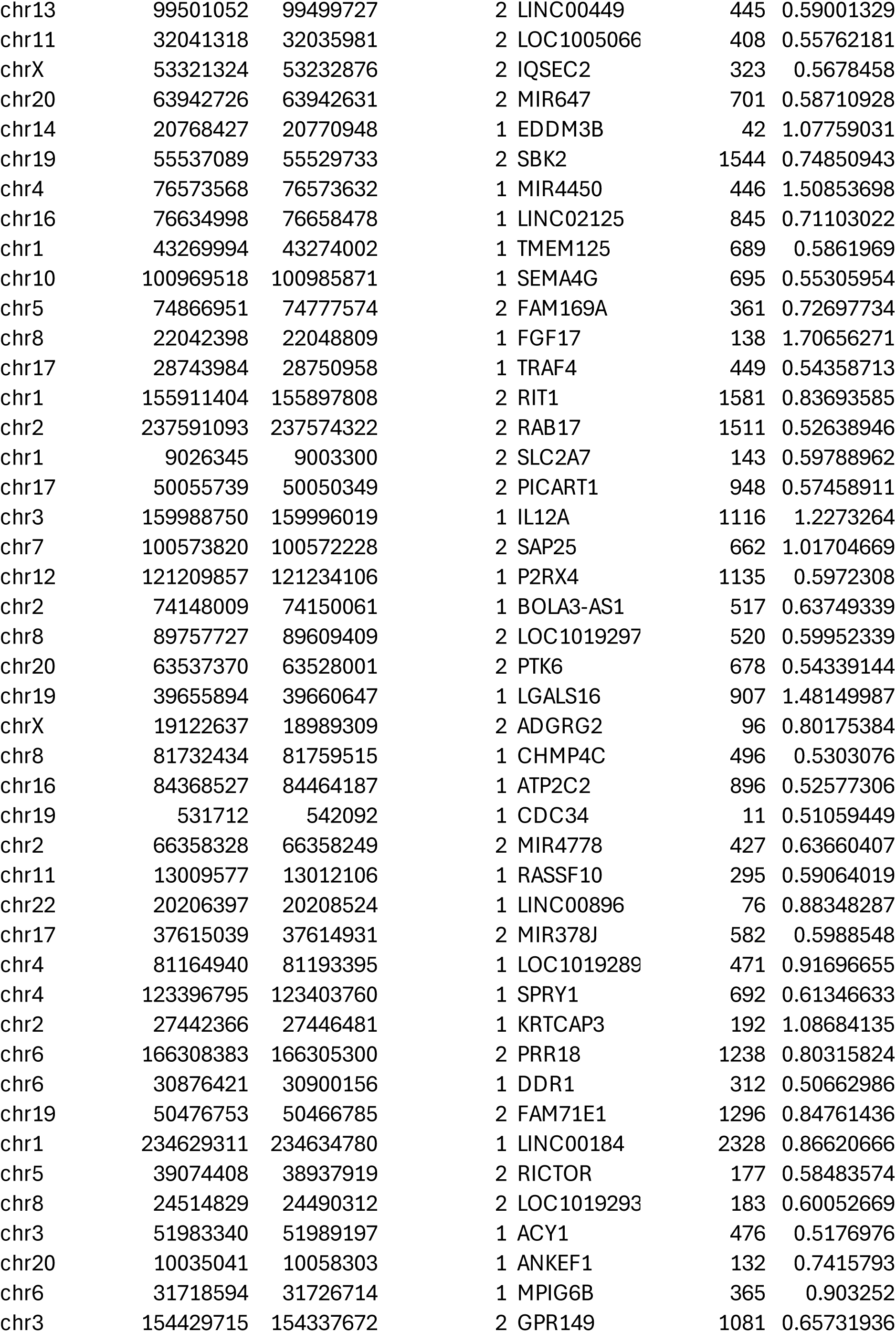
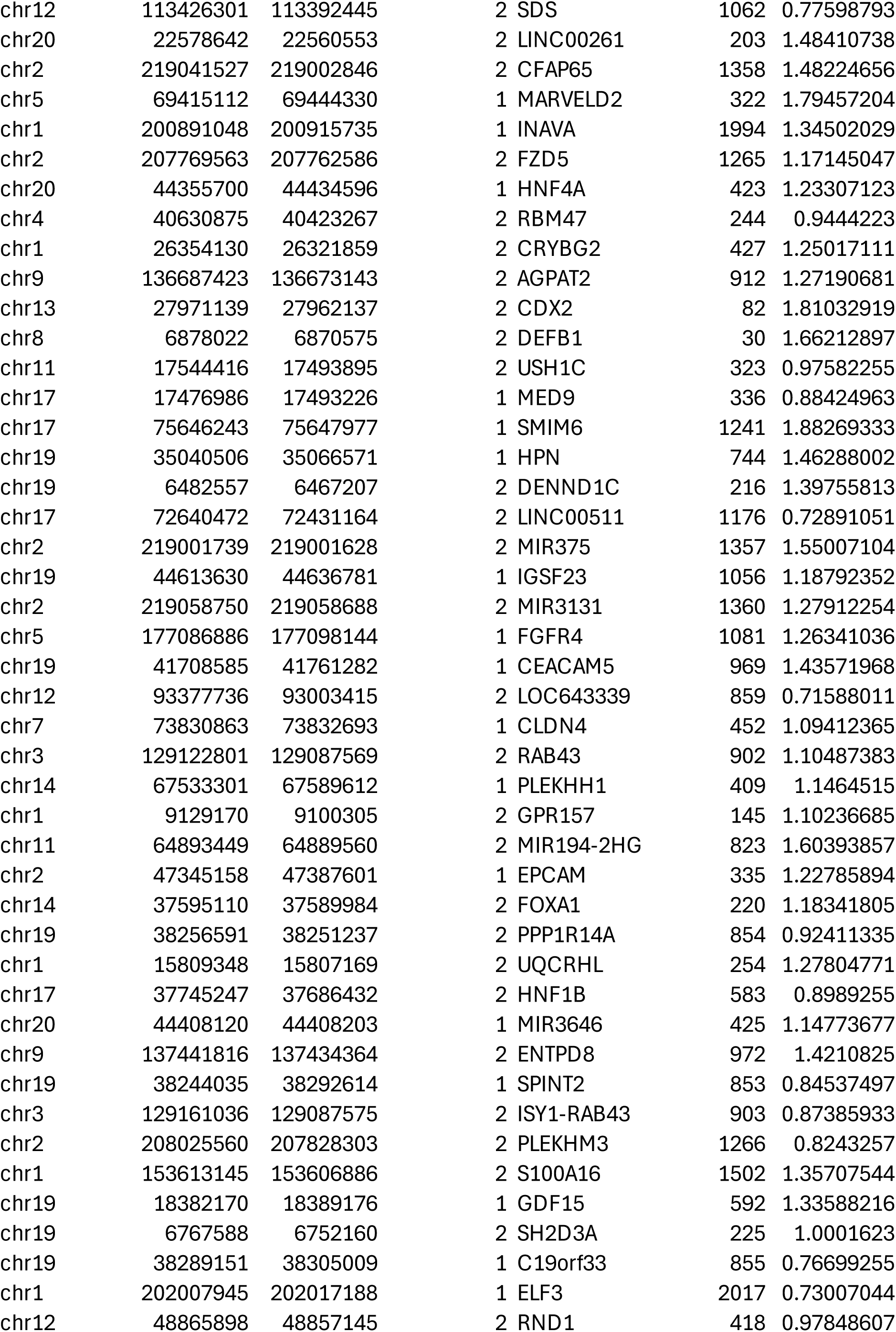
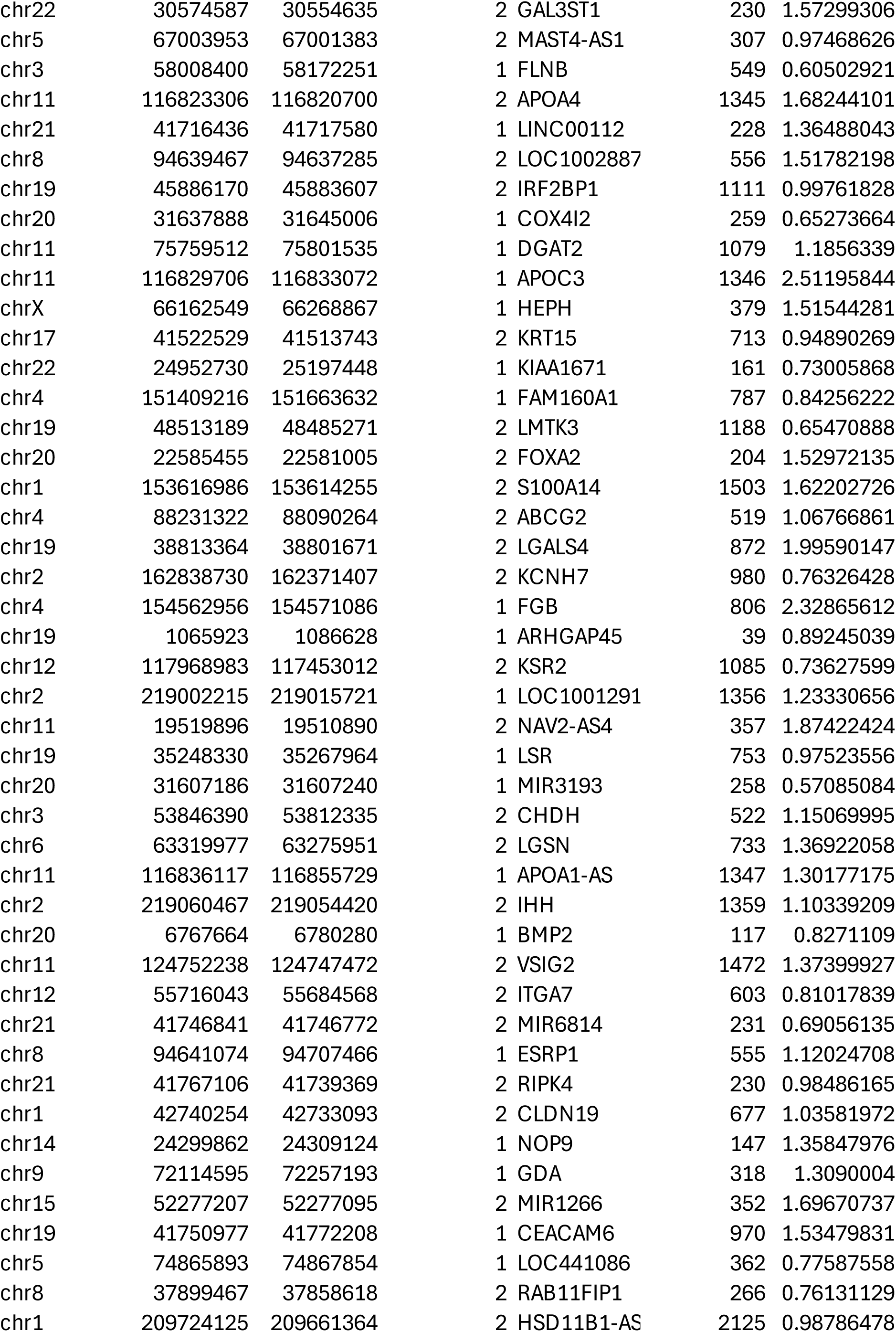
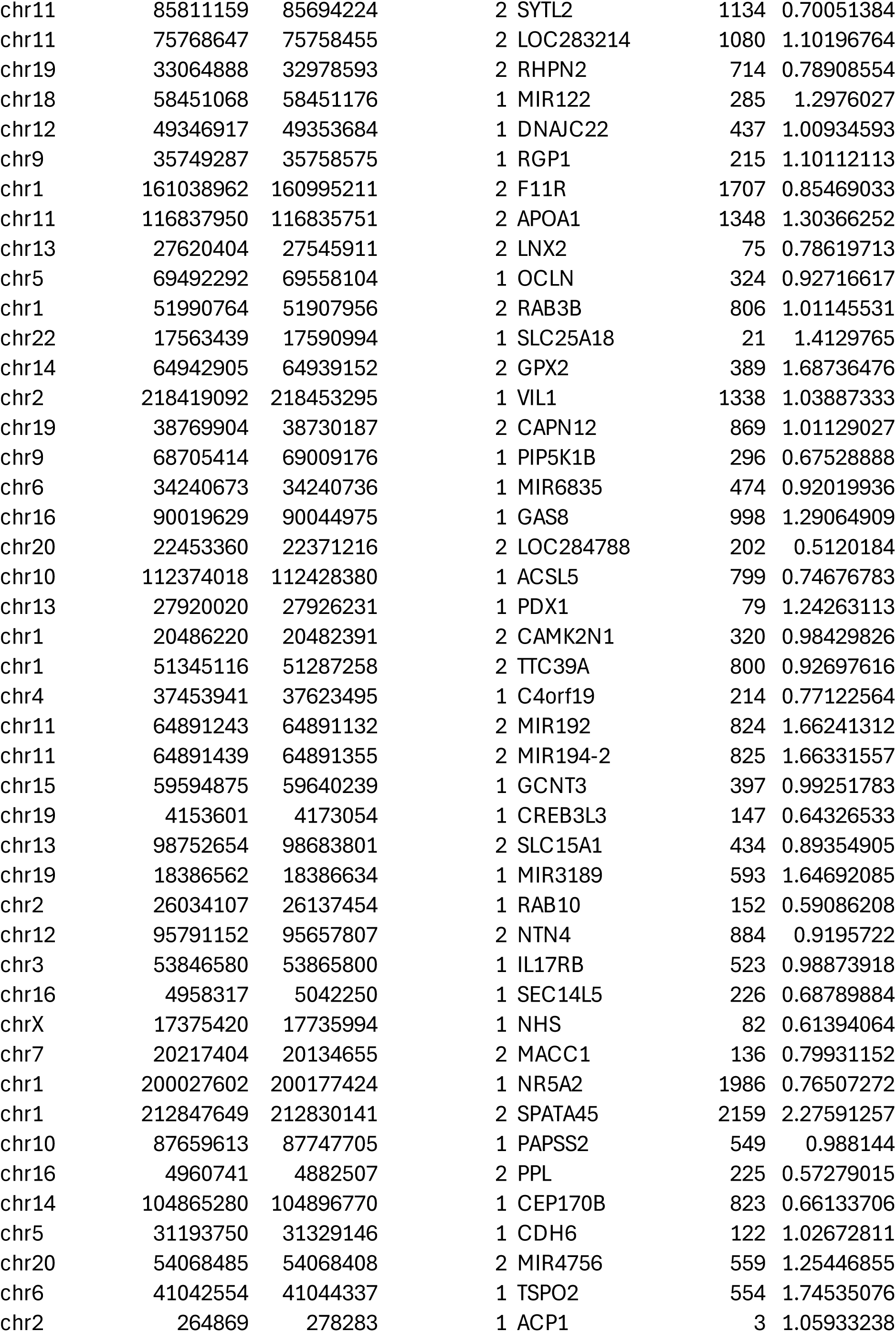
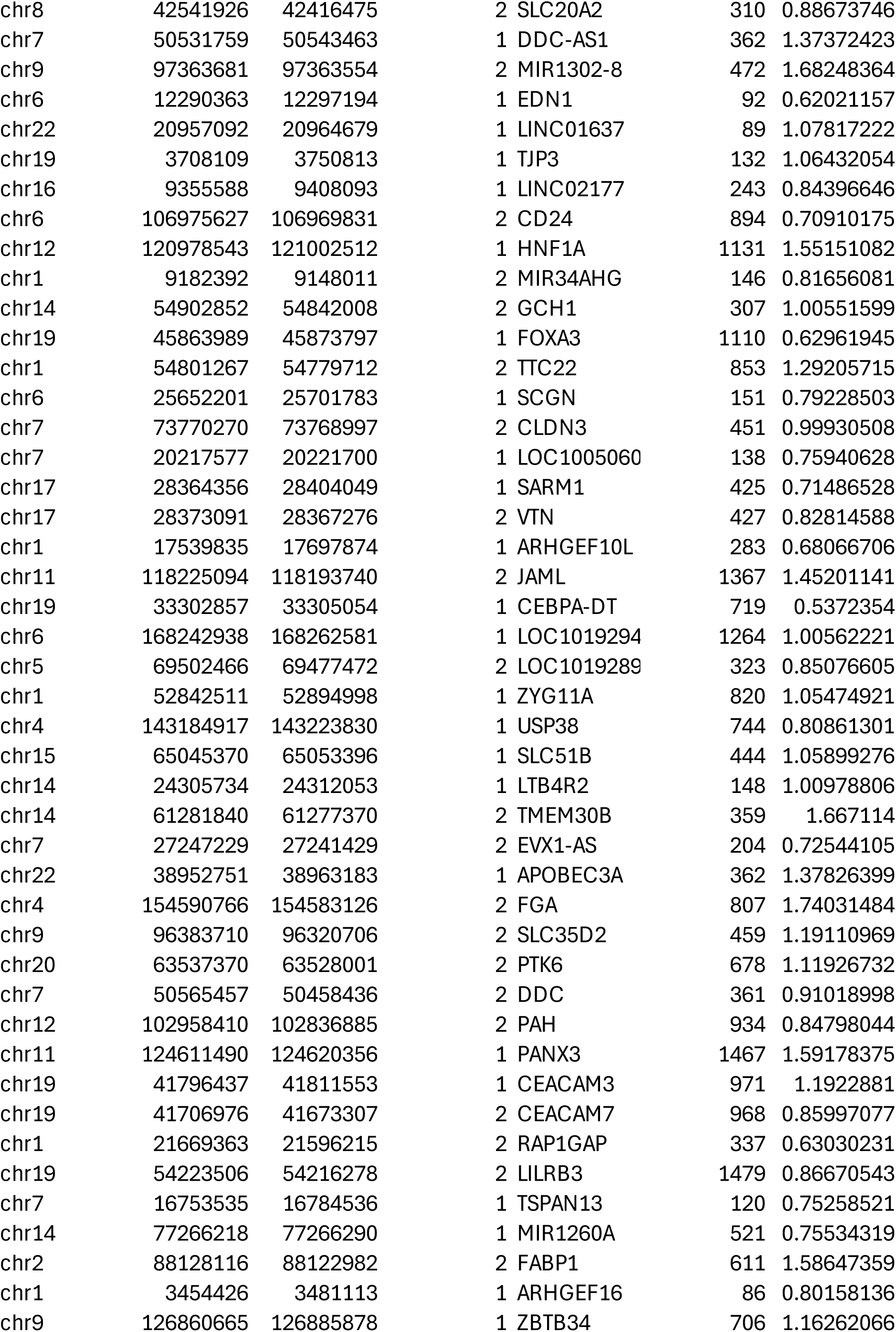
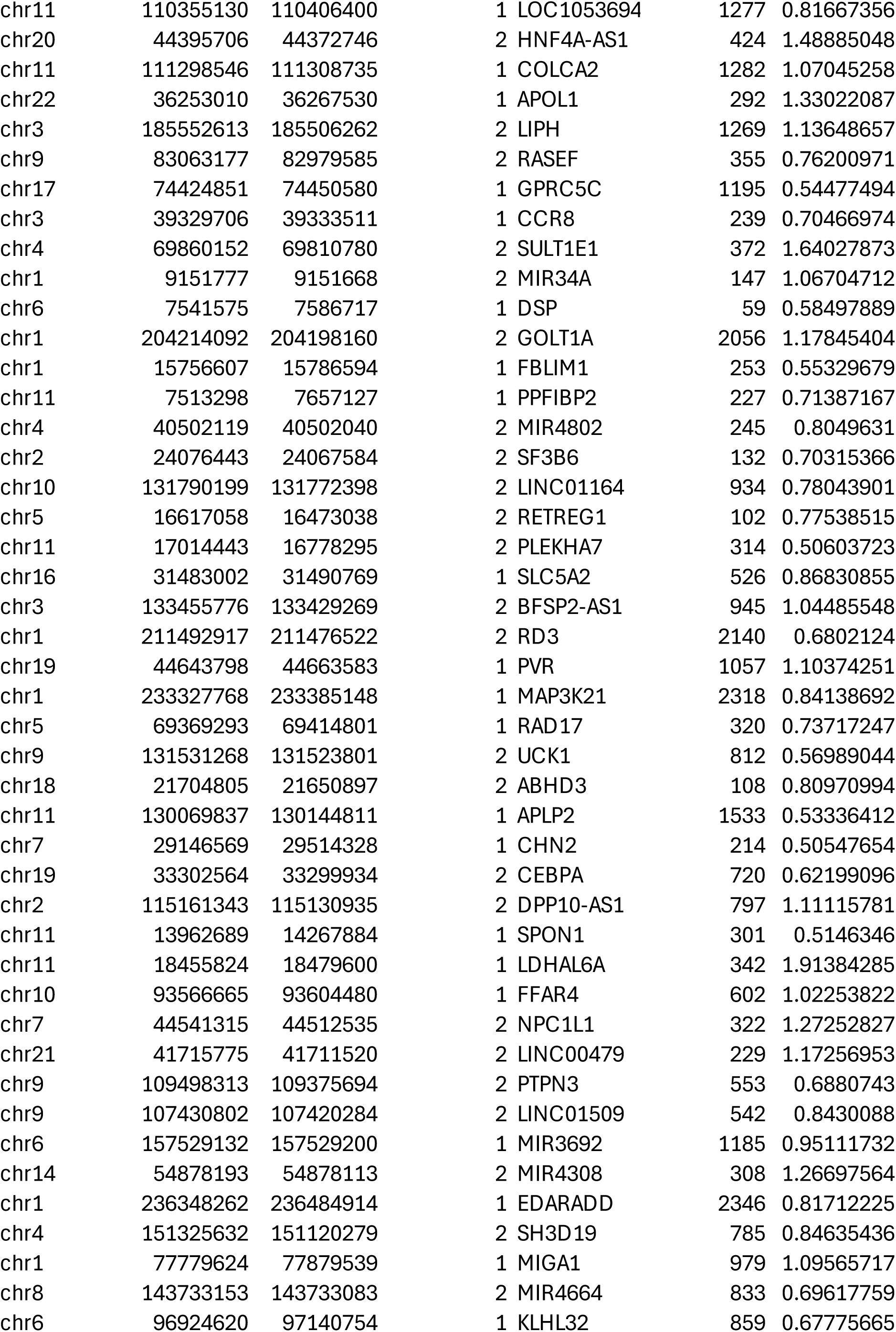
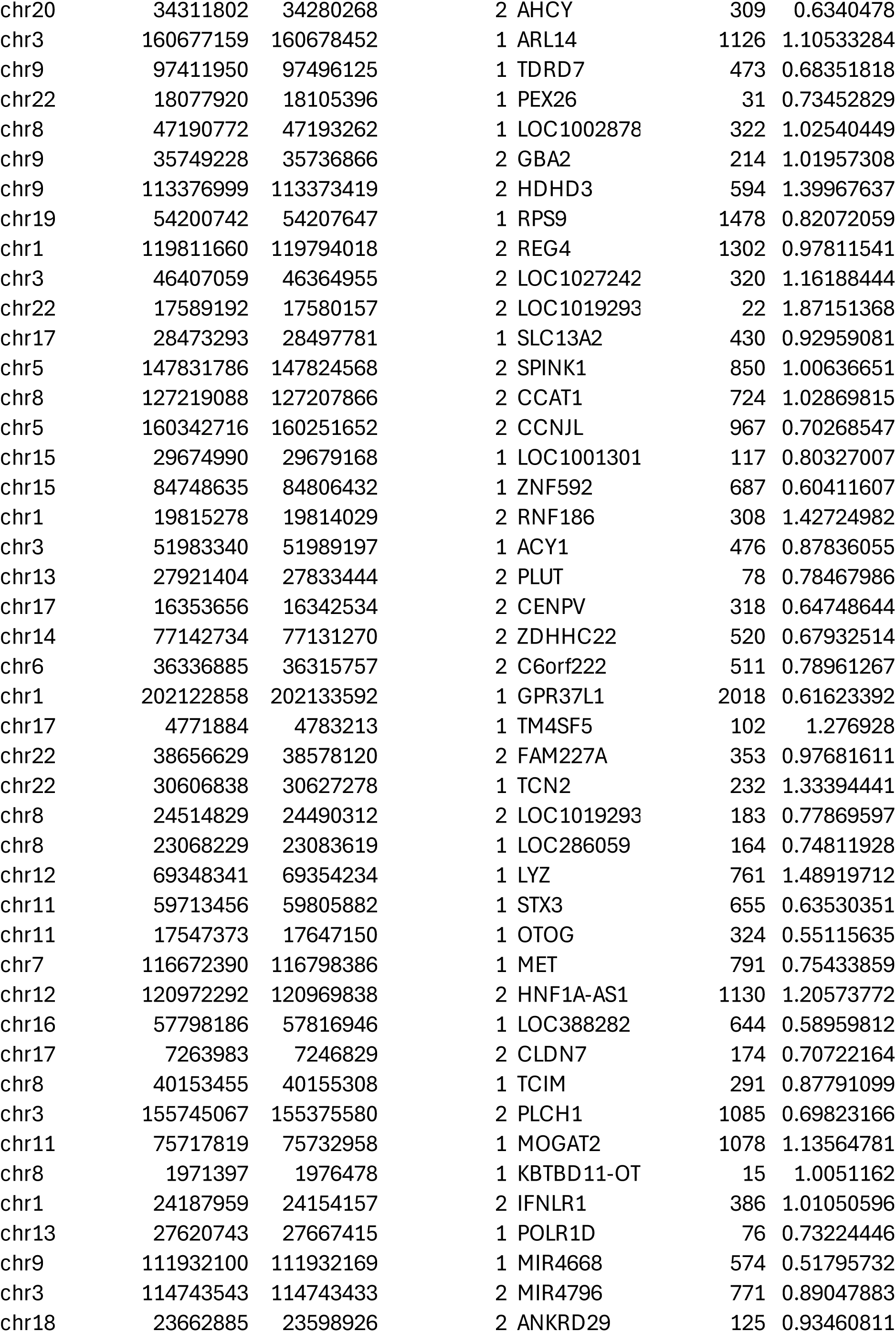
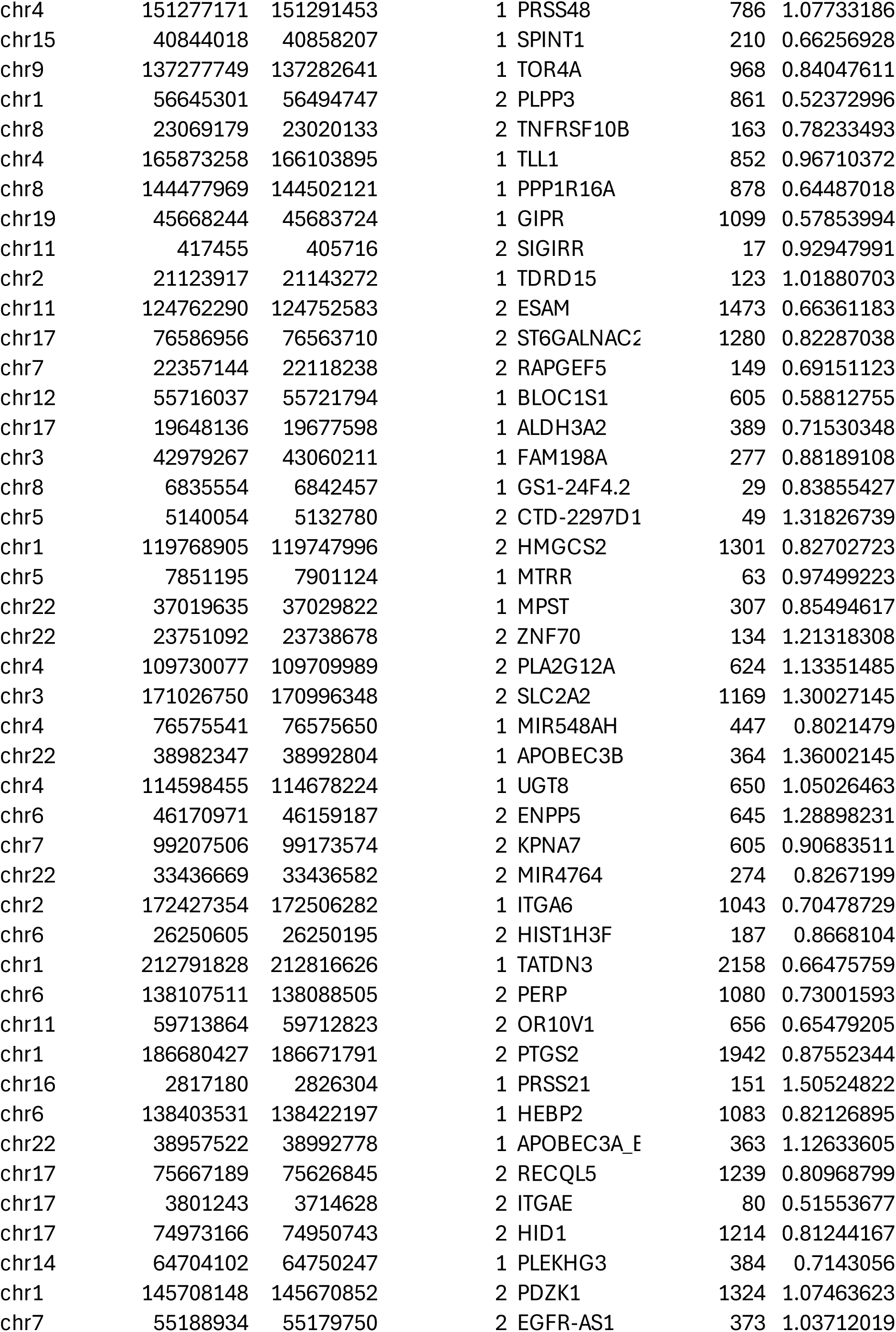
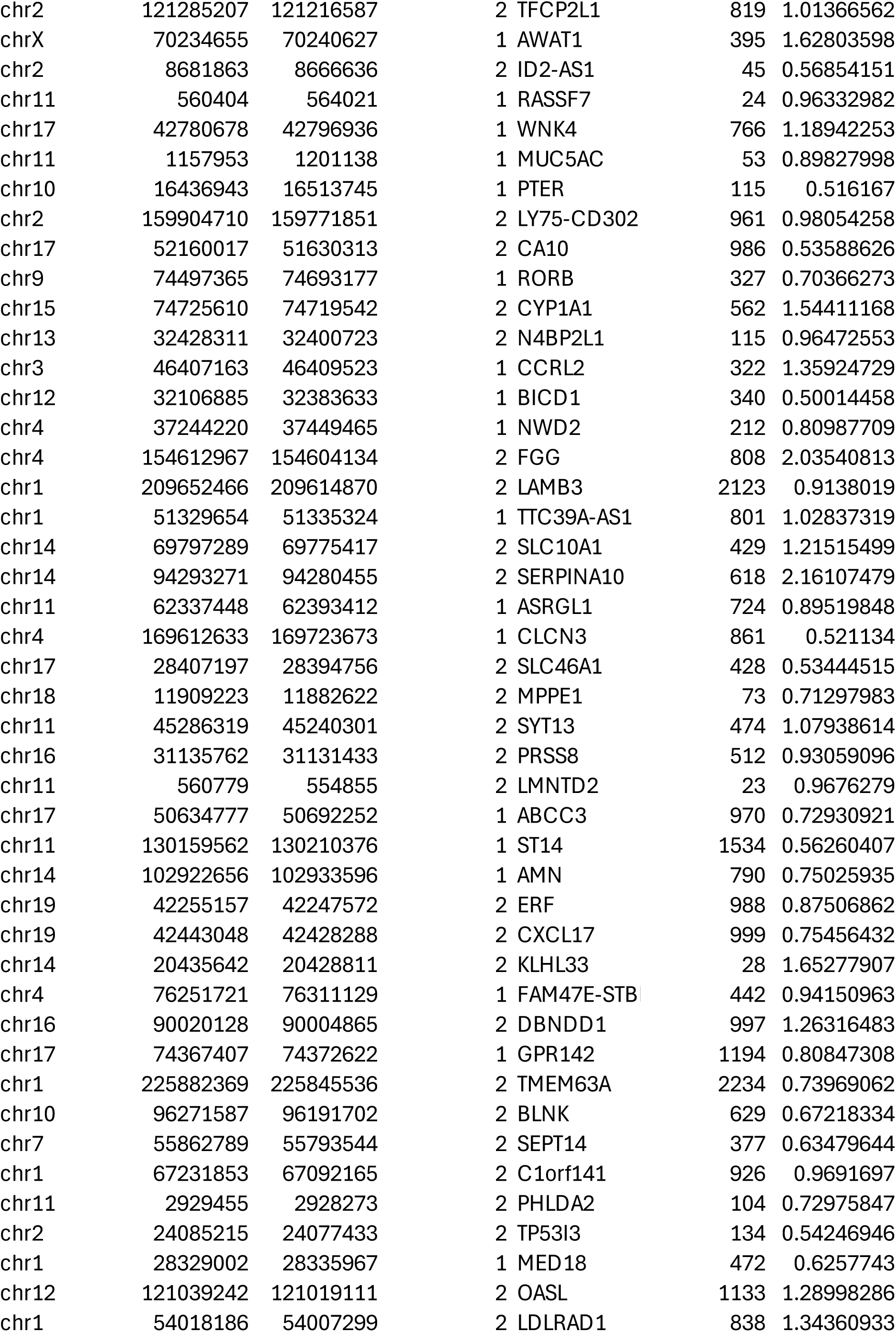
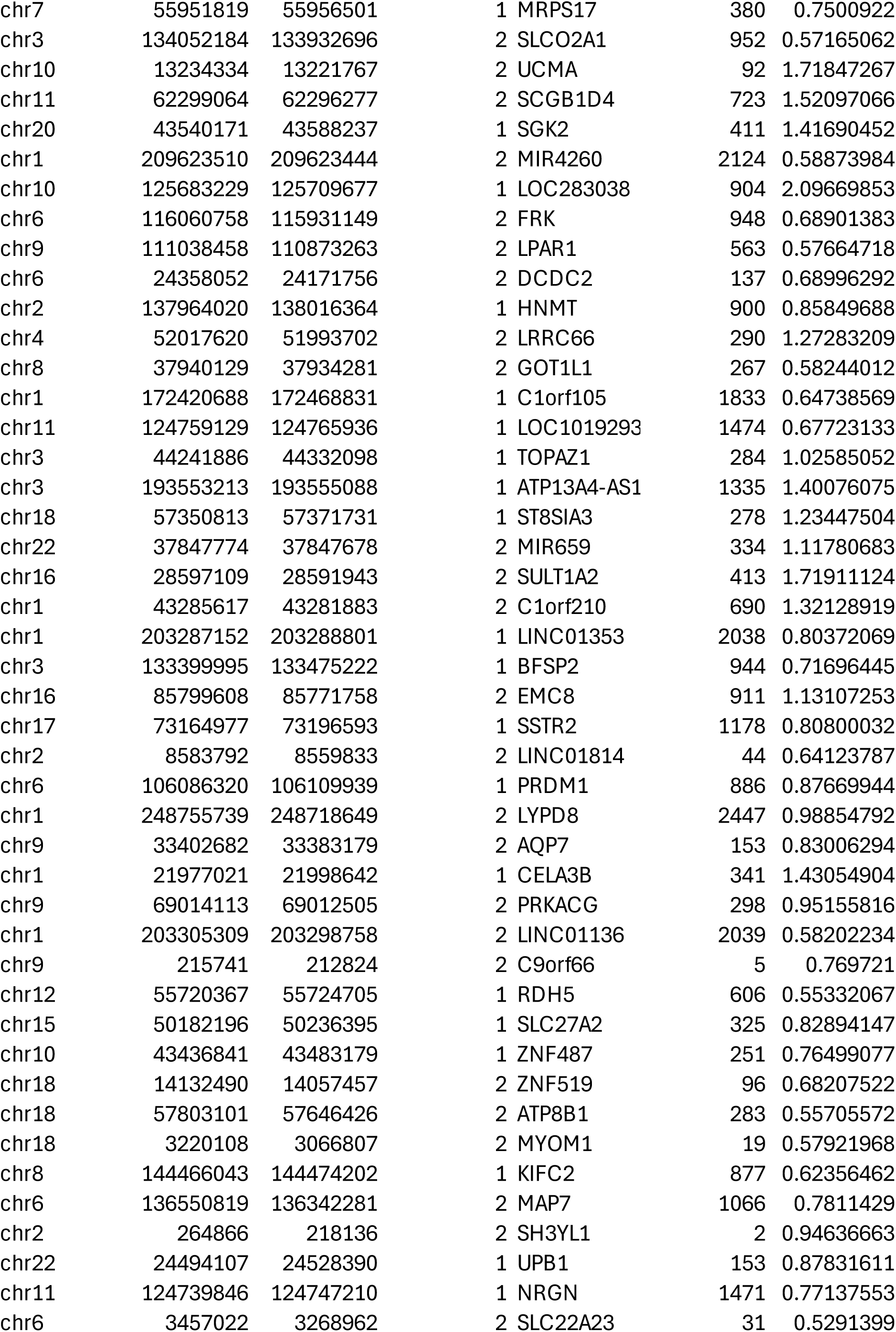
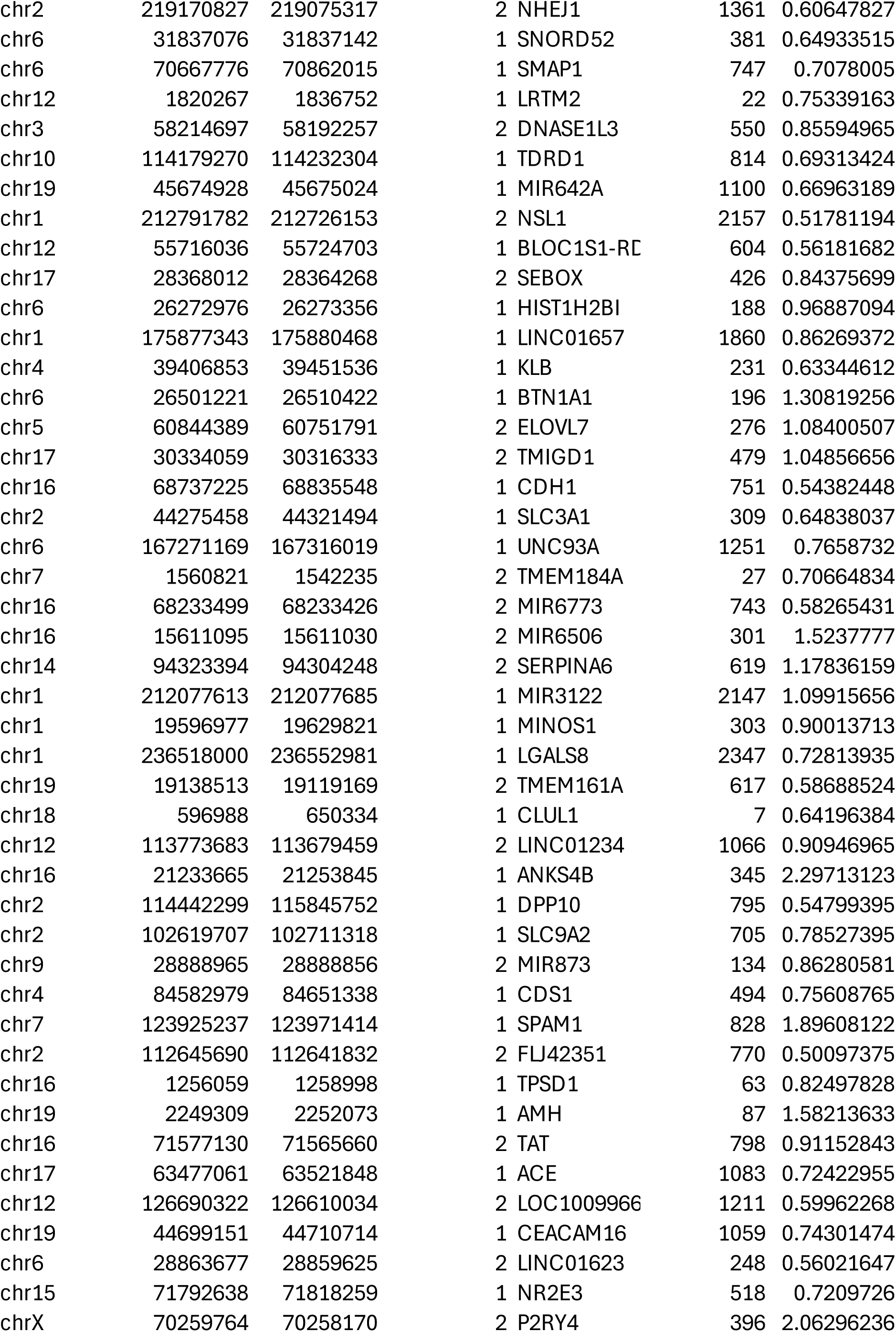
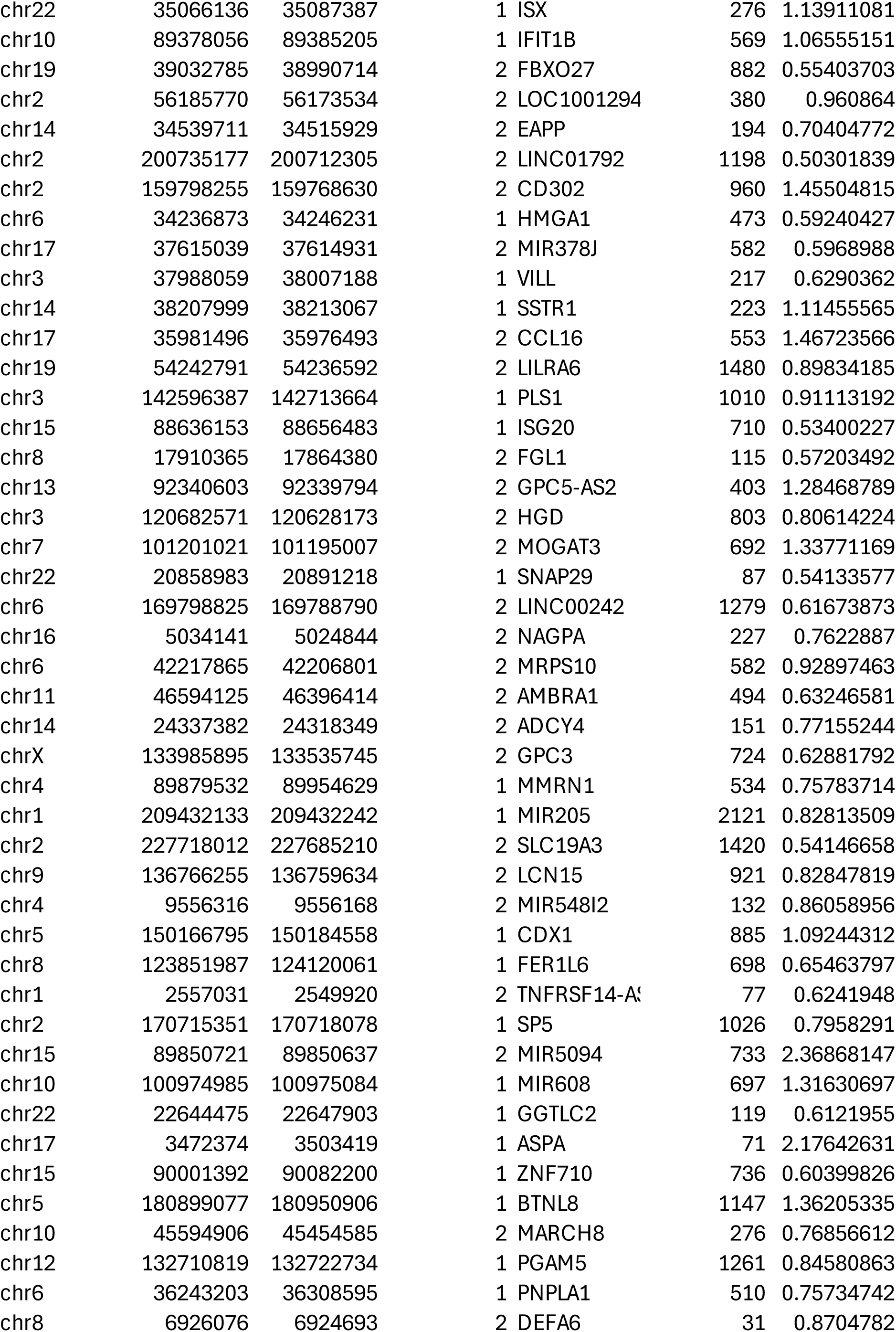
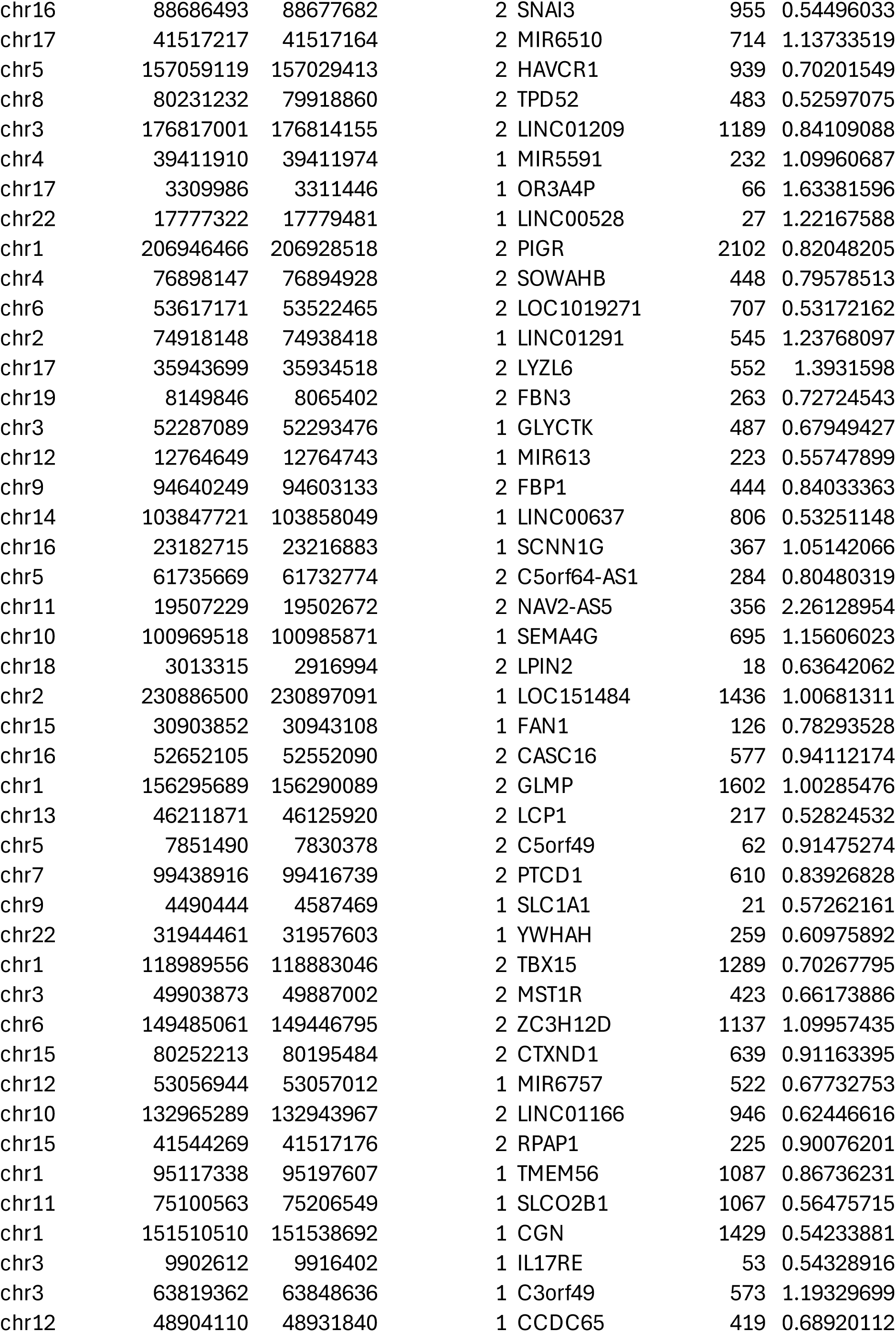
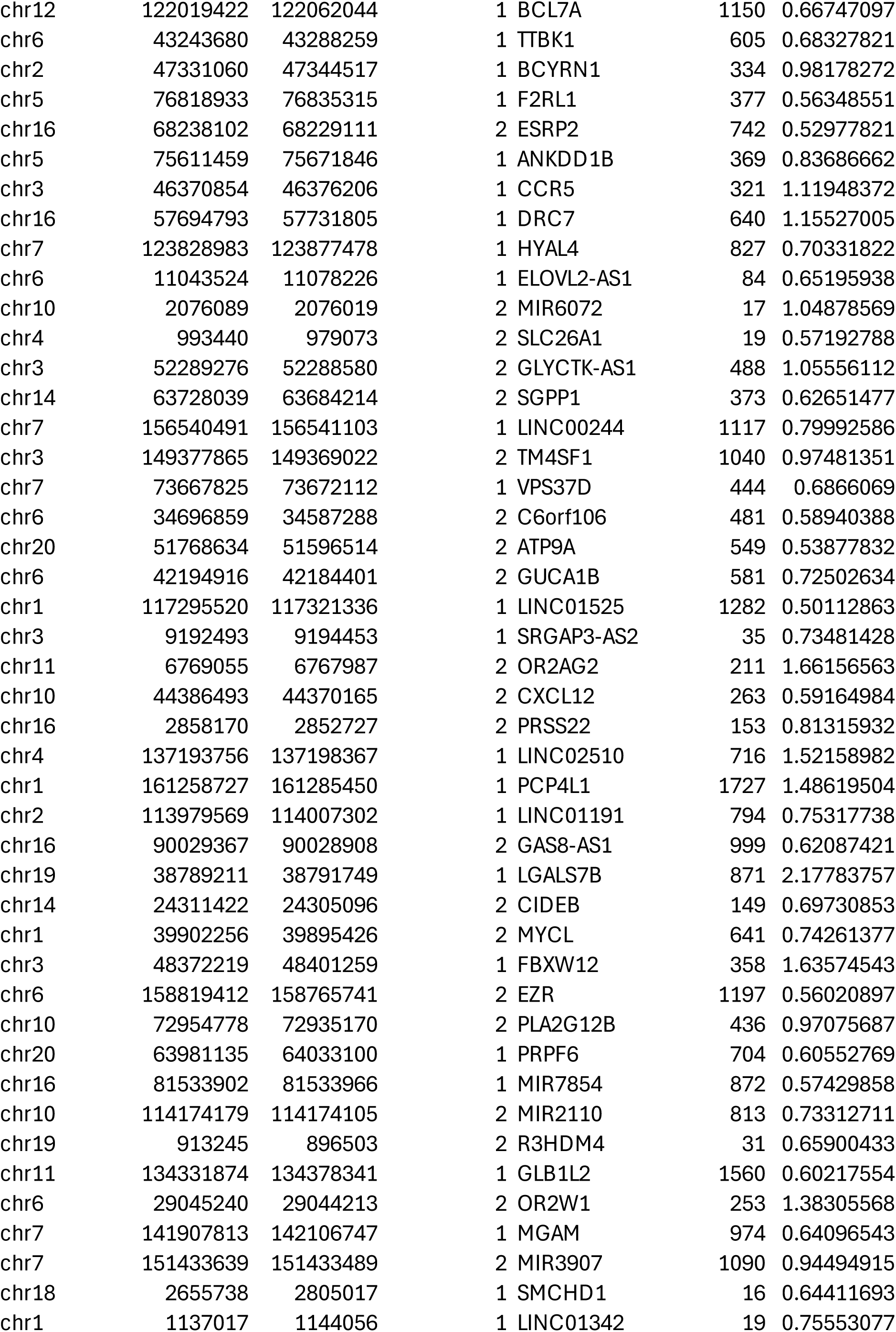
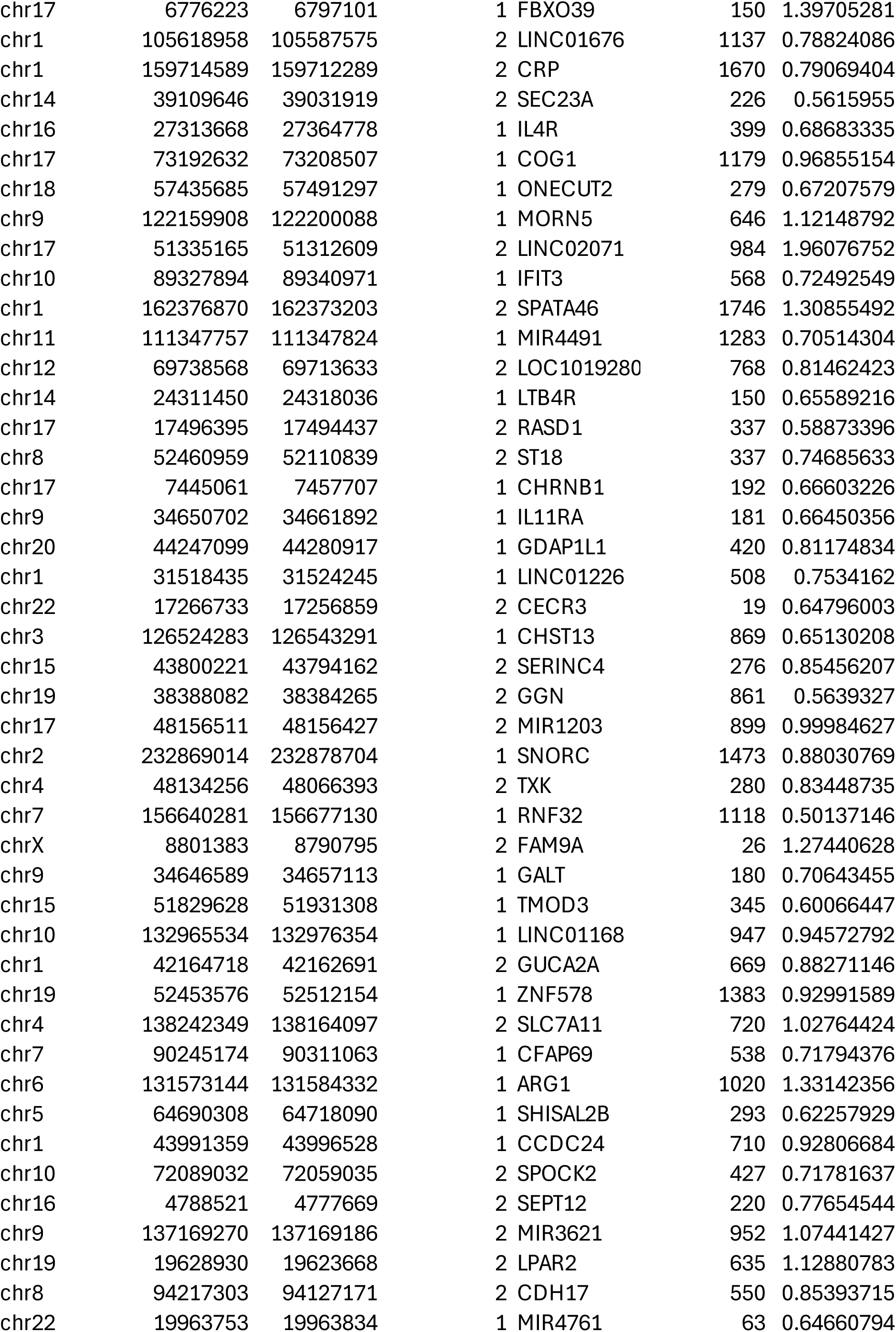
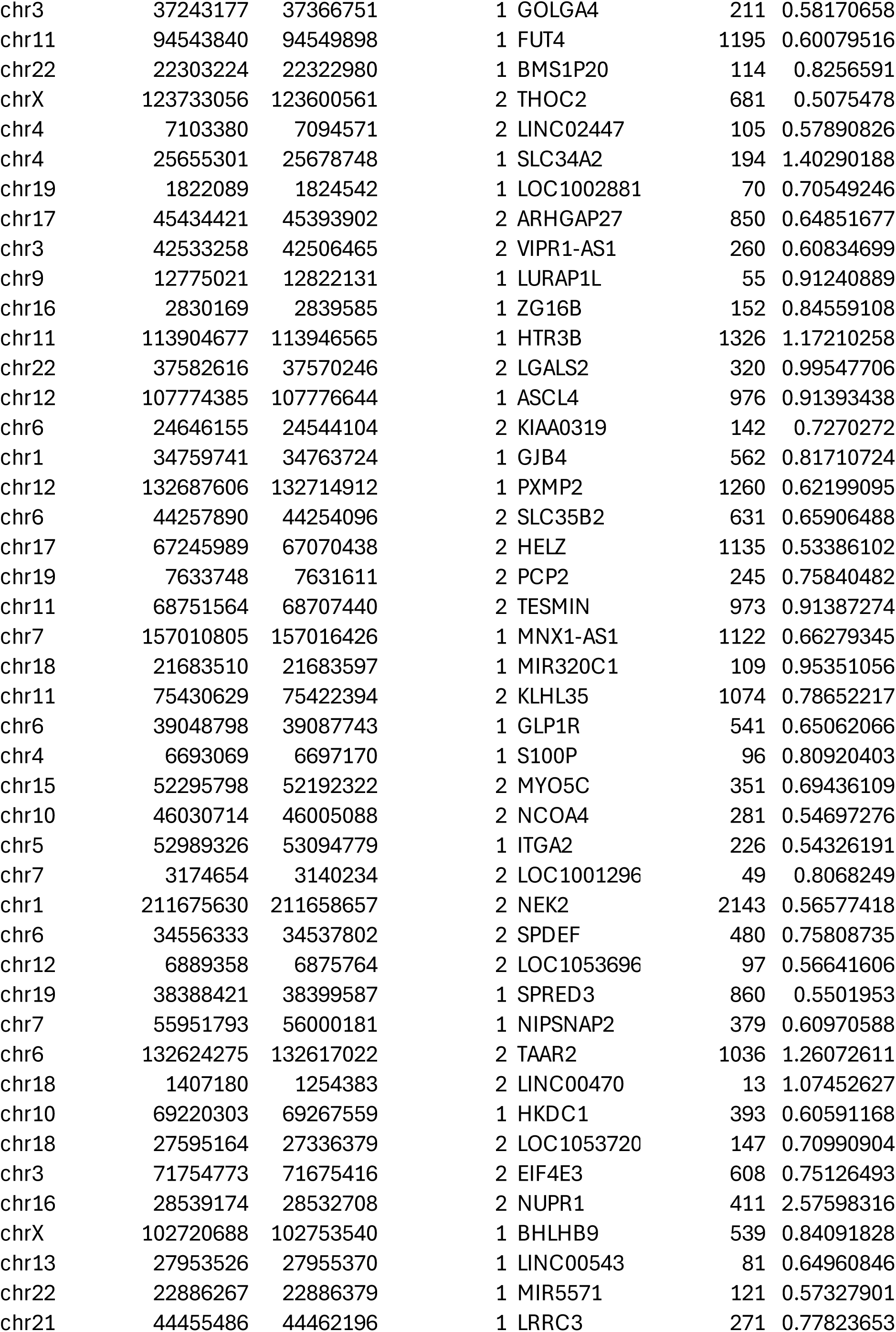
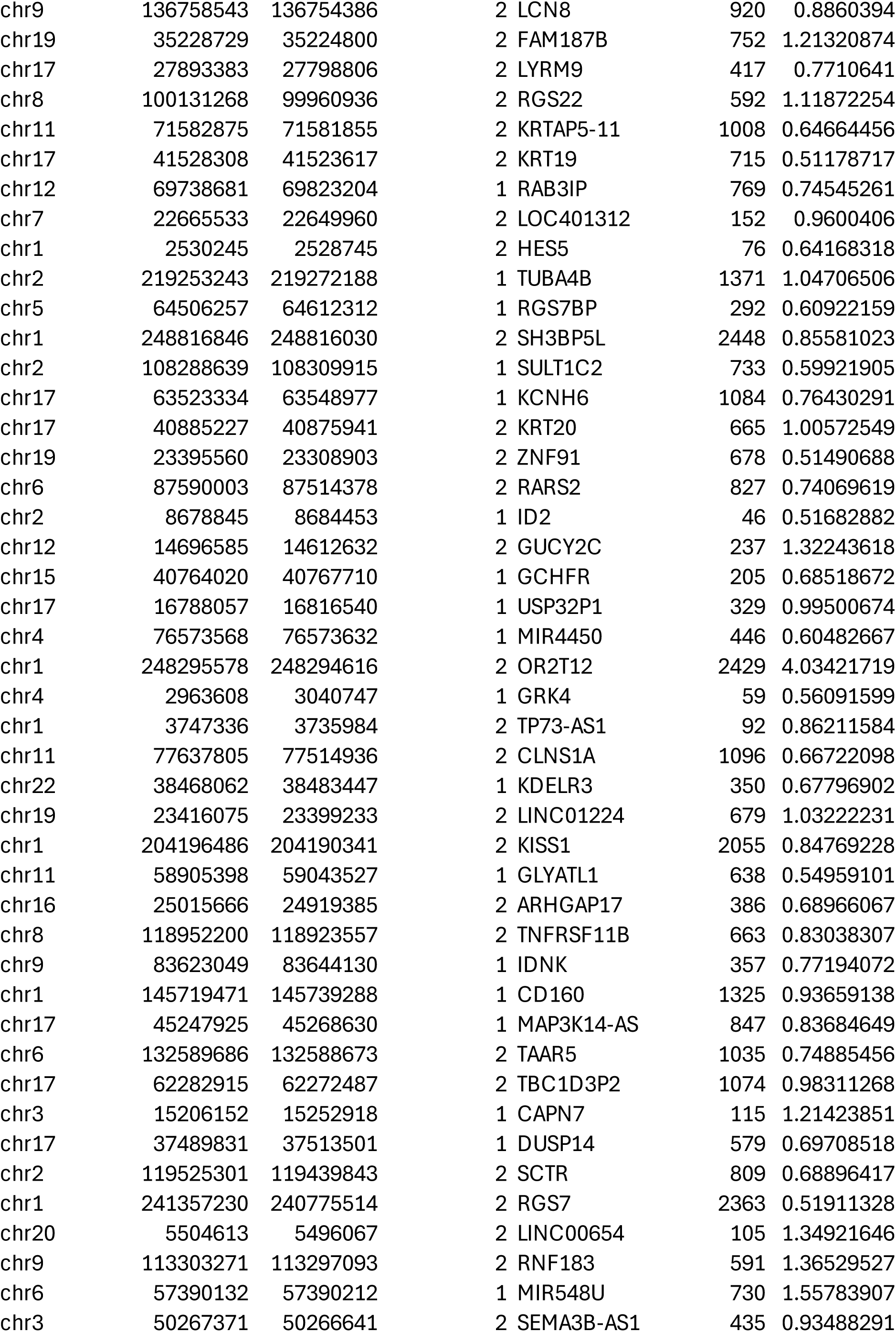
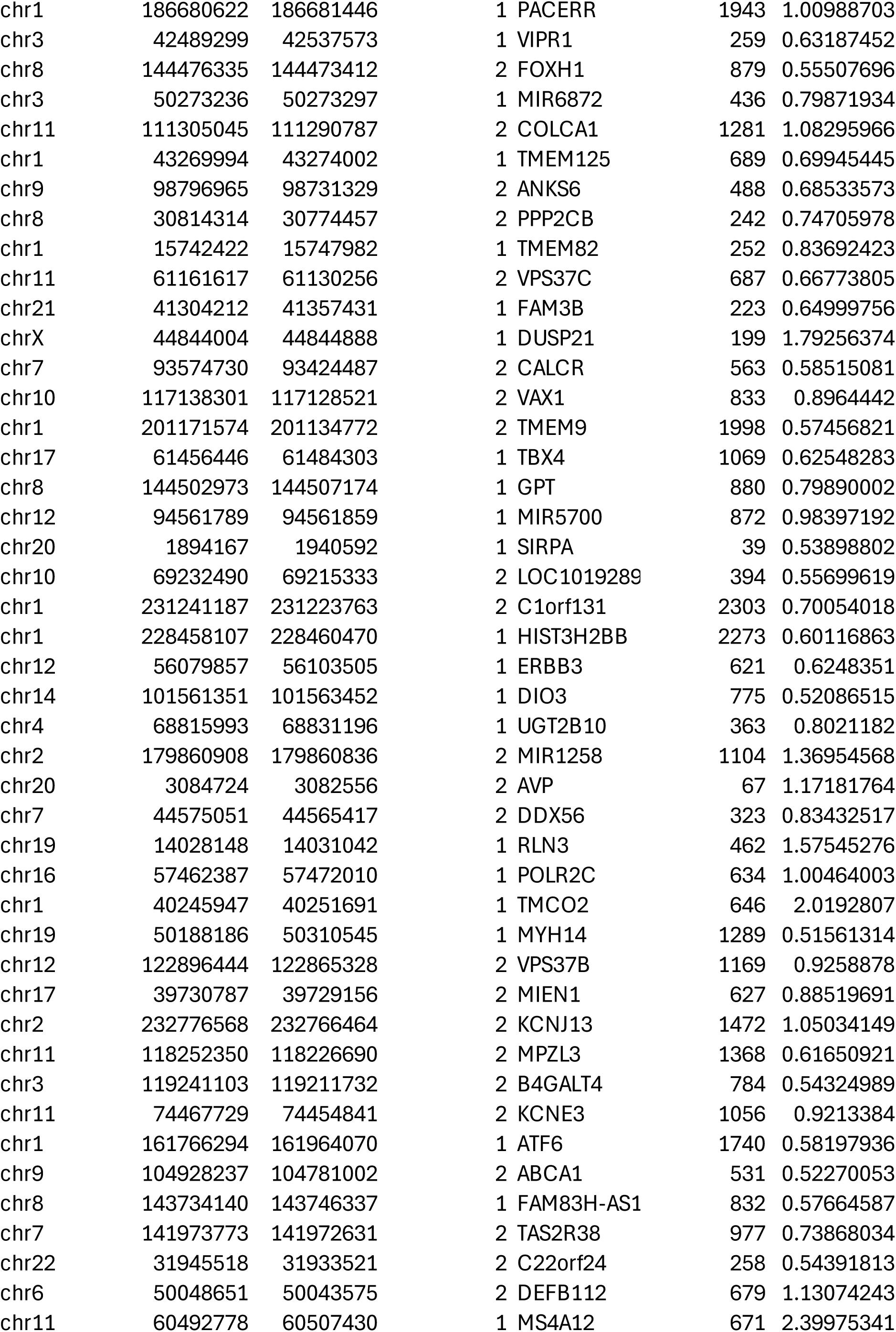
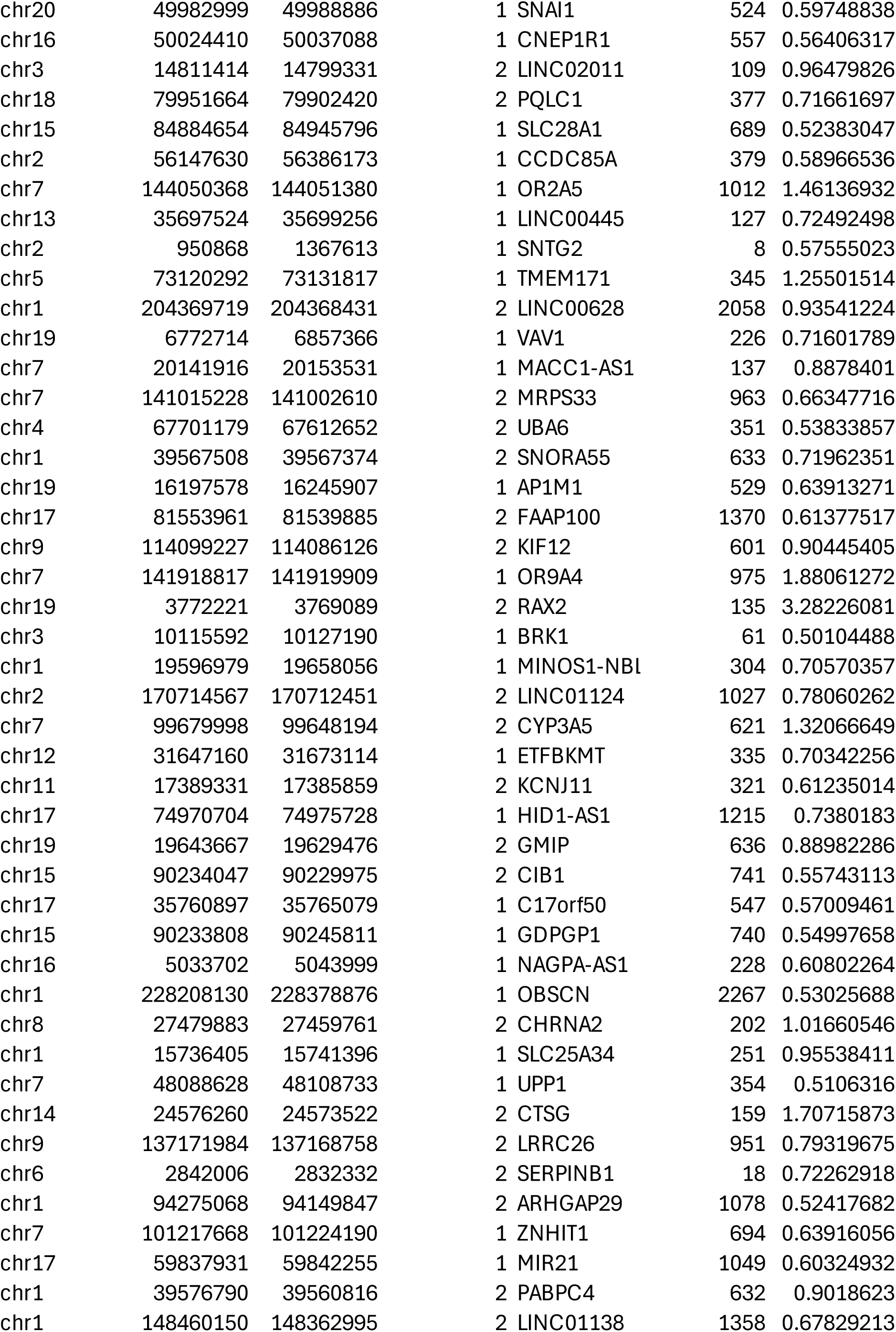
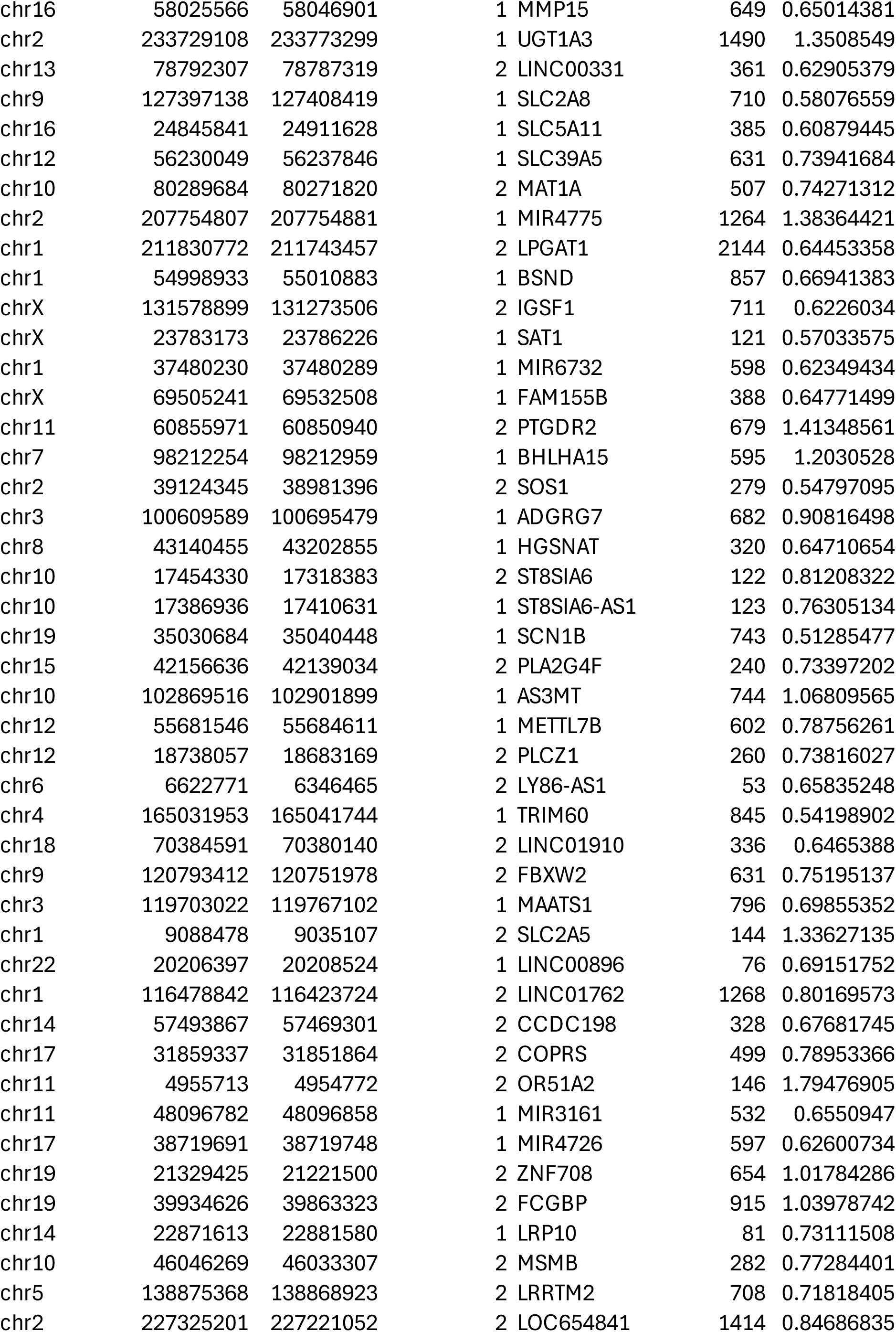
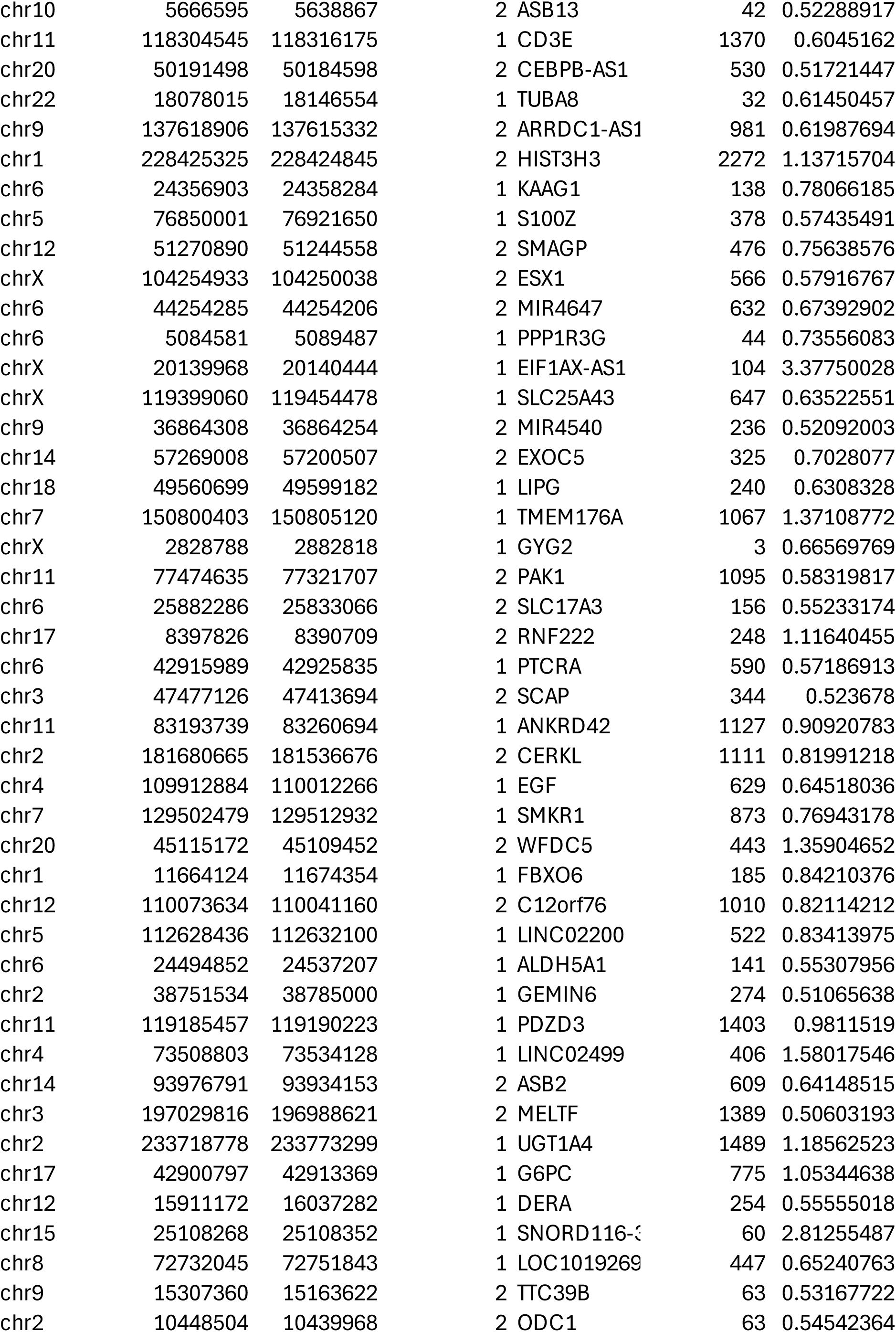
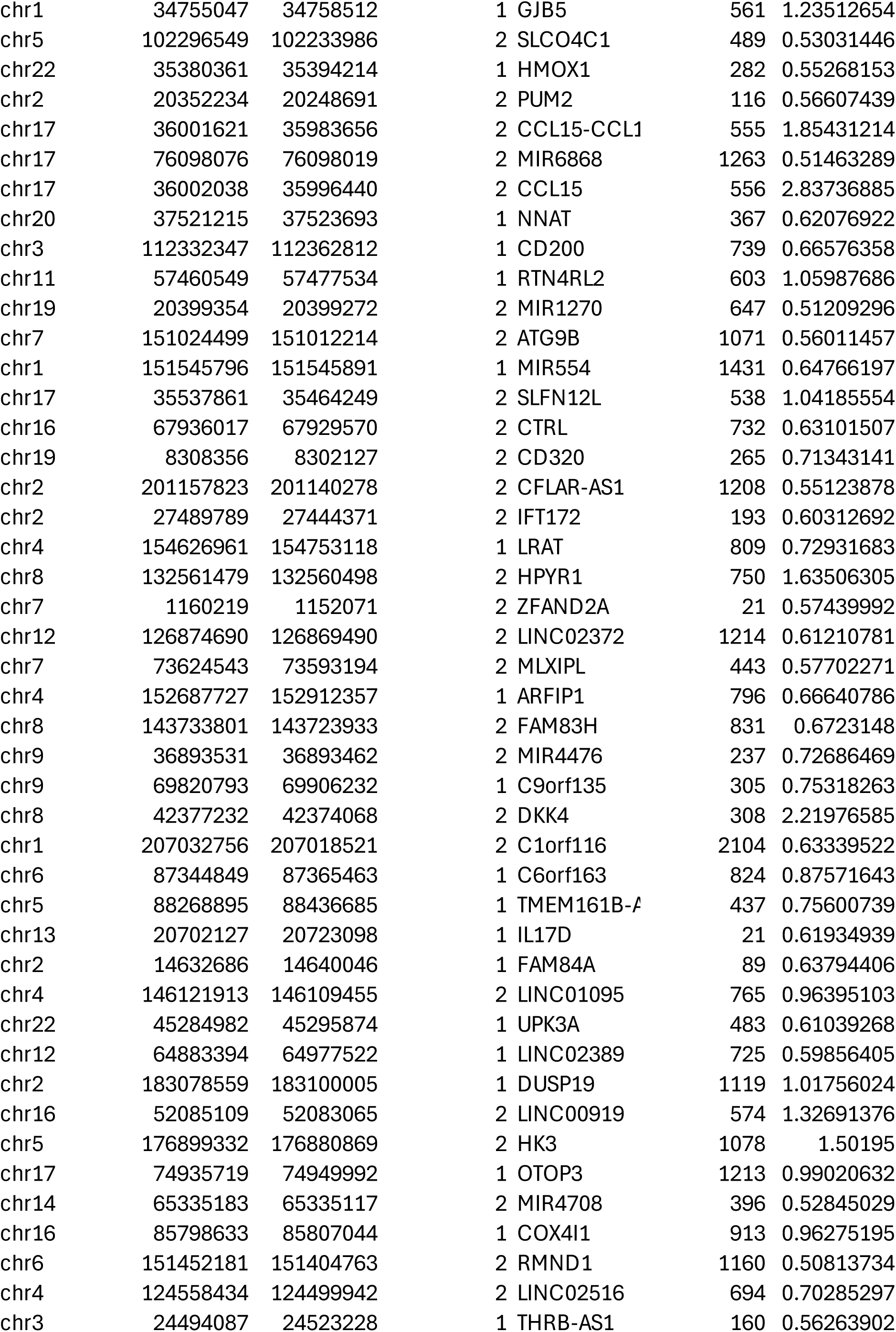
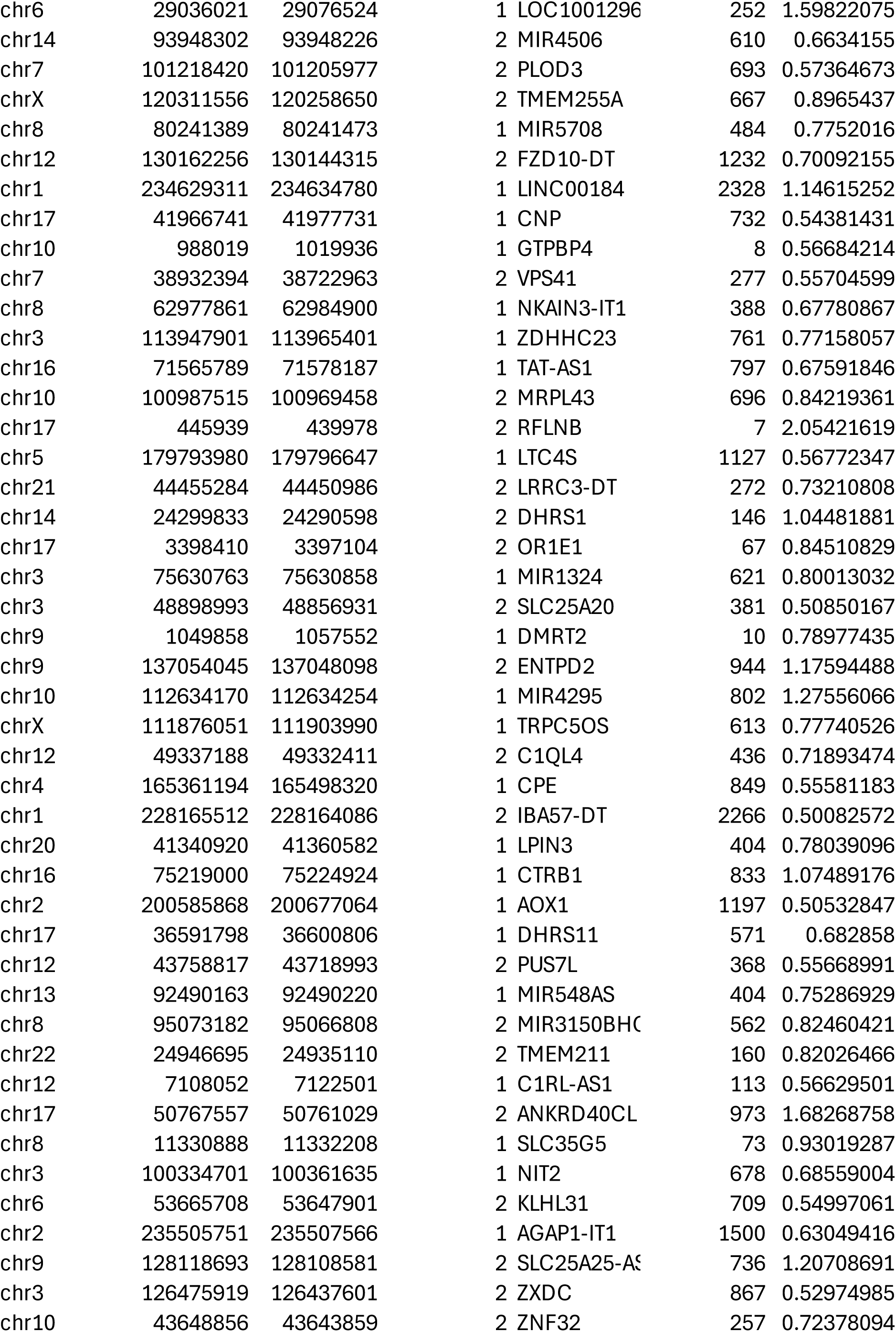
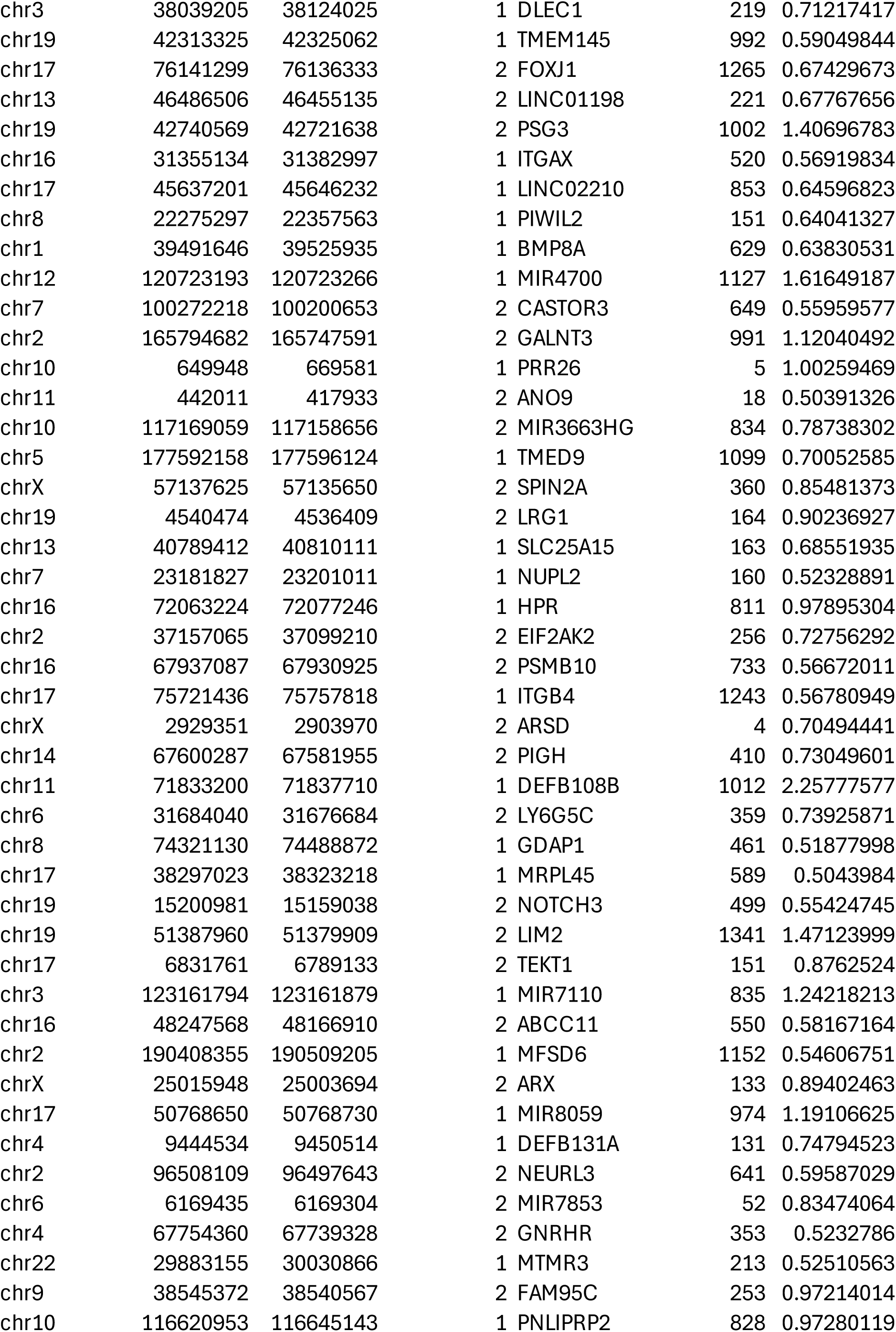
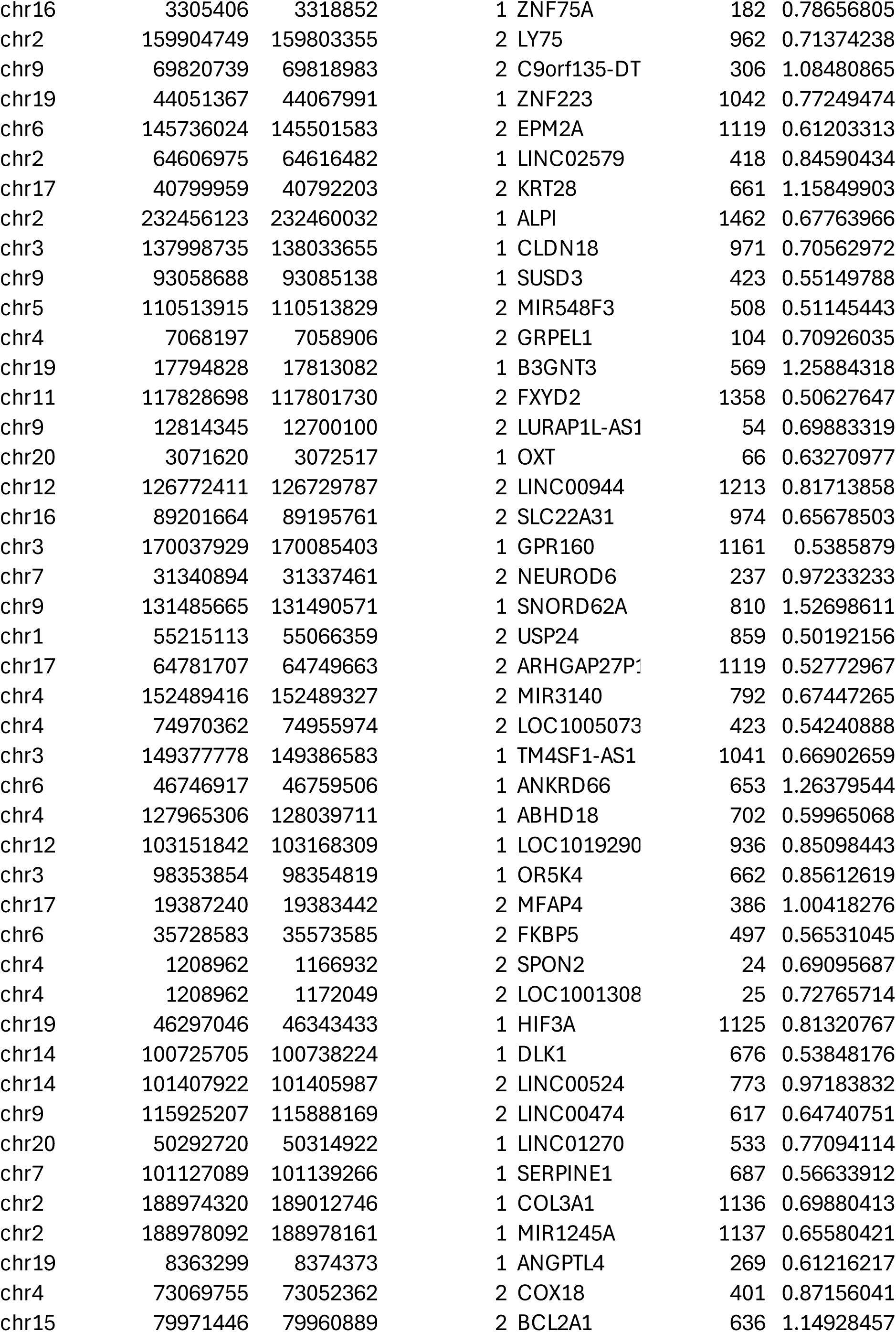
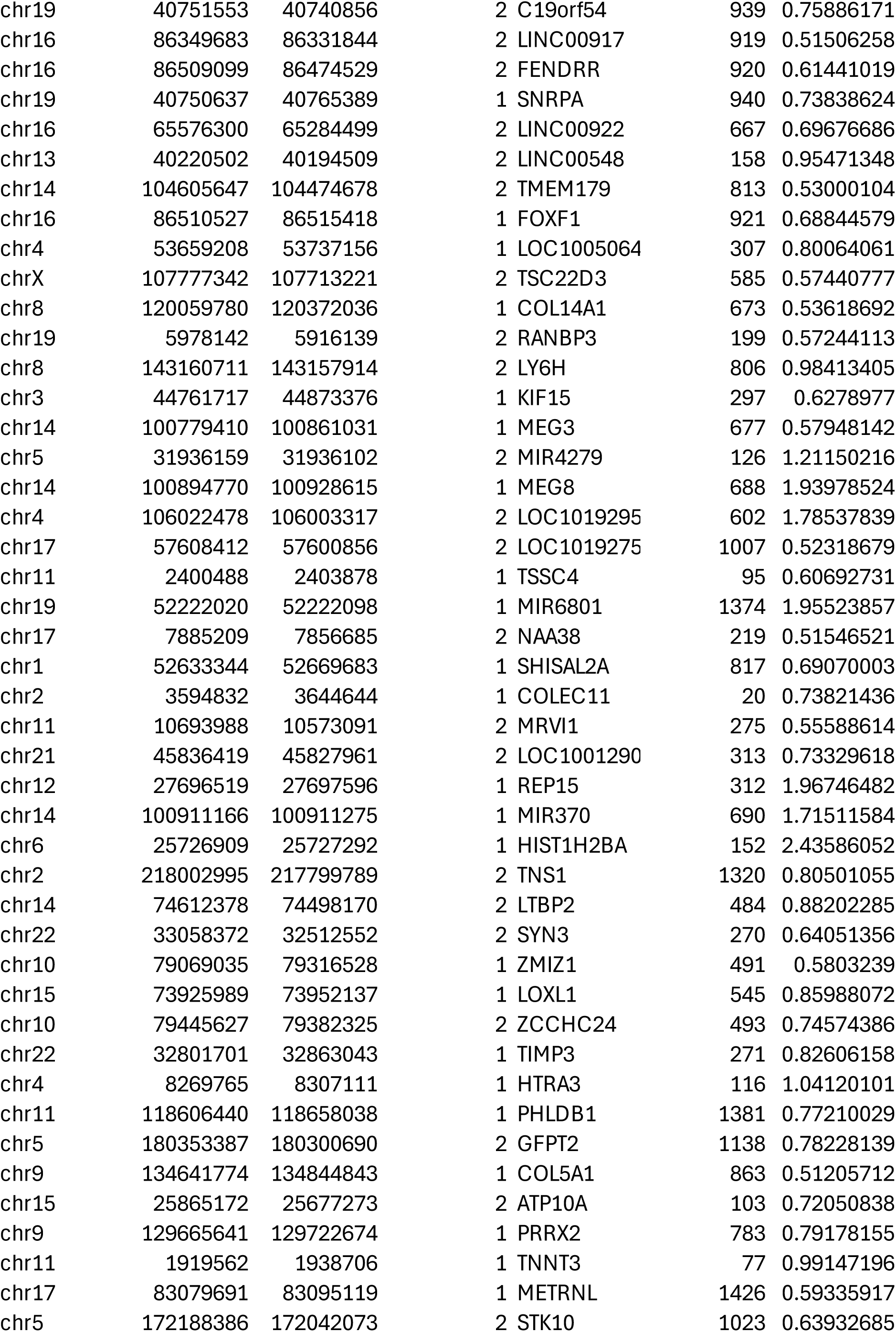
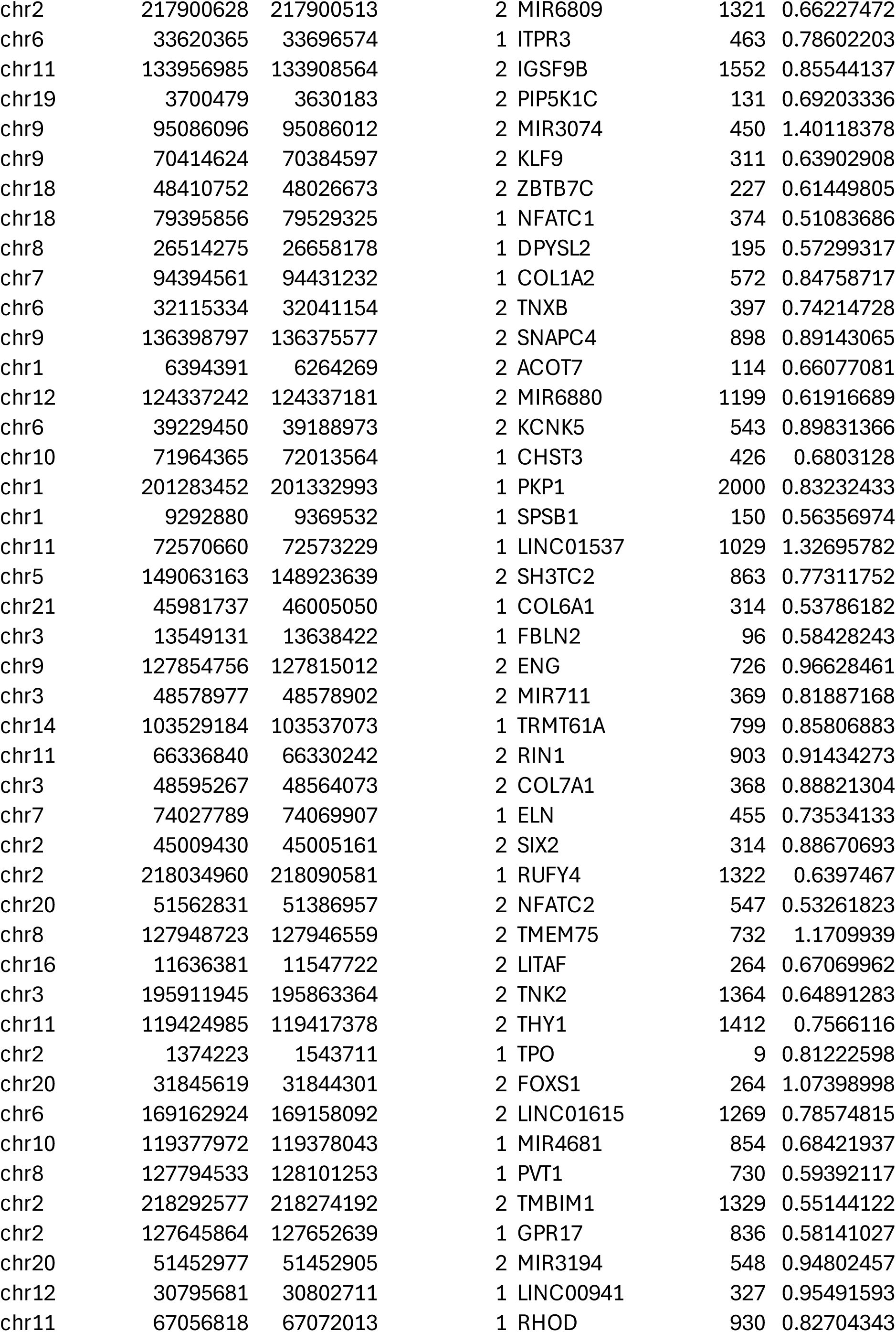
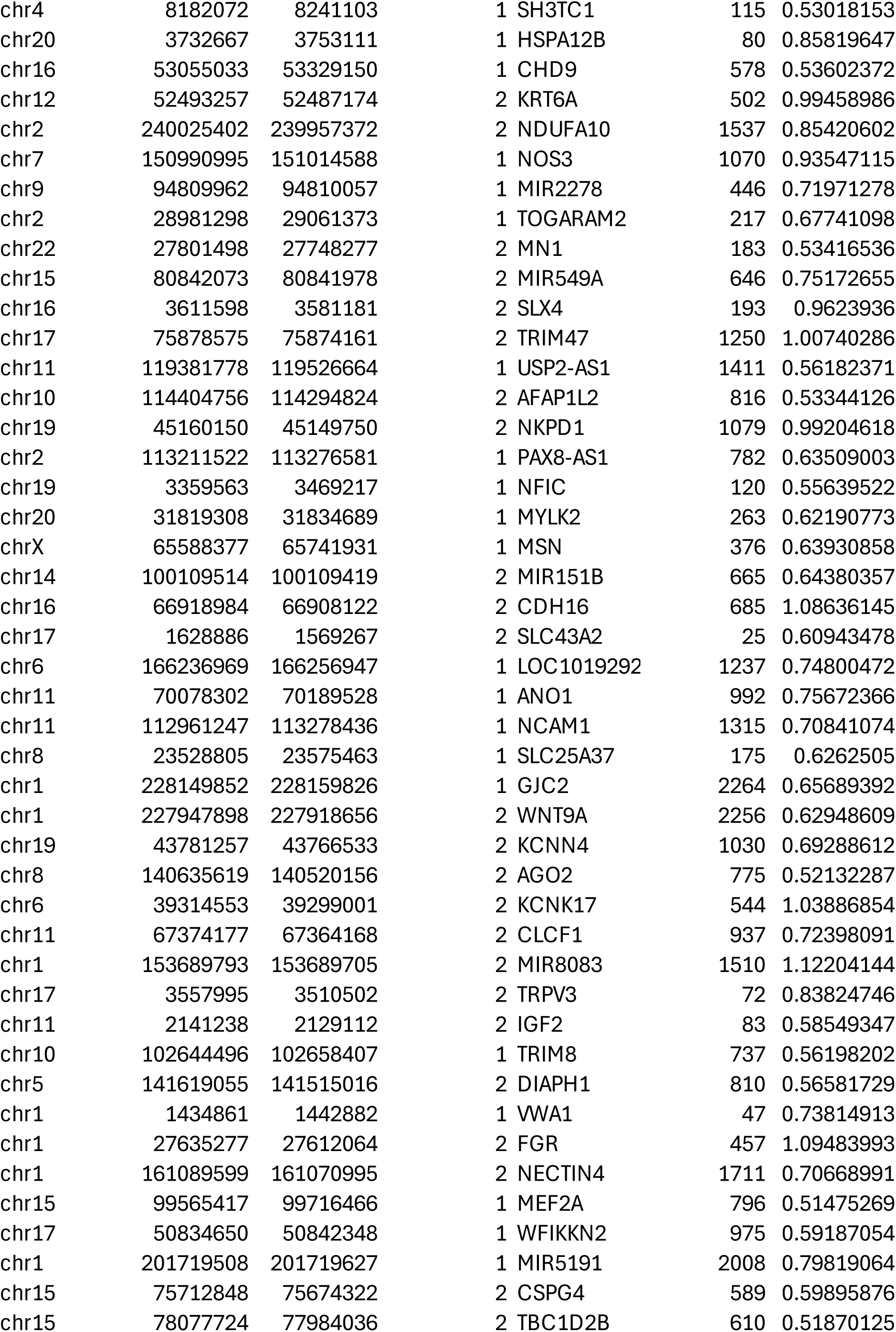
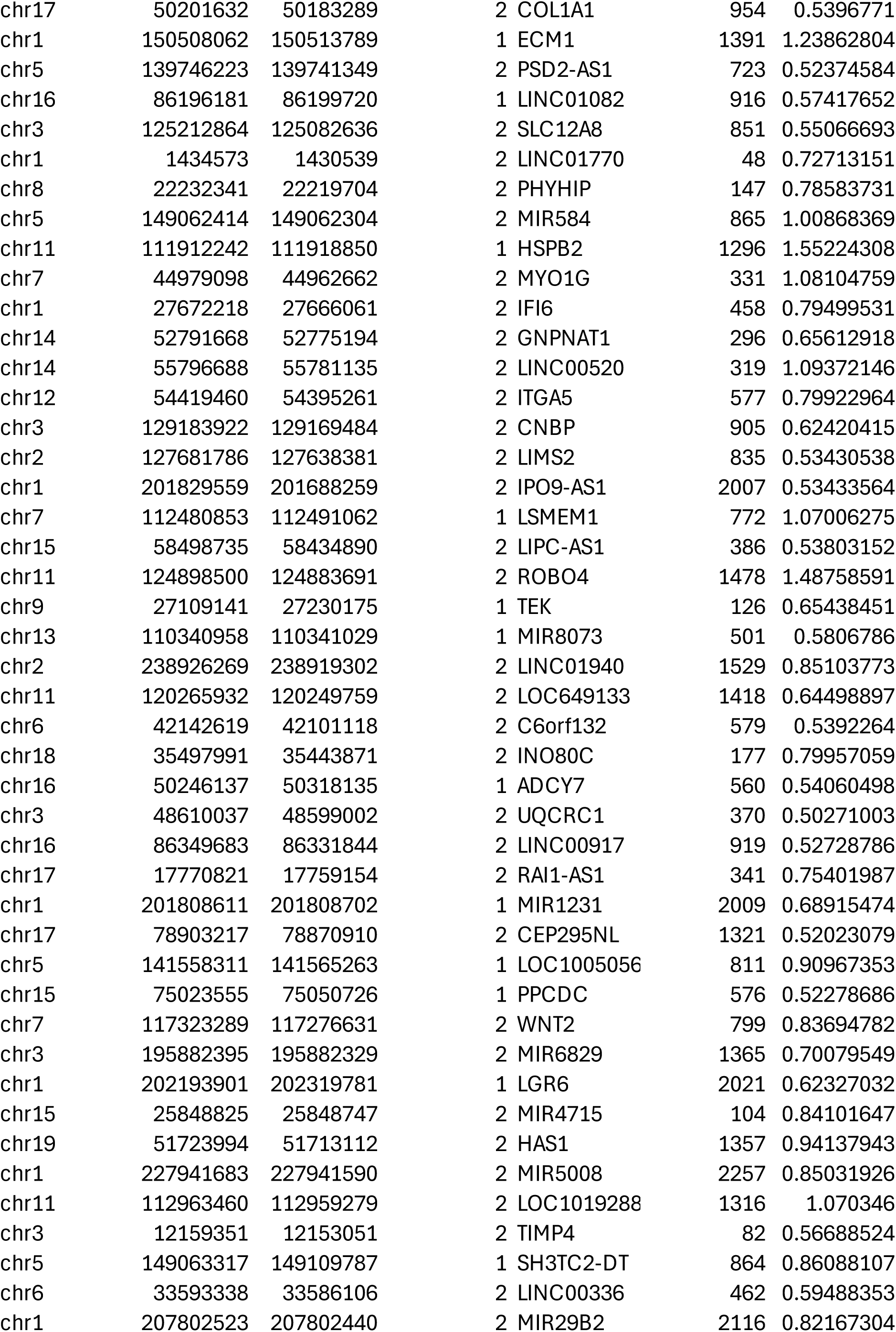
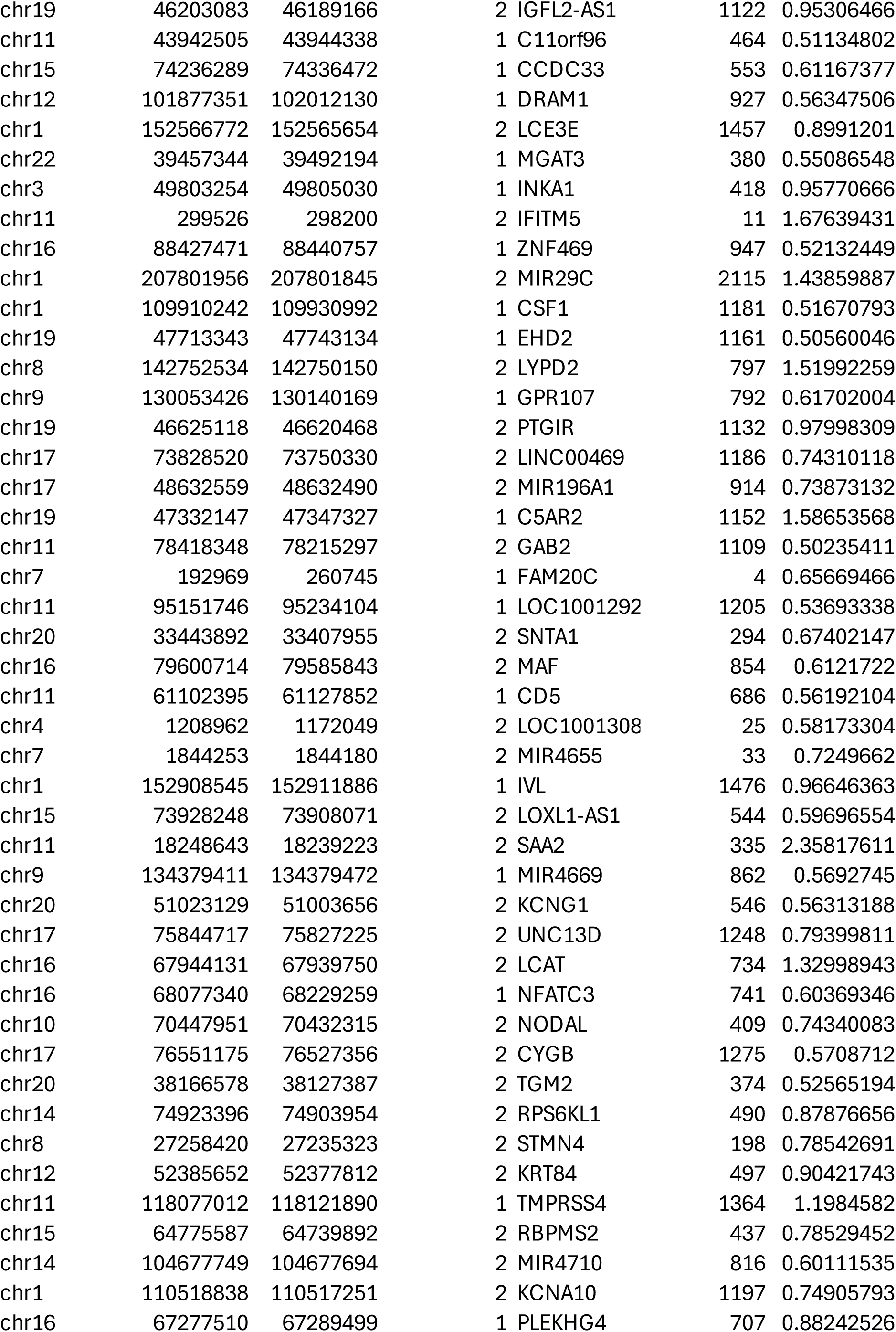
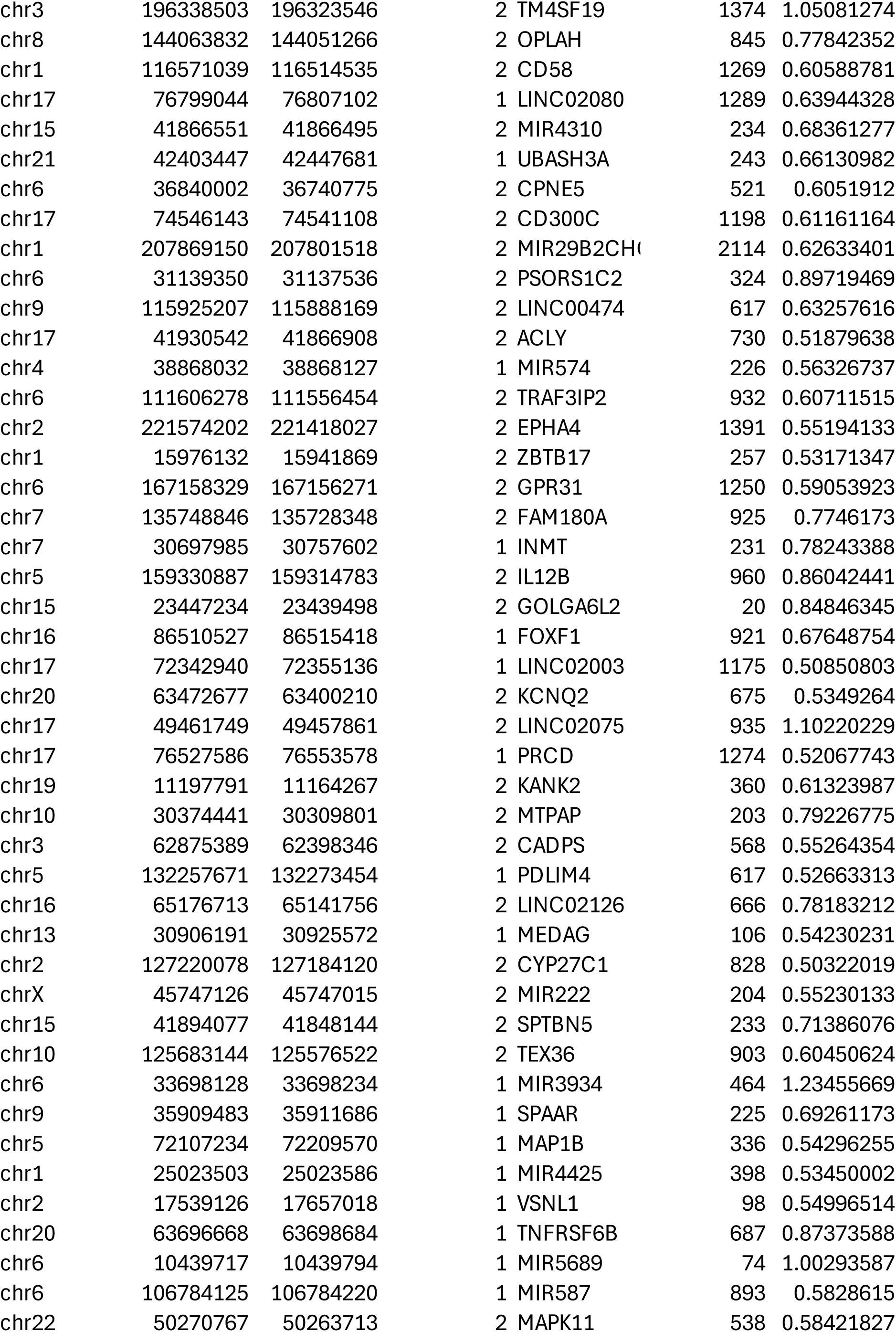
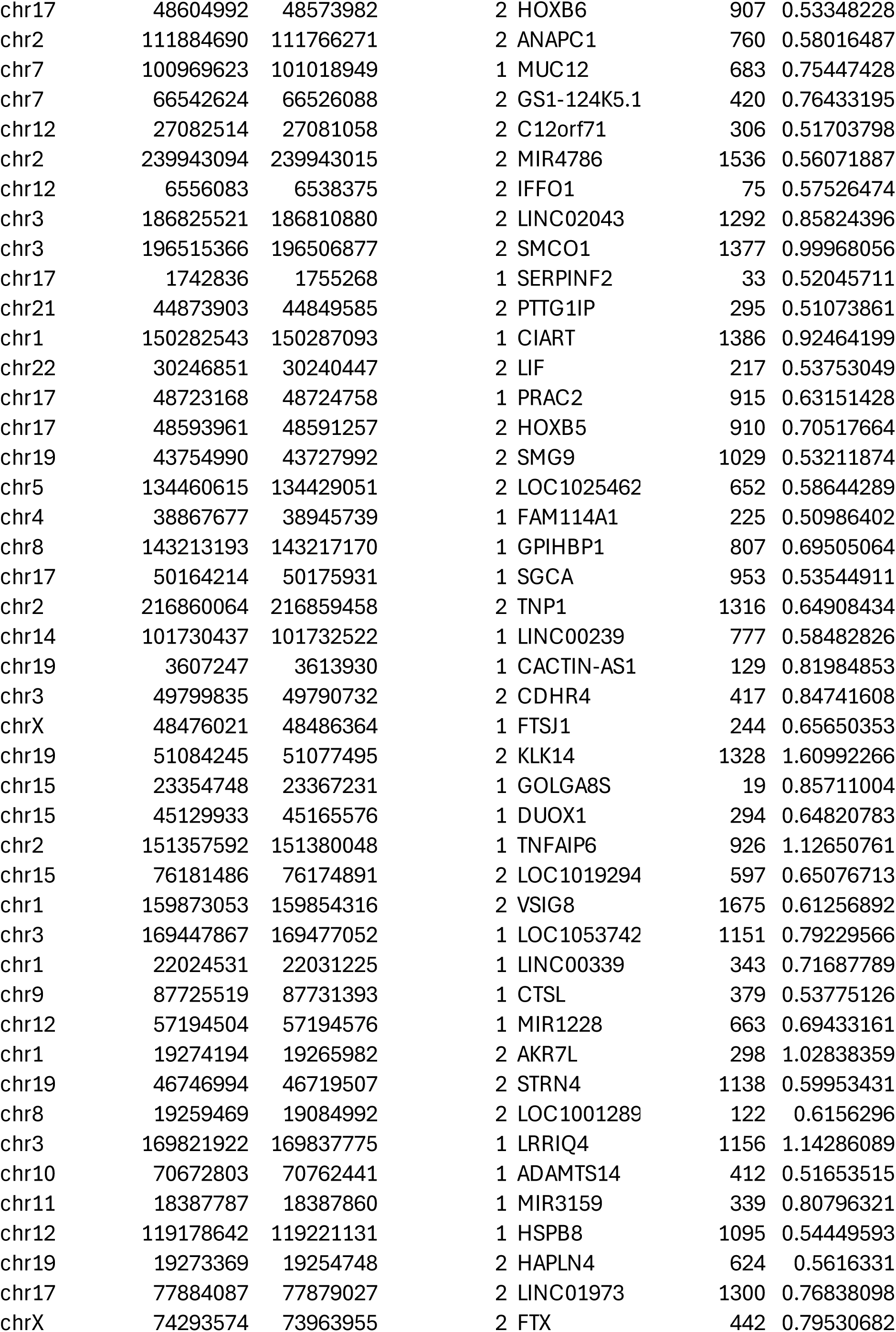
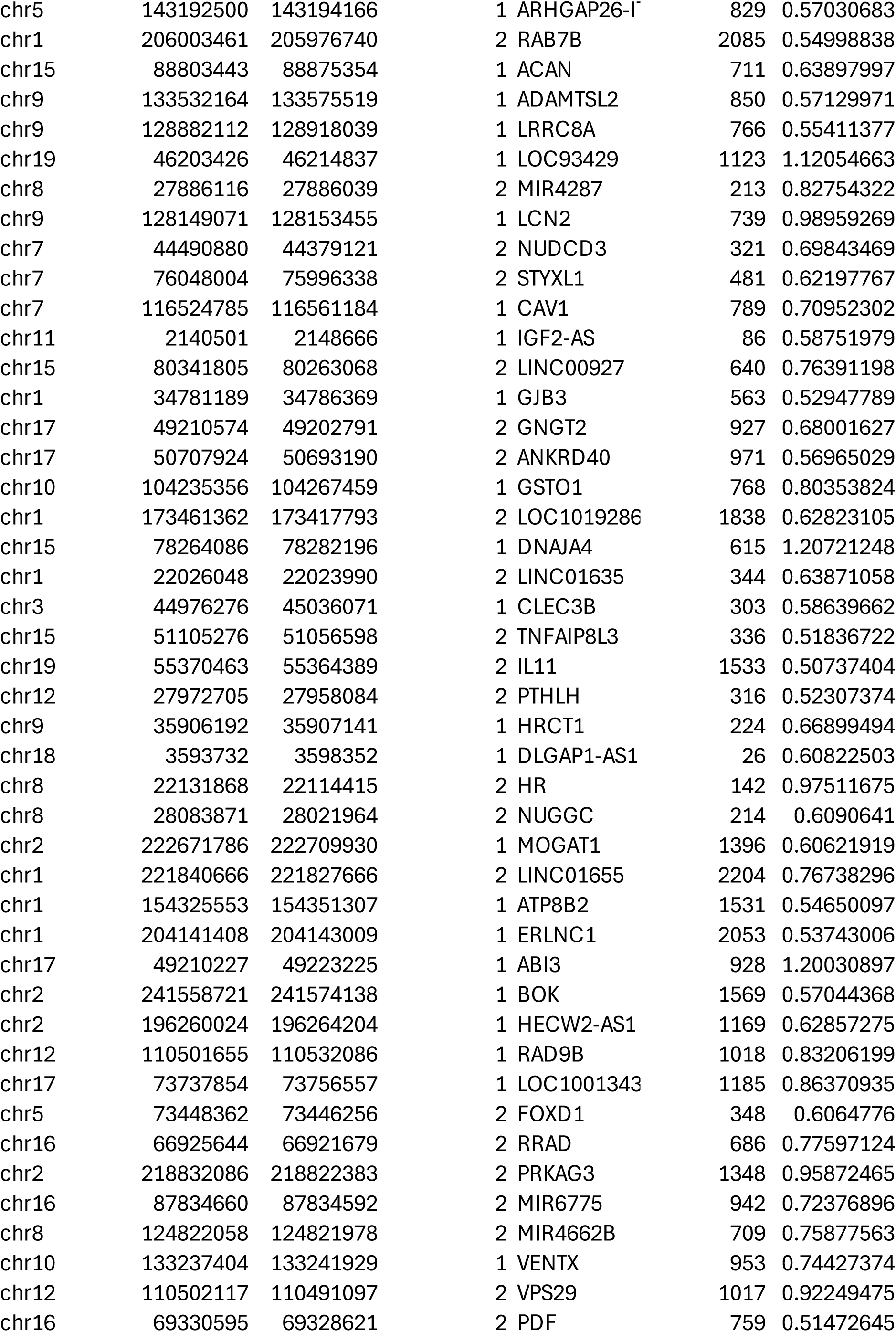
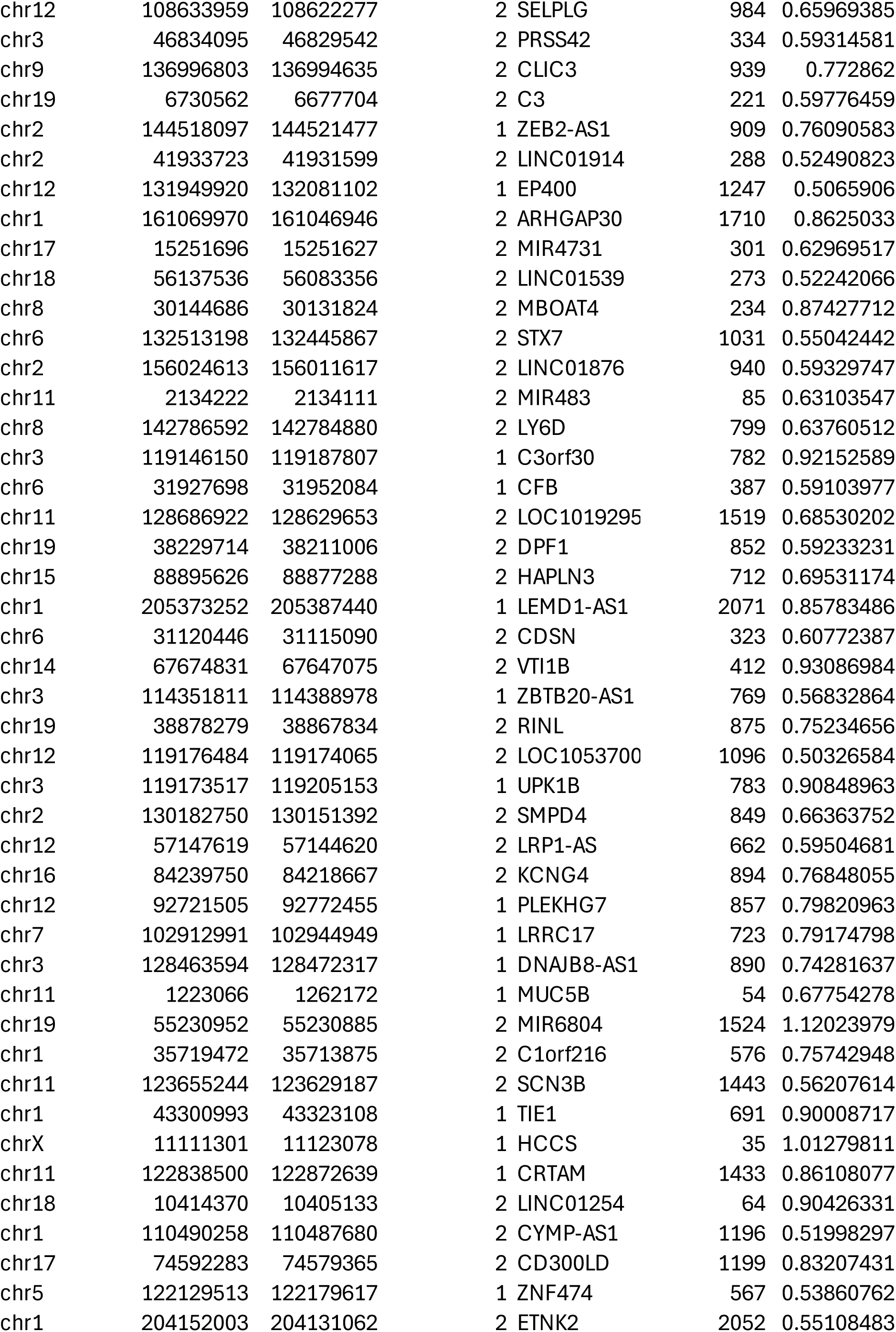
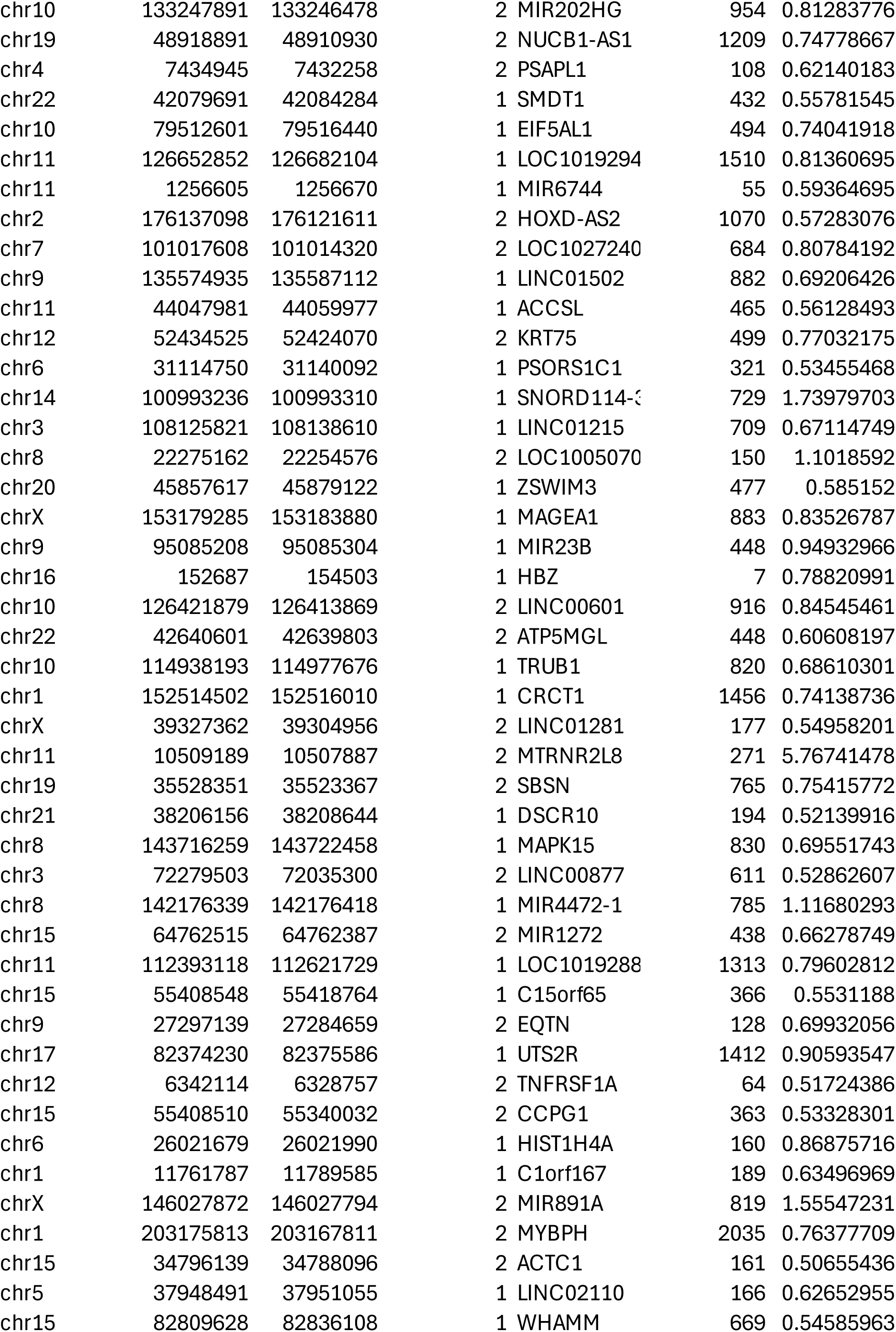
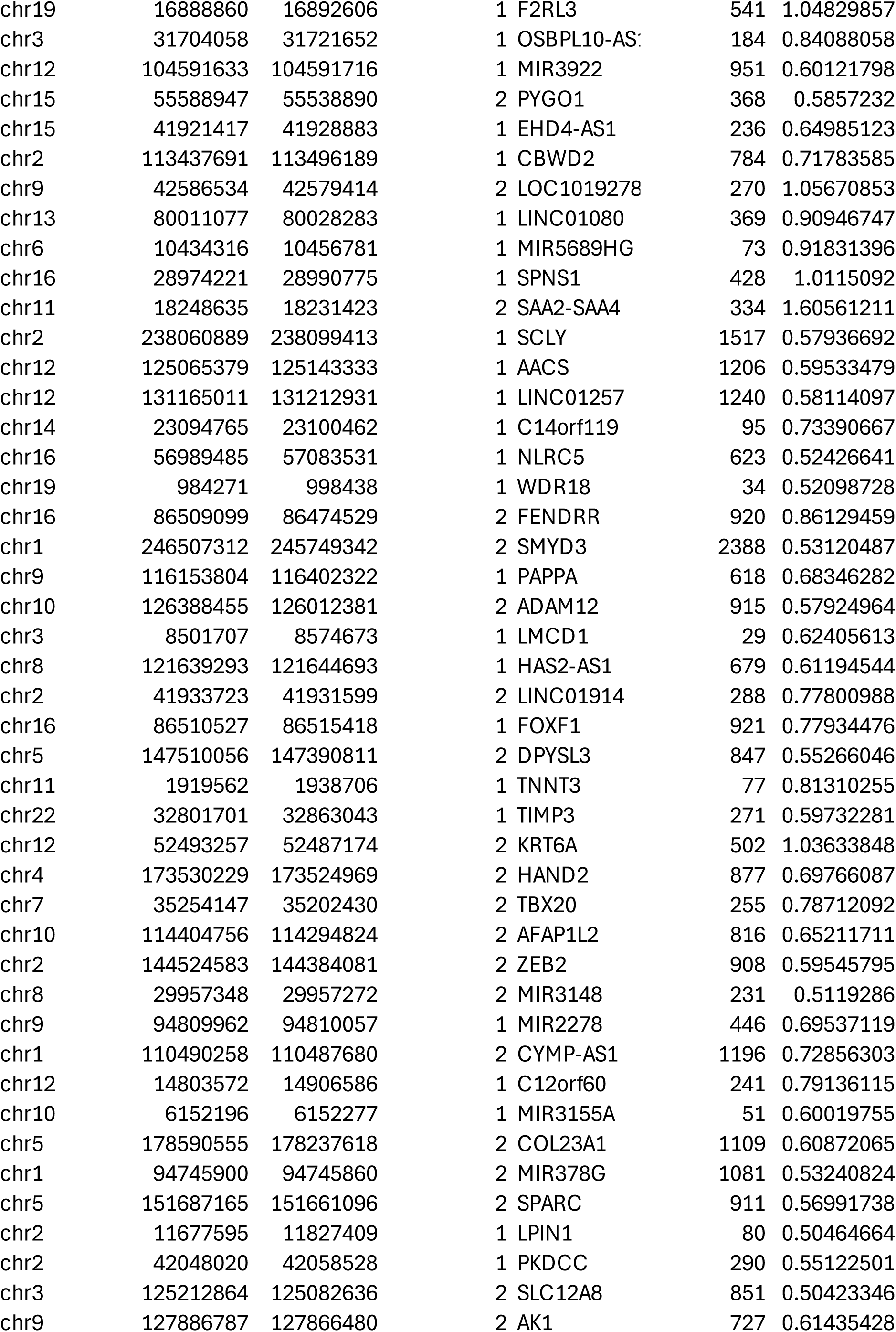
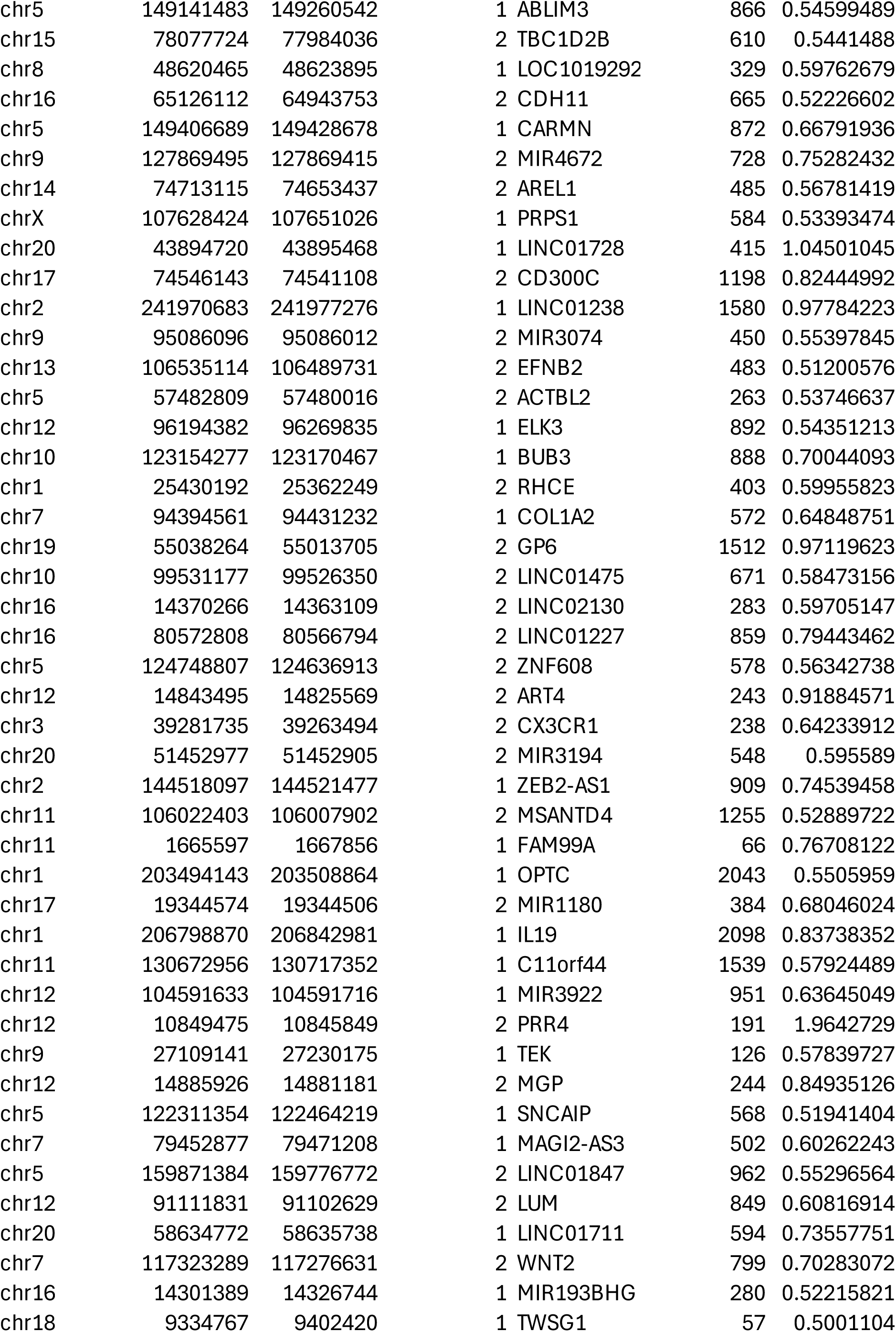
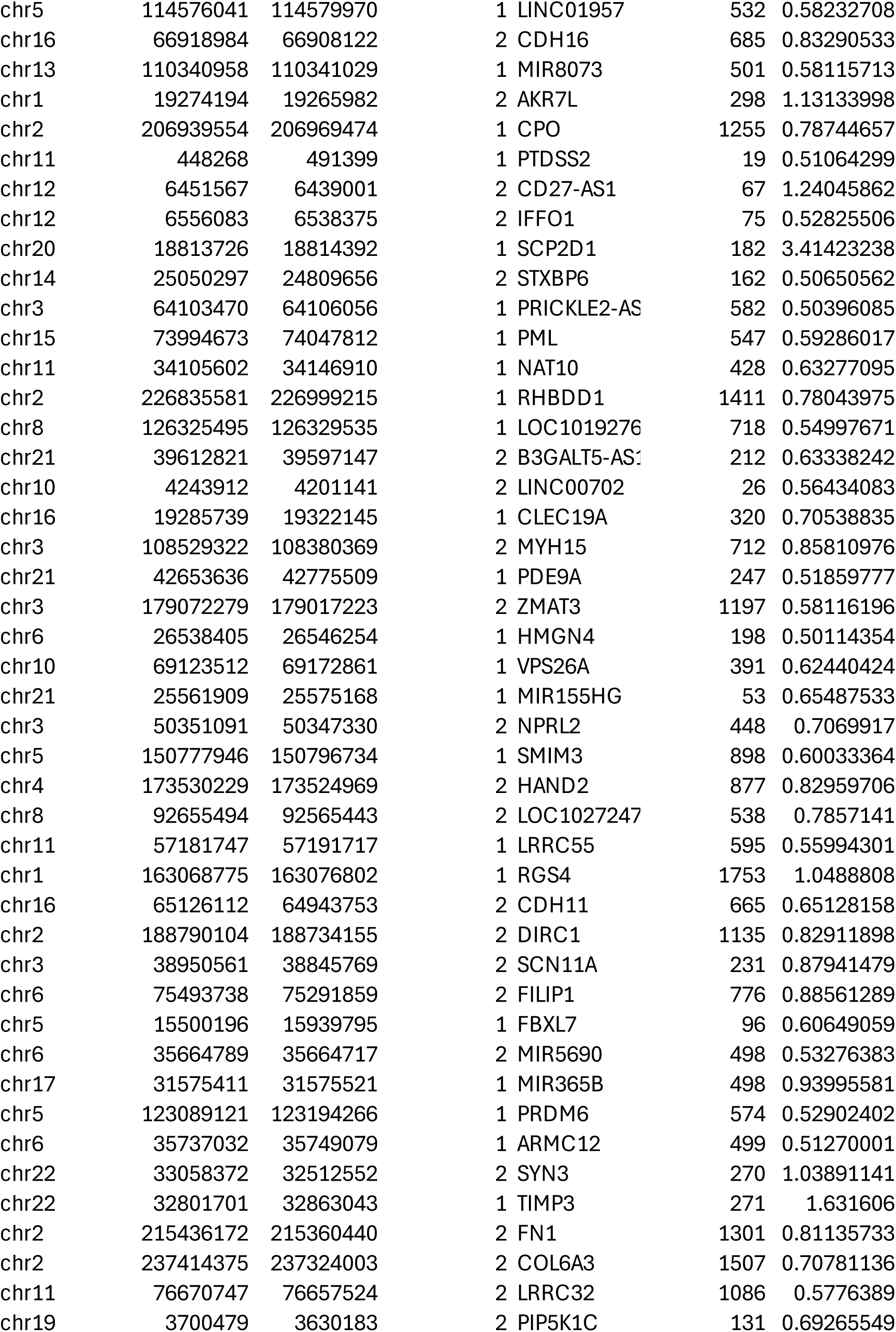
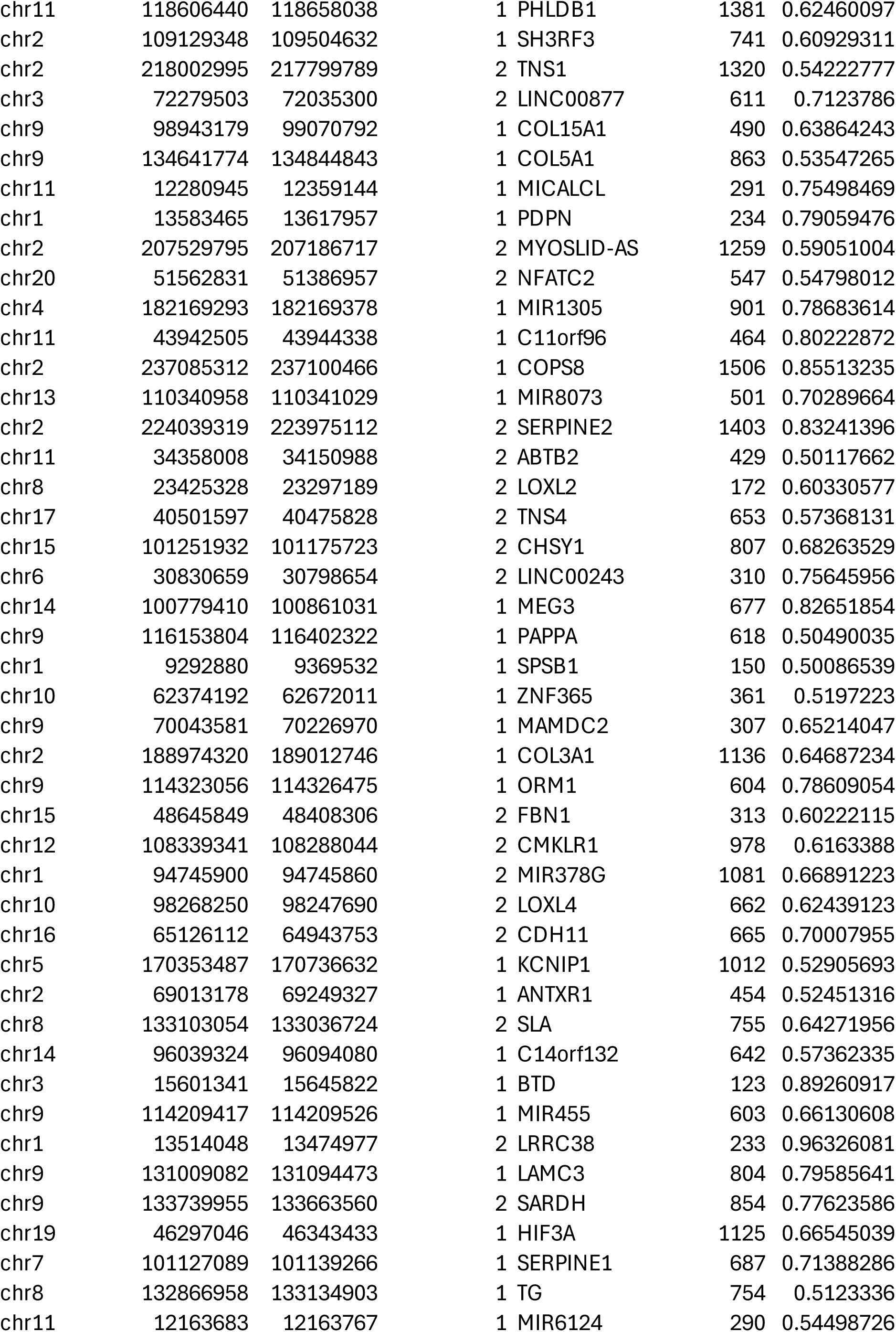
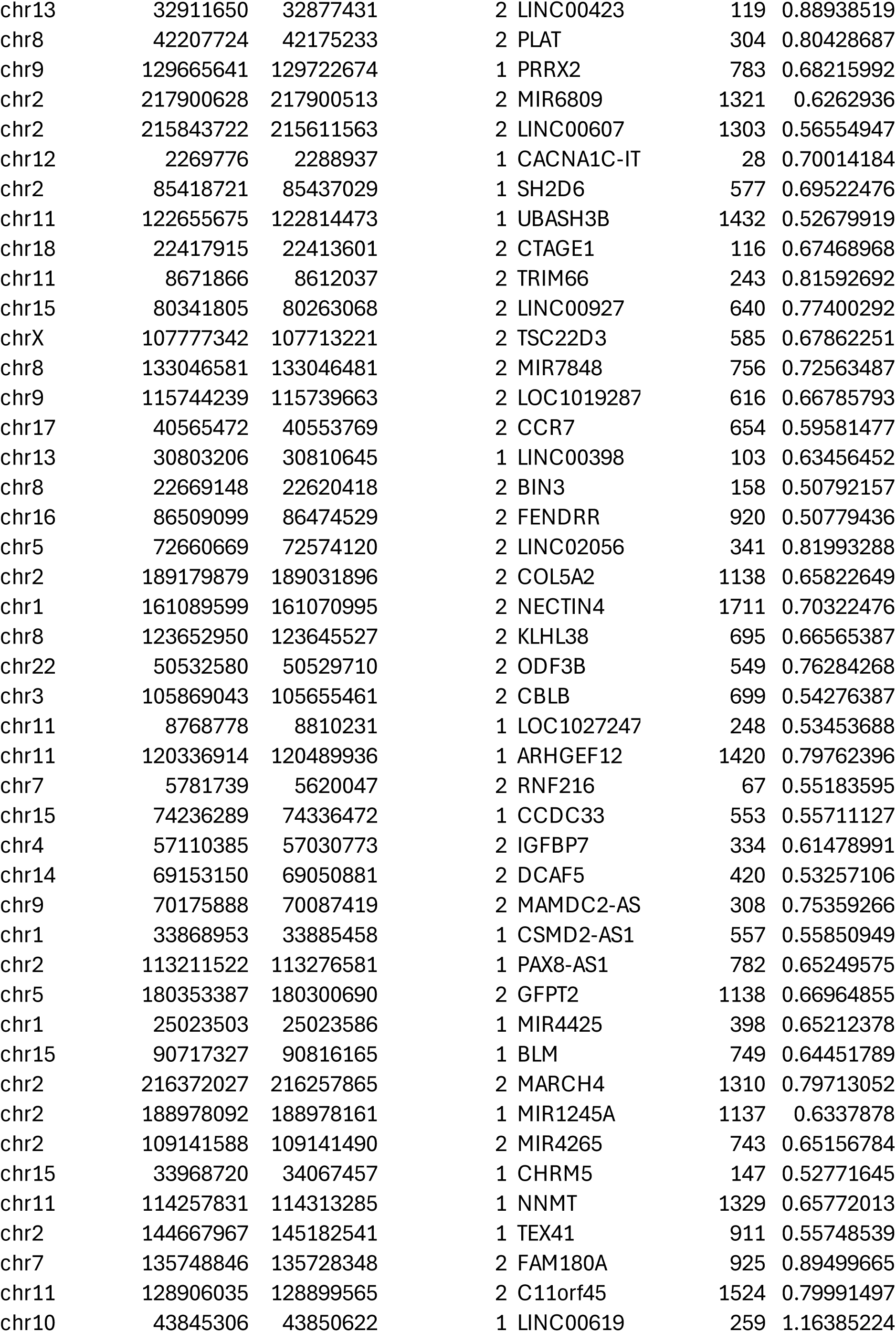
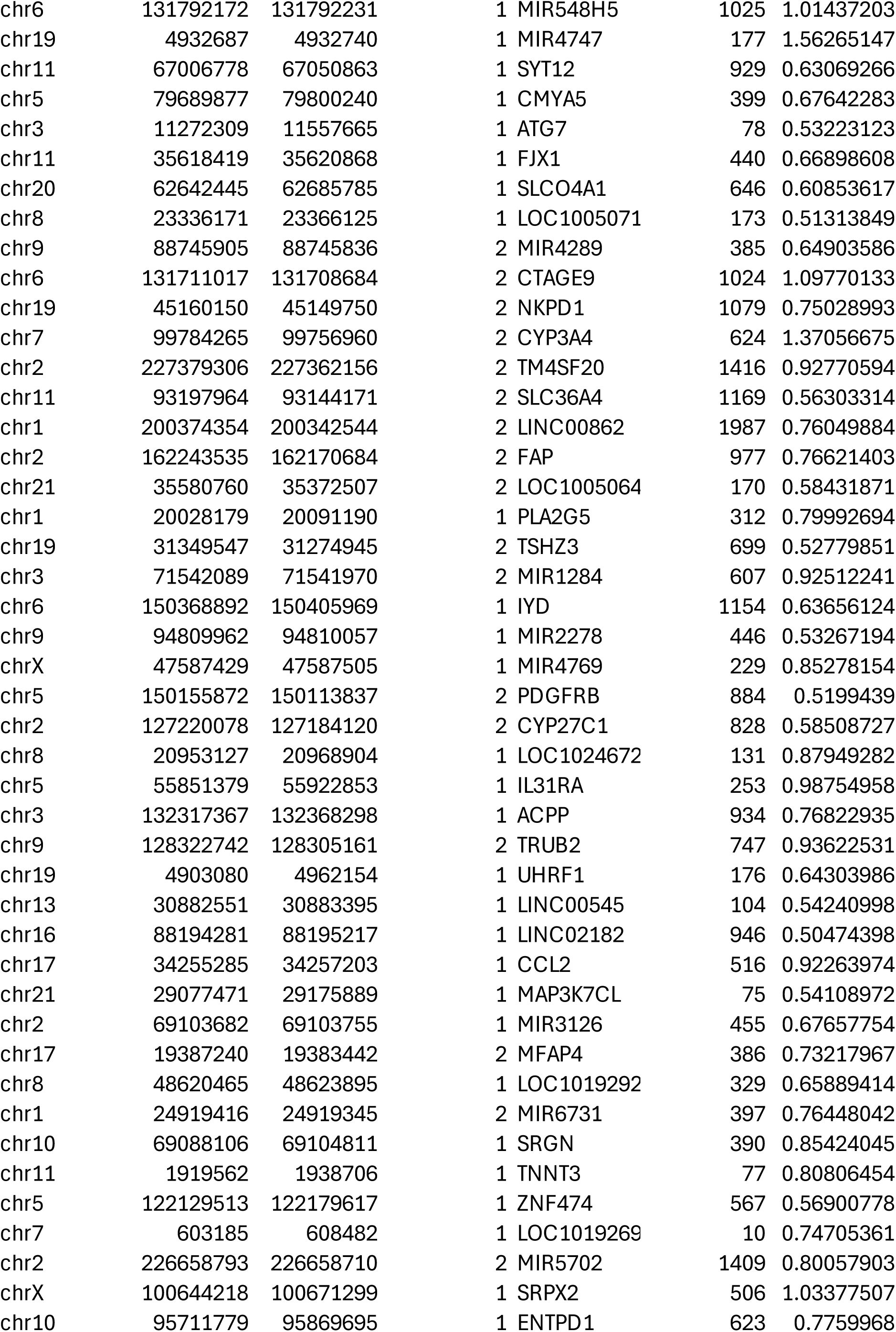
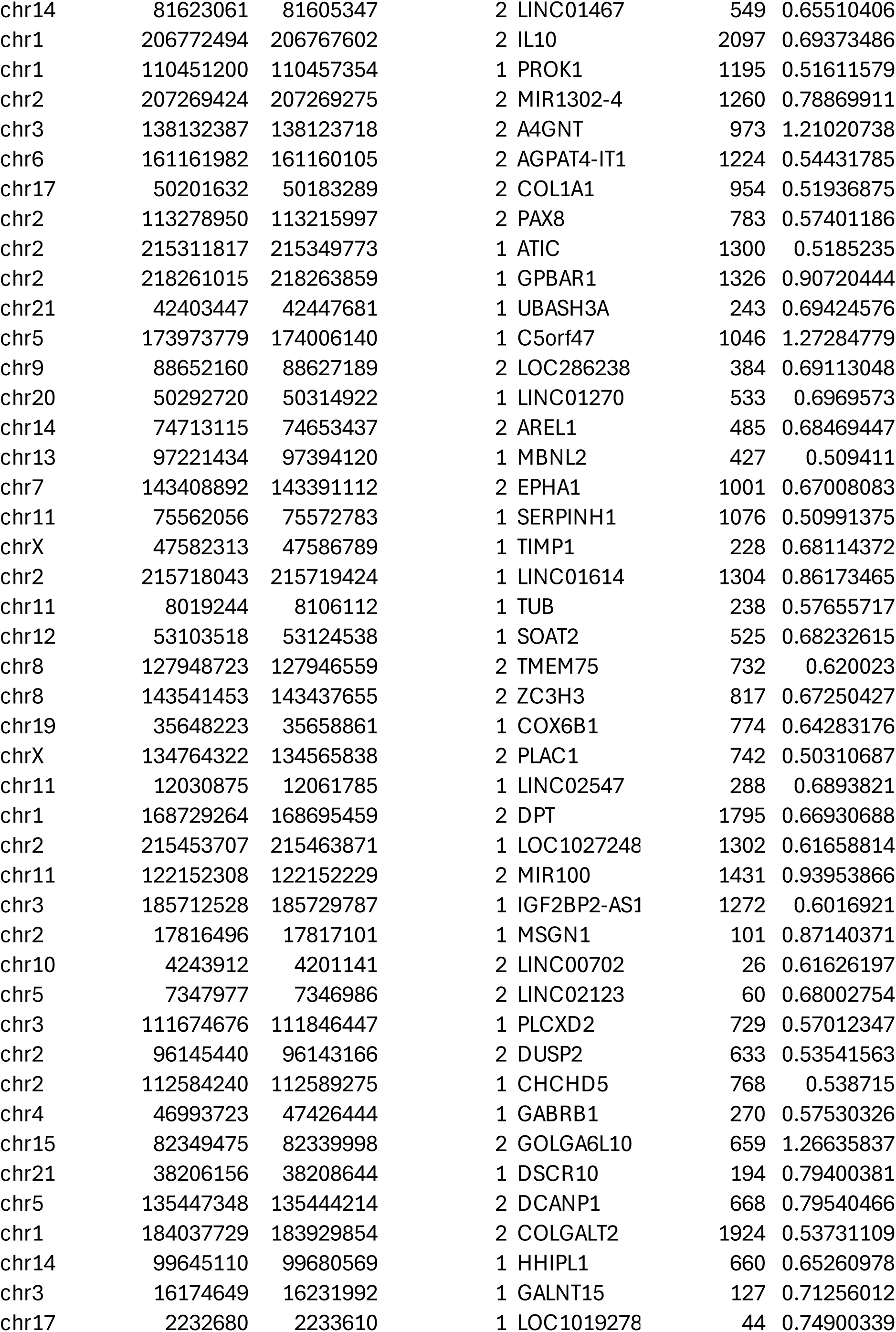
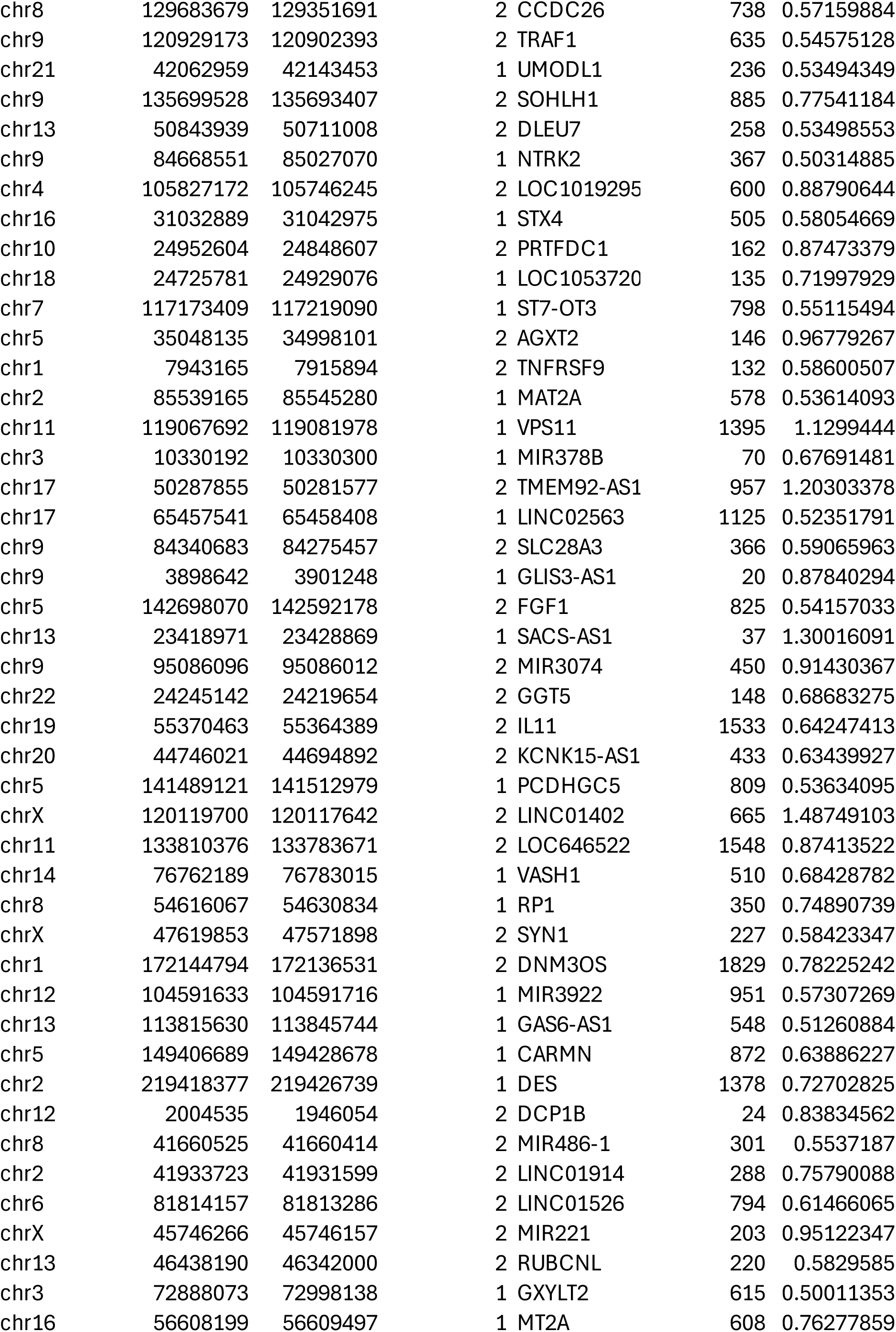
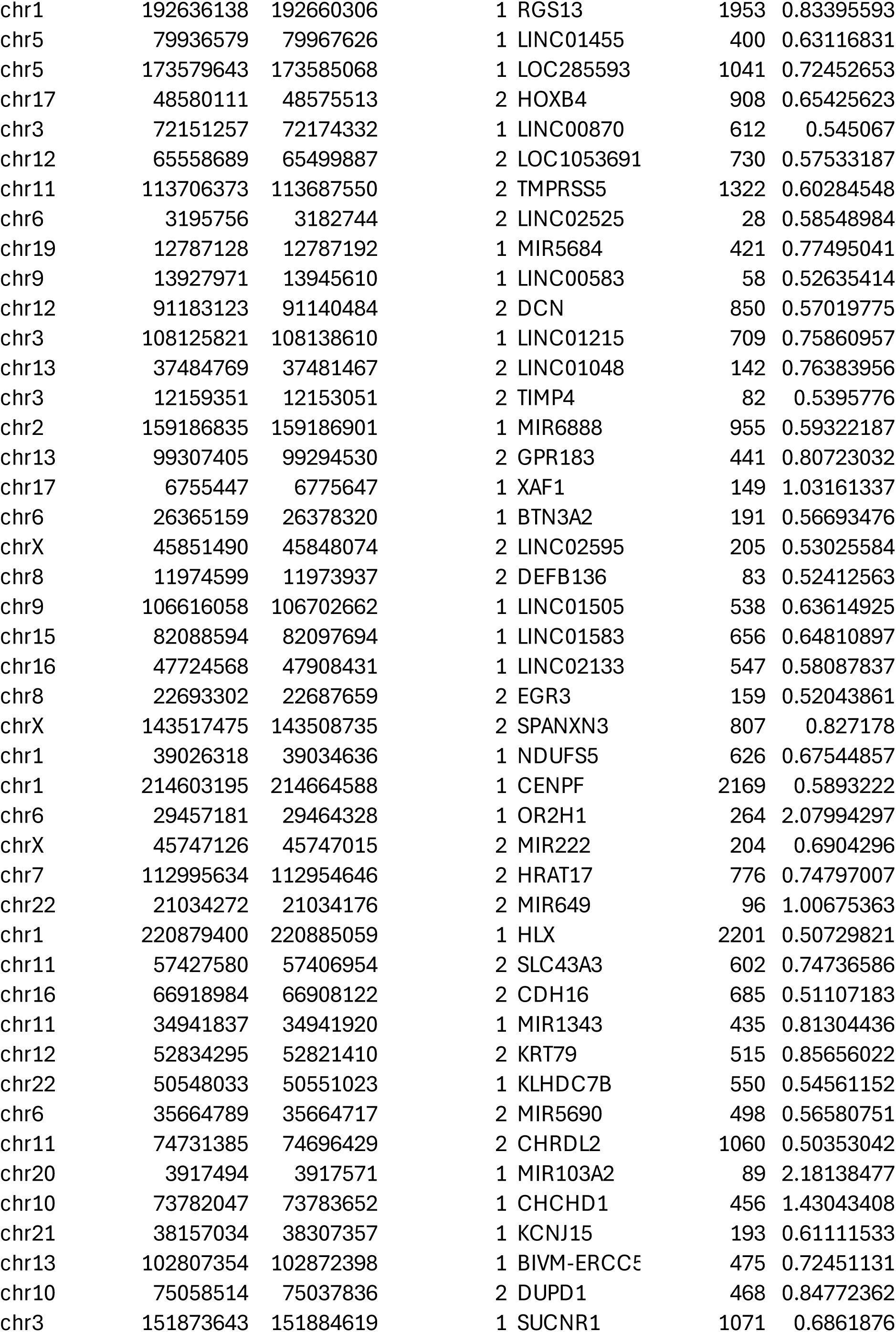
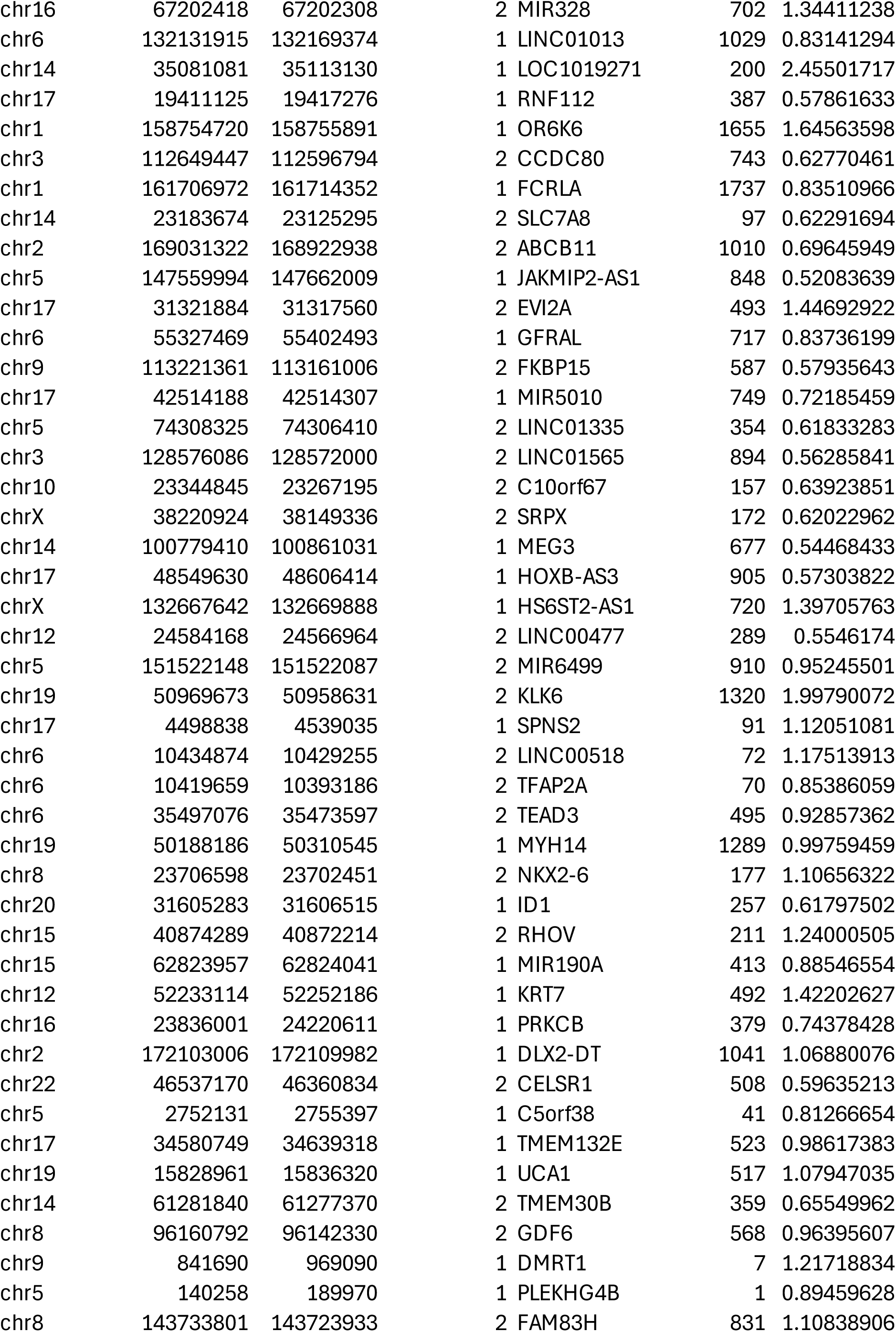
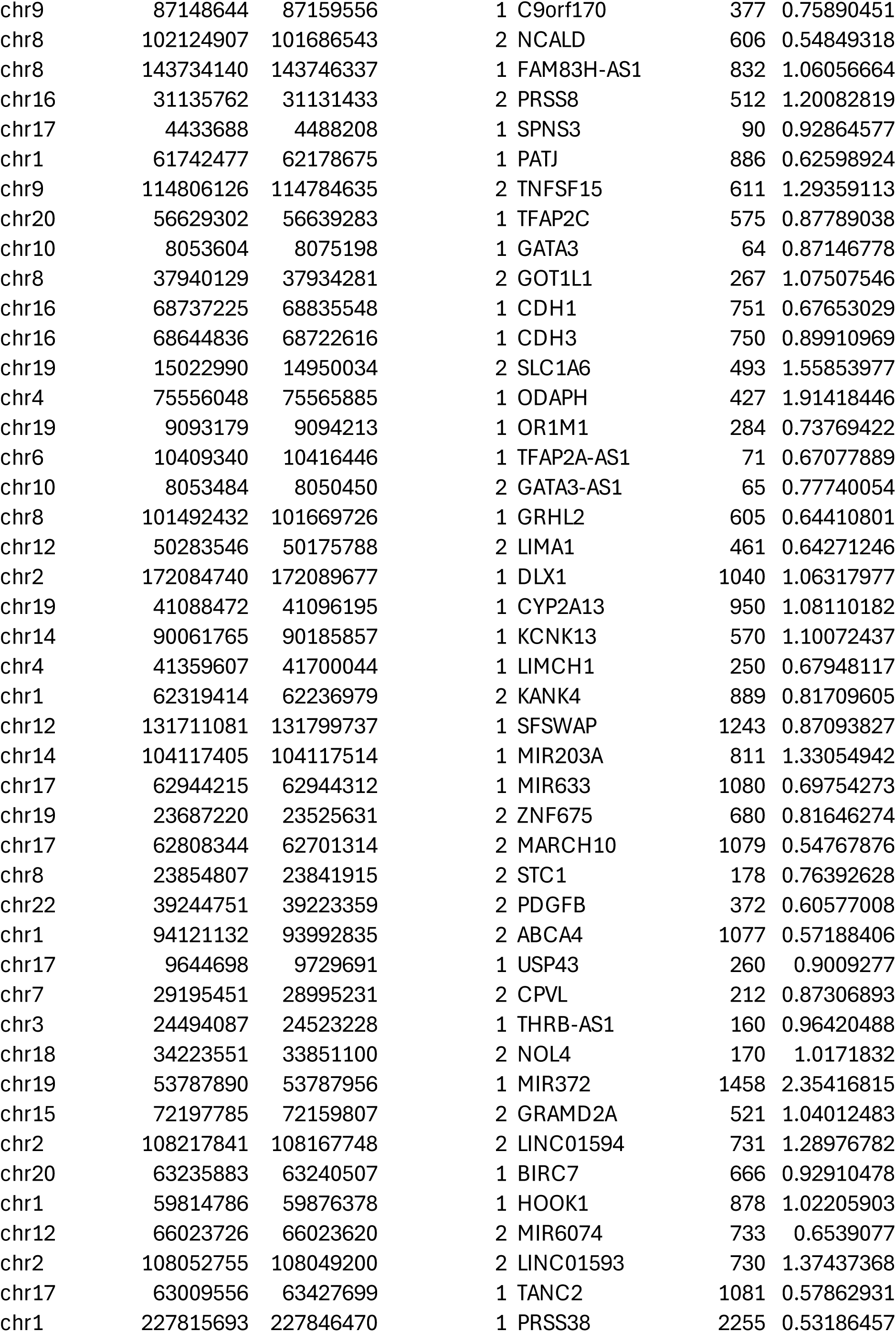
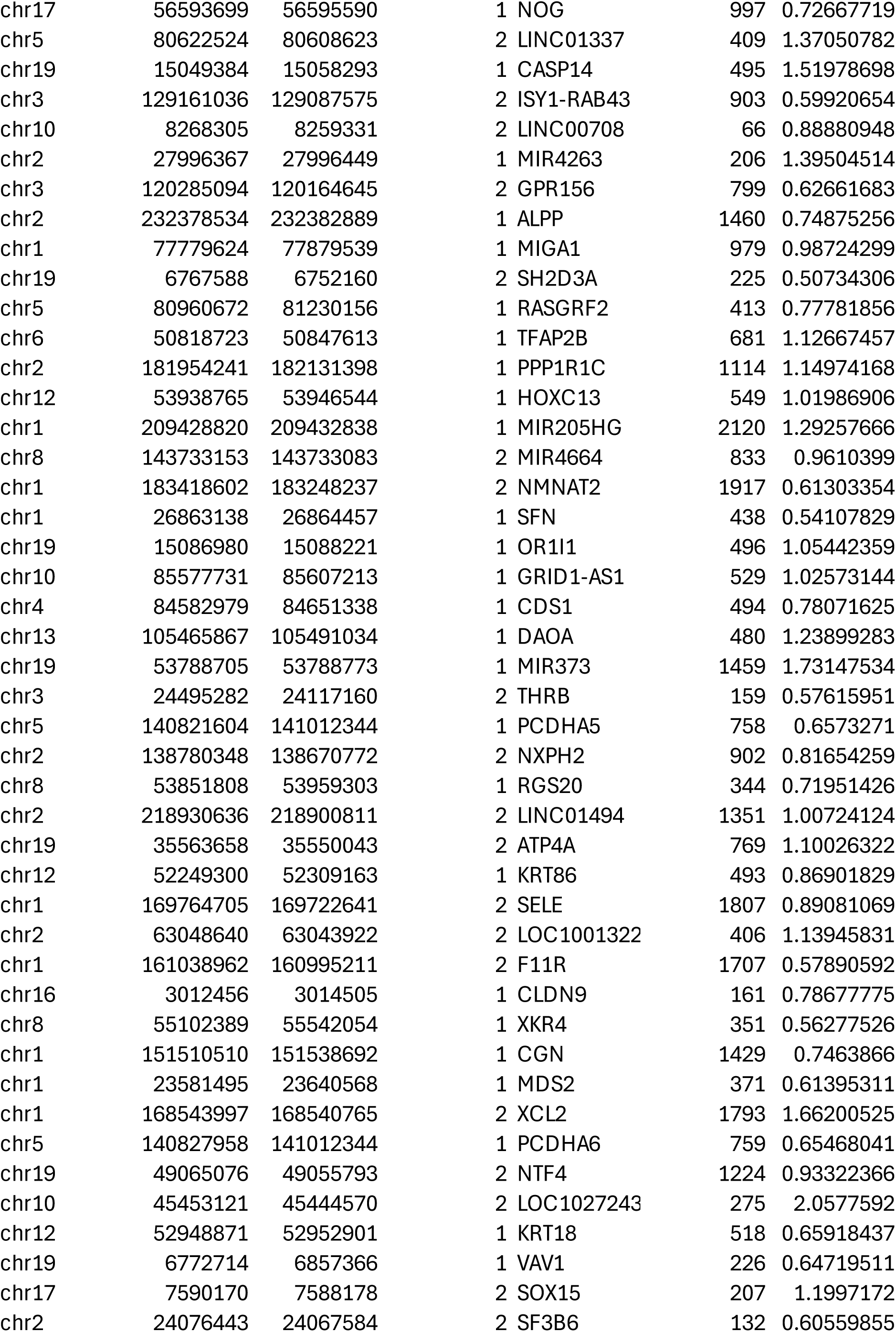
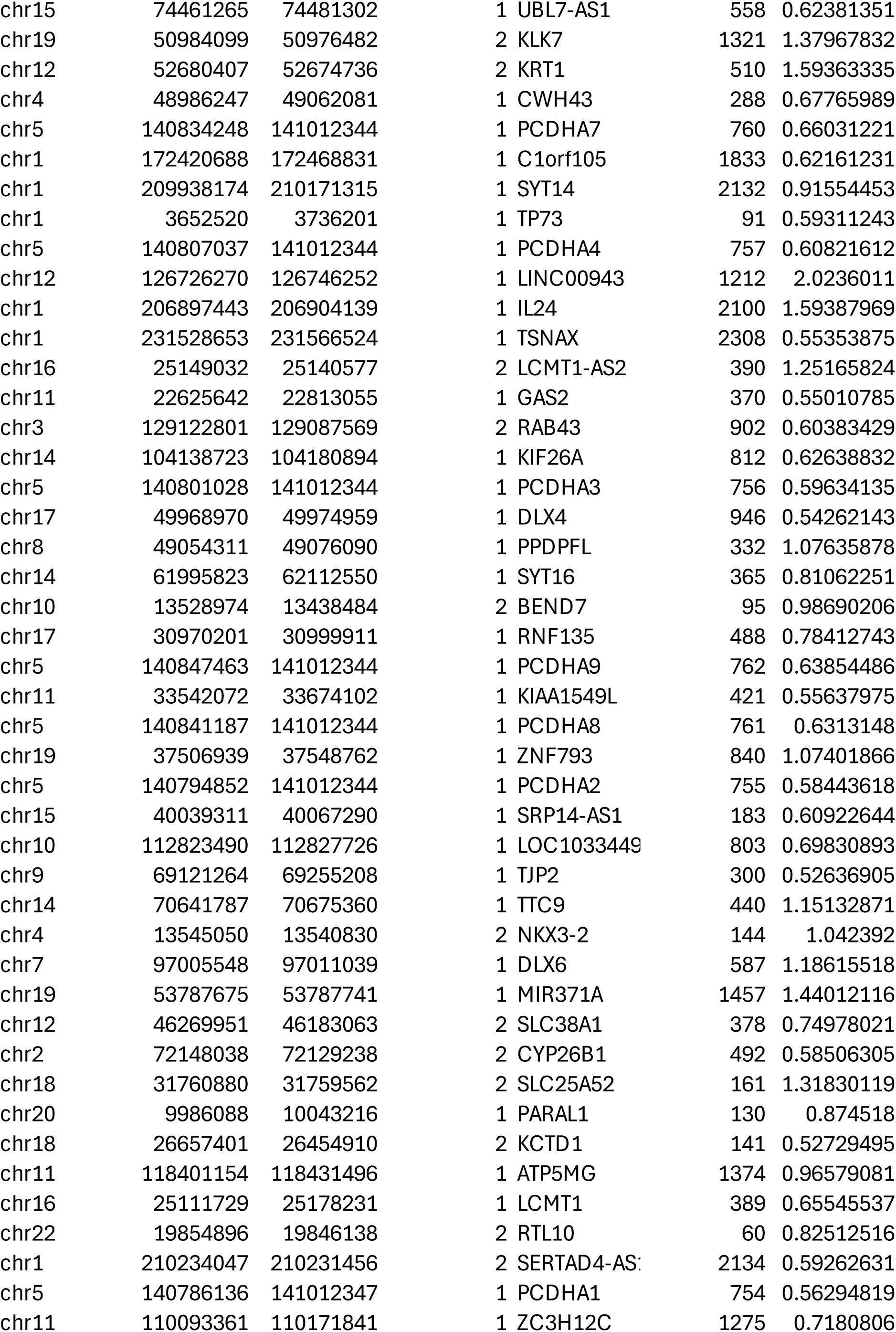
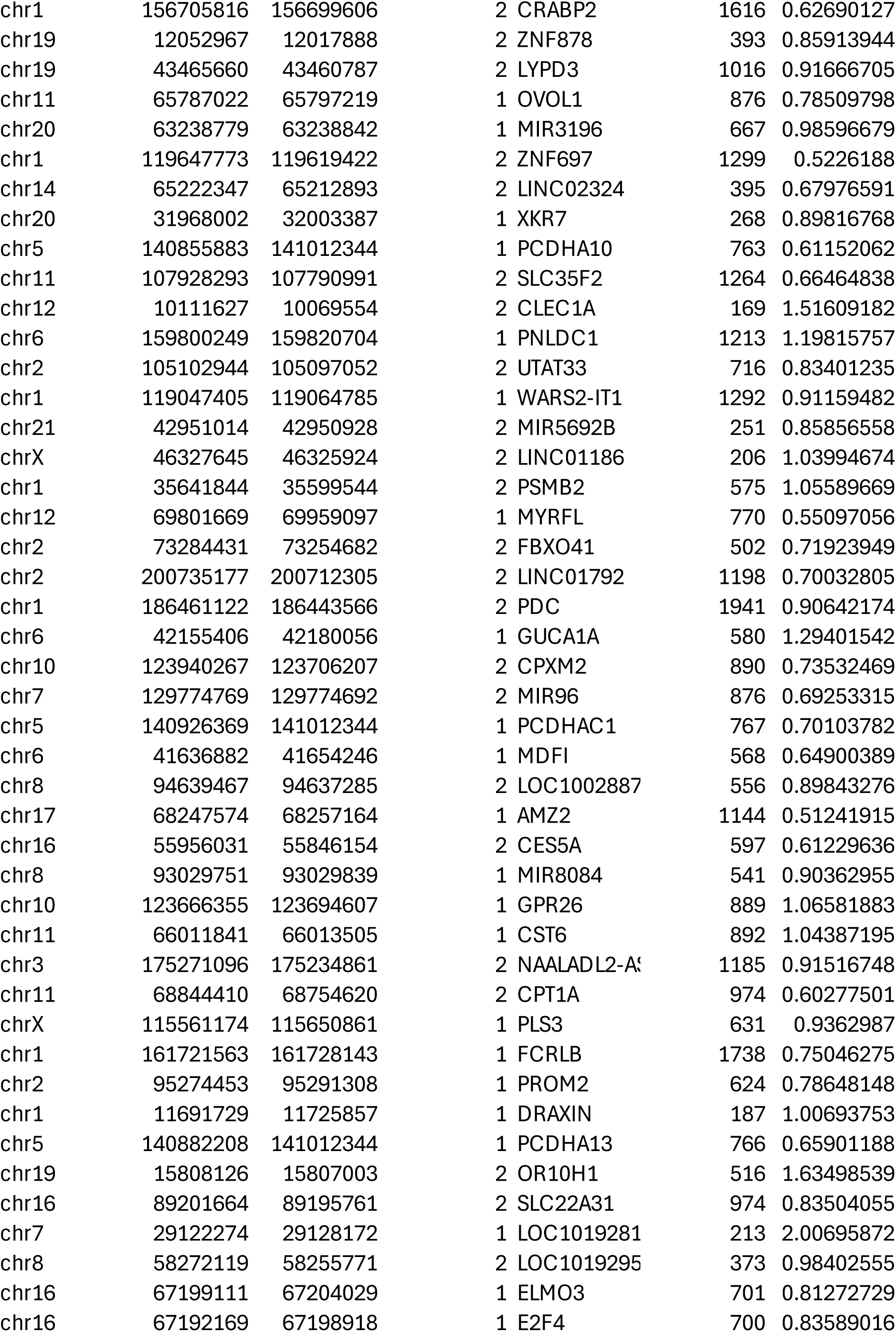
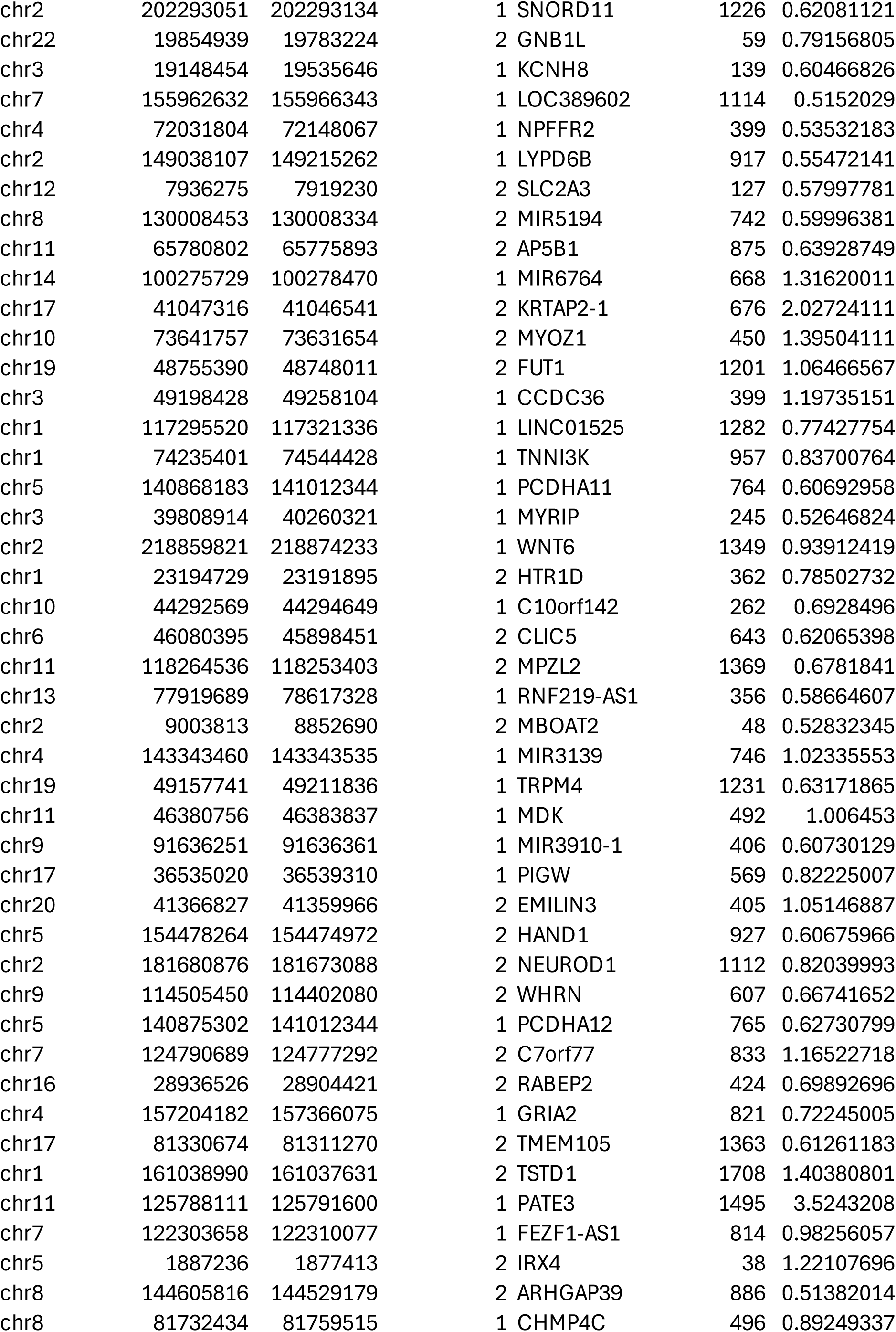
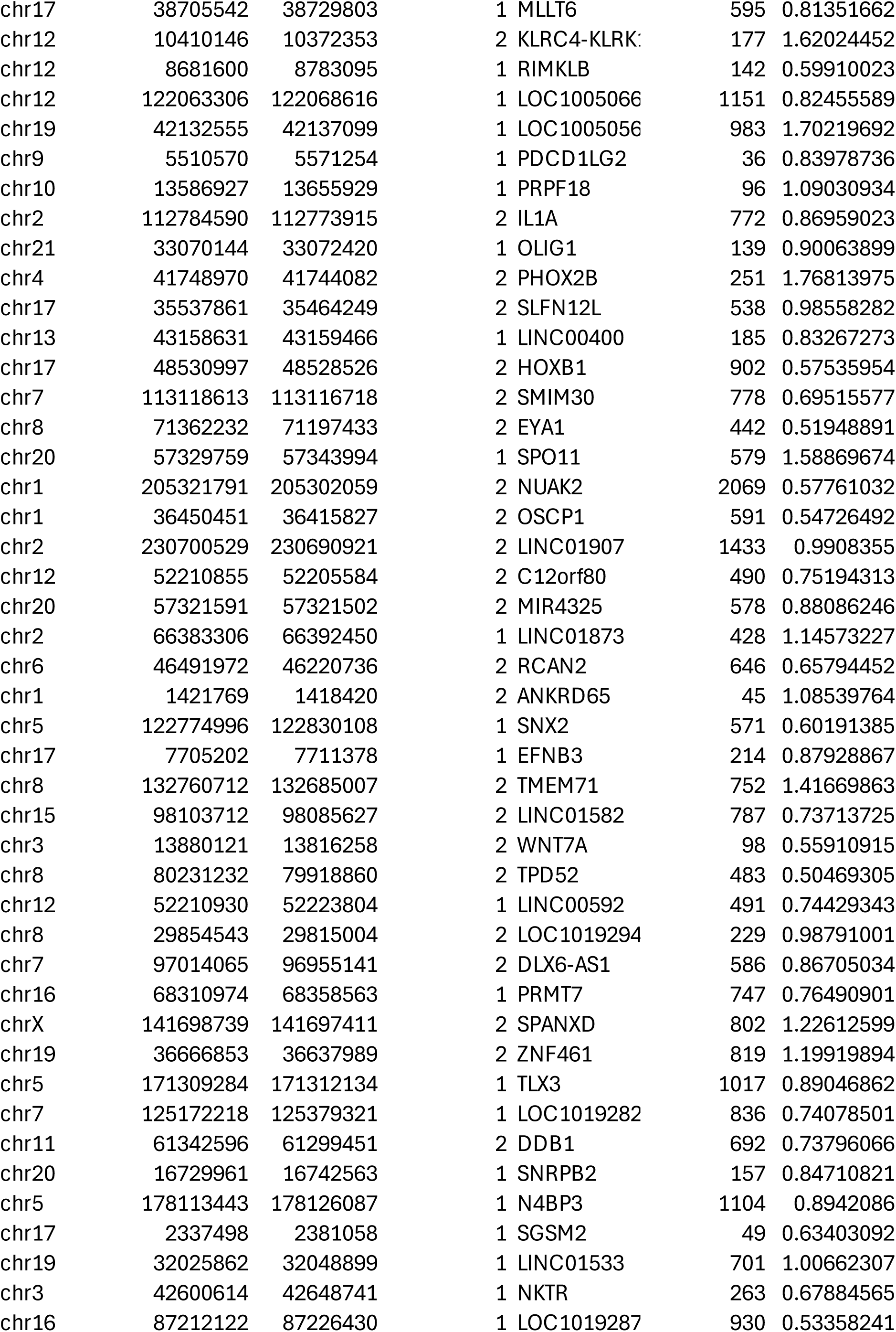
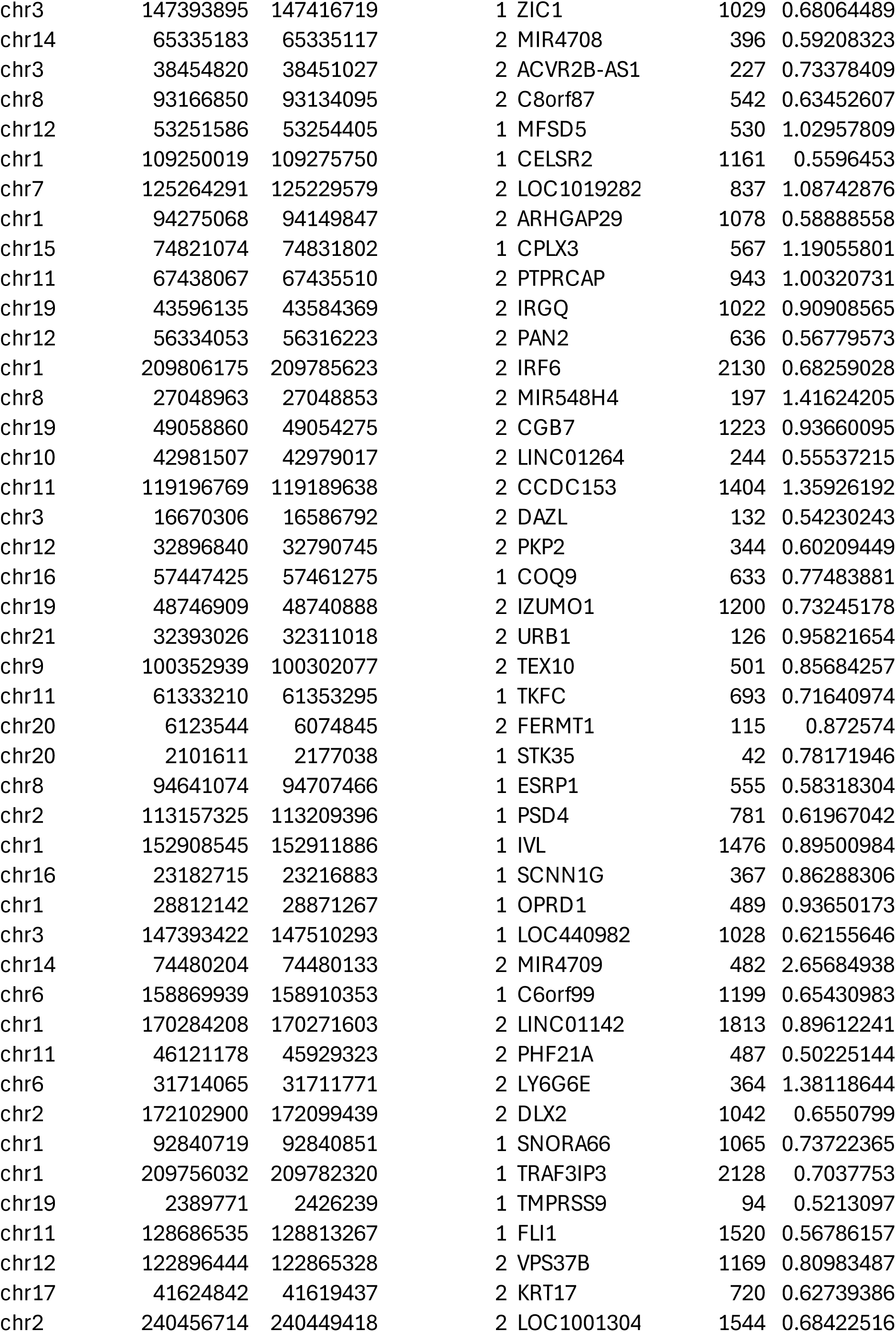
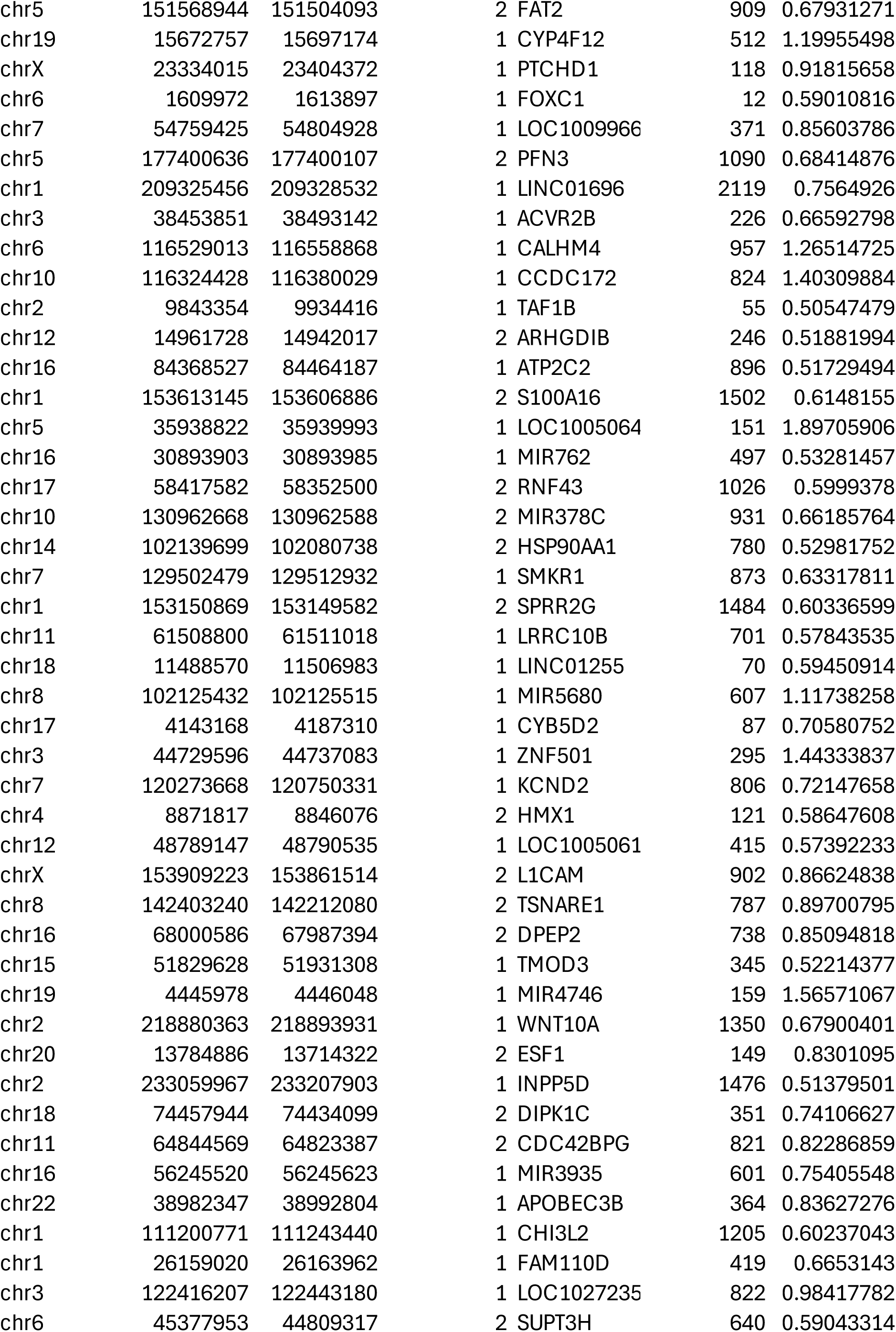
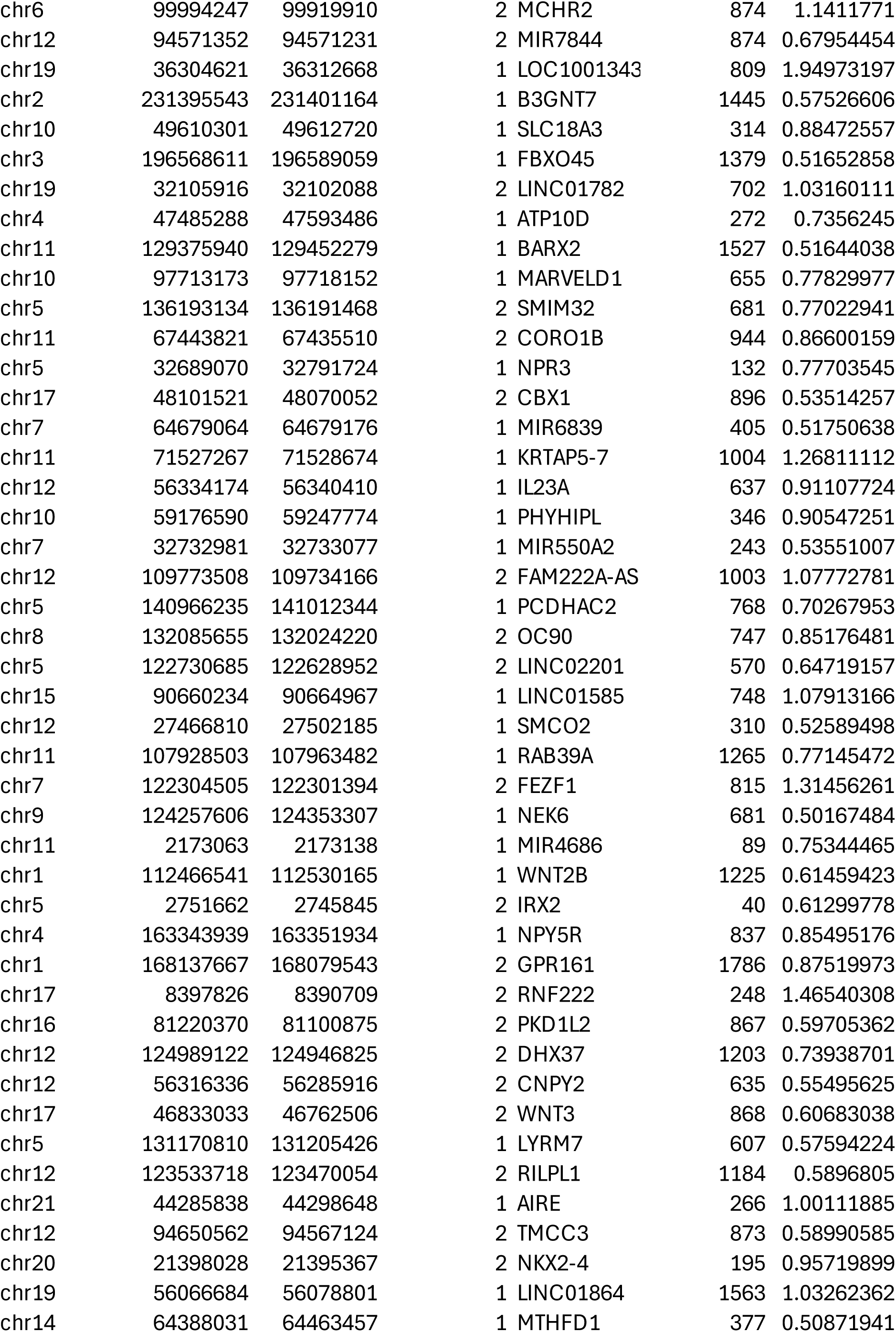
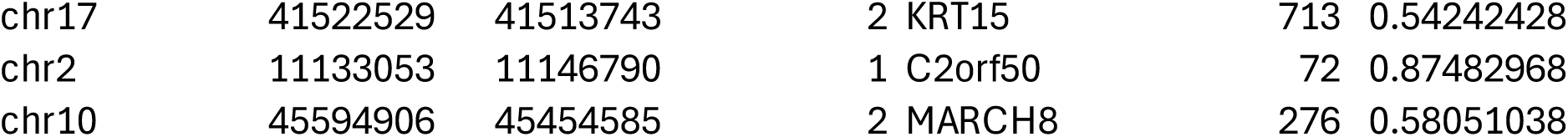
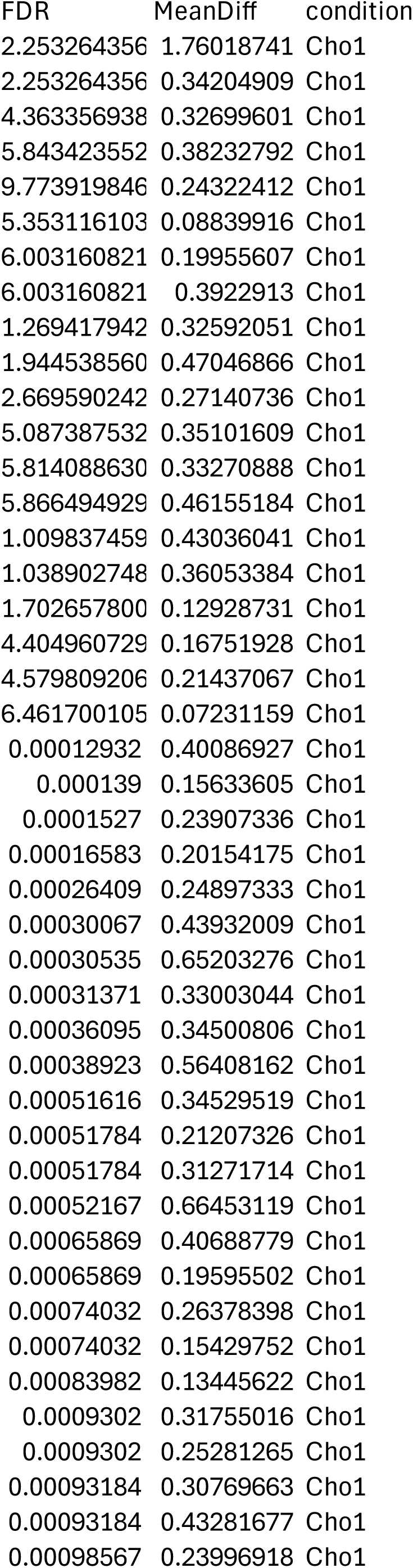
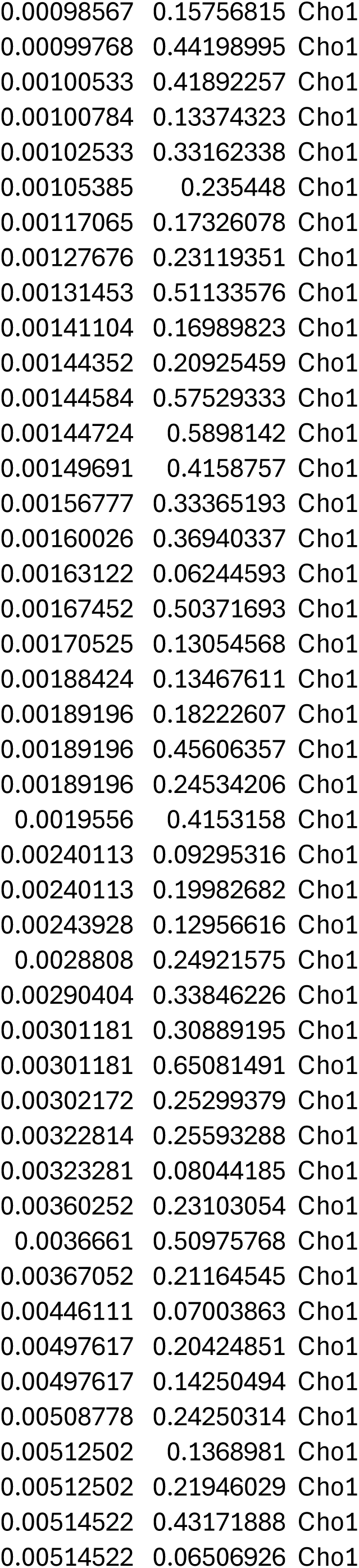
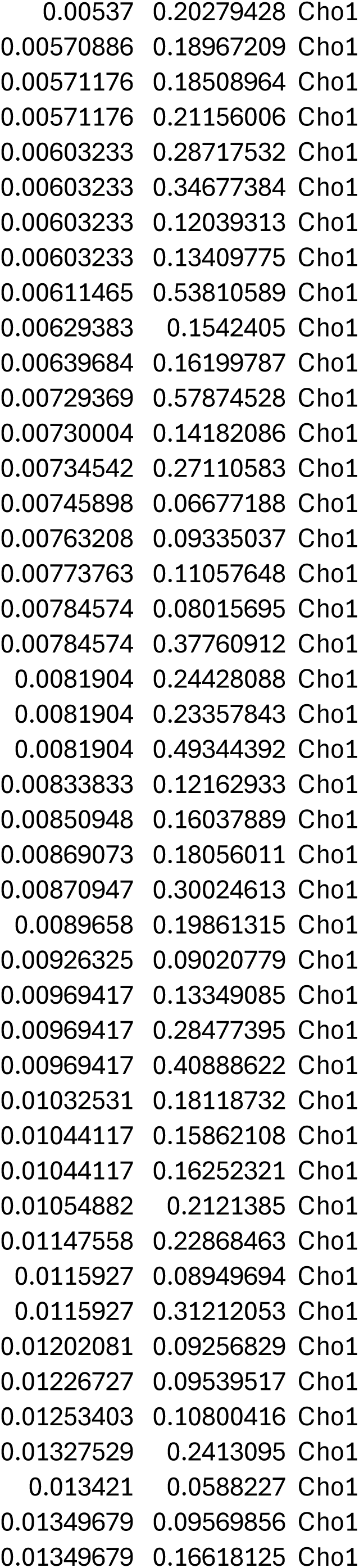
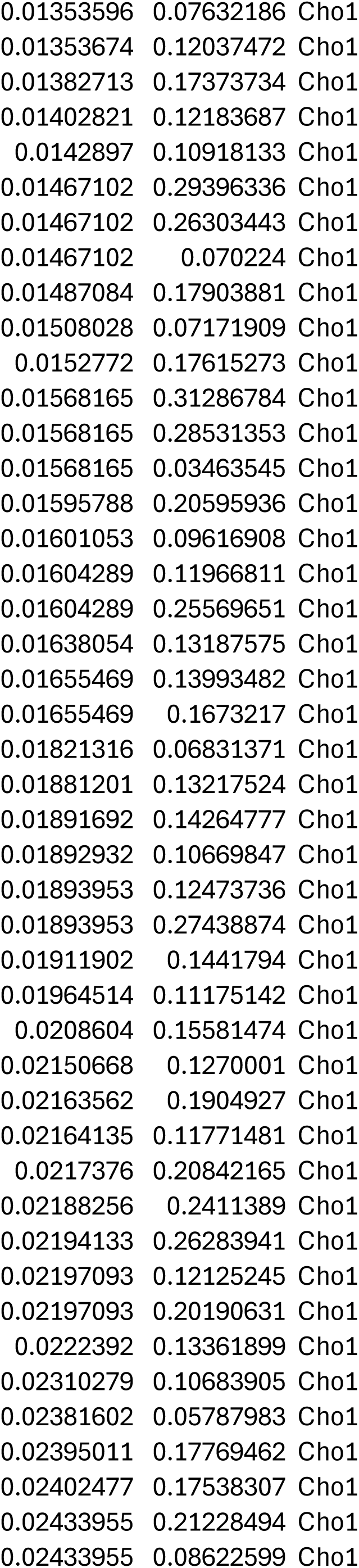
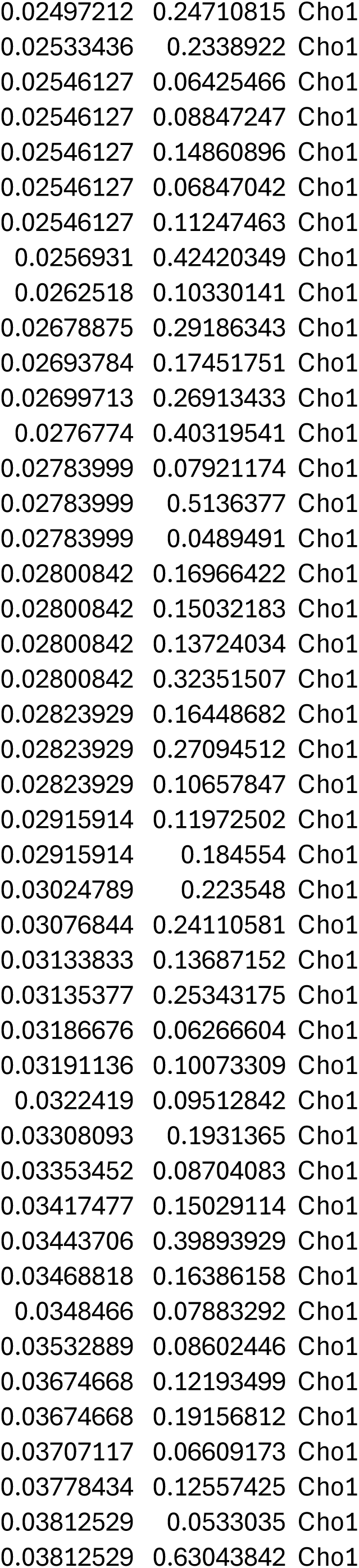
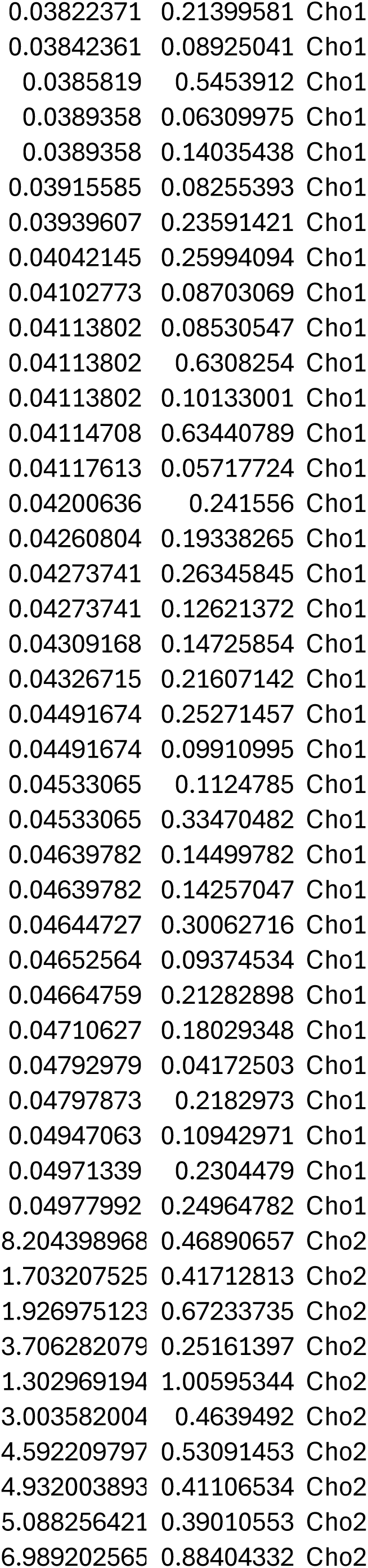
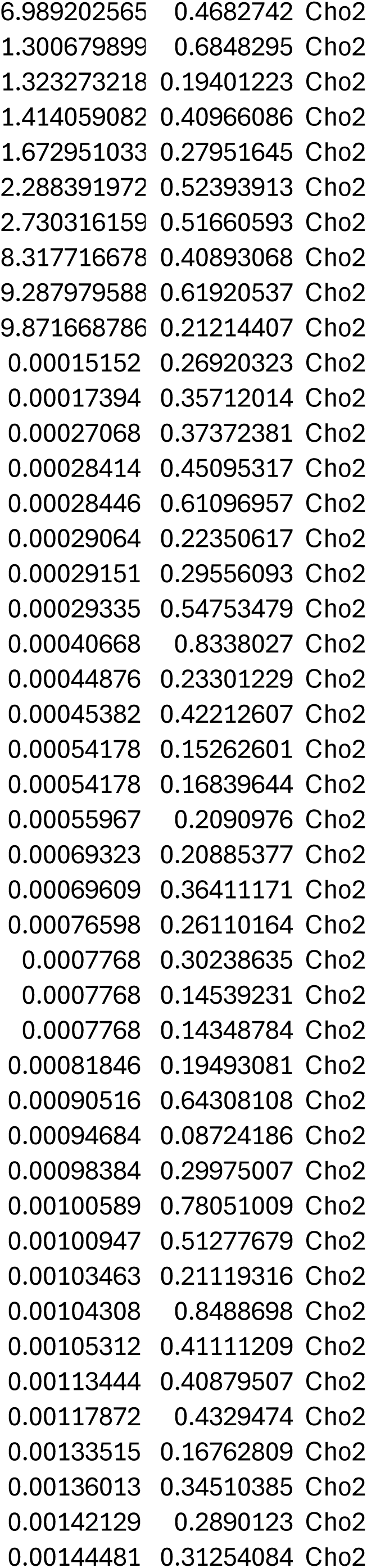
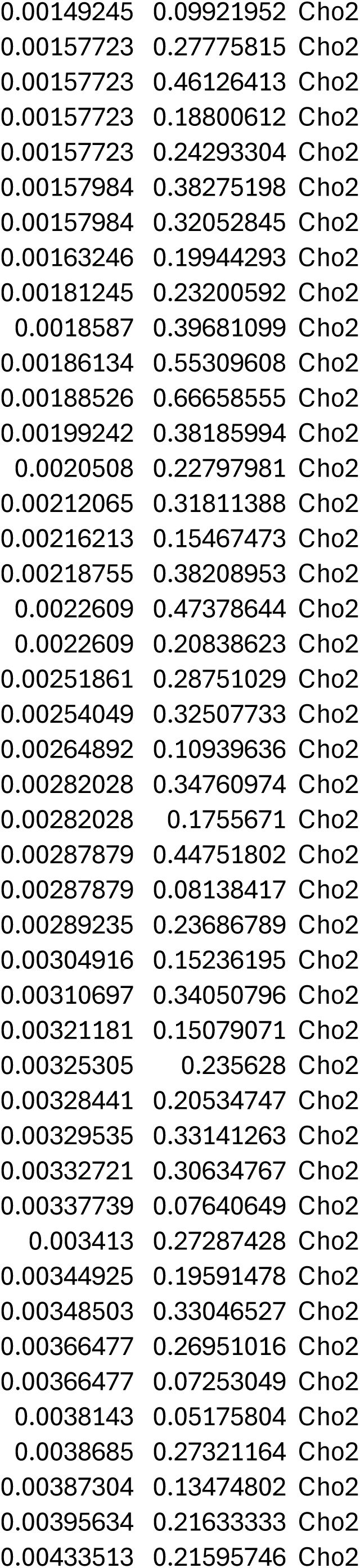
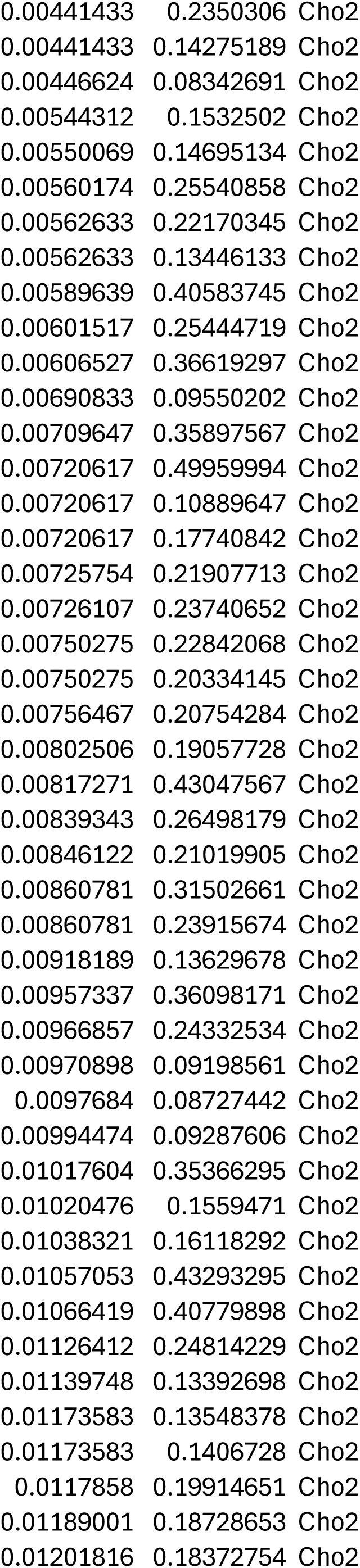
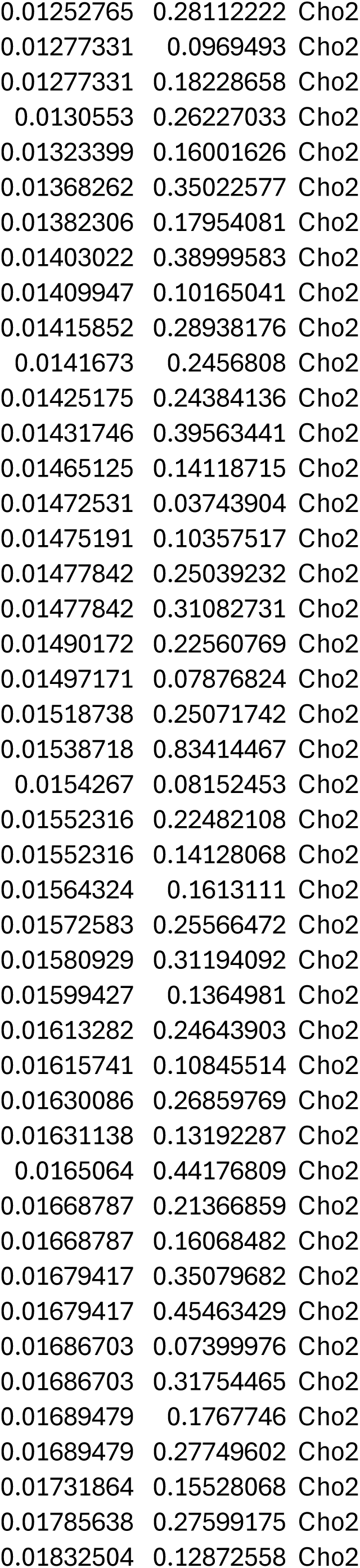
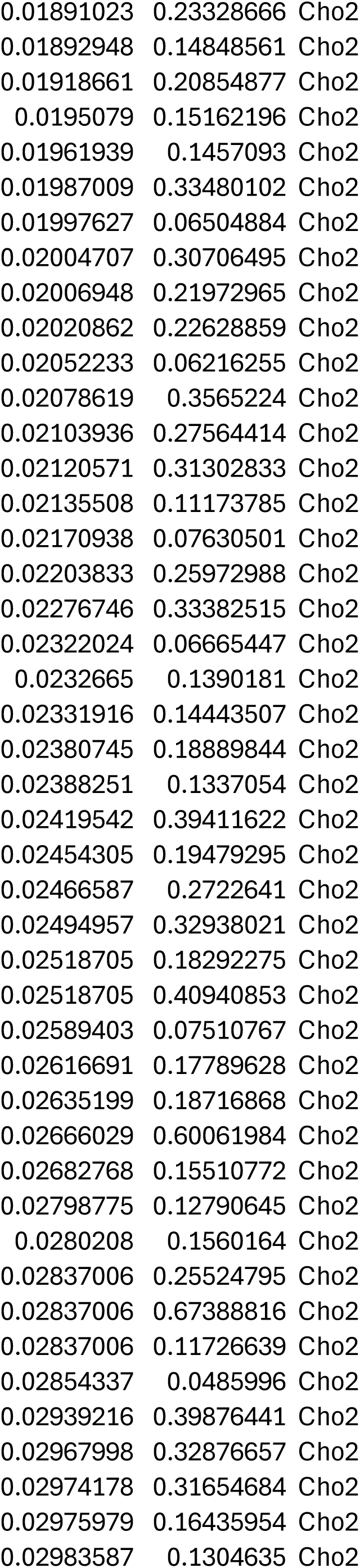
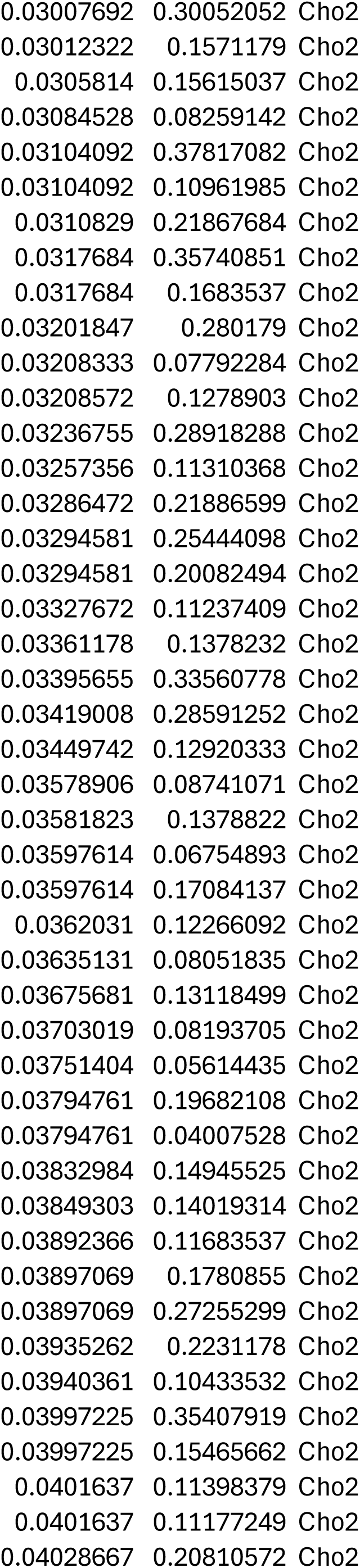
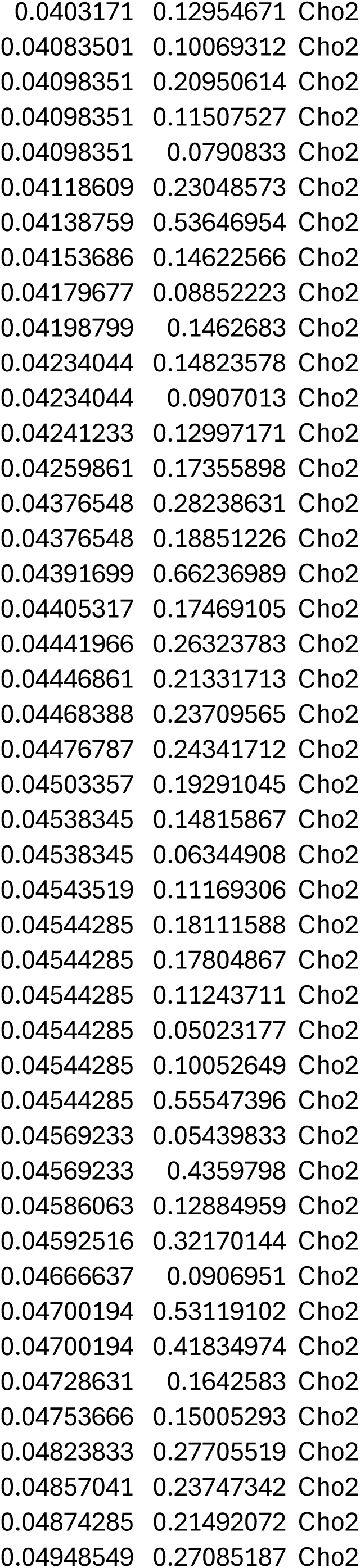
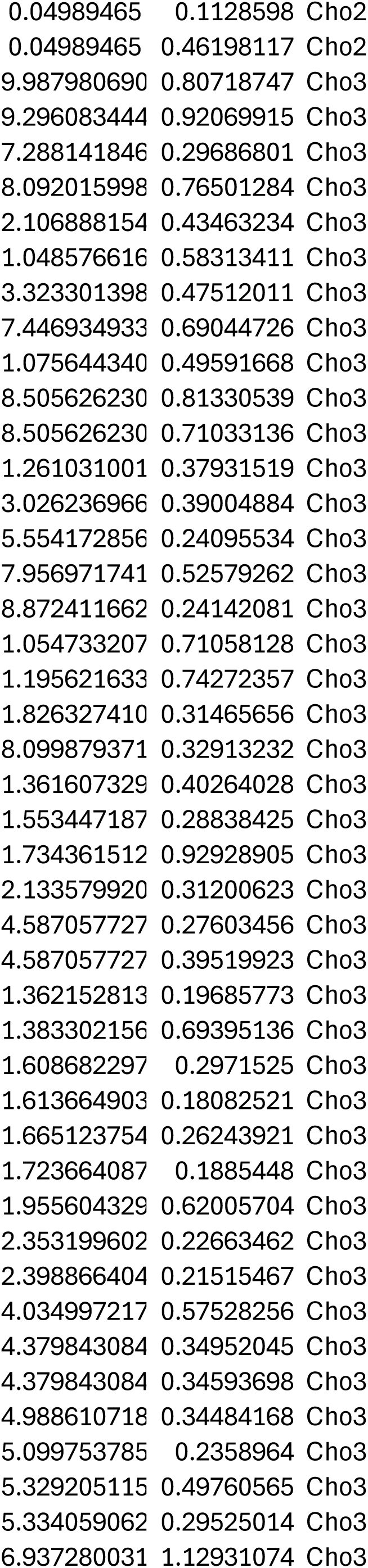
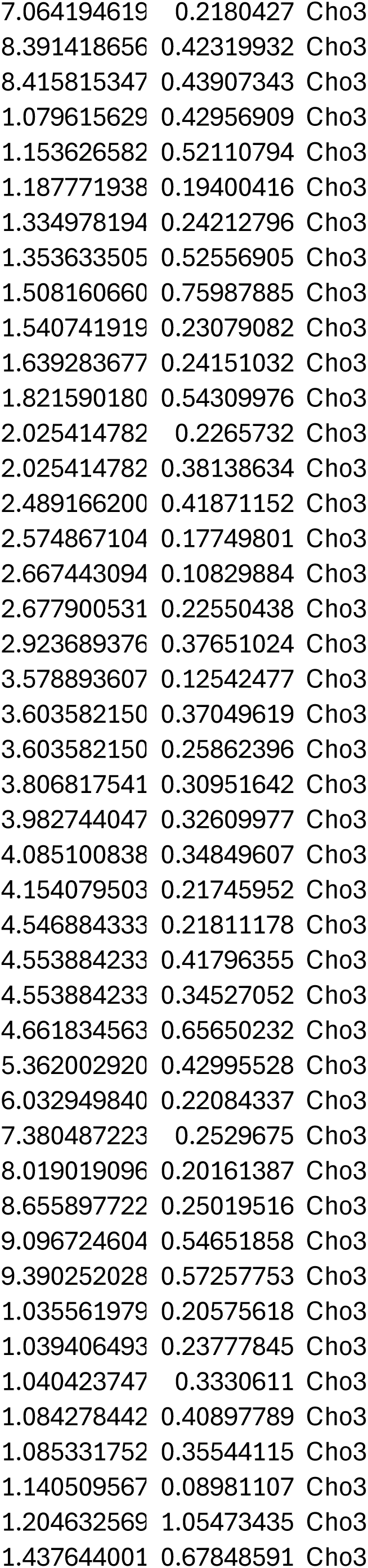
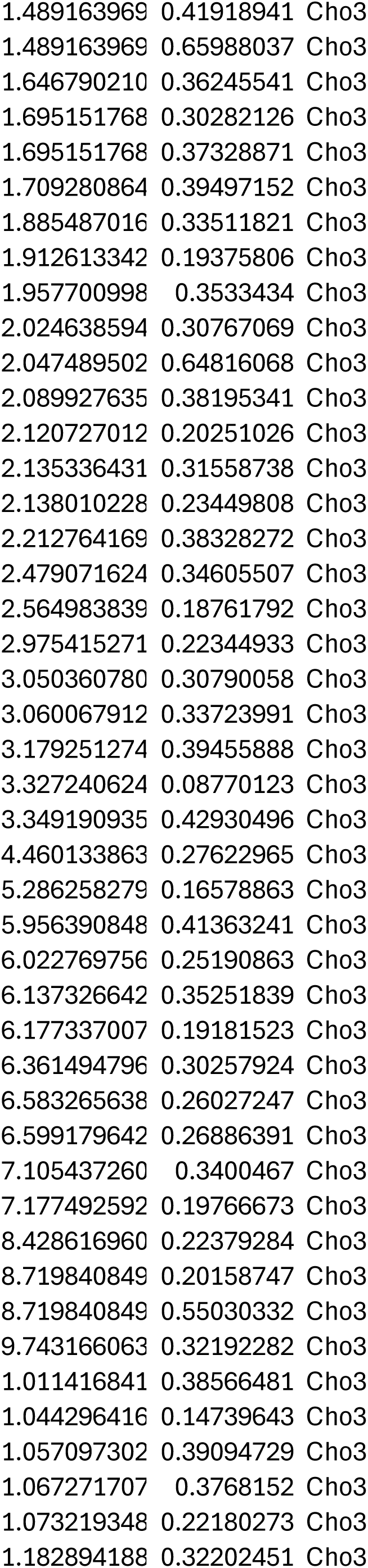
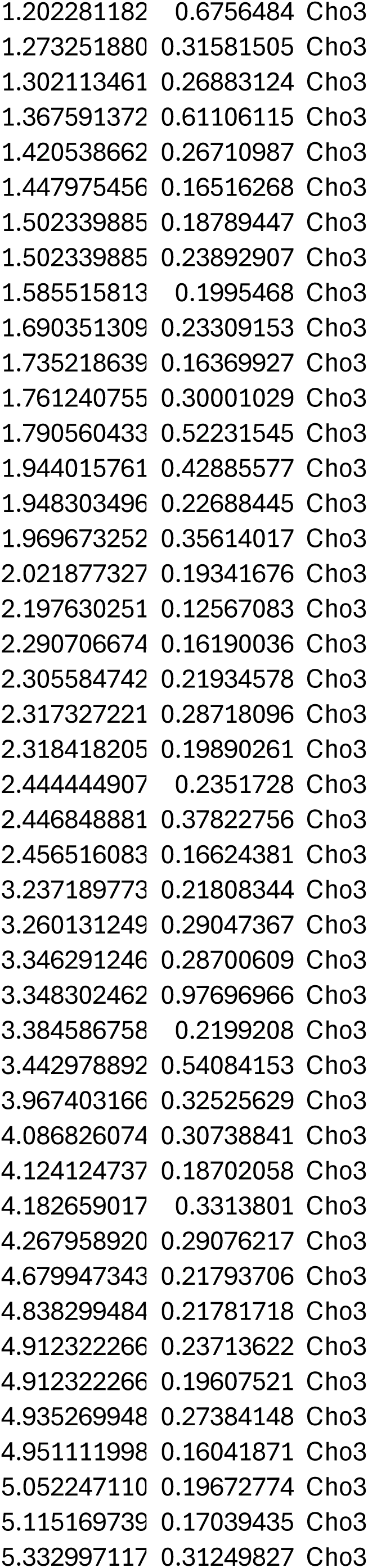
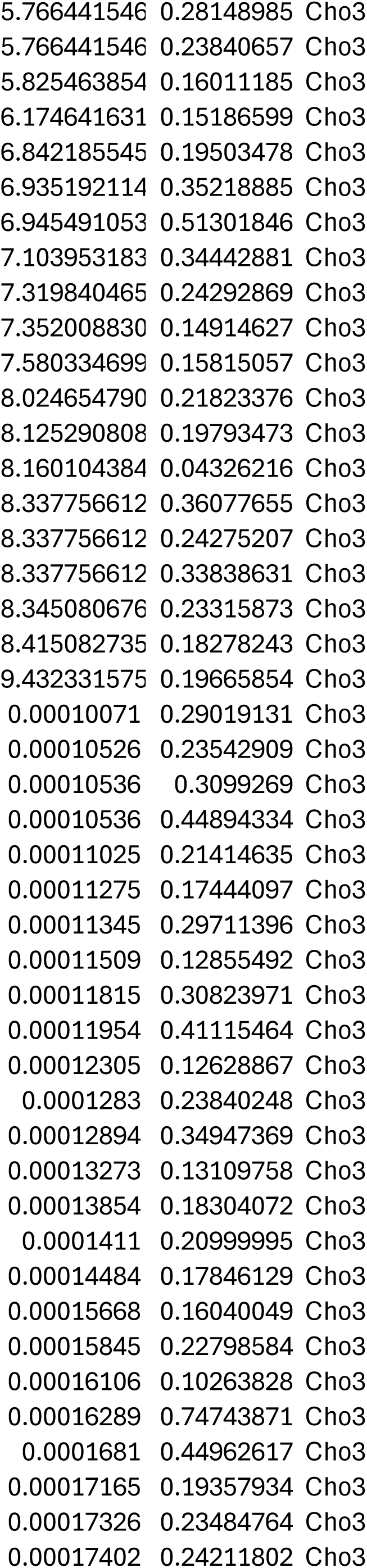
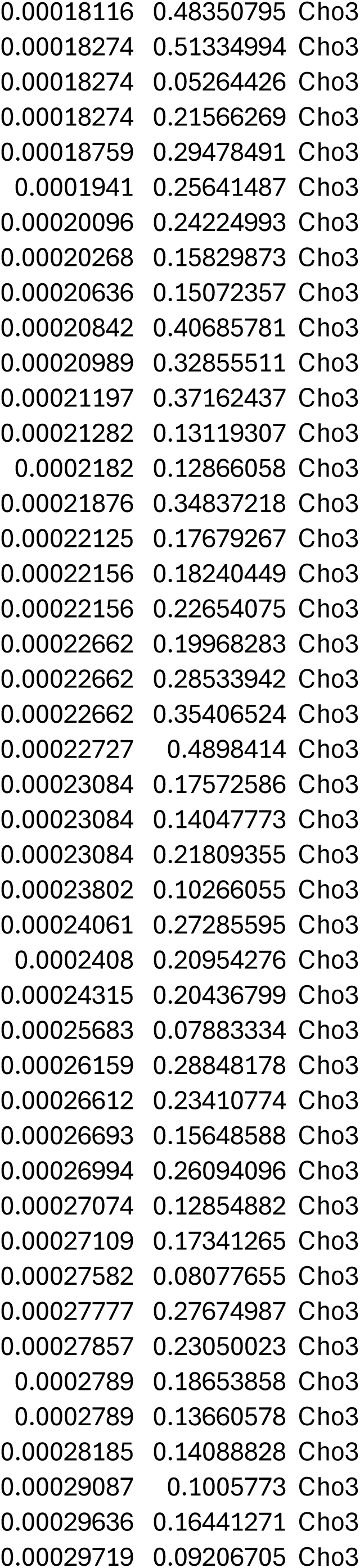
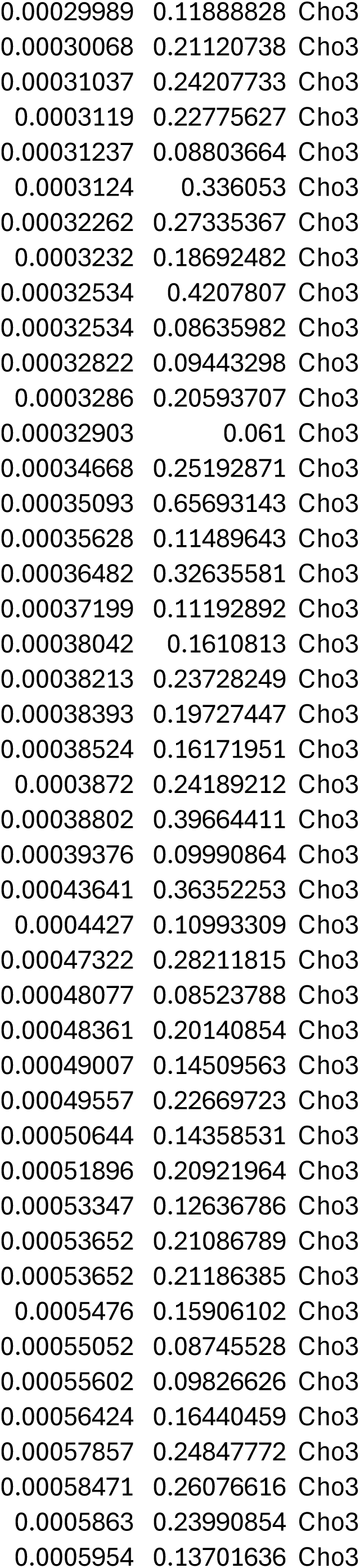
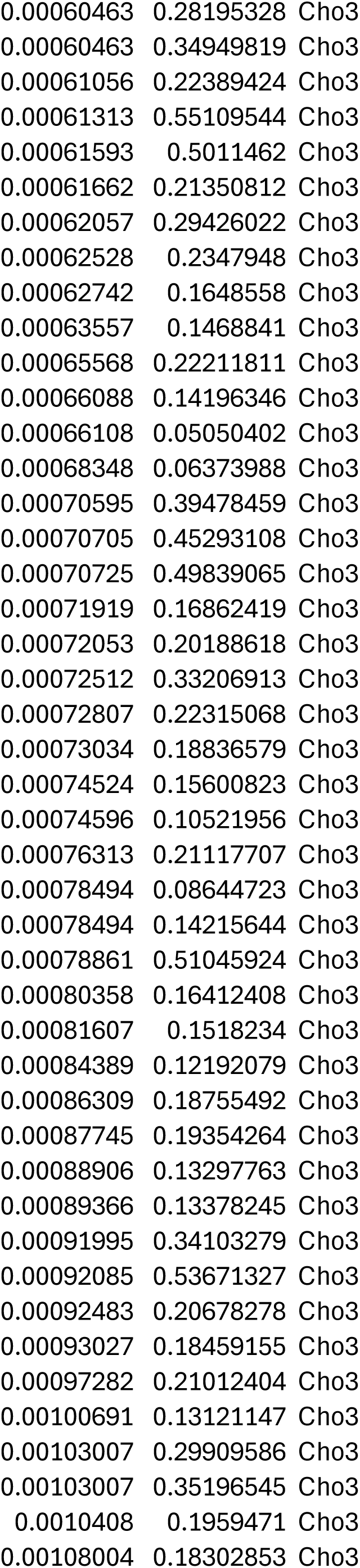
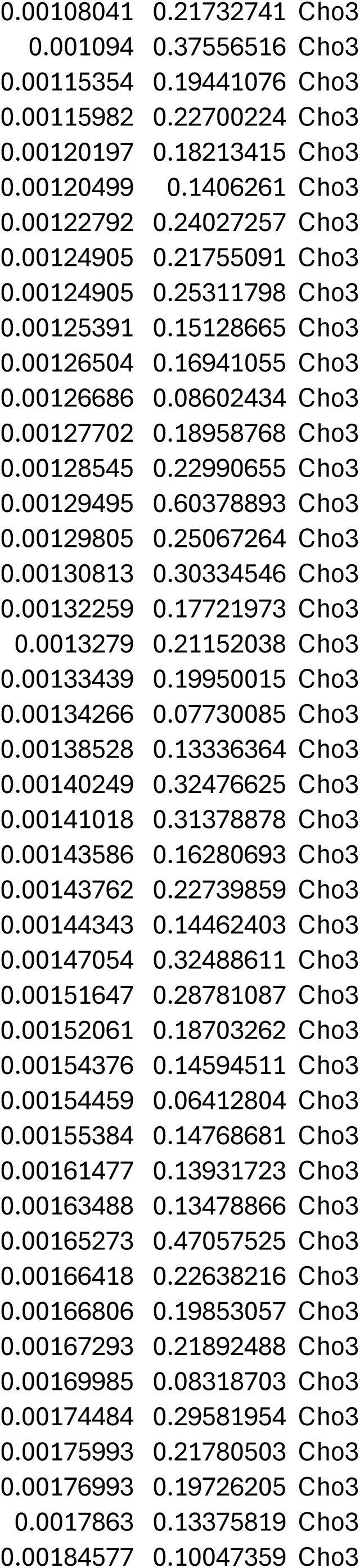
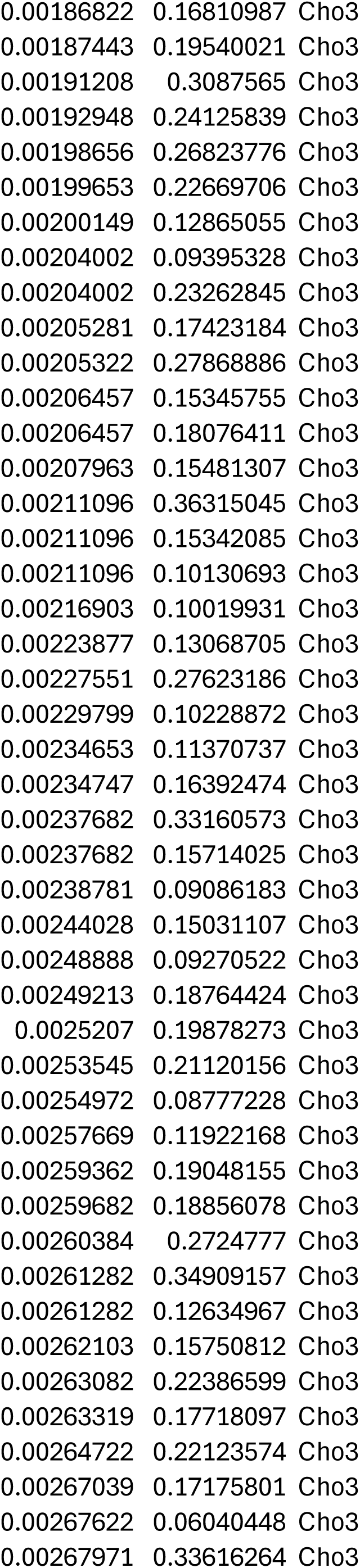
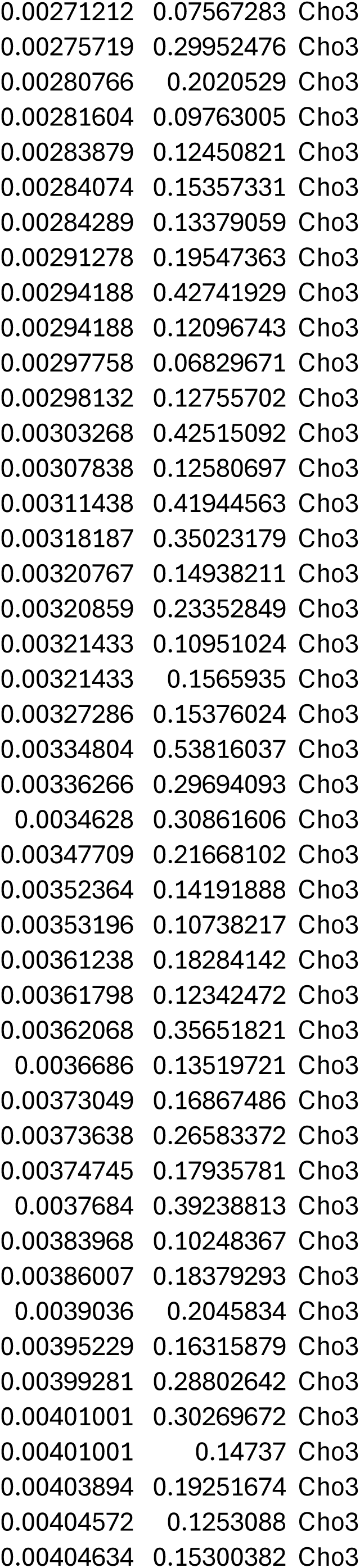
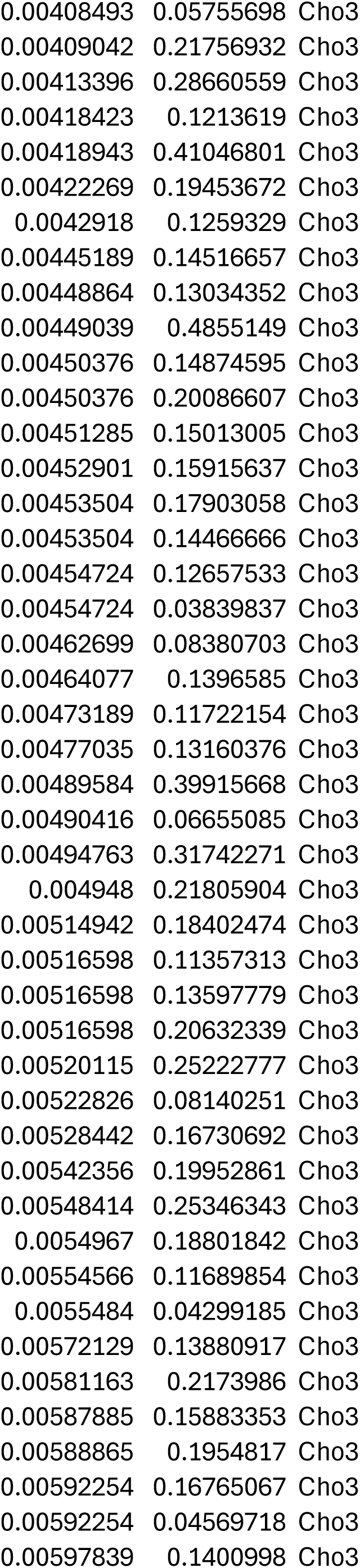
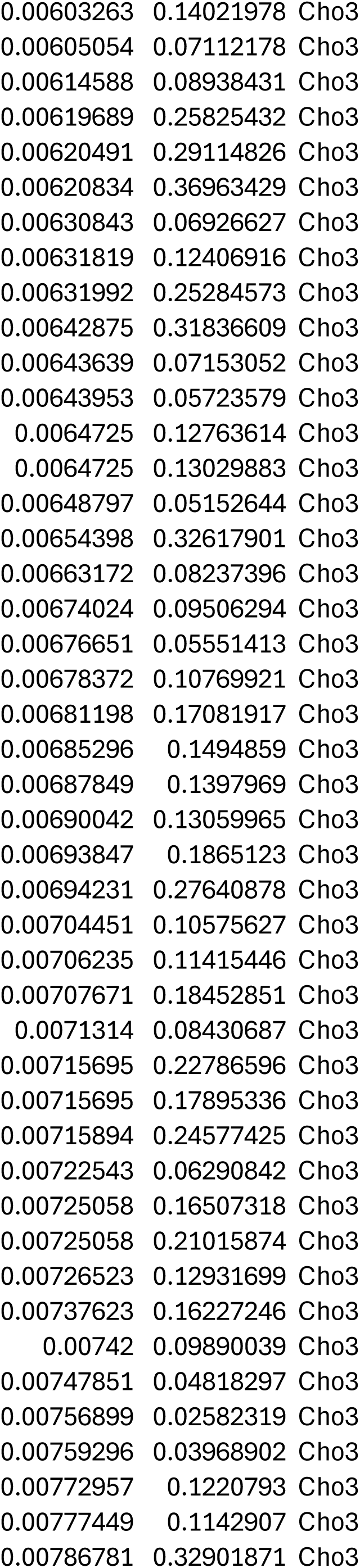
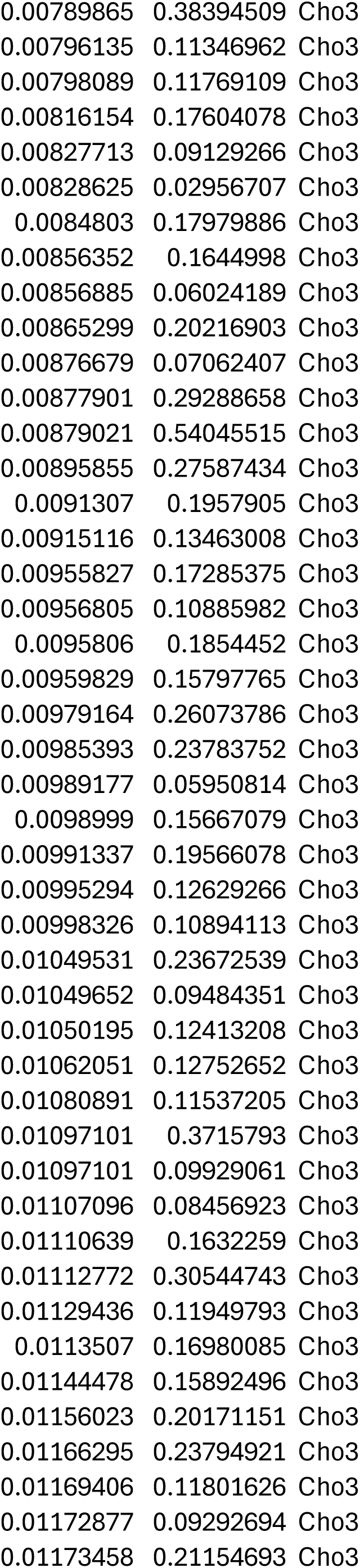
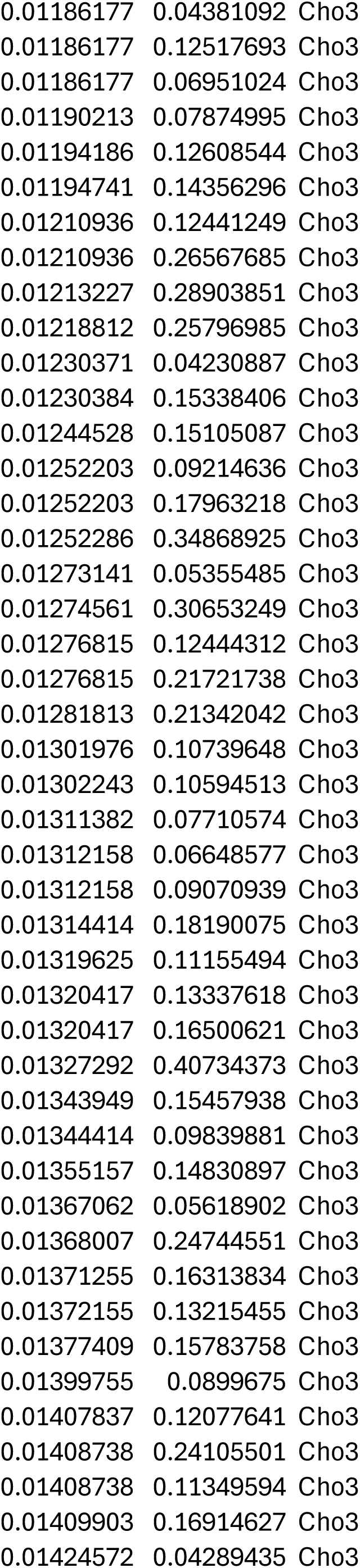
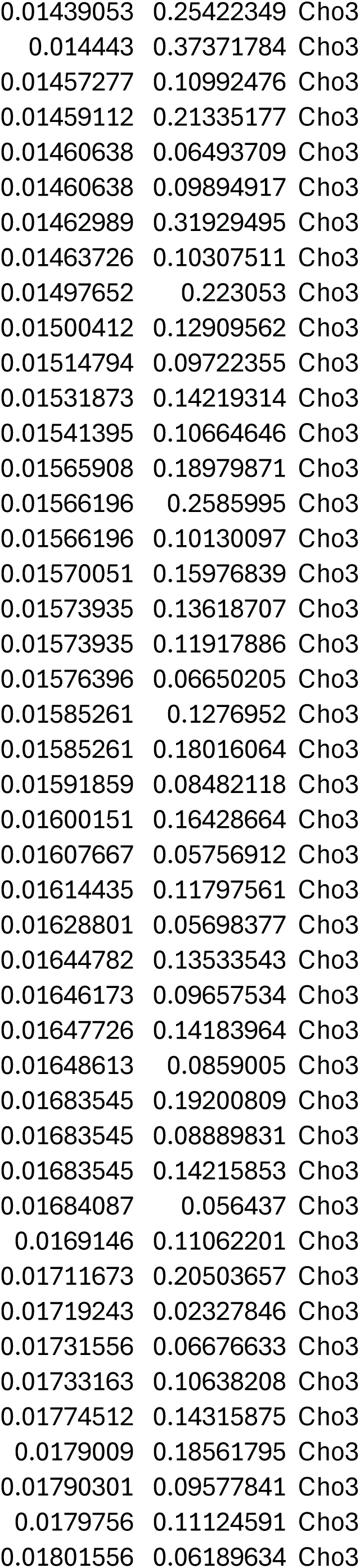
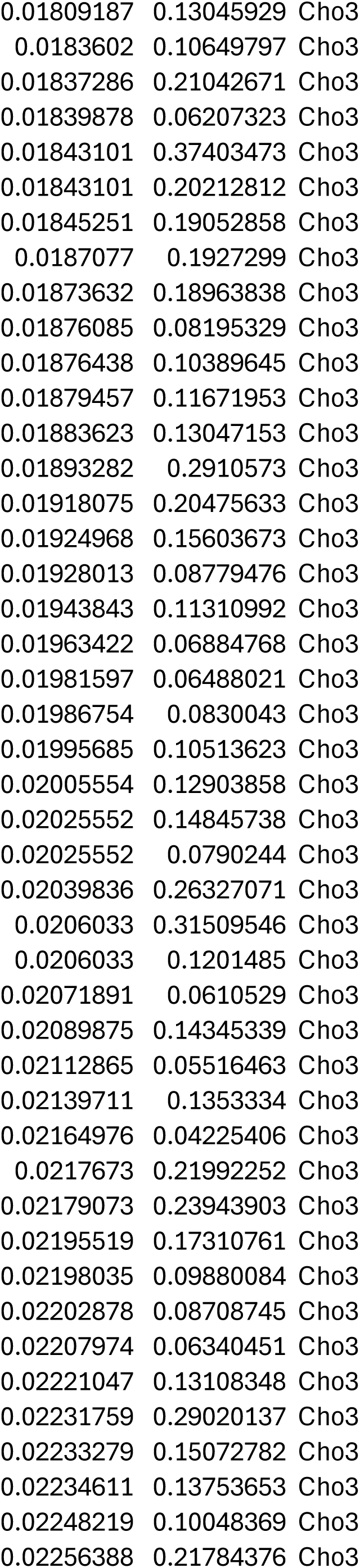
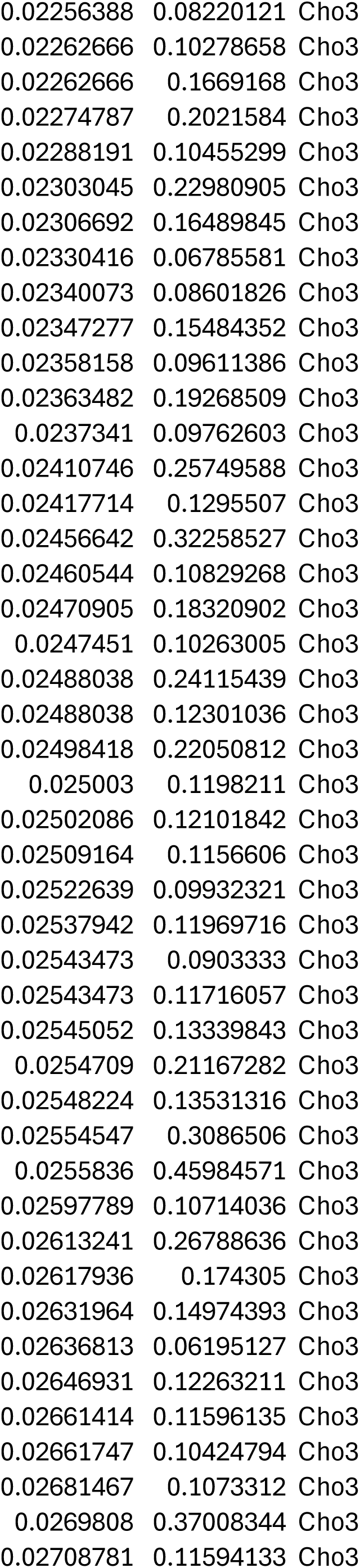
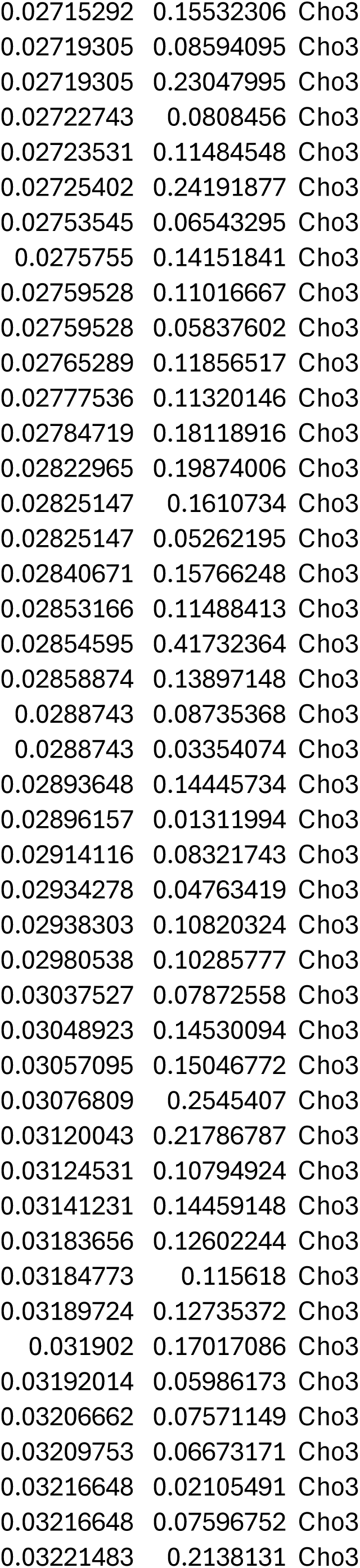
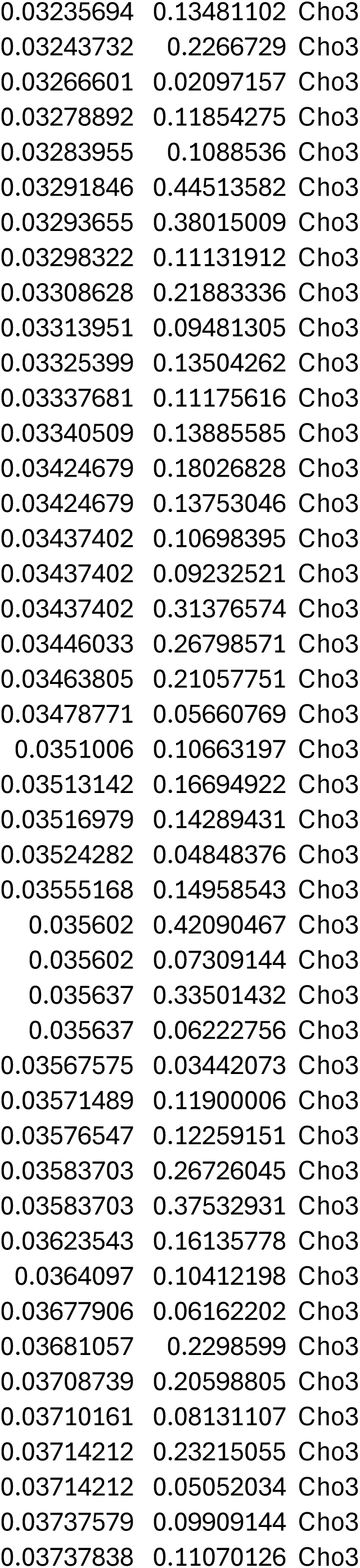
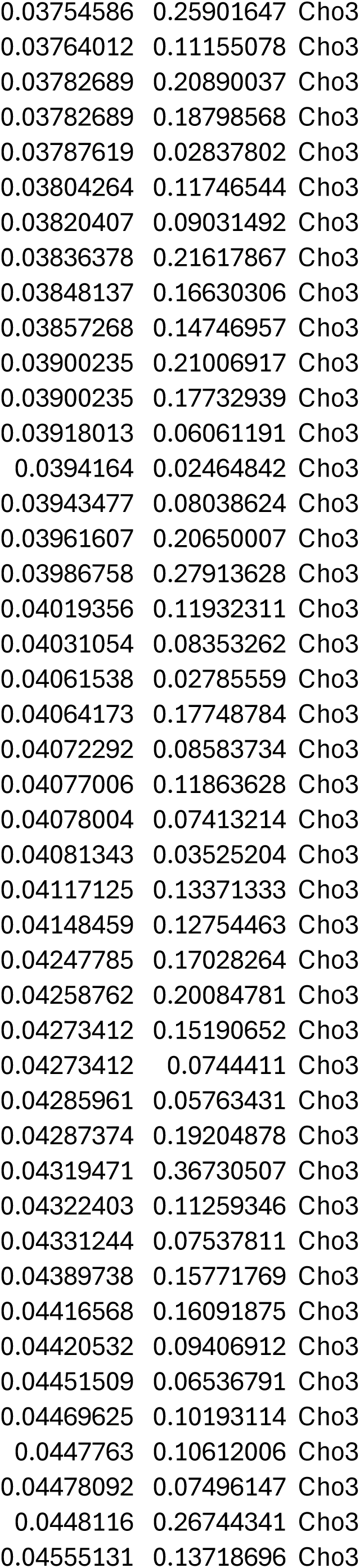
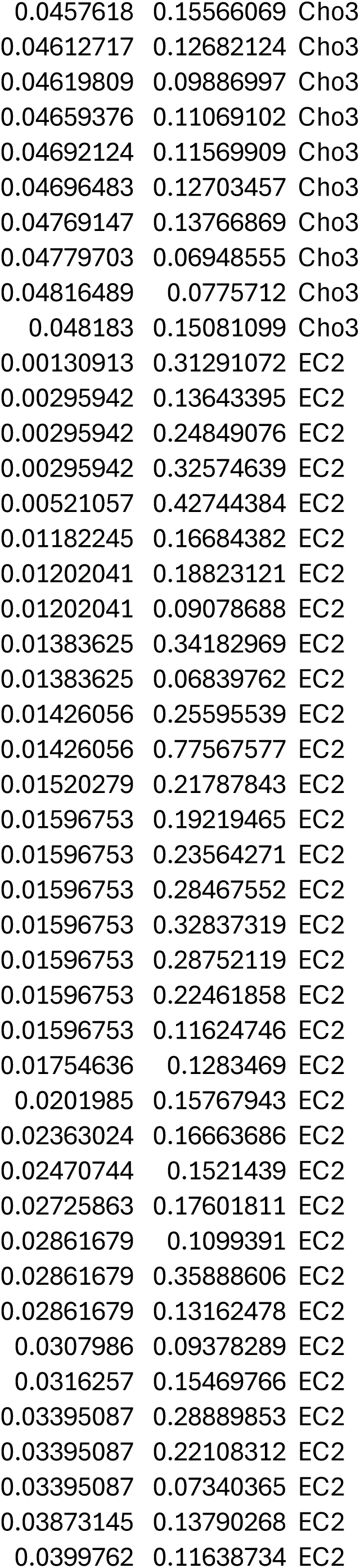
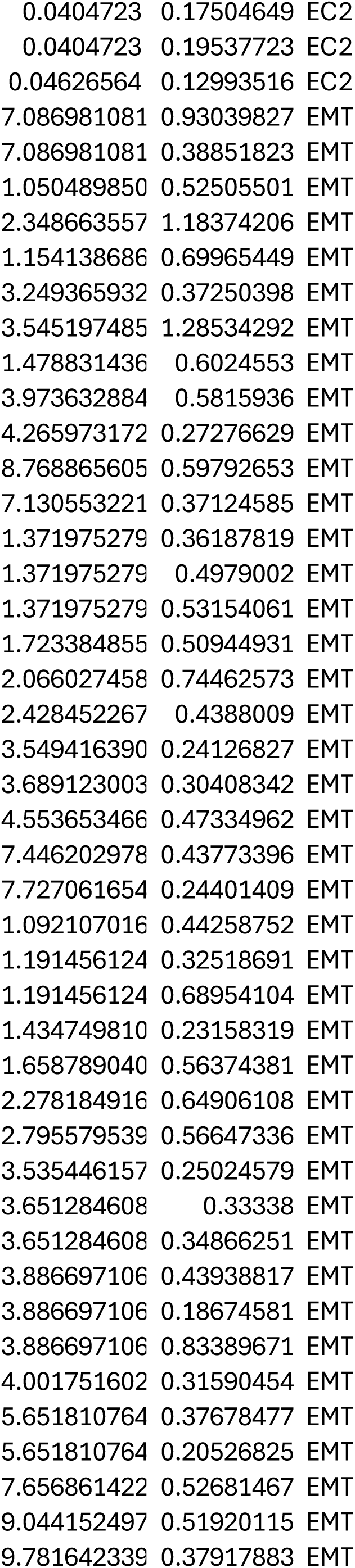
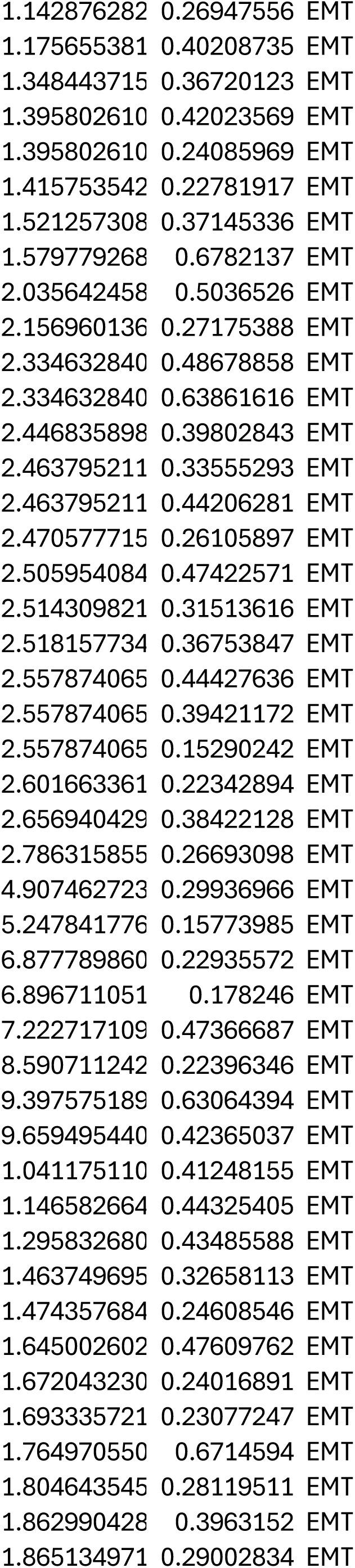
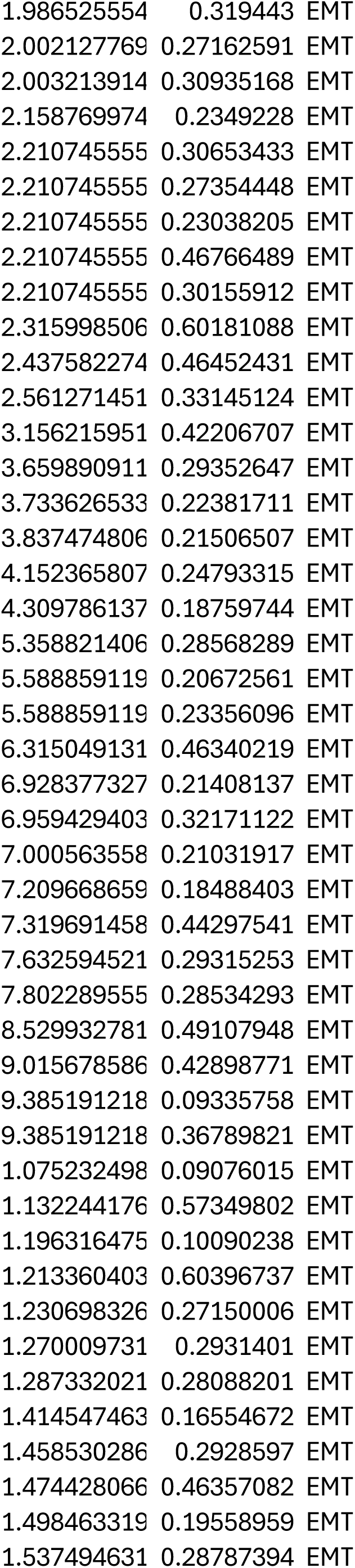
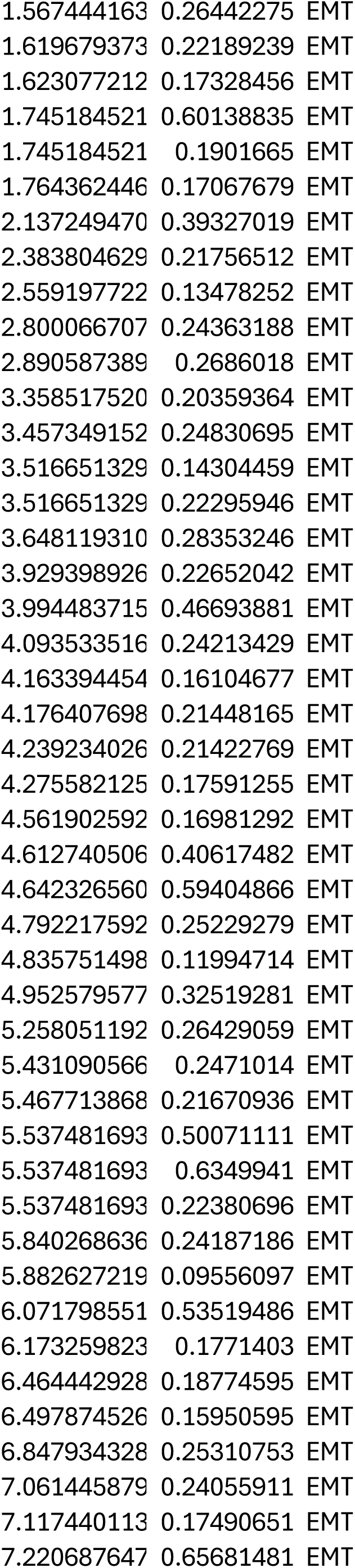
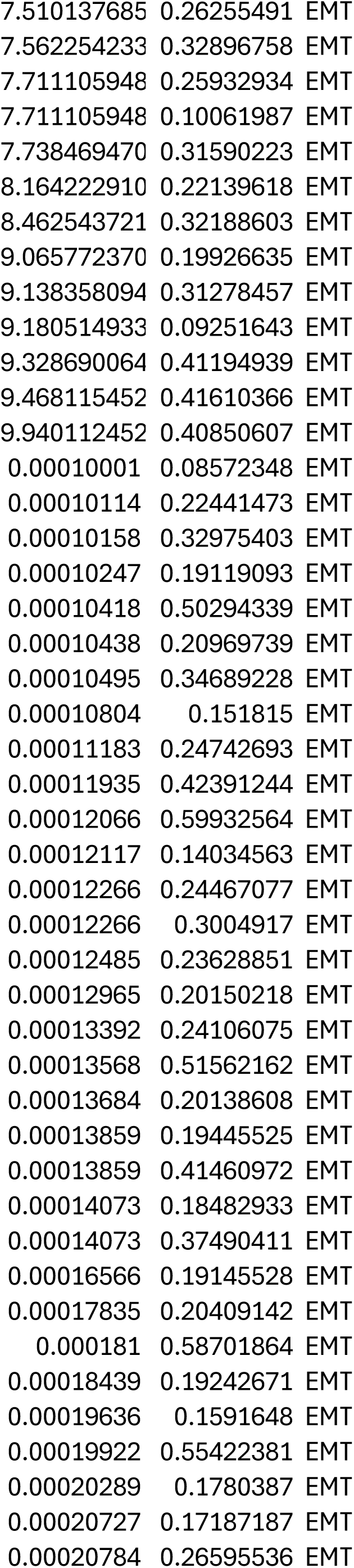
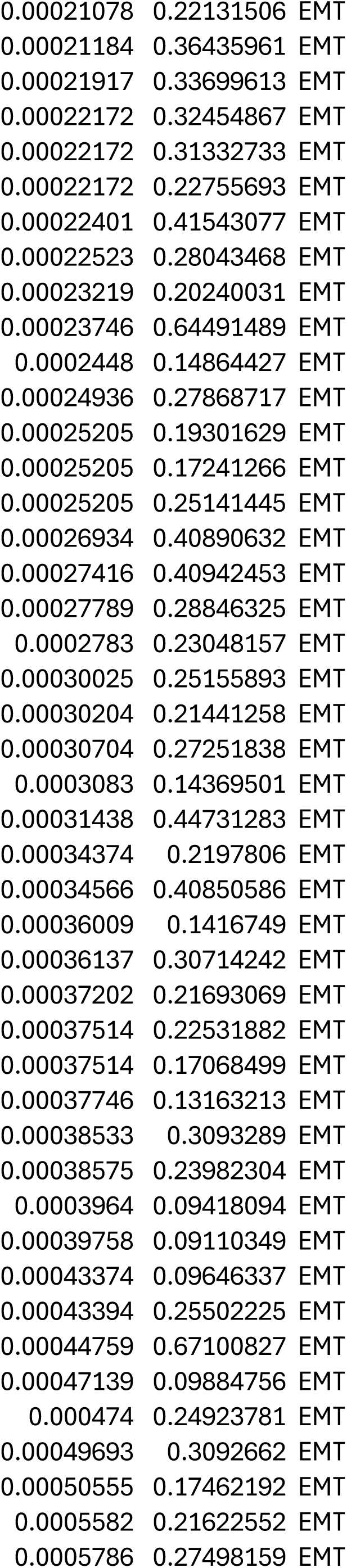
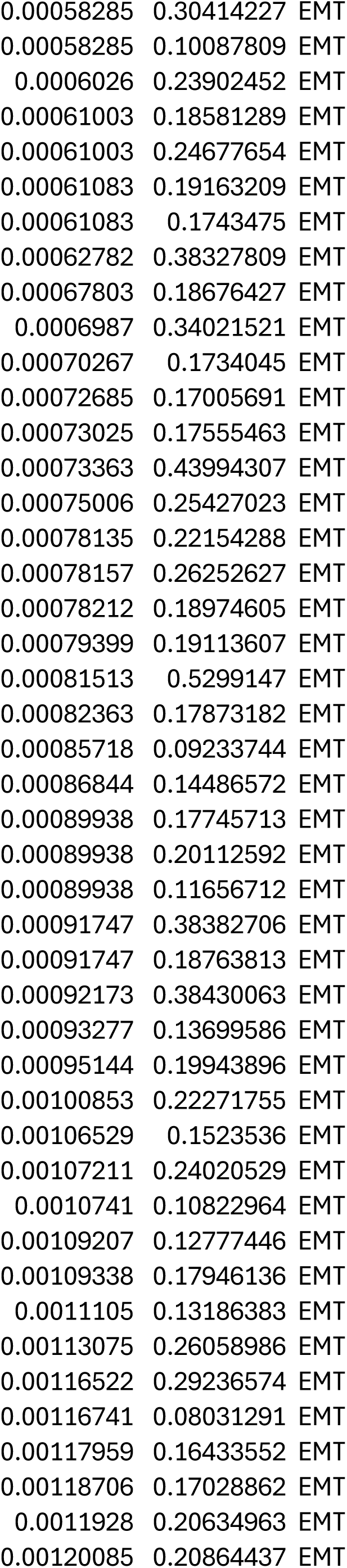
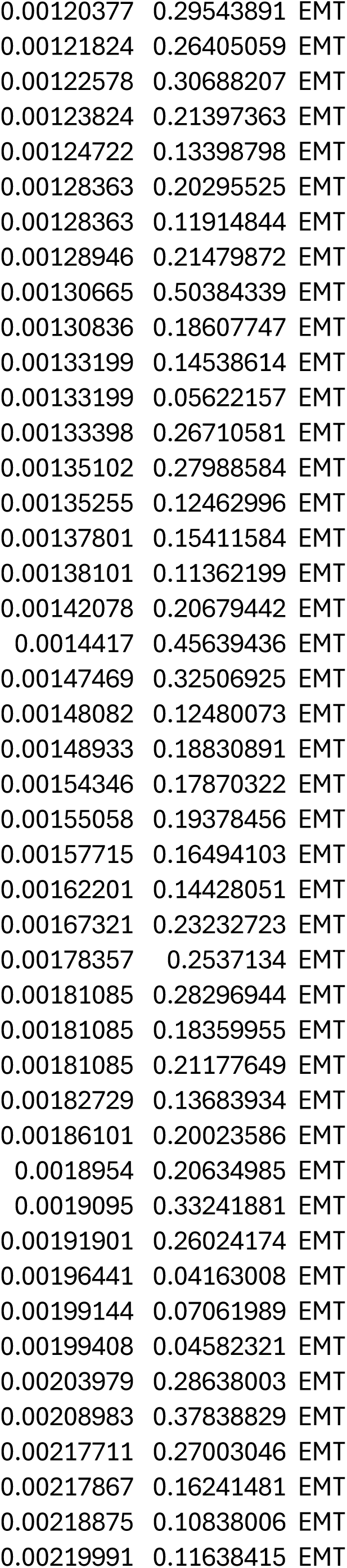
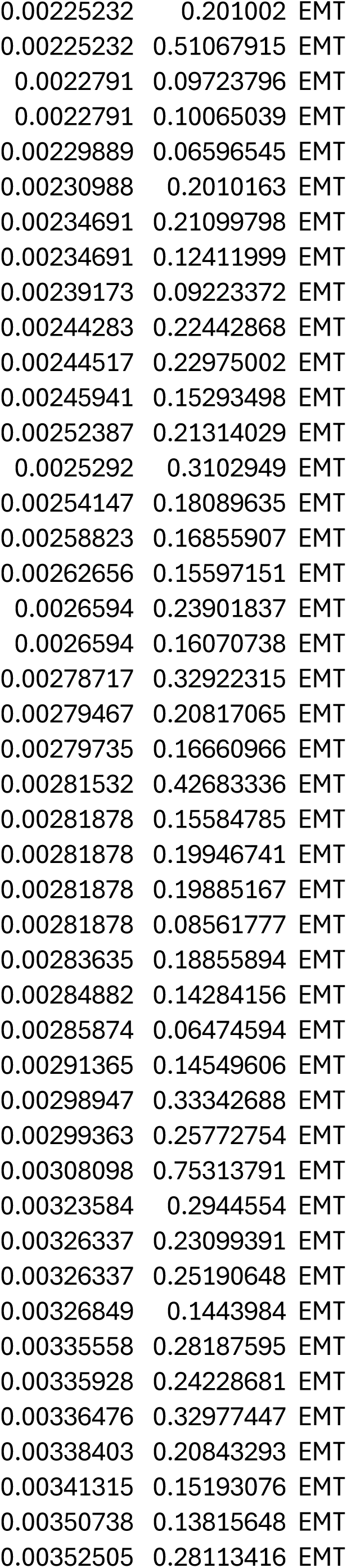
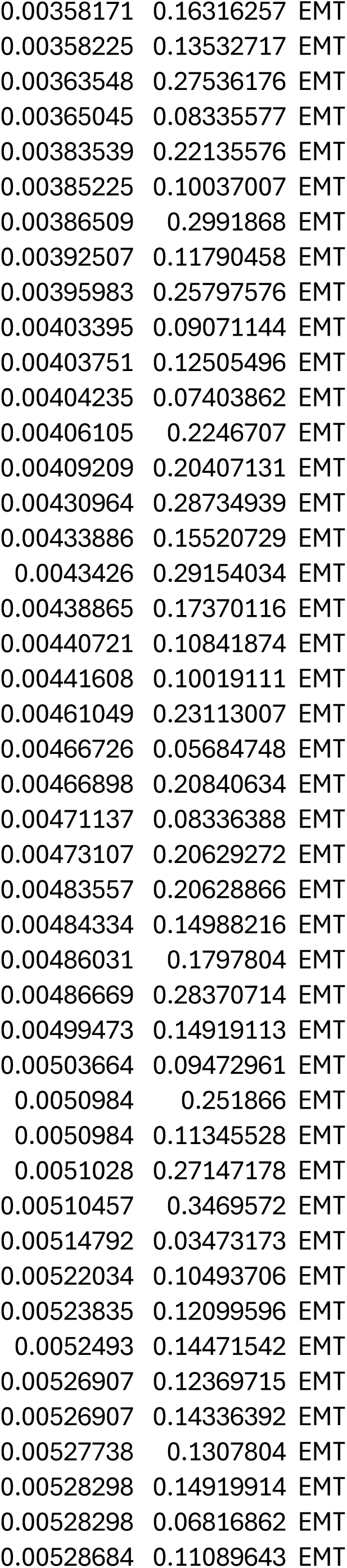
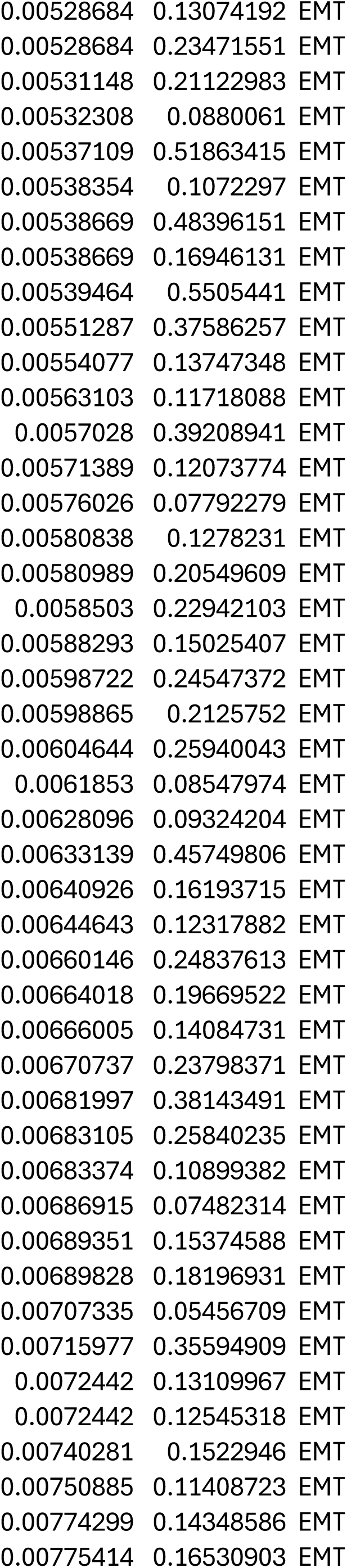
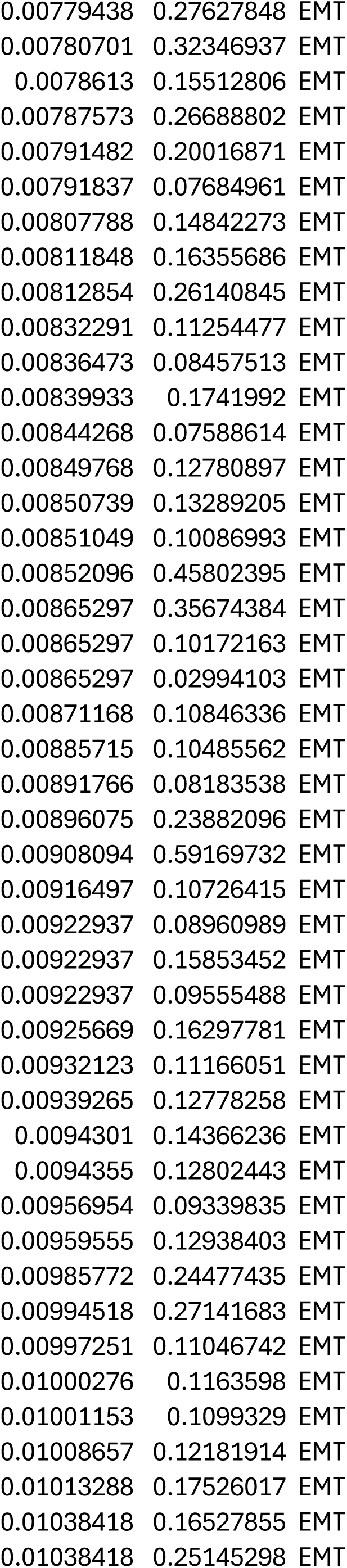
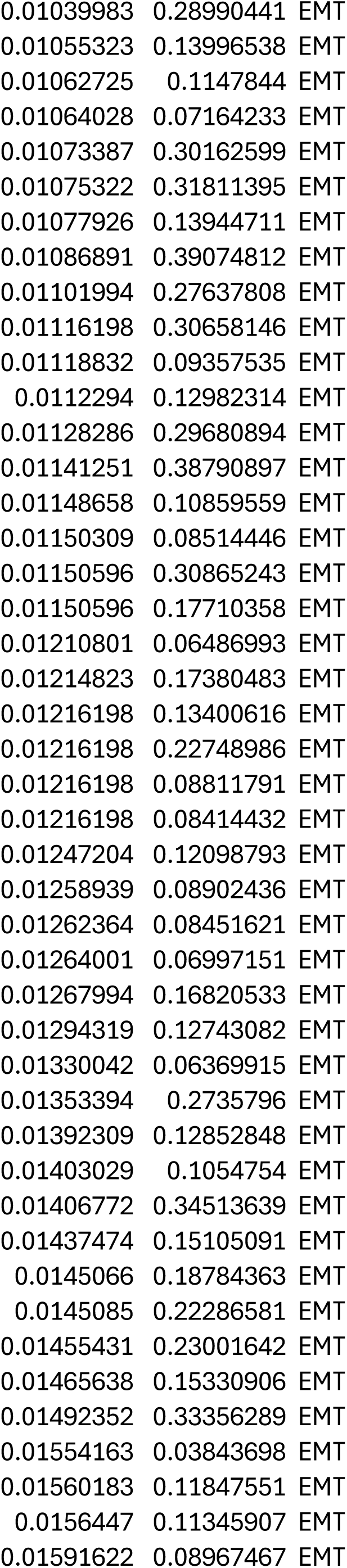
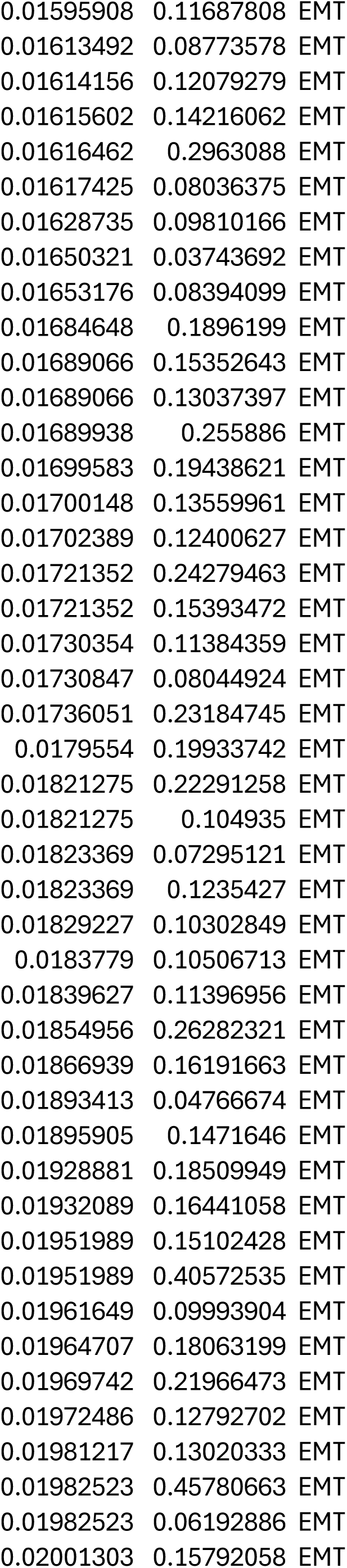
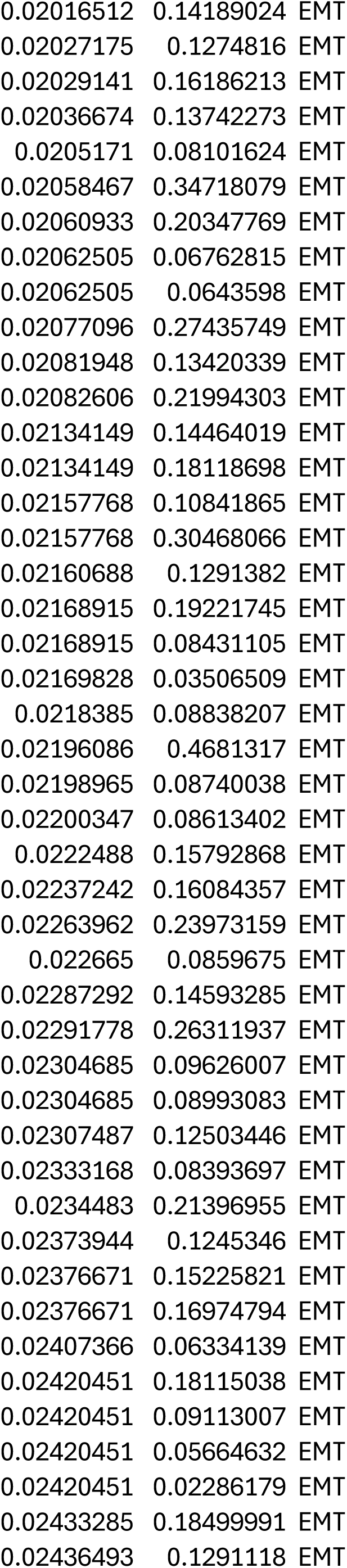
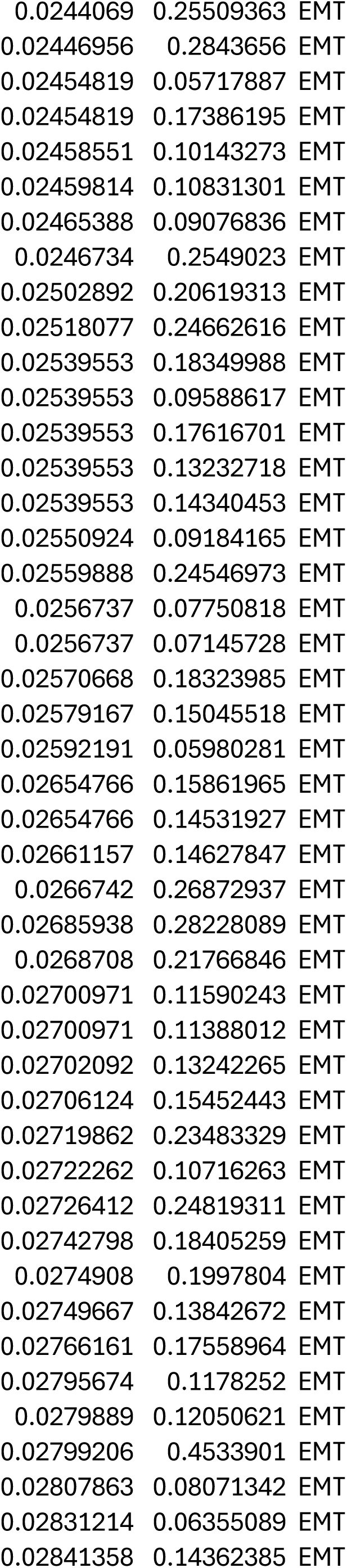
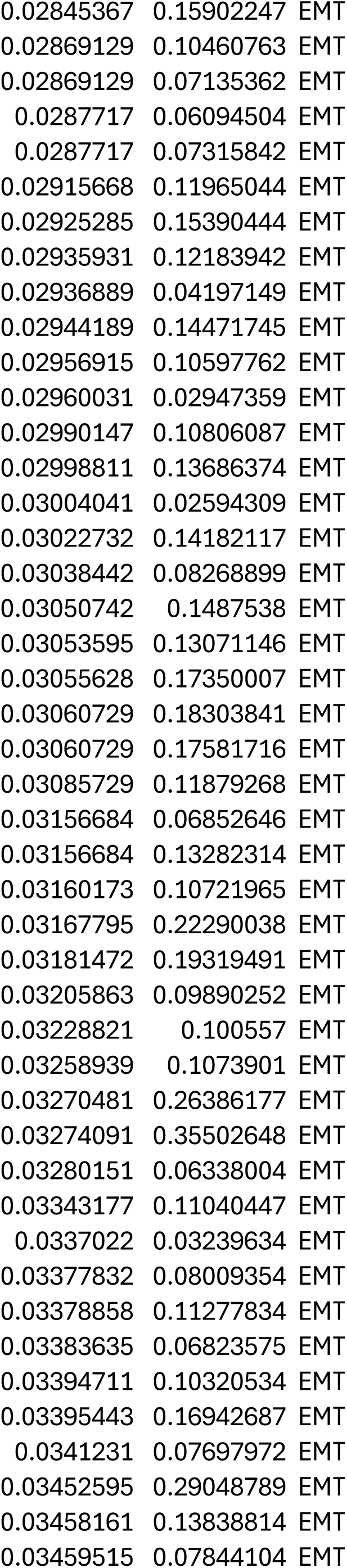
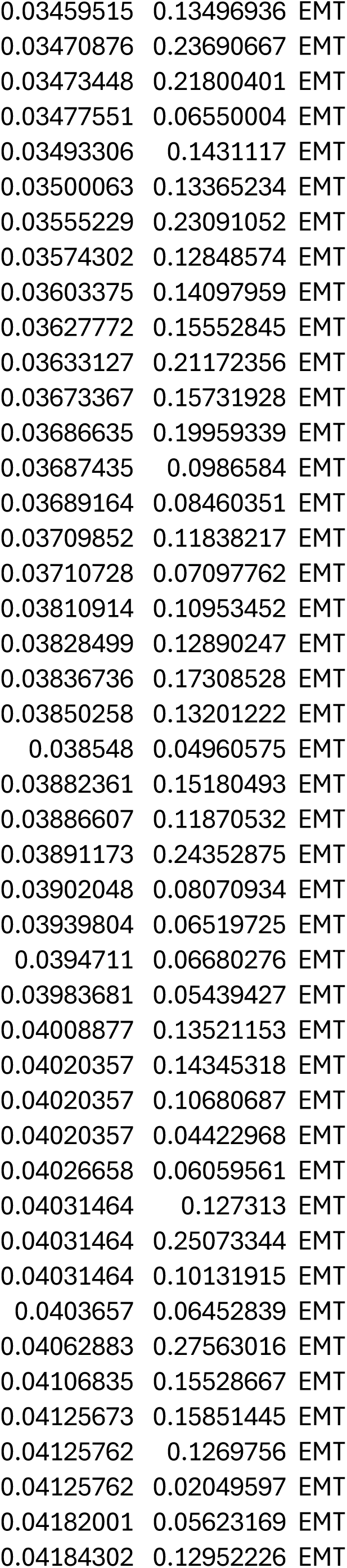
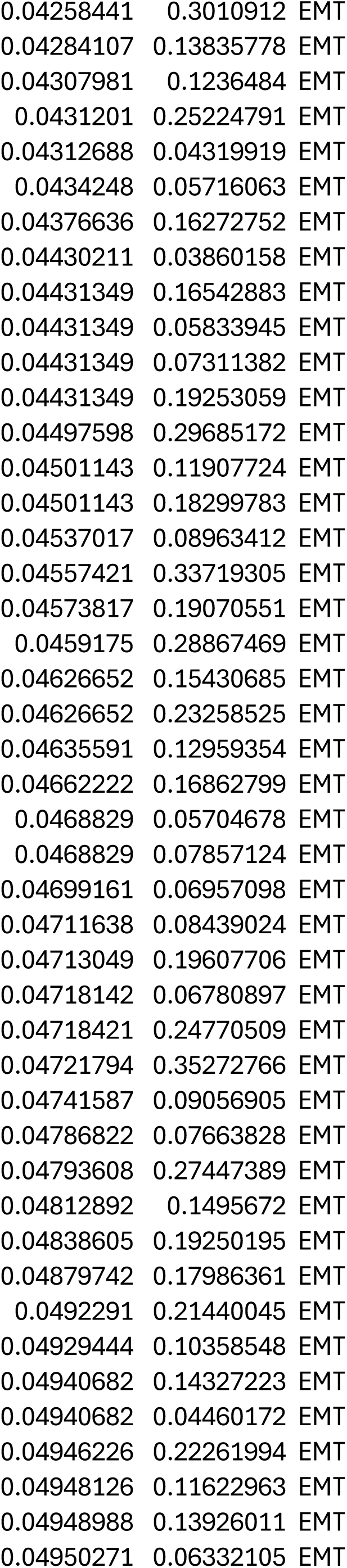
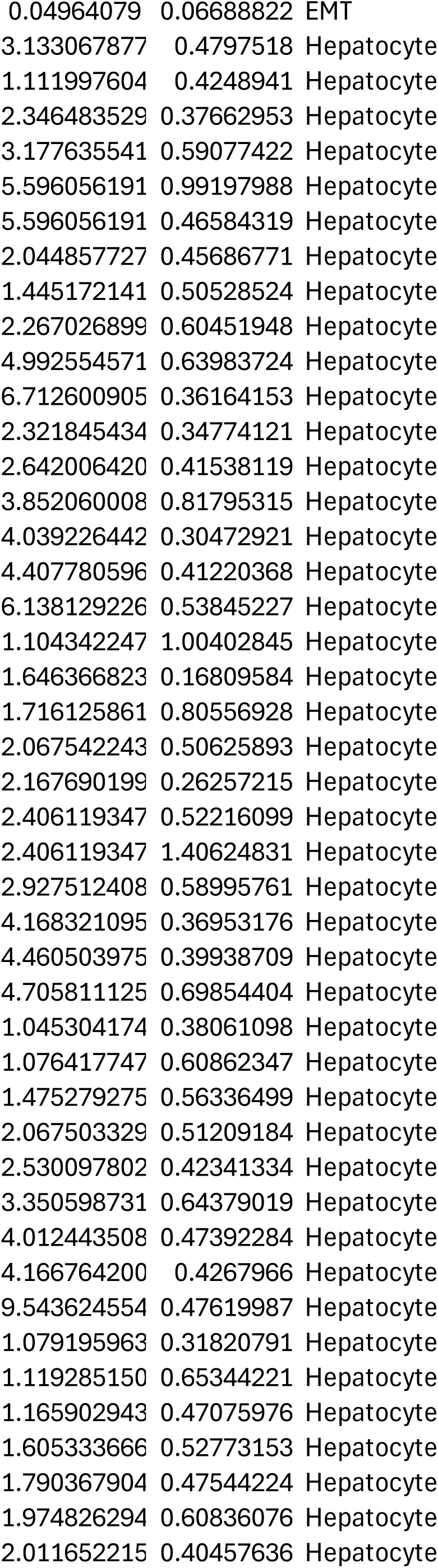
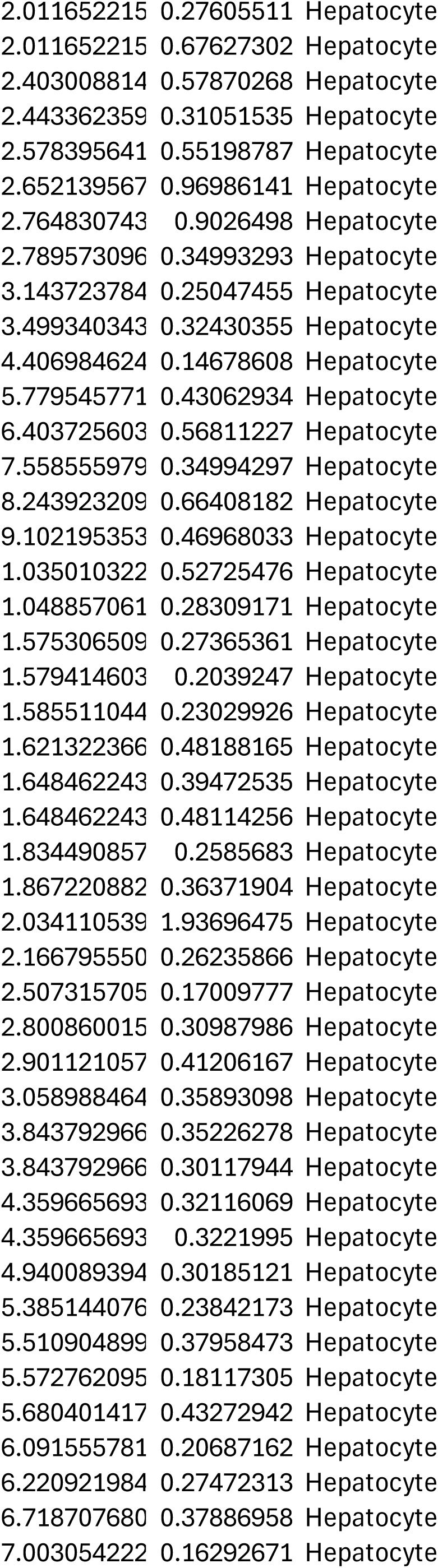
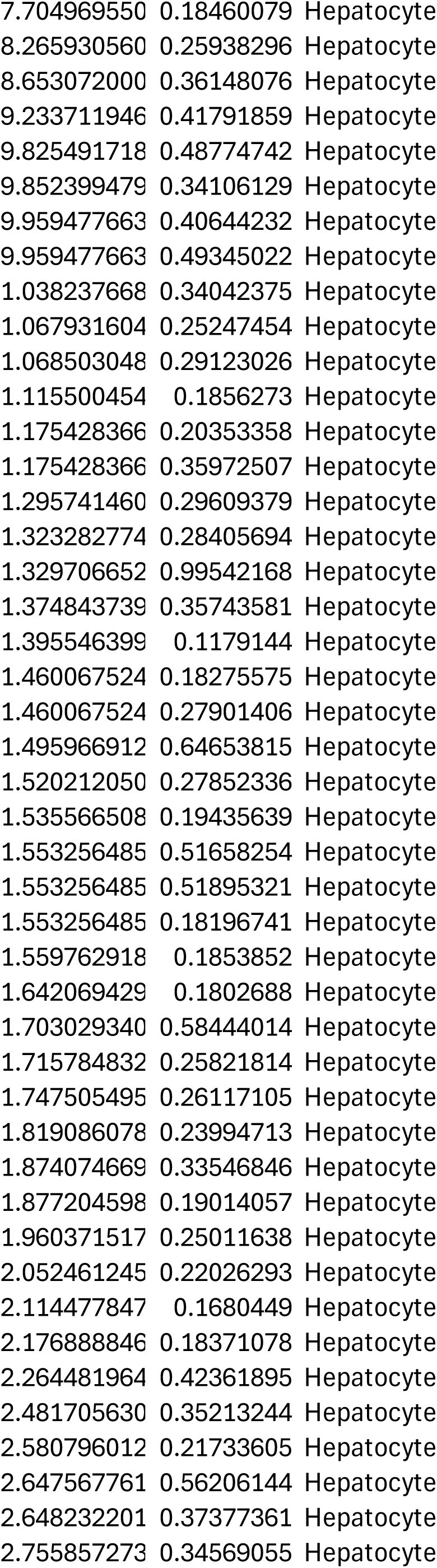
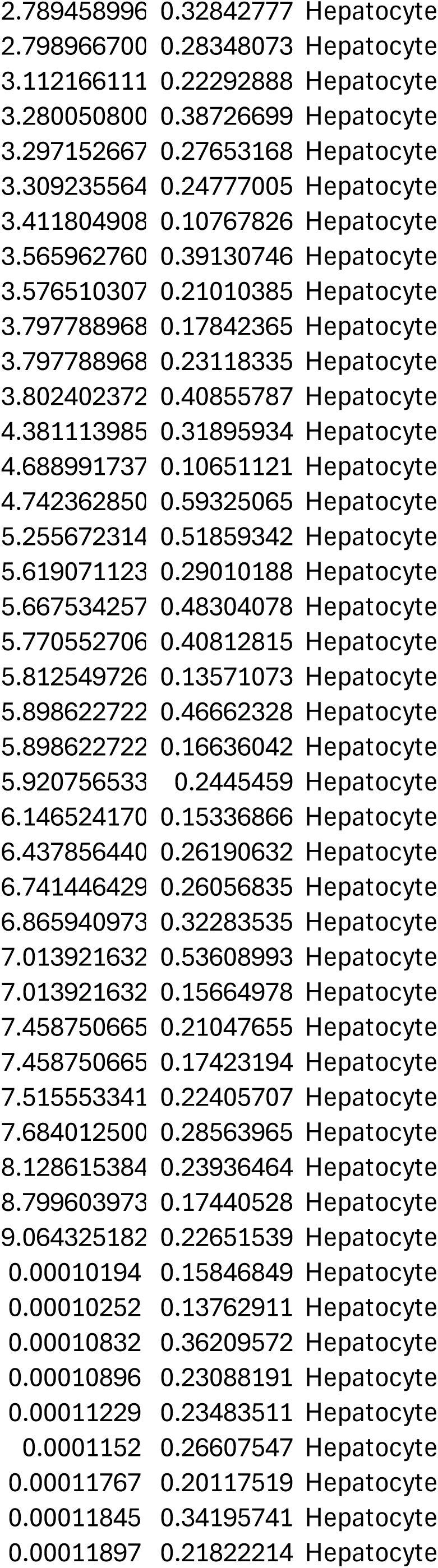
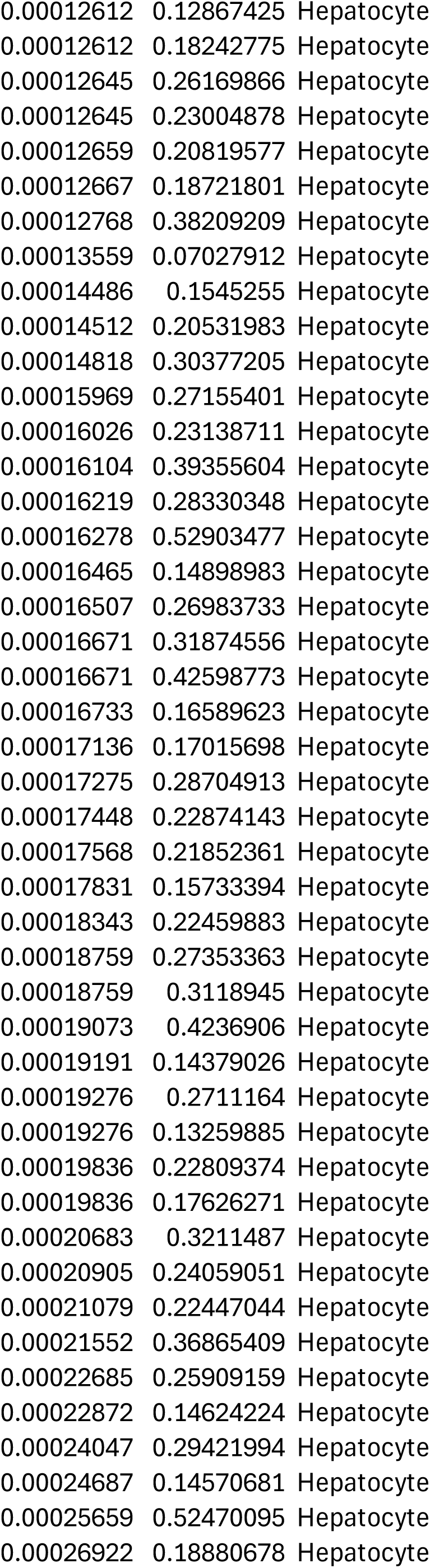
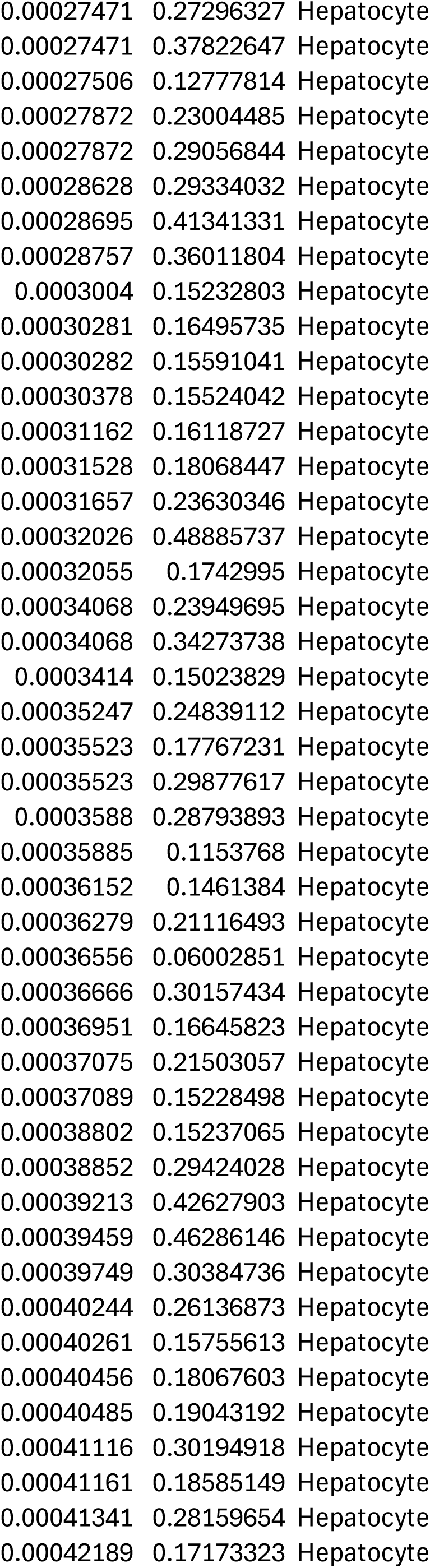
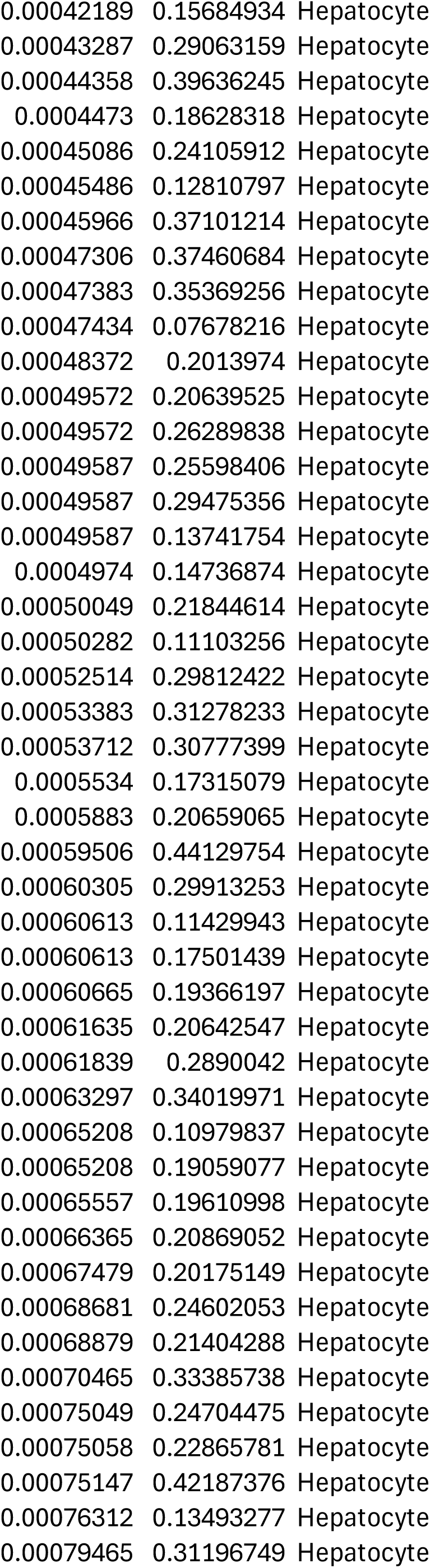
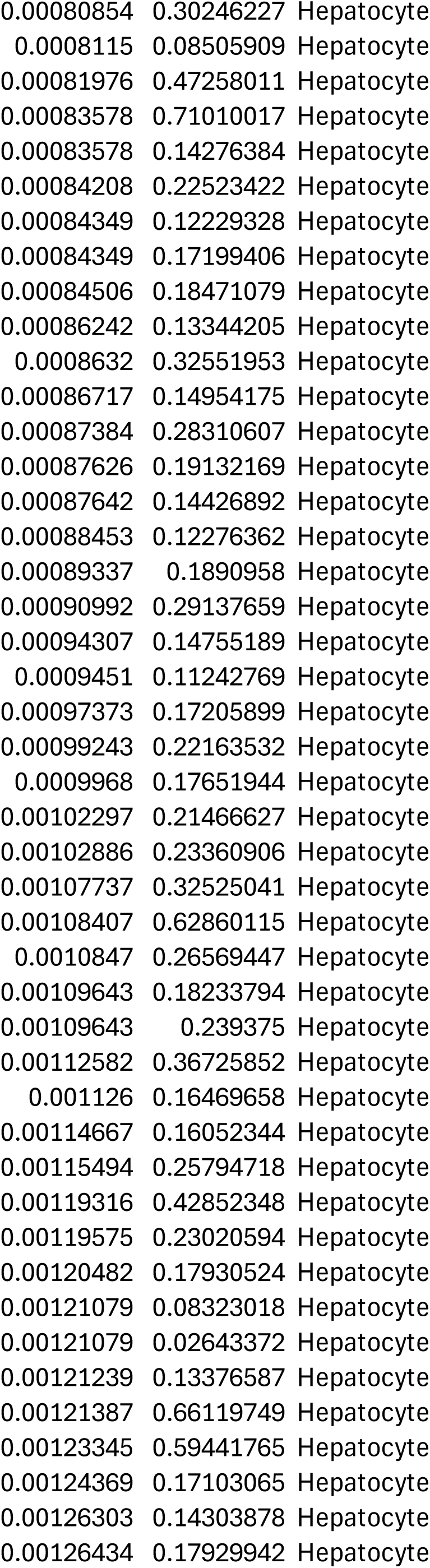
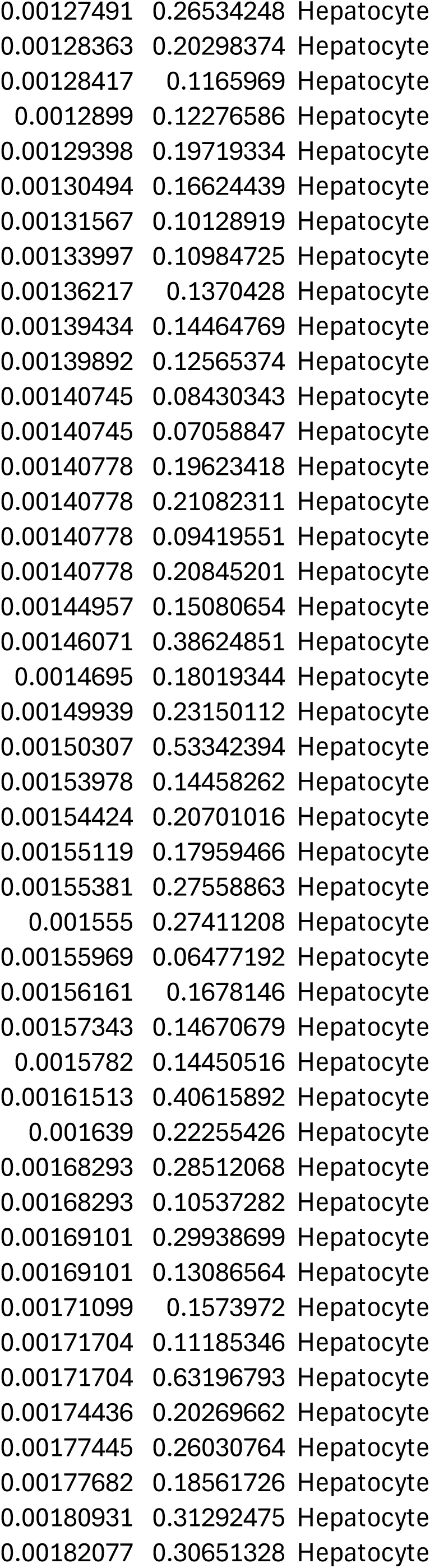
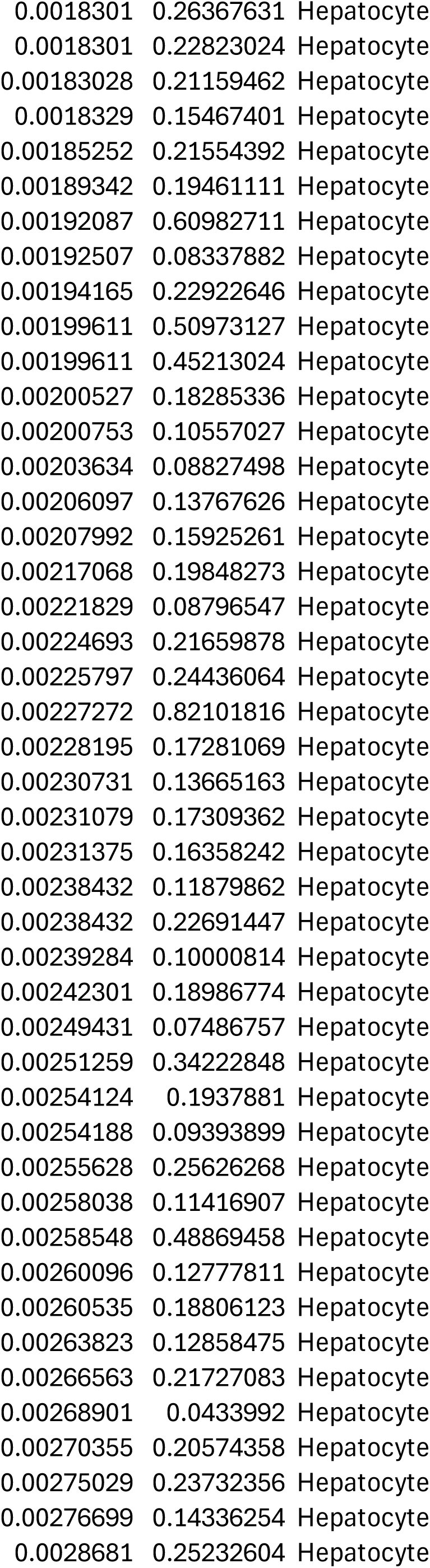
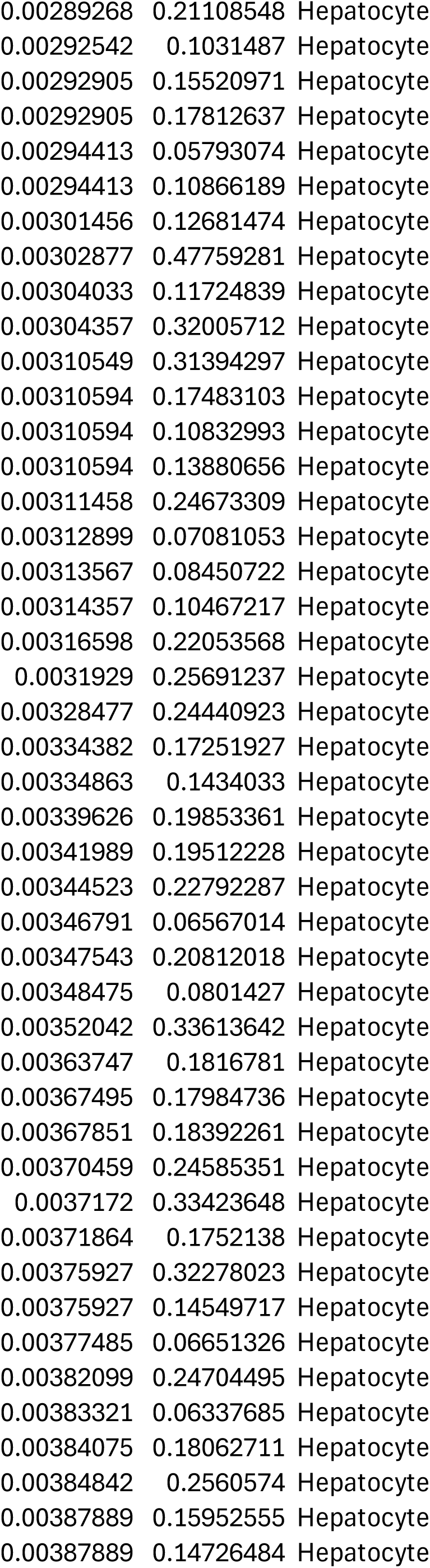
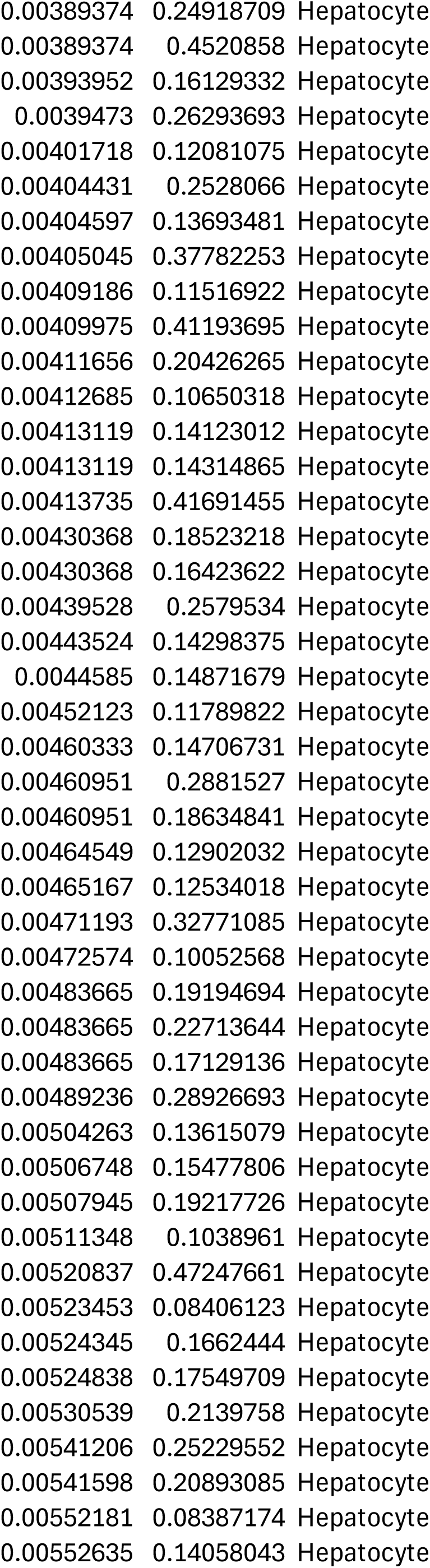
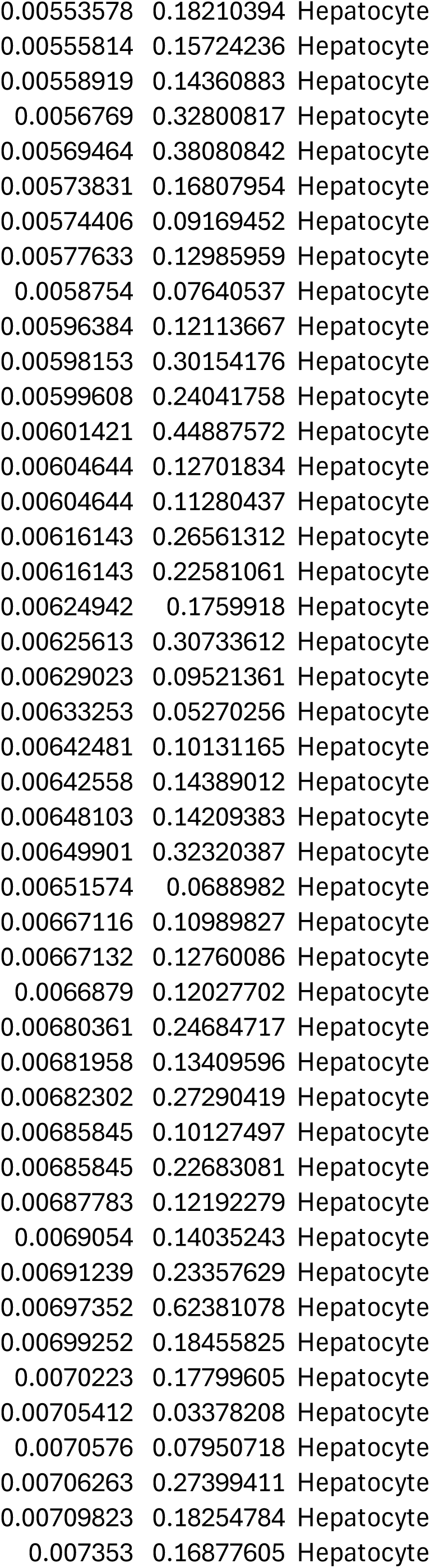
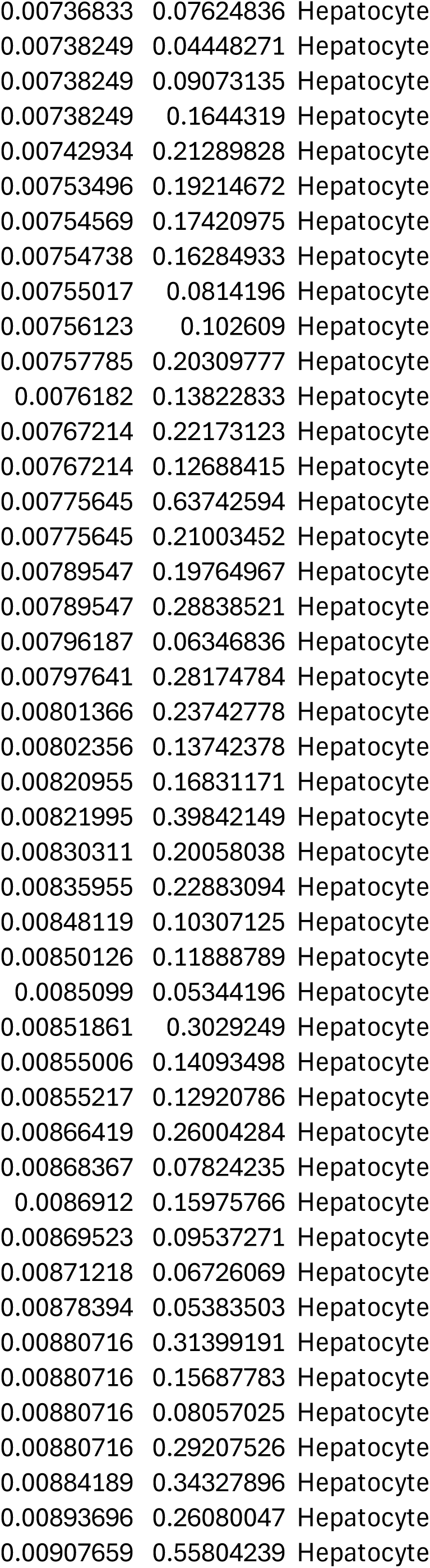
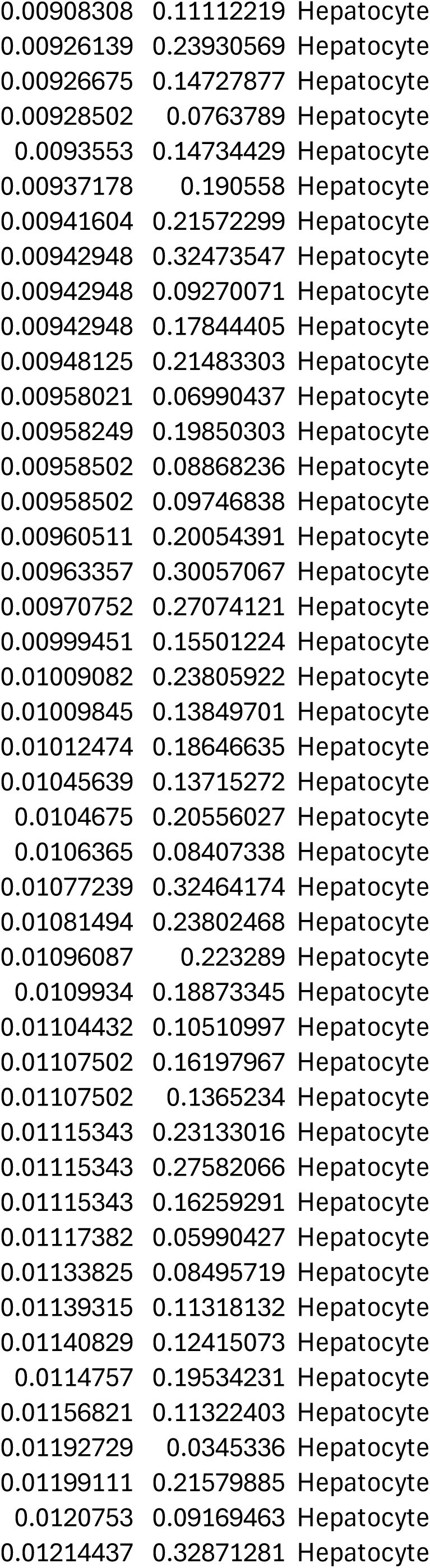
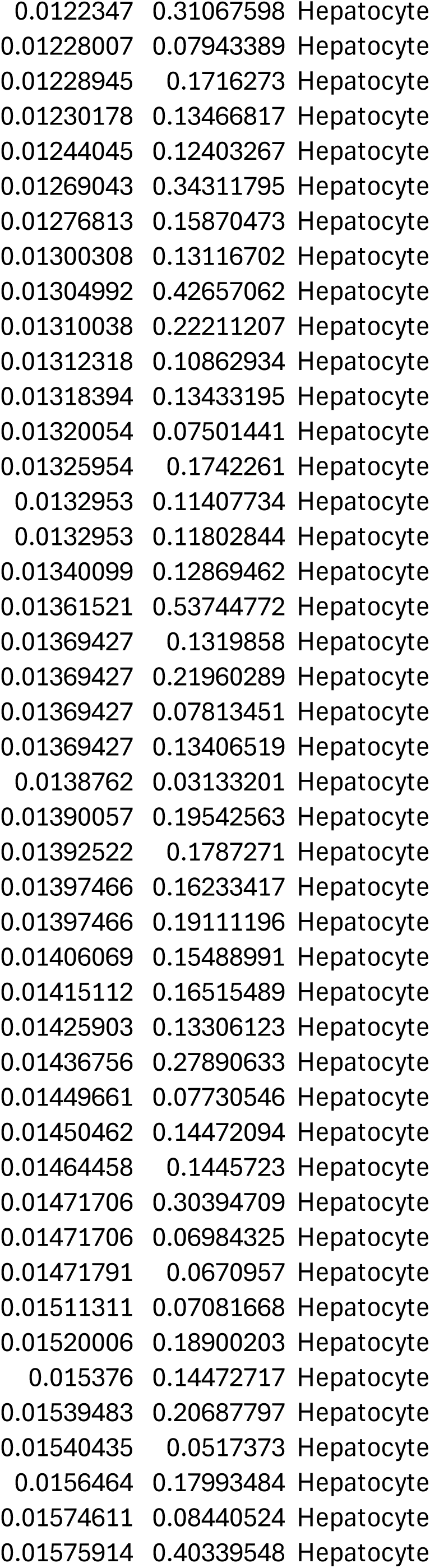
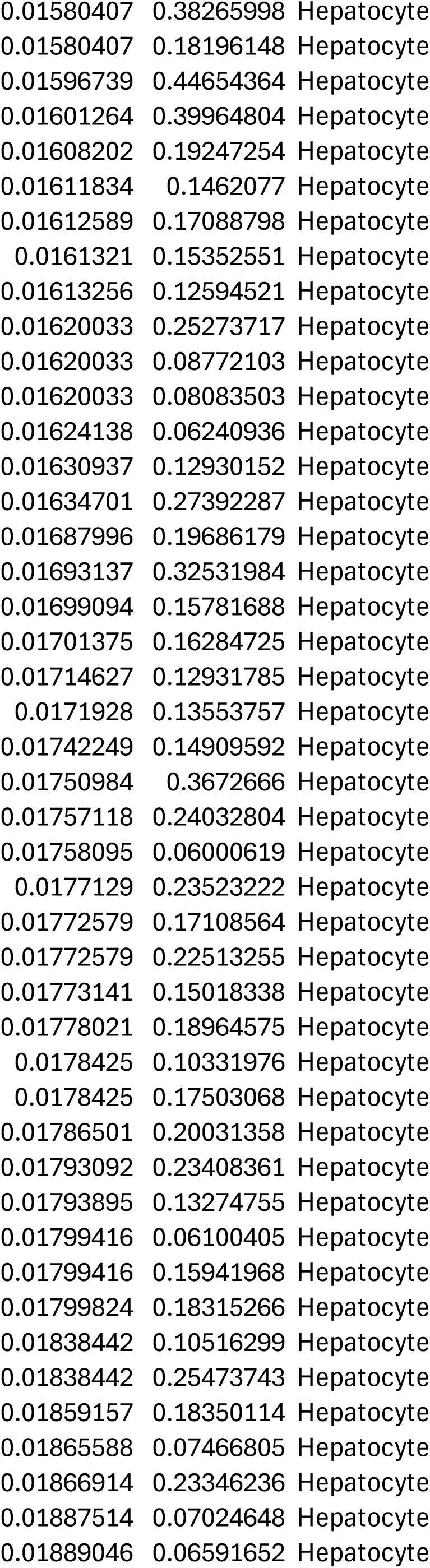
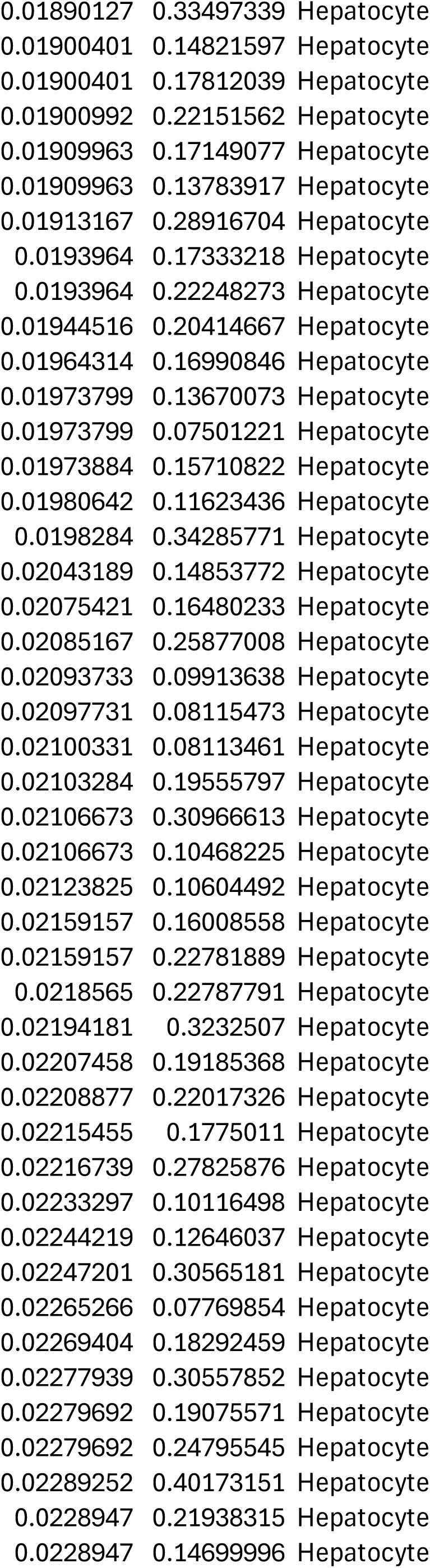
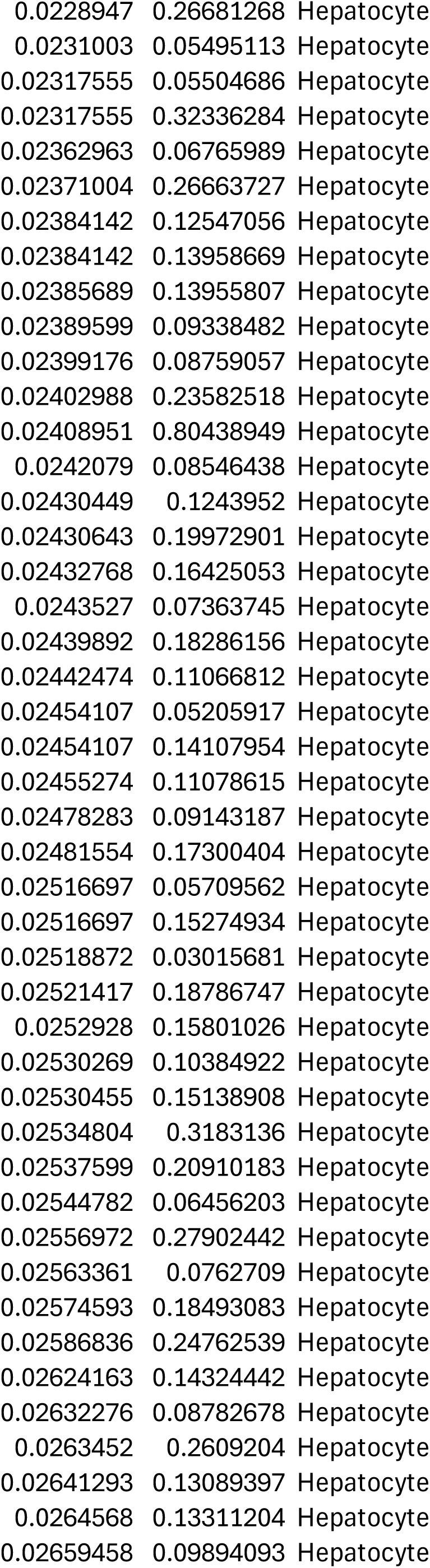
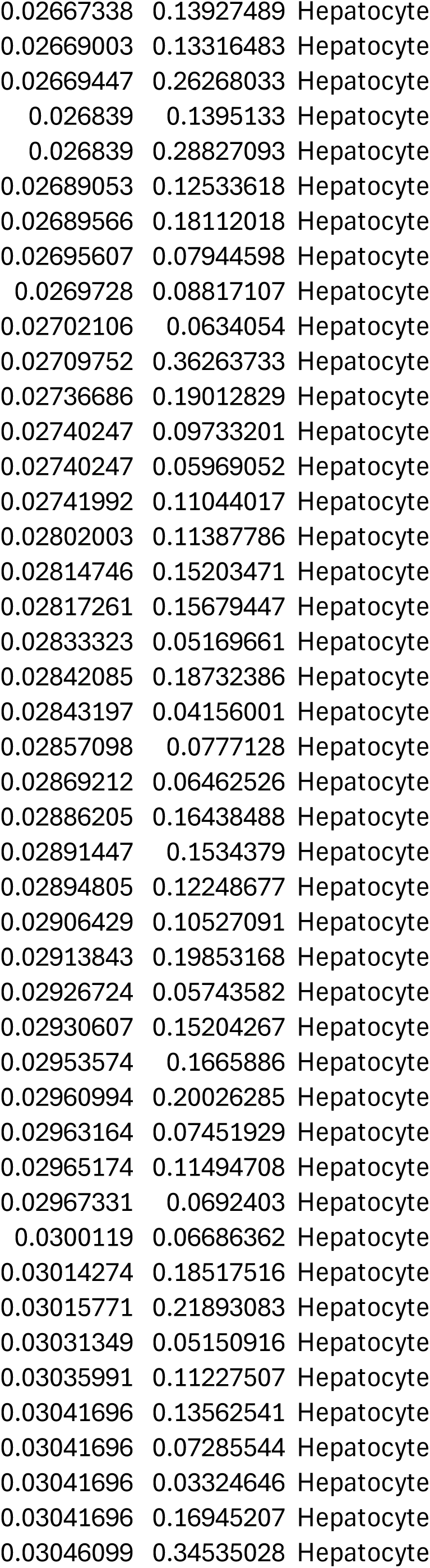
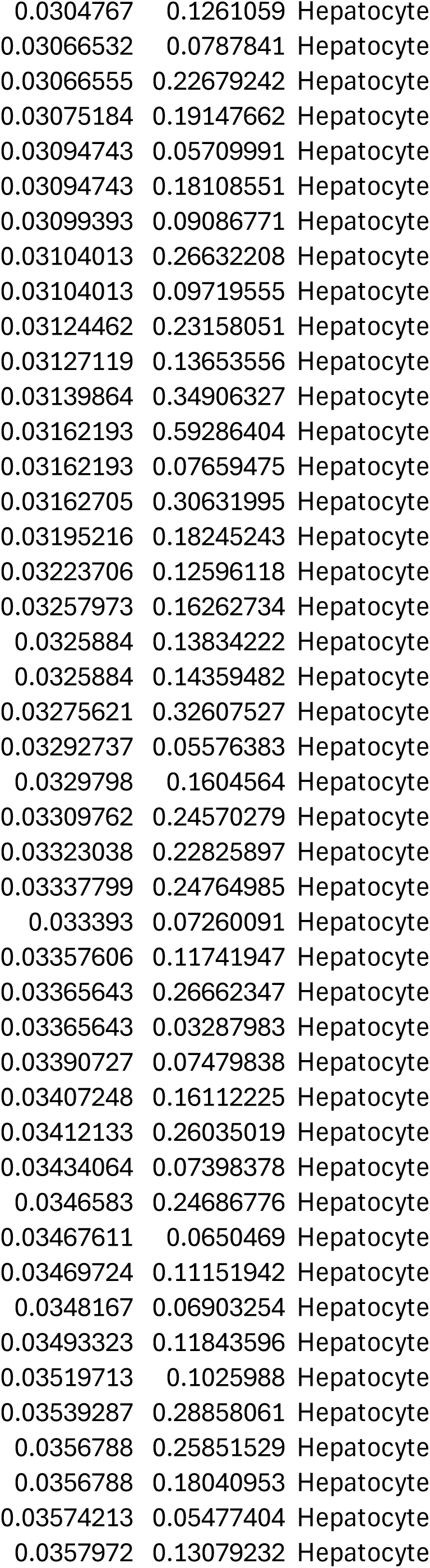
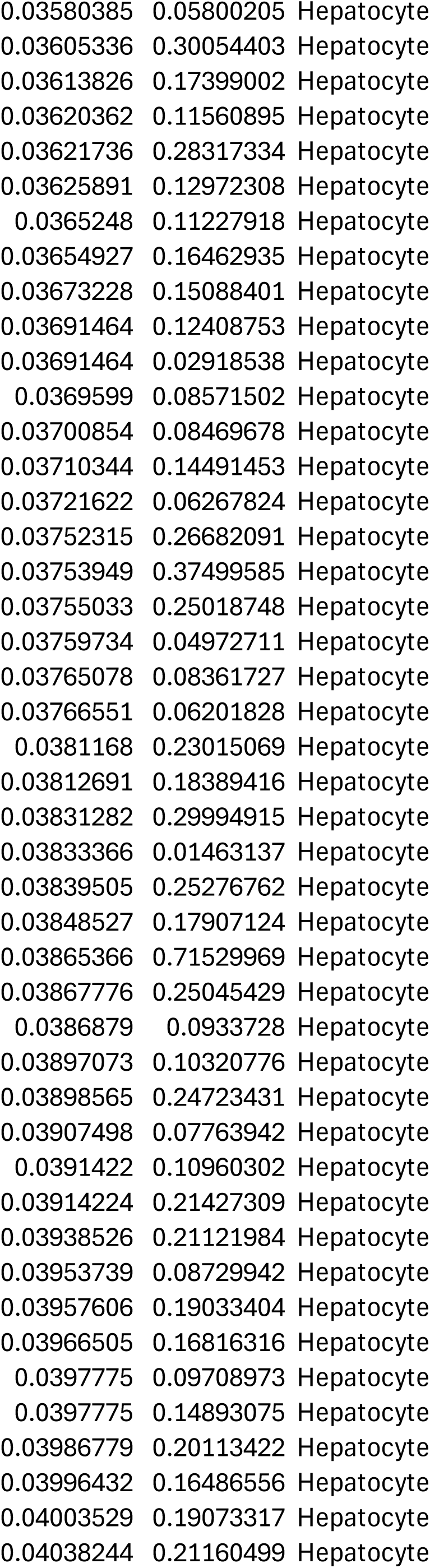
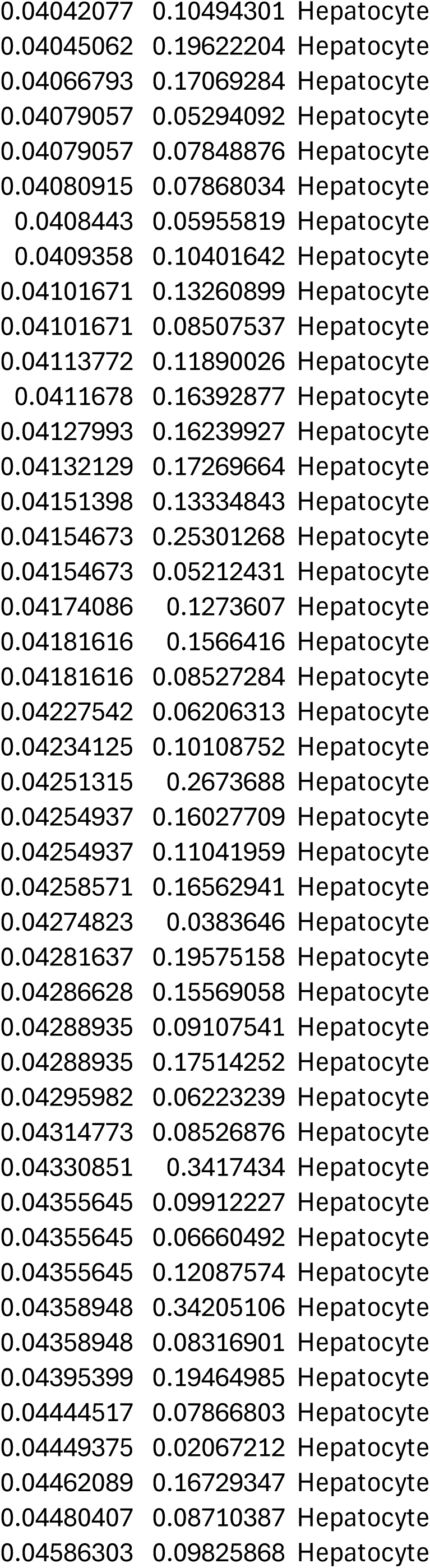
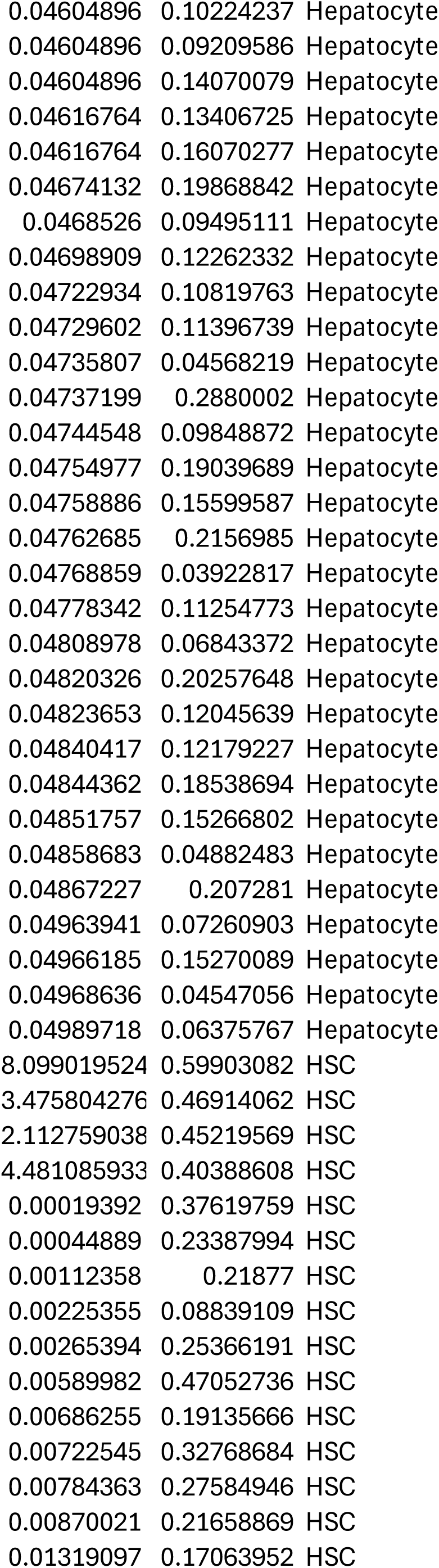
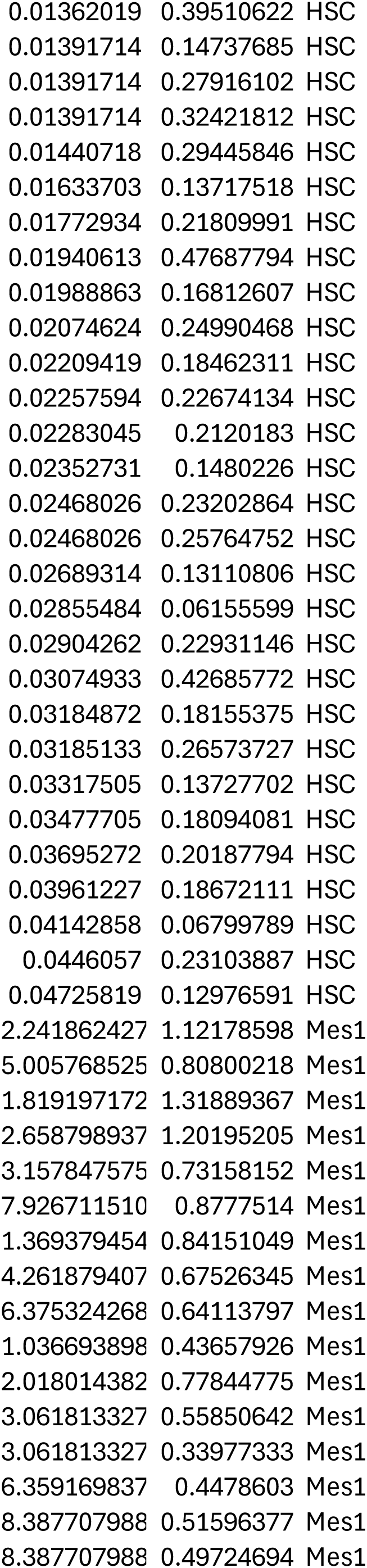
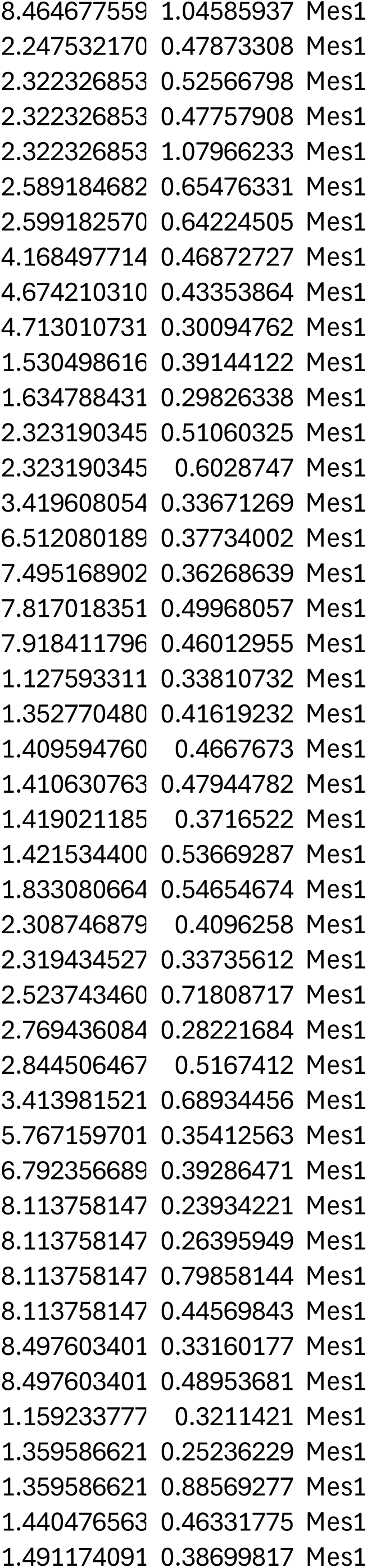
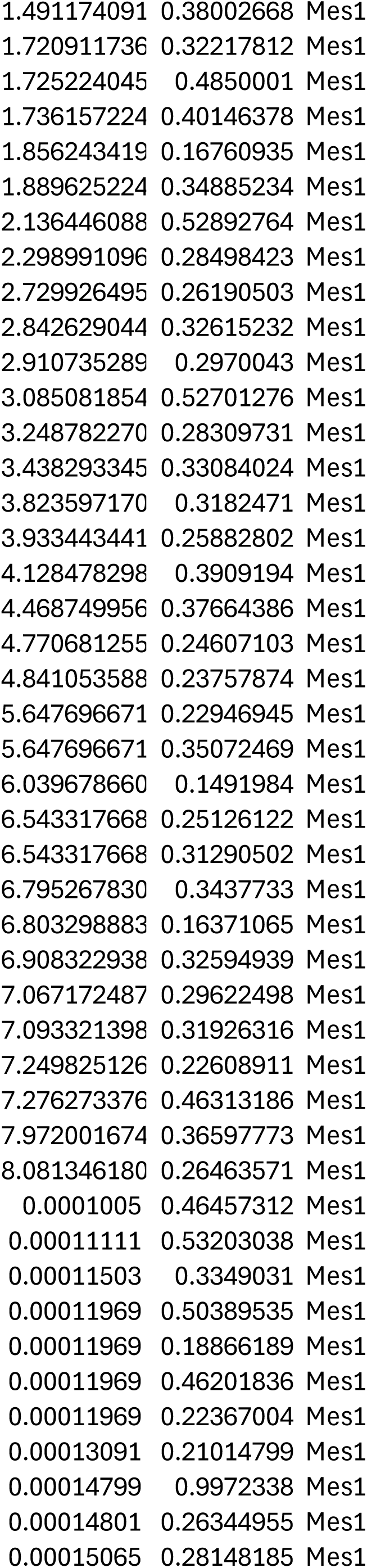
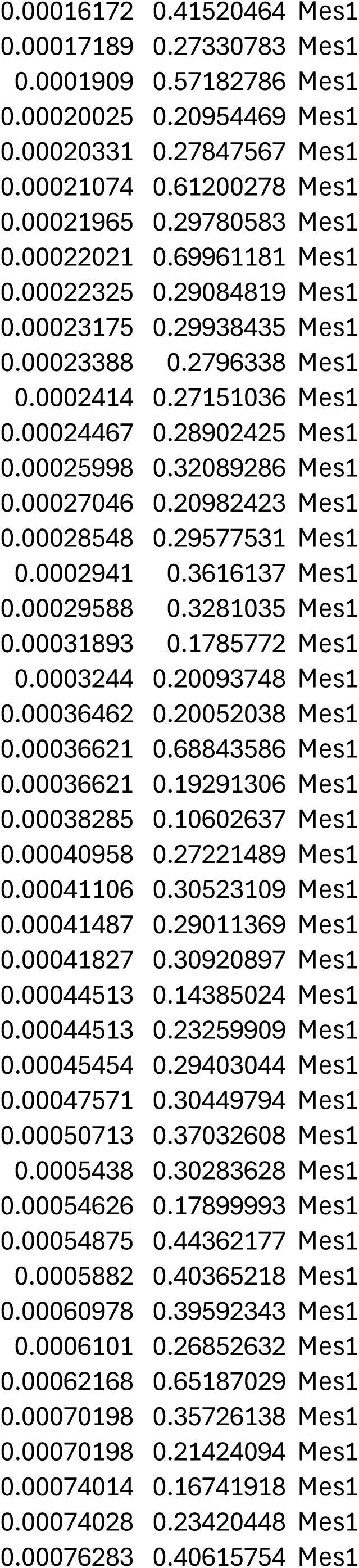
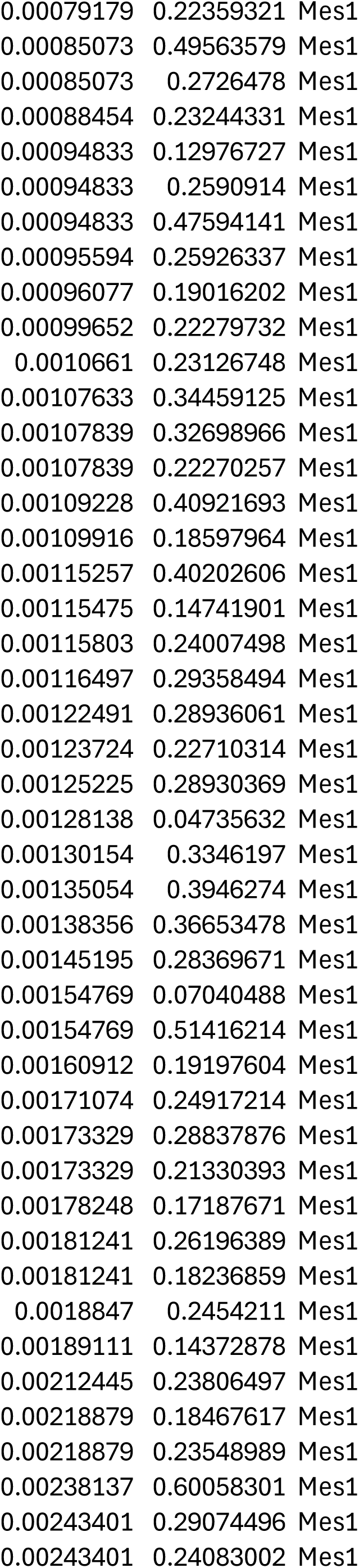
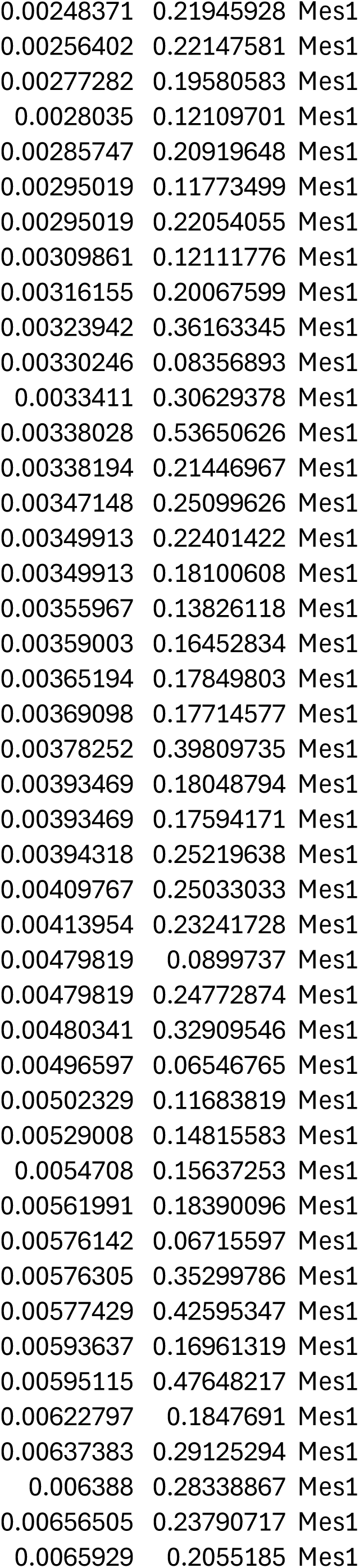
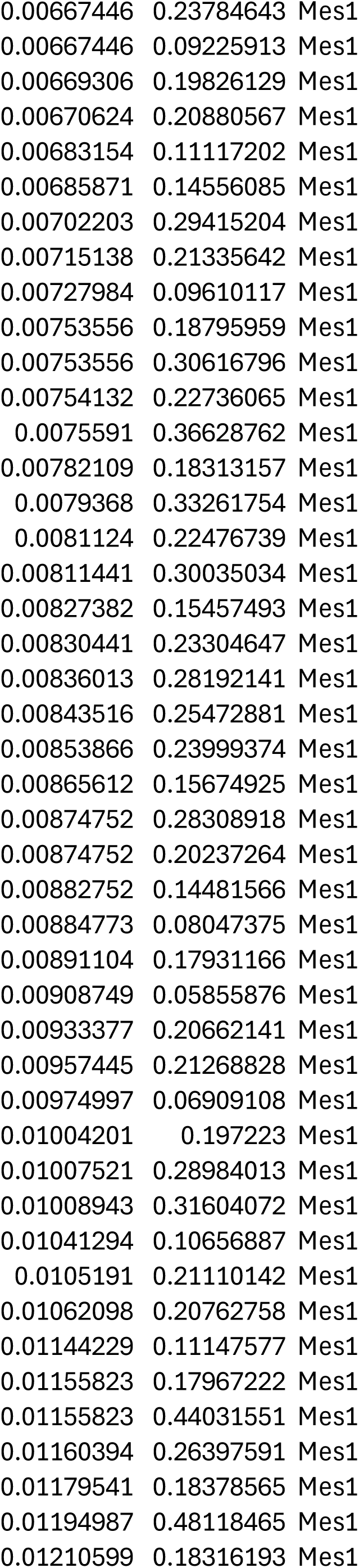
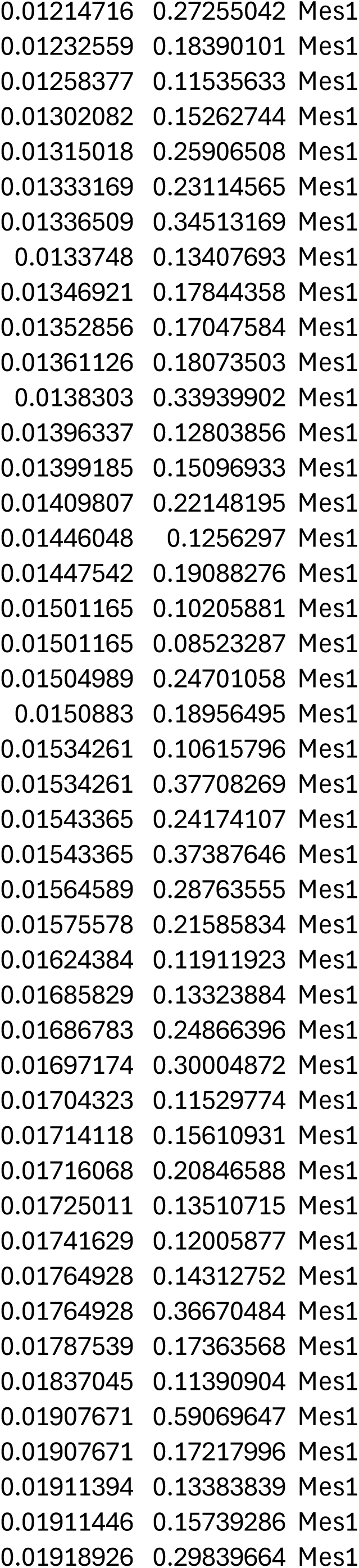
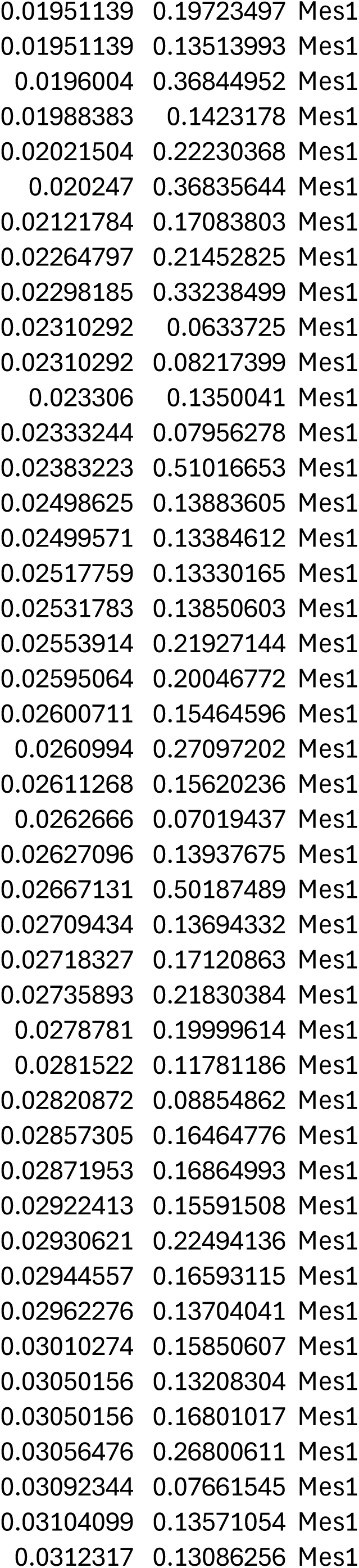
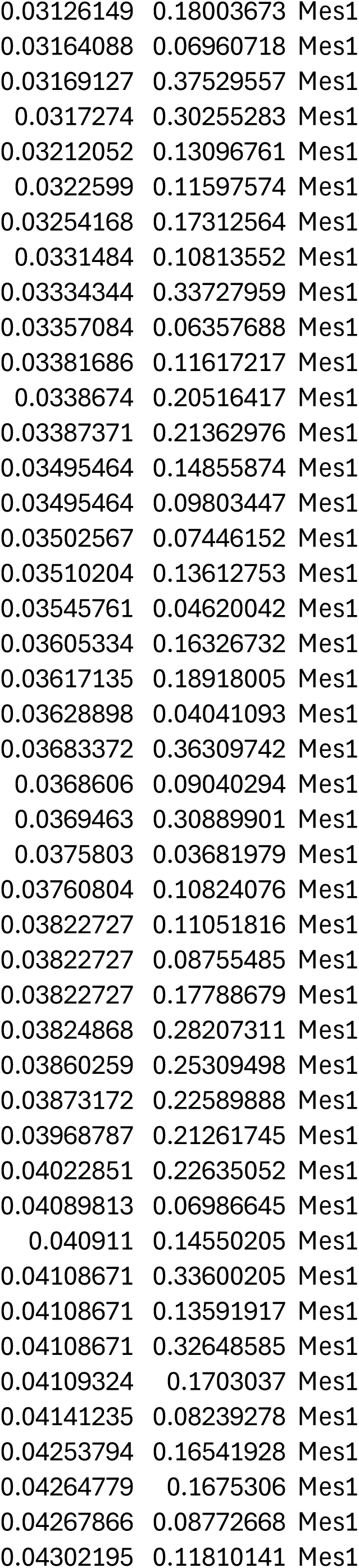
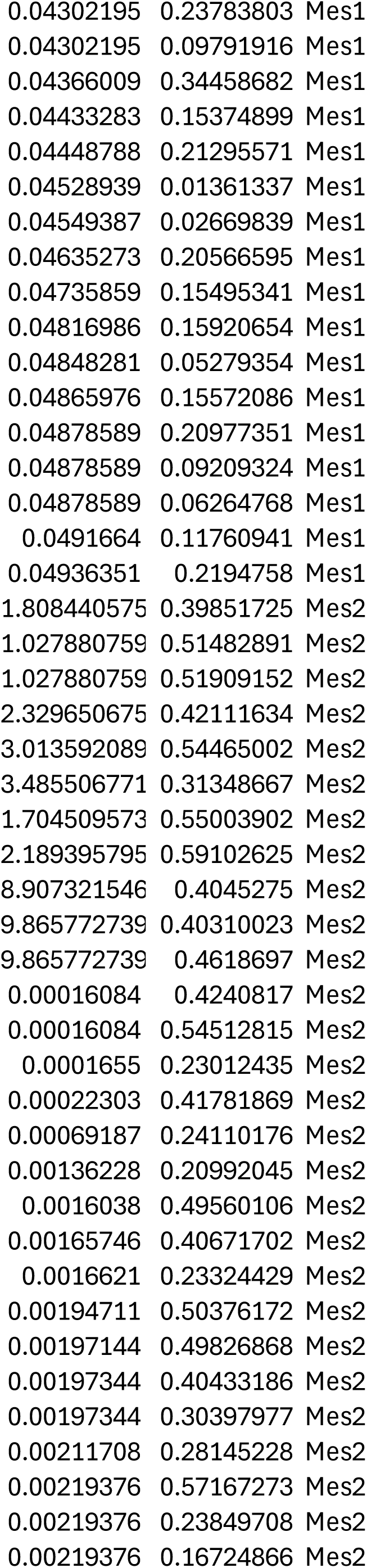
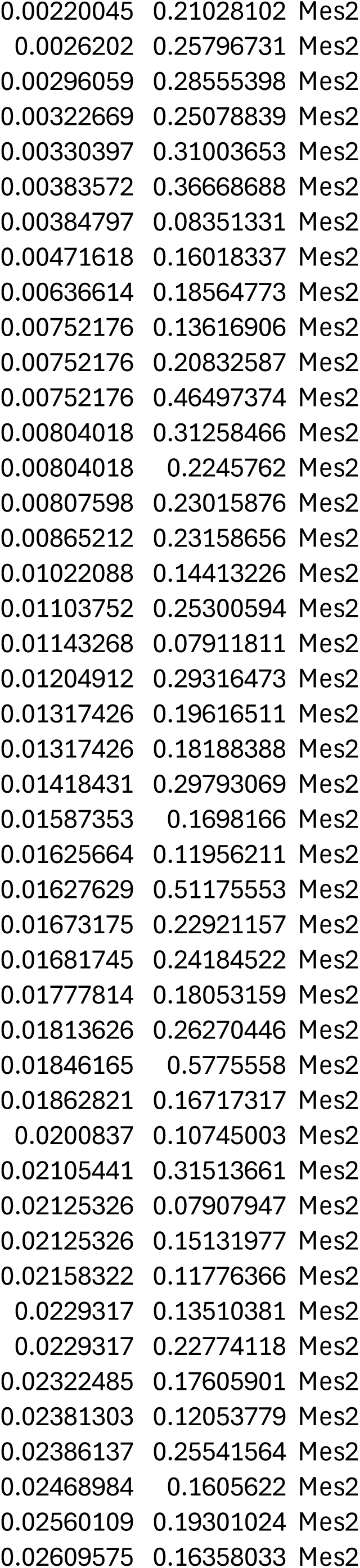
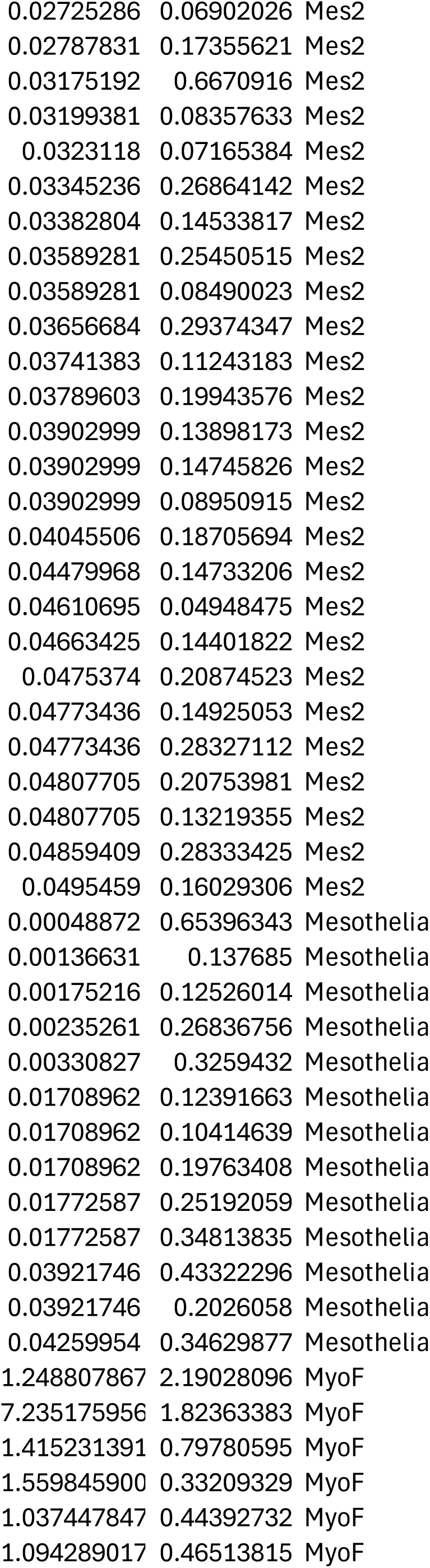
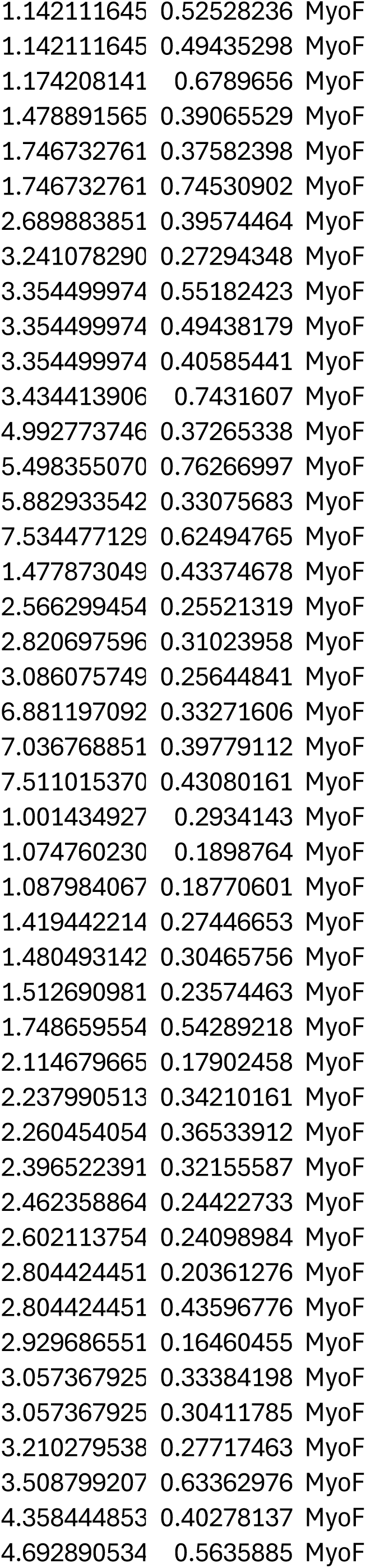
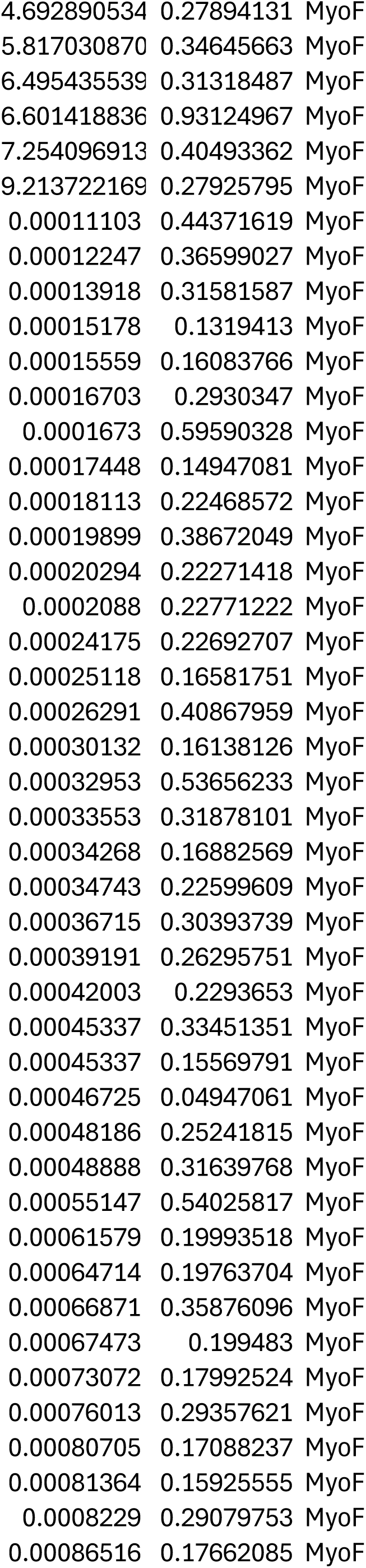
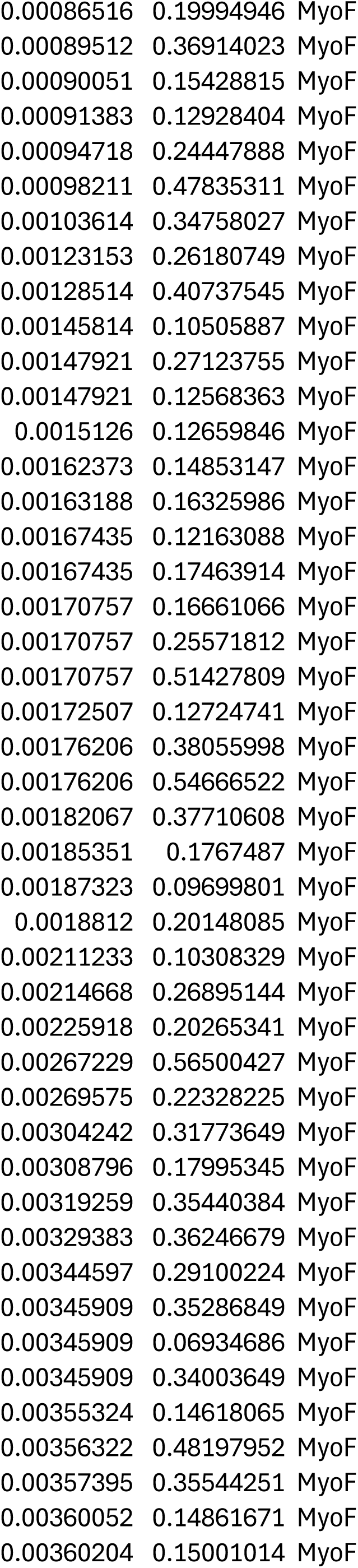
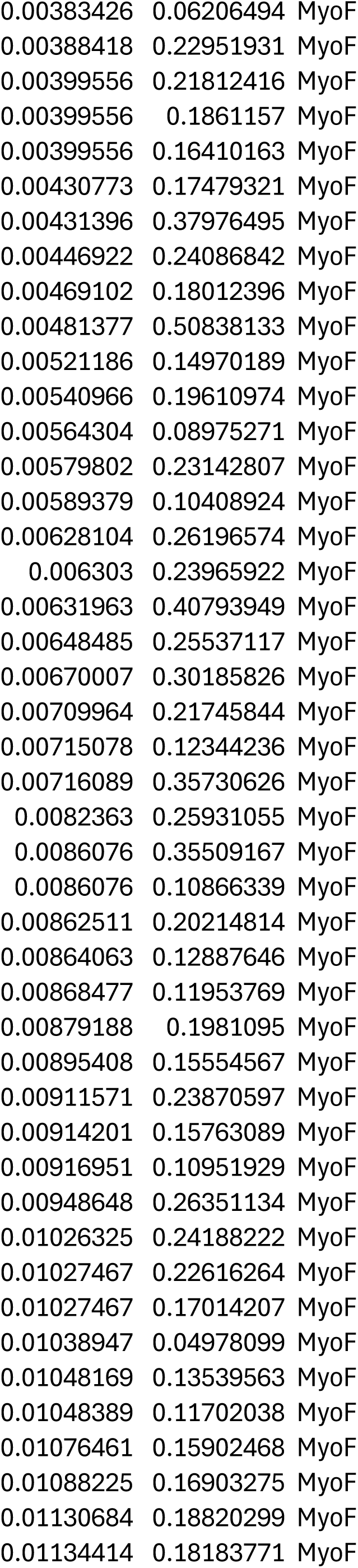
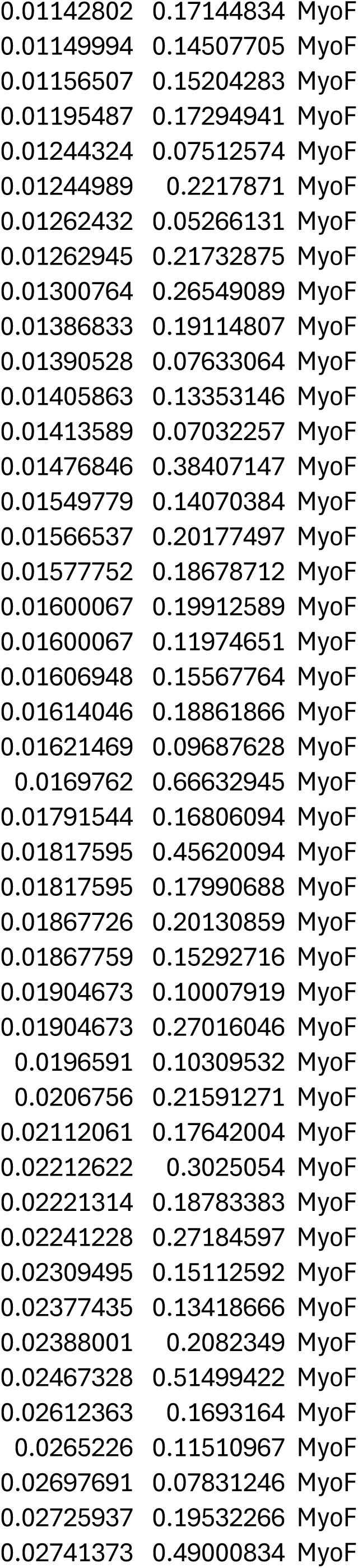
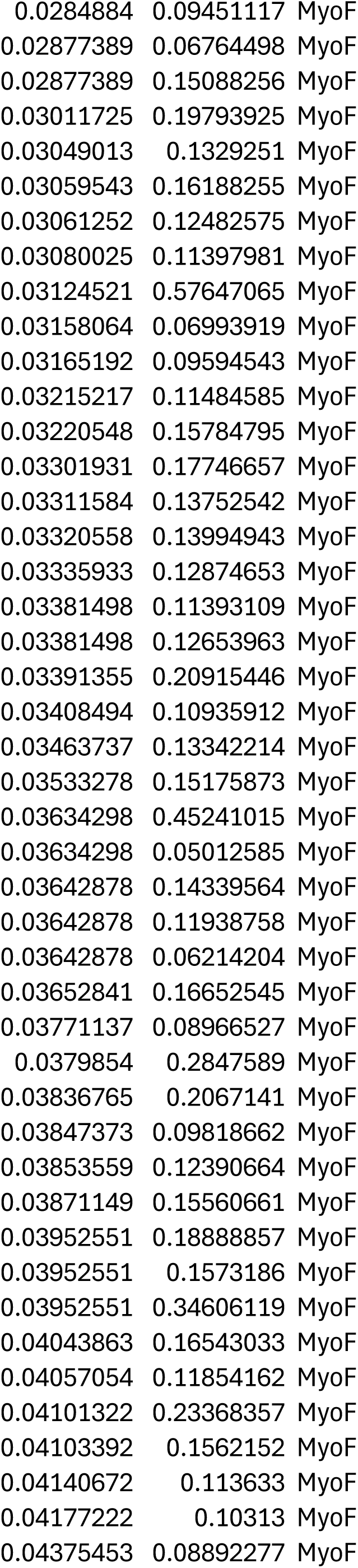
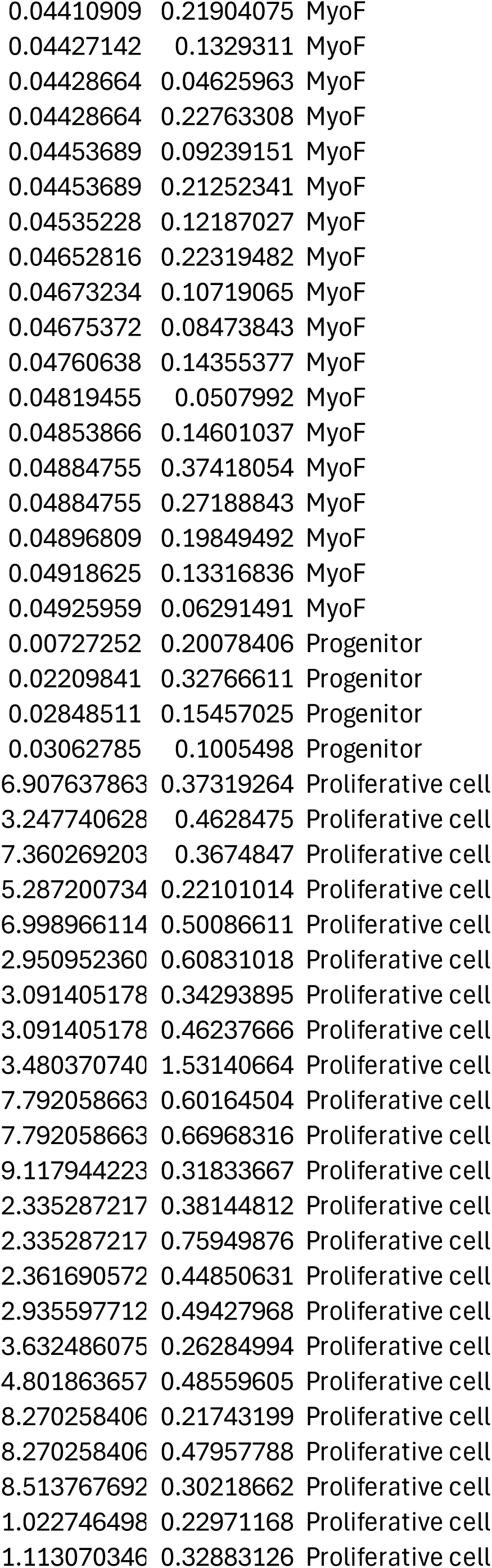
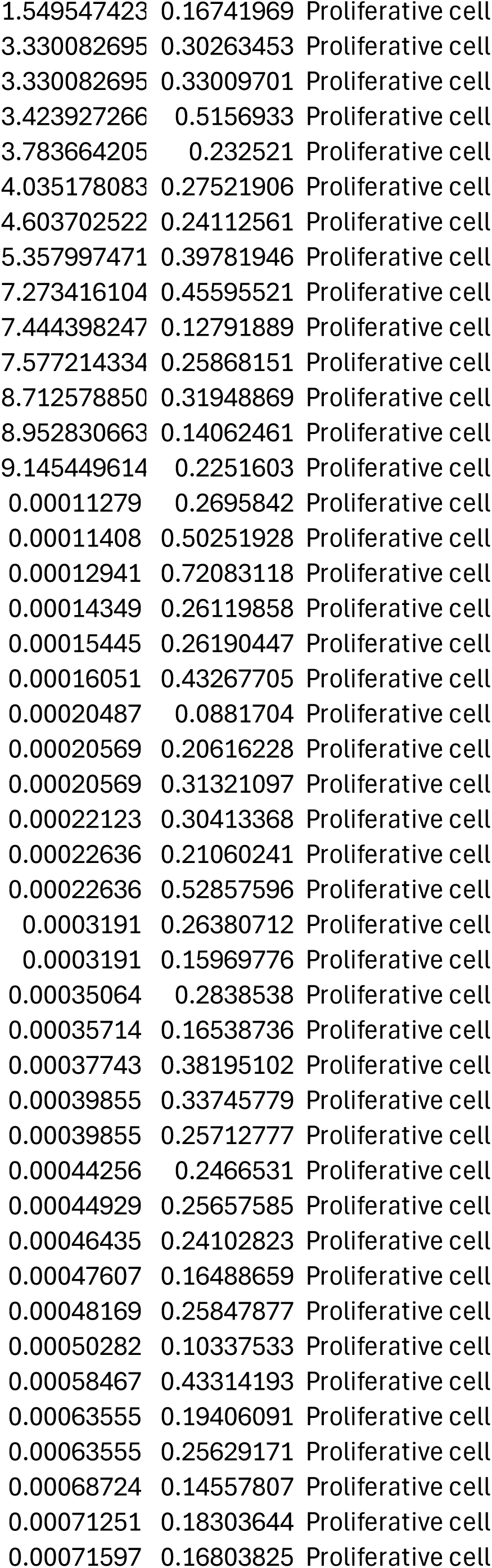
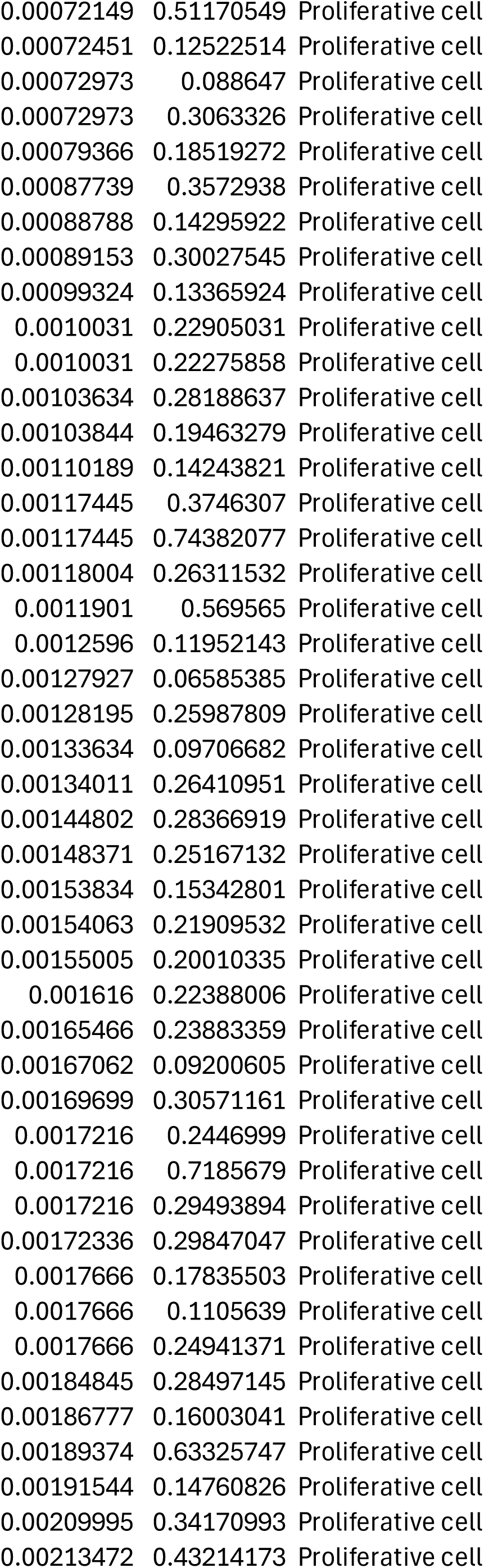
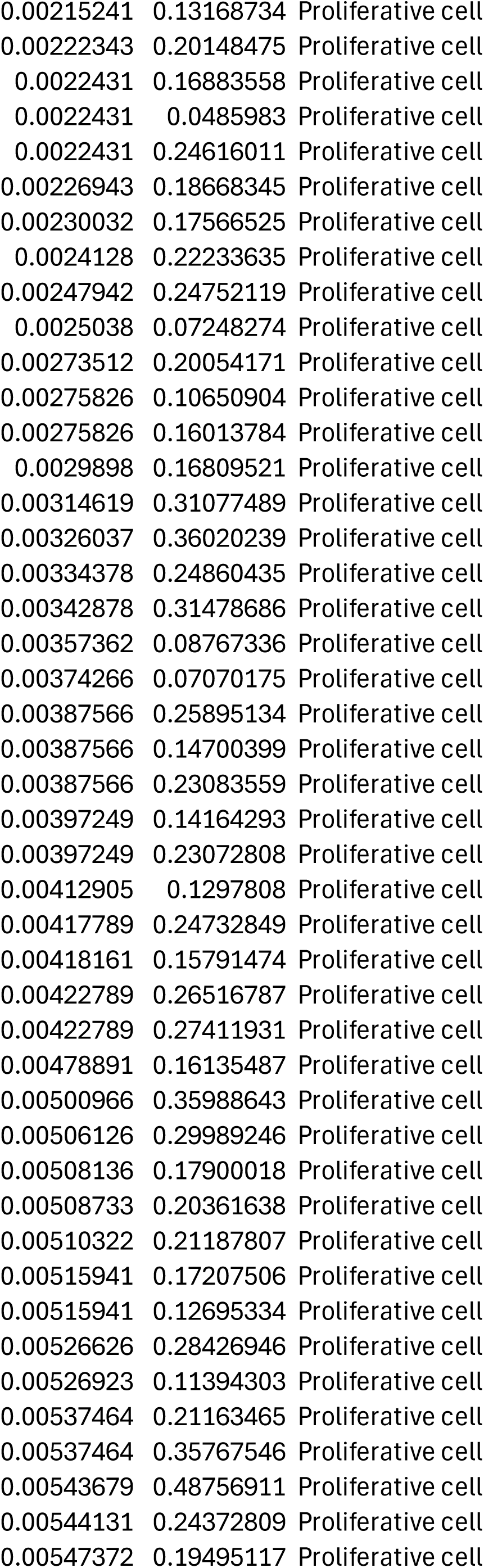
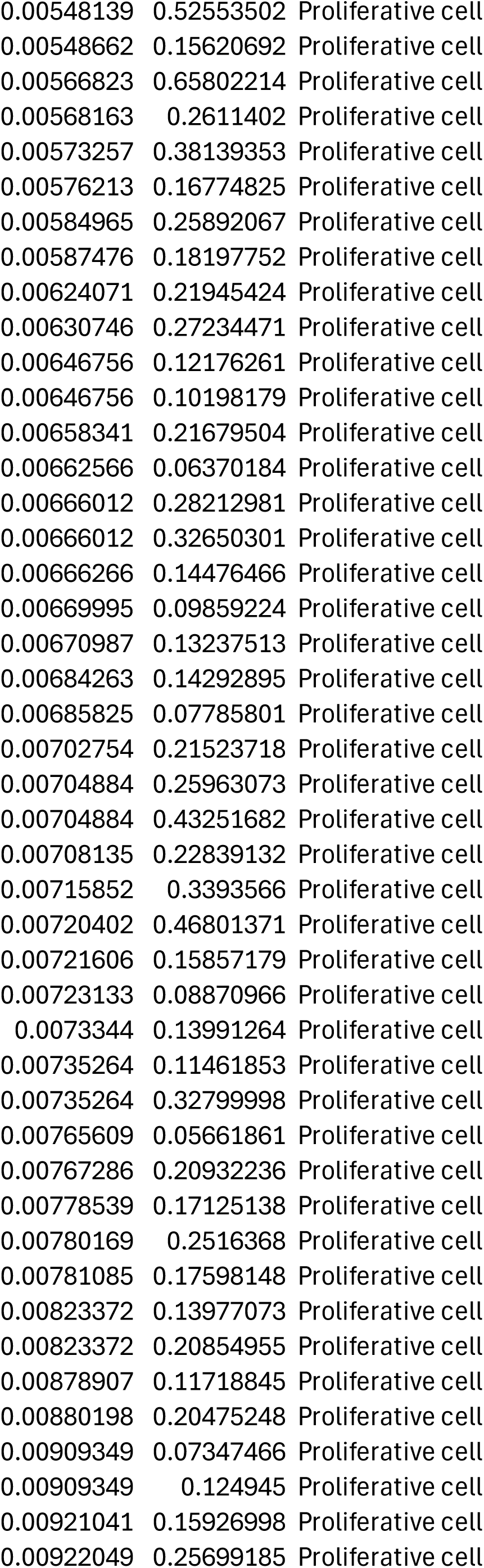
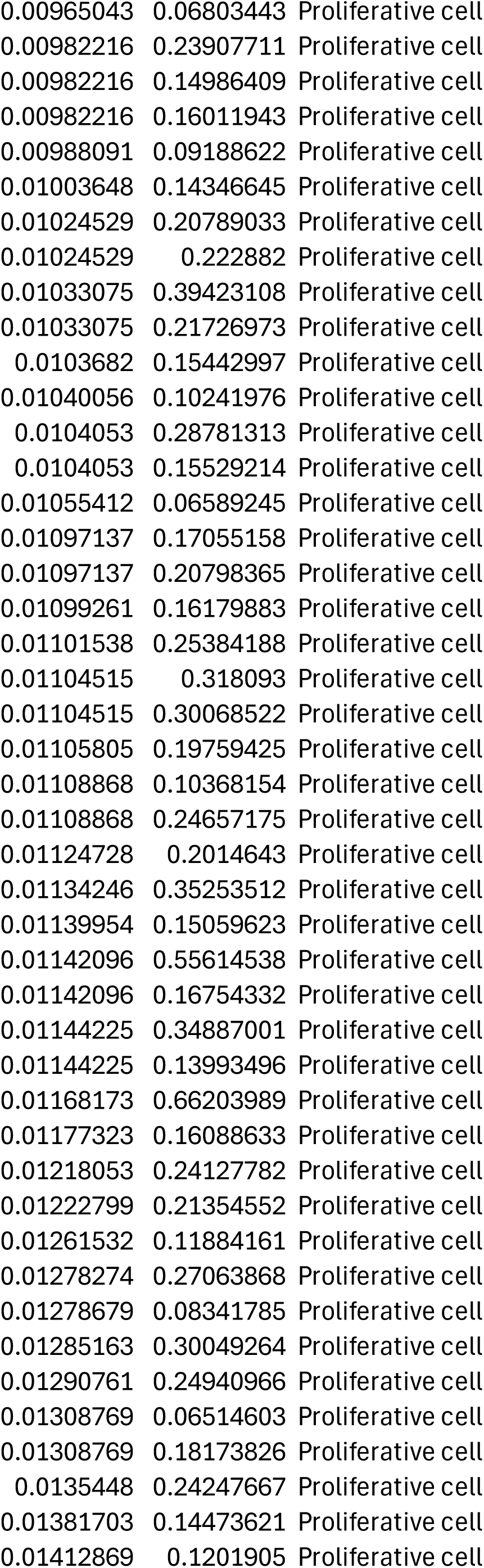
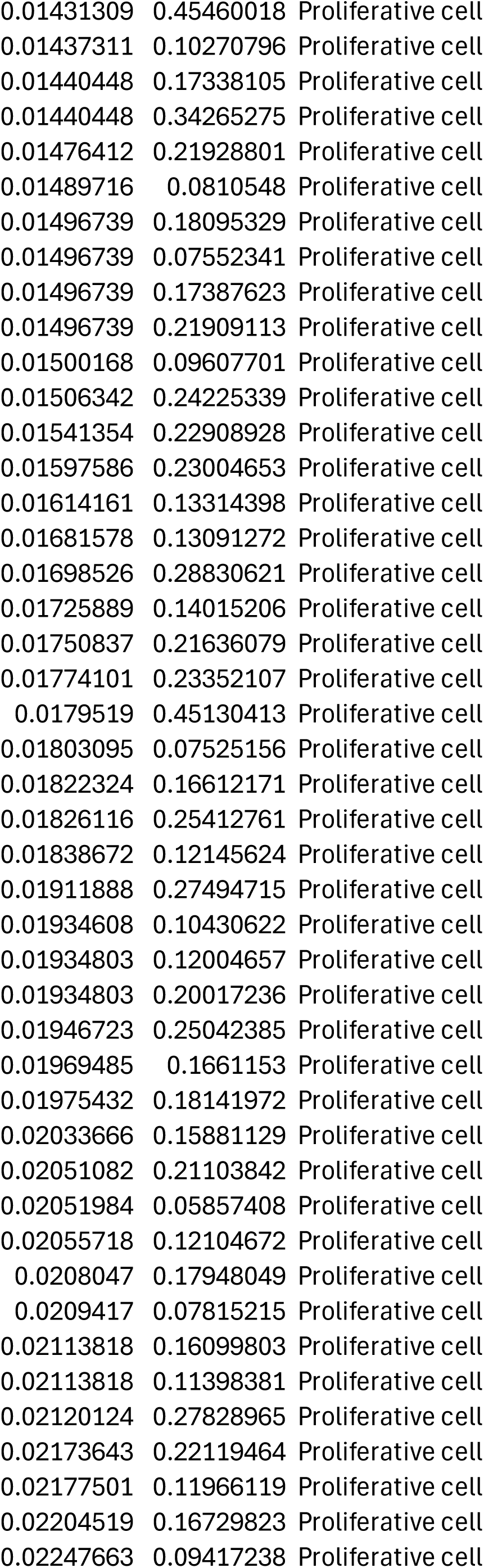
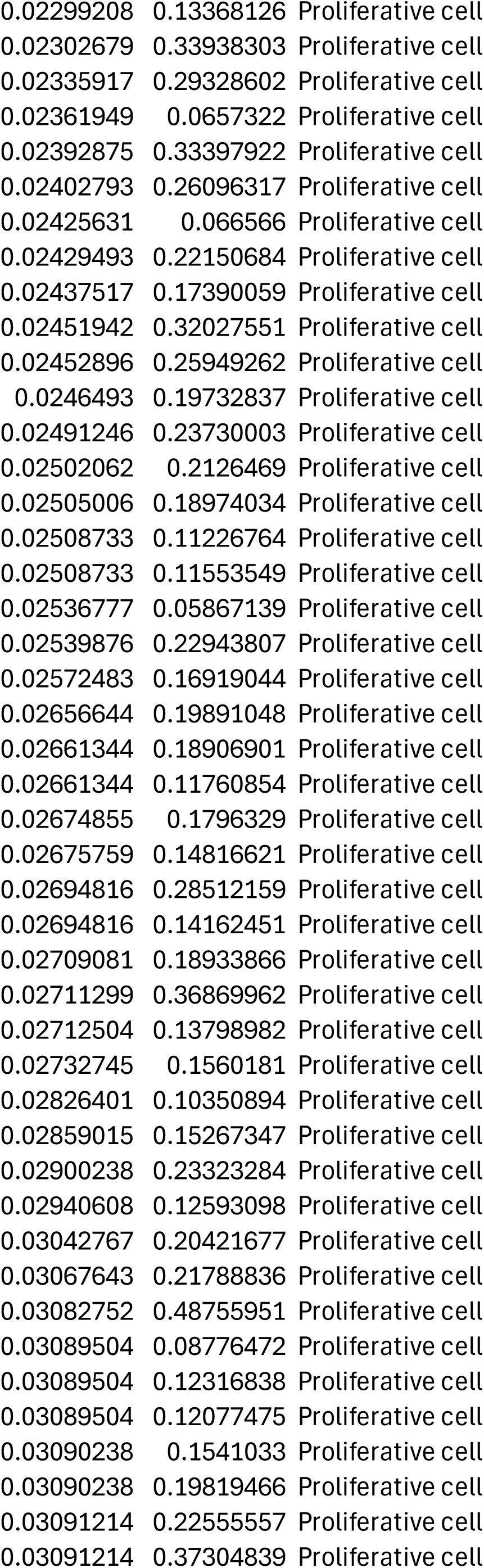
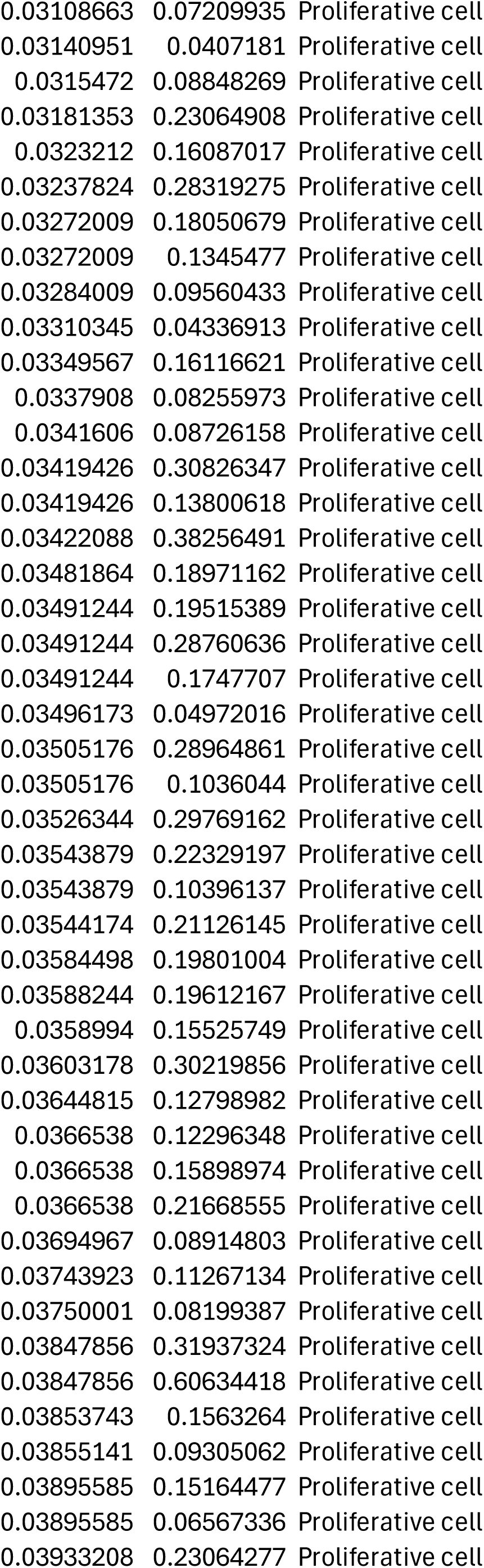
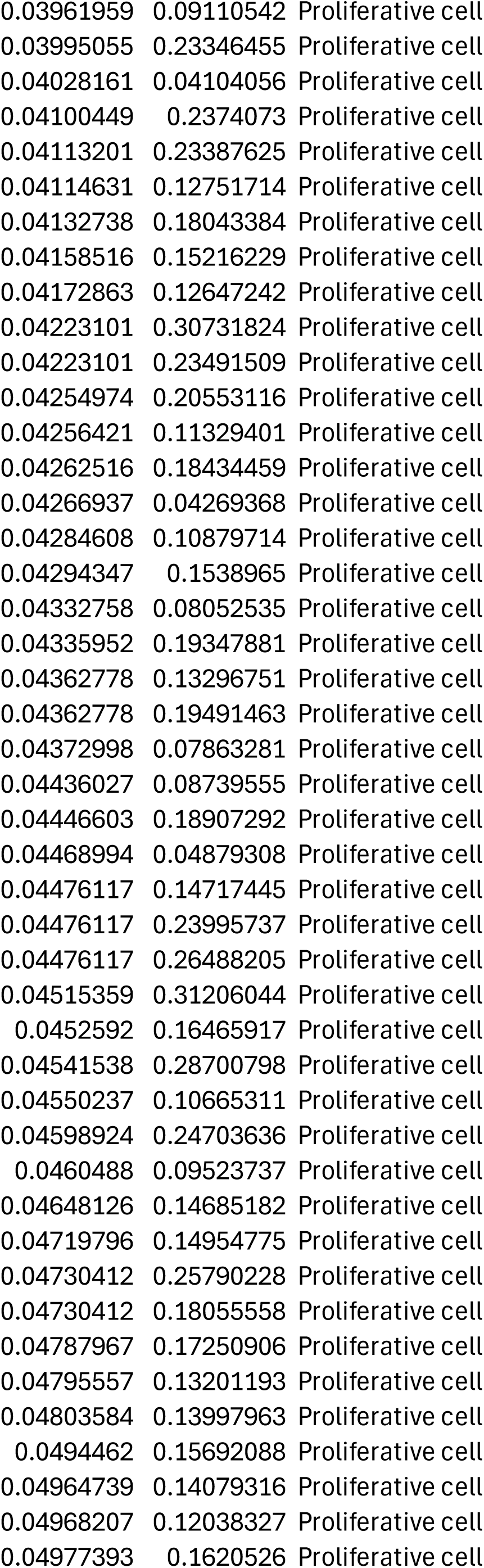
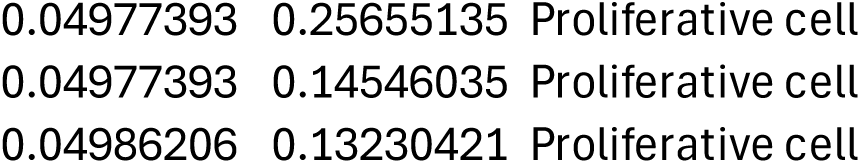

